# Wetland soil history shapes microbial community composition, while hydrologic disturbance alters greenhouse gas fluxes

**DOI:** 10.1101/2020.06.29.178533

**Authors:** Regina B. Bledsoe, Colin G. Finlay, Ariane L. Peralta

**Affiliations:** Department of Biology, East Carolina University, Greenville, NC 27858, USA; NewLeaf Symbiotics, St. Louis, MO 63132, USA

**Keywords:** 16S rRNA, carbon dioxide, denitrification, greenhouse gases, hydrology, metagenomics, methane, nitrous oxide, plant cover, plant-microbe interactions, soil mesocosms, soil redox

## Abstract

While wetlands represent a small fraction (∼5-10%) of the world’s land surface, it is estimated that one-third of wetlands have been lost due to human activities. Wetland habitat loss decreases ecosystem benefits, including improved water quality and climate change mitigation. These microbially mediated functions are dependent on redox conditions, which are altered by soil hydrology and the presence of plants. We tested the overarching hypothesis that while microbial community composition would be resistant to change due to long-term hydrologic history, key functions like greenhouse gas production would remain plastic and responsive to short-term environmental shifts. Using a mesocosm design, we manipulated the duration of hydrologic conditions (i.e., stable dry, stable flooding, and alternating wet/dry) and the presence of plants to induce soil redox changes in wetland soils. We measured soil redox status, used targeted amplicon and shotgun metagenomic sequencing to characterize microbial communities, and measured greenhouse gas production to assess microbial function. The eight-week hydrologic treatment shifted community composition but did not override the stronger effects of long-term hydrologic history. Methane and carbon dioxide fluxes were altered by short-term hydrologic treatment, with methane production favored in the wet treatment and carbon dioxide production favored in the dry treatment. Plant presence versus absence manipulation had little impact on soil microbiome composition or soil greenhouse gas production. The results highlight the resistance of microbial community structure shaped by historical hydrologic regimes, and emphasize that hydrologic conditions exert a stronger influence than plant presence on microbial composition and function. Predicting the outcomes of wetland disturbance and restoration requires an enhanced understanding of community stability and functional plasticity. Our results suggest that wetland hydrologic restoration can establish a stable microbial community that is resistant to environmental shifts, but microbial functions such as greenhouse gas emissions remain responsive to hydrologic disturbances, including flooding and drought.

## Introduction

Wetlands, despite comprising only 5-10% of terrestrial land (Kingsford et al. 2016), have experienced at least a 21% loss globally (Fluet-Chouinard et al. 2023). These ecosystems are vital for water quality improvement and climate change mitigation through carbon sequestration (Mitsch et al. 2015). The conversion of wetlands for urban and agricultural development leads to the loss of these essential functions (Zedler and Kercher 2005). Consequently, there is significant interest in their conservation, restoration, and construction to leverage their ecosystem benefits (Spieles 2022; Patel et al. 2021; Suhani et al. 2020). Key wetland functions include anaerobic microbial processes in saturated soils, which facilitate complete inorganic nitrogen removal via denitrification and suppress aerobic organic matter decomposition (Mcleod et al. 2011; Martínez-Espinosa et al. 2021). However, these anoxic conditions can also support methanogenesis, potentially decreasing long-term carbon storage and climate benefits (Bridgham et al. 2013). The microbial functions delivering these ecosystem services are amenable to management for enhanced wetland benefits, necessitating a deeper understanding of the controls on wetland microbial ecosystem functions (Peralta, Stuart, et al. 2014). This gap in understanding microbial controls on ecosystem functions contributes to the limited success of current wetland restoration efforts, highlighting the need for further research (Cadier et al. 2020).

To produce cellular energy, microorganisms catalyze a series of oxidation-reduction (redox) reactions (Barron 1939; Walz 1979; Seto et al. 2025). Aerobic respiration yields the greatest amount of energy and is expected under dry conditions, but as water fills soil pore spaces, diffusion of oxygen (O_2_) into the soil matrix decreases, causing a shift to a reducing environment (Truu et al. 2009; Mitsch et al. 2013; Marschner 2021). Reducing environments favor anaerobic respiration using terminal electron acceptors such as nitrate (NO_3_ ^−^), manganese, ferric iron, sulfate, and carbon dioxide (CO_2_) (Burgin et al. 2011; Marschner 2021). Therefore, measuring soil redox potential is useful for predicting which biogeochemical processes are likely to be carried out by soil microbes.

To manage microbial functions that support nitrogen removal and retention while reducing methane (CH_4_) production, it is important to understand how changes in hydrology influence microbial communities. Microbial taxa are adapted to specific hydrologic conditions in their environment, and temporal changes in soil moisture (Peralta, Ludmer, et al. 2014; Truu et al. 2009; de Nijs et al. 2019; Ricks and Yannarell 2023). Drought, drainage, and soil re-wetting shift soil redox potentials, microbial community composition, and metabolism, which can influence the relative rates of anaerobic processes such as denitrification and methanogenesis (Kim et al. 2008; Truu et al. 2009; Peralta, Ludmer, et al. 2014; Fairbairn et al. 2023).

If drying and re-wetting events are periodic, a cycle of activity and dormancy can maintain higher microbial diversity than stable environmental conditions (Peralta, Ludmer, et al. 2014). For example, many denitrifiers are facultative anaerobes, capable of rapidly switching between aerobic respiration and denitrification (Tiedje et al. 1984; Sun et al. 2019; Freddi et al. 2023). Methanogens are often dominant in prolonged anoxic conditions, such as flooded wetlands, and are responsible for converting CO_2_ or methyl groups to CH_4_ (Mitsch and Gosslink 2007; Hernández et al. 2017; Ferry 2011). As hydrologic changes occur, oxic-anoxic interfaces support methanotrophs, which transform CH_4_ to CO_2_, and denitrifiers (McDonald and Murrell 1997; Mitsch and Gosslink 2007; Conrad 2009). This dynamic interplay between microbial guilds under fluctuating hydrologic conditions is therefore crucial for key transformations in carbon and nitrogen cycles.

While hydrology is a primary determinant of soil redox potential, plants also modify the soil environment. The soil microenvironments in contact with plant roots are hotspots of microbial activity due to plant-mediated transport of nutrients, O_2_, and carbon belowground (Kuzyakov 2010; Philippot et al. 2013; Chanton 2005). Additionally, vegetation can alter soil redox potentials by facilitating transport of O_2_ into root zones (Chanton 2005; Sundberg et al. 2007) and transporting CH_4_ from root zones into the atmosphere (Carmichael et al. 2014; Hu et al. 2015).

Climate change is projected to increase the frequency of both flooding and drought in the southeastern US, while ongoing land use change continues to reduce wetland and forest cover (Apurv and Cai 2021; IPCC 2023; Easterling et al. 2017; Skeeter et al. 2019; Terando et al. 2014; Griffith et al. 2003; Potapov et al. 2022). These large-scale shifts in hydrology and vegetation represent a major disturbance to wetland soil microbiomes. Understanding and predicting the functional responses of these microbiomes is difficult because their stability is based on a complex network of biotic and abiotic interactions (Griffiths and Philippot 2013; Lennon et al. 2023; Treseder et al. 2012).

Microbial stability is defined by two key properties: resistance (insensitivity to disturbance) and resilience (the rate of recovery post-disturbance) (Shade et al. 2012; Pimm 1984). Following a disturbance, a shift in community function can be driven by two different mechanisms: compositional change (a shift in who is there) or phenotypic plasticity (a change in what the existing microbes are doing) (König et al. 2018; Philippot et al. 2021). Soil microbiome stability is also influenced by abiotic factors, including soil texture, substrate diversity, and nutrient availability (Griffiths and Philippot 2013). Functional and taxonomic characterization of microbiome responses to hydrologic and plant disturbance can help refine predictions of wetland greenhouse gas (GHG) emissions.

This study investigated the effects of hydrologic change duration (i.e., sustained dry, interim dry/wet, sustained wet) and vegetation presence on microbial community structure, soil redox conditions, and GHG production. We hypothesize that hydrologic history will have a greater impact on microbial community composition than eight weeks of hydrologic and plant presence/absence treatments. While we expect community composition to be resistant to change over eight weeks of incubation, we also hypothesize that short-term hydrologic and plant presence/absence treatments will impact GHG production through community phenotypic plasticity combined with a more limited shift in community composition. Predicted functional responses include increased methanogenesis in reducing conditions (wet and no plants) and increased aerobic respiration in oxidizing conditions (dry and plants).

To test these hypotheses, we conducted a mesocosm experiment in which we manipulated hydrology (i.e., dry, interim, and wet) and the presence of plants in soils collected from different hydrologic histories (i.e., dry, wet/dry transition, and saturated wet zones) within a restored coastal plain wetland. We measured soil nutrient concentrations, redox potentials, GHG fluxes, and characterized soil microbial communities using shotgun metagenomic and 16S rRNA gene amplicon sequencing. By imposing acute differences in hydrology and plant cover, we were able to test plant-microbe responses to long-term hydrological regimes versus short-term hydrological disturbances, and the associated net ecosystem greenhouse gas outputs.

## Methods

### Study site

We collected soil samples from the Timberlake Observatory for Wetland Restoration (TOWeR) located in the Albemarle Peninsula in Tyrell County, North Carolina (35°54’22” N 76°09’25” W; Figure 1). The site is part of the Great Dismal Swamp Restoration Bank, which is a 1,700 ha wetland consisting of 420 ha of mature forested wetland, 787 ha of forested wetland, 57.2 ha of drained shrub-scrub, and 440 ha of agricultural fields restored to wetland habitat in 2007 (Ardón et al. 2010; 2013; Morse et al. 2012). A major portion of the restoration effort was to remove drainage ditches and plant 750,000 saplings, including *Taxodium distichum, Nyssa* spp., *Salix nigra, Fraxinus pennsylvanica*, and *Quercus* spp. (Morse et al. 2012). A more recent survey indicates that the dominant plant species at the site is *Juncus effusus* L., along with *Euthamia caroliniana* (L.) Greene ex Porter & Britton, *Solidago fistulosa* Mill., and *Scirpus cyperinus* (L.) Kunth (Hopfensperger et al. 2014). The site is connected to the Albemarle Sound via the Little Alligator River, and the site’s position in the landscape increases the potential for saltwater intrusion to occur (Ardón et al. 2013). Even though there is little elevation change (-1 m to 2 m) across the restored wetland, there is a hydrologic gradient, which is driven by the position of the water table and represents upland dry, saturated wet, and transition dry/wet (“interim”) sites (Hopfensperger et al. 2014).

**Figure 1.**
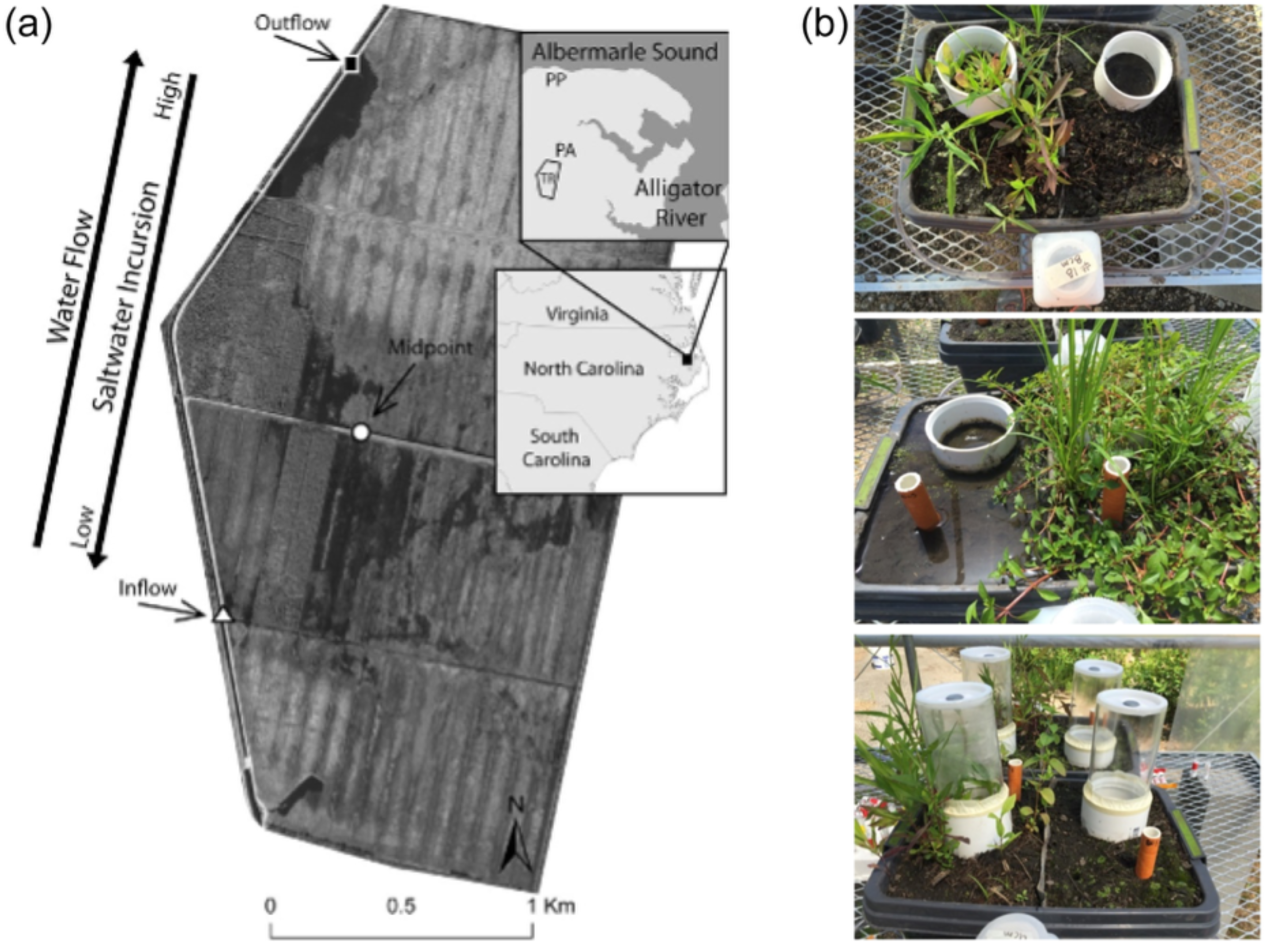
Field site and mesocosm set-up. Wetland mesocosms sourced from the Timberlake Observatory for Wetland Restoration (image modified with permission from Ardón et al., 2013) (a). Aerial view of wetland mesocosm setup, where the left side of the box represents plant treatment, and the right side represents no plant treatment, separated by a stainless mesh divider ((b), top, photo credit: R. Bledsoe). Collars were installed for measuring greenhouse gas concentrations. Also shown is a 1-liter bottle attached to each side of the mesocosm used to maintain water levels. Example of a wet treatment mesocosm with indicator of reduction in soils (IRIS) tubes installed ((b), middle, photo credit: R. Bledsoe). Example of mesocosm with prepared for greenhouse gas sampling ((b), bottom, photo credit: R. Bledsoe). Abbreviations: PP = Palmetto Peartree Preserve, PA = preservation area, TR = Timberlake Observatory for Wetland Restoration.

### Mesocosm experimental design

We used a mesocosm approach to assess the changes in the soil microbial community, redox potentials, and GHG production due to hydrologic changes and vegetation. We set up a mesocosm experiment using soil blocks collected from three sites within the TOWeR. Soils were collected from the restored portion of the site (12 years post-restoration), which had been used as agricultural fields for several decades before restoration (Ardón et al. 2010). We collected soils that represented three different hydrologic zones on the inflow side of the site at the following coordinates: 35.895917° N 76.165972° W (dry), 35.895778° N 76.165861° W (interim), and 35.895639° N 76.165556° W (wet). Six intact soil blocks (25 cm × 25 cm × 20 cm deep) were cut out from each of the three sites using landscaping knives and shovels. Soil hydrologic history was determined by water table levels at the time of collection (dry = 20 cm, interim = 10 cm, and wet = 0 cm below surface level). Soil blocks were contained in dark plastic containers of the same dimensions as the soil blocks (Figure 1).

To manipulate the presence of vegetation, the container was divided in half and separated by a root screen (20 µm stainless steel mesh). On one side, plants were allowed to grow during the experiment, while the ‘no plant’ side was maintained by careful removal of above- and belowground plant biomass. To manipulate hydrology within the mesocosm, vinyl tubing was inserted 3 cm from the bottom of the mesocosm container on each side and connected to a 1 L water bottle (Figure 1). Water levels inside the mesocosm were maintained by filling the 1 L bottle to the desired height using rainwater collected from a cistern. Finally, 10 cm diameter schedule 40 polyvinyl chloride (PVC) collars were inserted 5 cm below the surface as a sampling base for GHG chambers (described in subsection *GHG fluxes*). The mesocosm experiment was housed at East Carolina University (ECU) West Research Campus (Greenville, NC) under a covered hoop house with a shade cloth, which prevented precipitation and allowed 60% light penetration. Soils were collected on April 1, 2016, and prepared on May 29, 2016. Experimental mesocosm sampling commenced on June 13, 2016, and concluded on August 11, 2016. Temperatures ranged from 22.7 °C to 30.8 °C during the experiment. Because the hoop house was in a wetland approximately 150 km from the field site, within the same physiographic region, the temperature and relative humidity in the hoop house would have closely resembled field conditions. Blocking precipitation was essential for the controlled manipulation of hydrologic conditions.

Two weeks after installing PVC collars and setting up the plant treatment, we started hydrologic treatments. Hydrologic manipulation occurred over eight weeks to allow for multiple dry/wet transitions for the interim treatment. Mesocosms exposed to wet conditions were flooded by overhead watering and maintained using water reservoirs filled to maximum container height (18 cm). Mesocosms exposed to dry conditions were maintained using water reservoirs filled to a depth of 5 cm. The dry/wet (interim) treatments fluctuated between flooded and dry conditions, as described above, every two weeks, starting with a wet treatment and ending with a dry treatment.

All mesocosm plants originated from the standing plant community and seed bank (described in Hopfensperger et al. 2014). We did not monitor plant community composition, but we did observe general differences in mesocosm plant communities based on hydrologic treatment. Continuously flooded mesocosms were often dominated by aquatic and semi-aquatic plants like *Ludwigia palustris*, while interim and dry mesocosms contained a higher proportion of sedges and *J. effusus*.

### Measuring integrated soil redox status

We measured soil redox status at the beginning and end of the experiment using Indicator of Reduction in Soils (IRIS) tubes (InMass Technologies, Figure 2, Appendix S1: Figure S1). The IRIS tubes are constructed from 1.5 cm diameter schedule 40 PVC pipe coated in iron oxide (Fe(III)) paint (Rabenhorst 2008). In oxidizing conditions, Fe(III) appears as an orange-red paint; but when exposed to reducing conditions, Fe(III) is reduced to Fe(II), which dissolves in solution, leaving a white clearing on the tube (Jenkinson and Franzmeier 2006; Rabenhorst 2008). At the beginning of the experiment, two IRIS tubes (12 cm depth) were installed in each mesocosm: one on the plant side and one on the bare soil side. The IRIS tubes were incubated in mesocosms for two weeks before removal and analysis. After we removed the IRIS tube, a non-coated PVC pipe was used to fill the hole. Two weeks before the end of the experiment, the non-coated PVC pipe was removed and replaced with a new IRIS tube to measure soil redox status at the end of the experiment.

**Figure 2.**
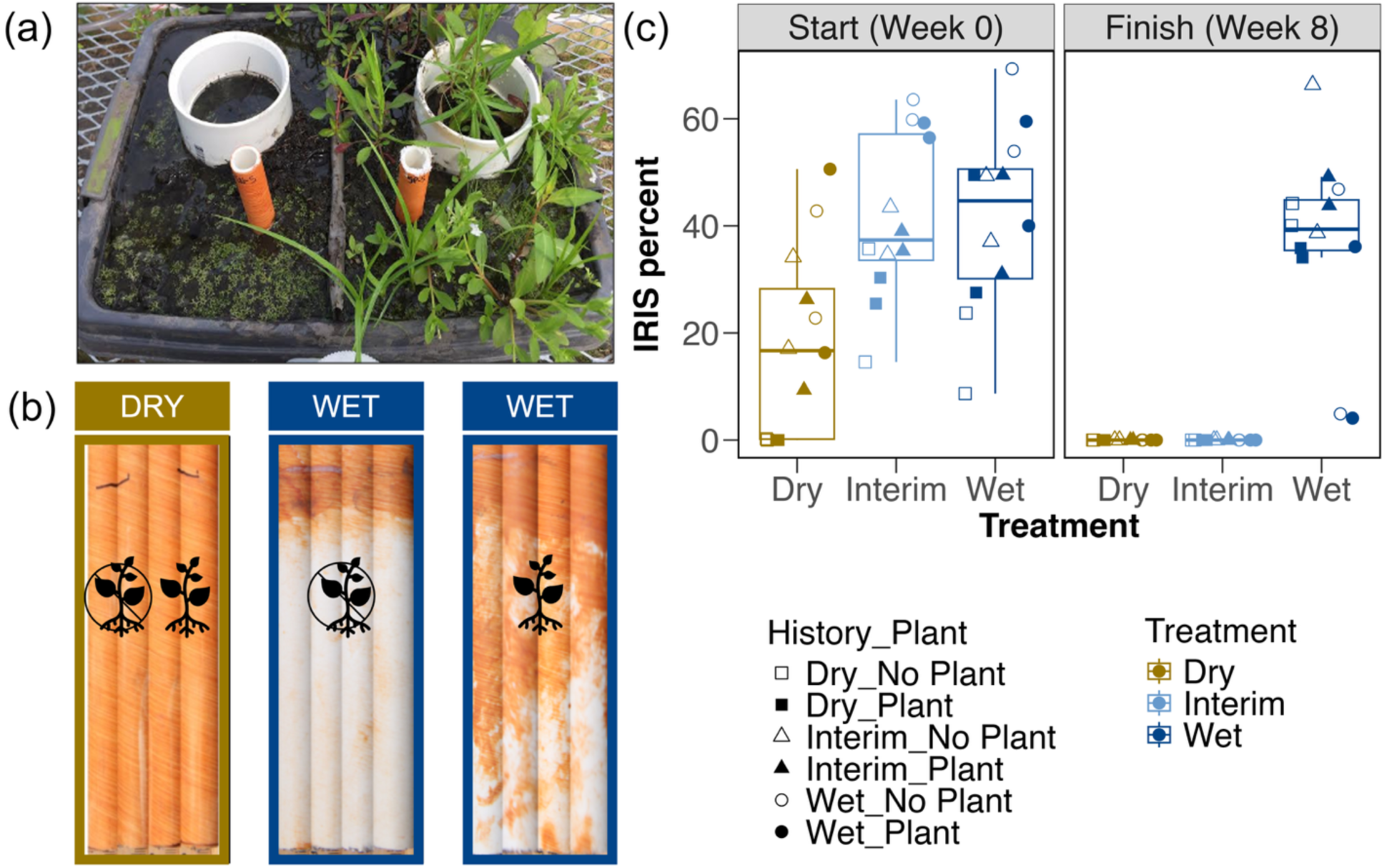
Indicator of reduction in soils (IRIS) tubes mesocosm image (photo credit: R. Bledsoe) (a), IRIS tube example images (photo credit: R. Bledsoe) (b), and boxplots summarizing redox status at the beginning and end of the experiment (c). Percent paint removed from IRIS tubes at the start of the experiment (week 0) and the end of the experiment (week 8). Example IRIS tubes used for ImageJ analysis post-incubation (b) with plant icons crossed out represent IRIS tubes from mesocosms with plants removed, and plant icons represent IRIS tubes from mesocosms with plants present. Box plots summarize percent paint removed from IRIS tubes at start (week 0) and finish (week 8) of experimental incubation. Color represents hydrologic treatment: brown indicates dry, light blue indicates interim, and dark blue indicates wet. Individual samples are plotted as points and colored according to their hydrologic treatment. Point shape represents hydrologic history (square = dry history, triangle = interim history, circle = wet history). Closed points represent mesocosms with plants, while open points represent mesocosms without plants. Boxplots summarize median, first and third quartiles, and two whiskers extending |≤| 1.5 * interquartile range.

We quantified the surface area of Fe(III) paint removed from IRIS tubes. First, we imaged the entire tube by taking four pictures and then stitched the photos into a composite using *GIMP2* (v2.8.14) (Kimball et al. 2014). Next, we identified areas of artificial paint removal (i.e., scratches from installing or removing tubes) and manually filled these pixels. Then, using *ImageJ* (v1.48) (Schneider et al. 2012), we converted all colored pixels to black. We compared the number of white pixels to the total pixels to determine the percentage of paint removed. Interpretation of redox status is based on the percent paint removed from a 10 cm section of tubing and summarized as follows: 0% not reducing, 1-5% probably not reducing, 5-10% possibly reducing, 10-25% and >25% definitely reducing (Rabenhorst 2008). For regression analyses, the percent of paint removed was treated as a continuous variable.

### Soil physicochemical characteristics

Soil properties were determined from soils collected during the destructive sampling of mesocosms at the end of the experiment. We collected six soil cores (3 cm diameter, 10 cm depth) from each side of the box (i.e., plant and no plant) and combined cores collected from one side into a composite sample (i.e., one plant composite and one no plant composite), passed soils through a 4 mm sieve, and homogenized samples before subsampling for soil analyses. For each sample, we measured gravimetric soil moisture by drying 20-30 g of field-moist soil at 105 °C for at least 24 hours. Approximately 5 g of field-moist soil was extracted with 45 ml of 2 M KCl, and extractable ammonium (NH_4_ ^+^) and NO_3_ ^−^ ions were colorimetrically measured using a SmartChem 200 auto analyzer (Unity Scientific, Milford, Massachusetts, USA) at the ECU Environmental Research Laboratory. To determine total carbon and total nitrogen (TC, TN), a subsample of air-dried soil was finely ground and sieved through a 500 µm mesh and analyzed using an elemental analyzer (2400 CHNS Analyzer; Perkin Elmer, Waltham, Massachusetts, USA) at the Environmental and Agricultural Testing Service Laboratory, Department of Crop and Soil Sciences, North Carolina State University. A second subsample of air-dried soil was sent to Waters Agricultural Laboratories, Inc. (Warsaw, NC) and analyzed for pH, phosphorus, potassium, magnesium, sulfur, manganese, iron, and humic matter, using standard Mehlich III methods (Mehlich 1984; Mylavarapu et al. 2014).

### Greenhouse gas (GHG) fluxes

We examined the effects of hydrology and vegetation on GHG production. We measured GHG fluxes on June 13, 2016, two weeks after hydrologic treatments were established, and then every two weeks until August 11, 2016, for a total of five sampling events. The GHG collection chambers were 20 cm in height with a diameter of 8.25 cm and constructed from clear acrylic tubing. GHG fluxes were measured using transparent chambers, thus representing net fluxes of heterotrophic and autotrophic plant and microbial activity. Chambers were sealed on one end with silicon and a pipe cap with a 33 mm septum installed as a gas sampling port. At the time of sampling, chambers were placed on top of preinstalled PVC collars, and the seals were taped to prevent diffusion from the seam (Hoffmann et al. 2018). A gas sample was collected from the chambers at four time points: 30, 60, 120, and 180 minutes after the chambers were attached. Initial ambient (T_0_) samples were not obtained before chamber closure. Instead, they were collected from the mesocosm incubation field site at the ECU West Research Campus as part of a 16-month effort to sample coastal plain wetland GHGs, spanning from August 2021 to December 2022. Monthly average ambient GHG concentrations were calculated and used to represent the T_0_ concentration for mesocosm flux measurements within each corresponding month. To collect gas samples, we used a needle attached to a 20 mL syringe, mixed headspace gas three times by pulling and depressing the plunger, and then collected a 20 mL sample. Gas samples were equally distributed between two 3 mL glass Exetainer vials (Lampeter, Wales, UK) fitted with screw top and dual-layer septa. Samples were stored upside down at room temperature in a dark location and were analyzed within 96 hours of collection.

We analyzed GHG concentrations using a Shimadzu gas chromatograph (GC-2014) fitted with an electron capture detector to detect N_2_O and a flame ionization detector with a methanizer to measure CH_4_ and CO_2_. Calibration standards are listed in Appendix S1: Table S1. If a sample measured above the calibration curve, we diluted the sample, remeasured the concentration, and accounted for the dilution factor when calculating the final concentrations. After the first two time points, calibration curves were adjusted to accommodate larger GHG concentrations (Appendix S1: Table S1). Since all unknown concentrations were analyzed within the bounds of their calibration curve, results were unaffected by adjustments to the calibration curves.

We calculated GHG fluxes using the *gasfluxes* package implemented in the R Statistical Environment and RStudio (RStudio 2024.12.0+467, Rv4.4.2) (Fuss and Hueppi 2024; R Core Team 2024). The best model fit among HMR, robust linear, and linear was selected for each flux (Appendix S2).

### Bacterial and archaeal community analyses

We assessed the soil microbial community composition at the beginning (‘start’, based on soils collected from the IRIS tube installation) and end of the experiment (‘final’, based on soils collected during destructive sampling). We collected ‘final’ soil samples using a standard soil probe (3 cm diameter, 10 cm depth) from the same location that we collected GHG samples. We extracted genomic DNA from the soils using the Qiagen DNeasy PowerSoil Kit and diluted the DNA to 20 ng/µL. This genomic DNA was used as template in PCR reactions with barcoded primers (bacterial/archaeal 515FB/806R primer set) originally developed by the Earth Microbiome Project to target the V4 region of the bacterial 16S subunit of the ribosomal RNA gene (Caporaso et al. 2012; Apprill et al. 2015; Parada et al. 2016). PCR reactions and sequencing library preparation are described in Appendix S1: Supplemental Methods. We sequenced the pooled libraries using the Illumina MiSeq platform using paired-end reads (Illumina Reagent Kit v2, 500 reaction kit) at the Indiana University Center for Genomics and Bioinformatics Sequencing Facility.

Sequences were processed using the *mothur* (v1.42.0) MiSeq pipeline (Kozich et al. 2013; Schloss et al. 2009). We assembled contigs from the paired-end reads, quality trimmed using a moving average quality score (minimum quality score 35), aligned sequences to the SILVA rRNA database (v132) (Quast et al. 2013), and removed chimeric sequences using the VSEARCH algorithm (Rognes et al. 2016). We created operational taxonomic units (OTUs) by first splitting sequences based on taxonomic class and then binning them into OTUs based on 97% sequence similarity. Since amplicon sequencing was conducted using short reads (250 bp) and not assembled genomes, we used an OTU-based 3% distance threshold to avoid splitting the same bacterial genome into distinct clusters (using amplicon sequence variant (ASV) of a single base difference). An OTU-based approach also limits over-inflated numbers of ASVs introduced by PCR bias and sequencing errors (Schloss 2021). In addition, employing an OTU- or ASV-based approach is known to result in similar broadscale community composition patterns (Glassman and Martiny 2018).

### Shotgun metagenomic analyses

Based on amplicon sequencing, we identified a subset of samples representing the most distinct microbial communities for shotgun metagenomic sequencing. A description of sample processing and metagenomic sequencing can be found in Peralta et al. (2020). Briefly, we chose a set of baseline and post-flooding/drying treatment samples. We sourced mesocosm baseline conditions from relatively dry (water level at about -20 cm, n = 4) and relatively wet (water level at about 0 cm, n = 4) field locations. At the end of the eight-week experiment, we selected a subset of samples from the hydrologic treatments (prolonged drying or wetting only) and plant treatments (presence or absence of vegetation) (n = 16). After genomic DNA extraction using the Qiagen DNeasy PowerMax soil kit, samples were sent to the U.S. Department of Energy (DOE) Joint Genome Institute (JGI) for sequencing and analysis (GOLD study ID Gs0142547 and NCBI BioProject accession number PRJNA641216). Plate-based DNA library preparation and subsequent Illumina sequencing, according to published protocols (Peralta et al. 2020), resulted in 583.2 Gbp of raw sequence data. Raw data were processed using the DOE JGI Metagenome Annotation Pipeline IMG/M v.5.0.9 (Chen et al. 2017; Huntemann et al. 2016; Markowitz et al. 2014).

From the metagenomes, we curated functional gene sets using IMG/M’s integrated Kyoto Encyclopedia of Genes and Genomes (KEGG) module list. For statistical analyses, we focused on the following four functional gene sets associated with biogenic greenhouse gas fluxes: (1) Denitrification (KEGG Module M00529), (2) Methanogen (M00617), (3) Central Carbohydrate Metabolism (M00001-M00011, M00307-M00309, M00580, and M00633), and (4) Prokaryotic Cytochrome C Oxidase (CcO) (M00155).

### Statistical Analyses

All statistical analyses were performed in the R statistical environment (RStudio 2024.12.0+467, Rv4.4.2) (R Core Team 2024). Before multivariate statistical analyses, we normalized sample-to-sample variation in sequence depth by taking the relative abundance of each OTU. OTU data were then rarefied using *vegan::rrarefy* (Oksanen 2015). Likewise, functional gene counts were converted to relative abundance by dividing the count of each functional gene by the total number of functional gene counts in each respective functional gene category. We examined beta diversity by visualizing bacterial community responses to hydrologic history (field conditions), hydrologic treatment (contemporary dry/wet treatments), and plant presence/absence using principal coordinates analysis (PCoA) of bacterial community composition based on Bray-Curtis dissimilarity. We also used PCoA to visualize the Bray-Curtis dissimilarity of functional gene categories according to hydrologic history, hydrologic treatment, and plant presence/absence. We used permutational multivariate analysis of variance (PERMANOVA) to determine differences between bacterial communities among hydrologic history, hydrologic treatment, and plant presence. Hypothesis testing using PERMANOVA was performed using the *vegan::adonis* function (Oksanen 2015). Unique taxa representing each hydrologic history and hydrologic treatment were determined by Dufrene-Legendre indicator species analysis using the *labdsv::indval* function (Roberts 2023). Finally, soil properties were compared against bacterial community and functional patterns (based on Bray-Curtis dissimilarity) using the *vegan::envfit* function (Oksanen 2015). Soil properties with *p* ≤ 0.05 were represented on the PCoA plot of 16S rRNA community composition as vectors scaled by their correlation to microbial community patterns. The GHG fluxes and soil properties with *p* ≤ 0.1 were plotted as vectors, scaled by their correlation, on PCoA plots of functional gene composition.

We constructed linear mixed effects models with sampling plot (mesocosm) and sampling date as random effects to determine the importance of the fixed effects of hydrologic history (field conditions), hydrologic treatment (contemporary dry/wet treatments), and plant presence on GHG fluxes using the *lme4* R package (Bates et al. 2015). Then, we used Akaike’s Information Corrected Criterion (AICc) model comparisons to determine the simplest combination of fixed effects needed to explain the most variance in GHG fluxes (Hurvich and Tsai 1993; Gorsky et al. 2019). We did not use the interaction between the fixed effects hydrologic history and hydrologic treatment because combinations would have resulted in a sample size of *n* = 2 for each group. Lastly, to determine the proportion of variance explained by fixed effects (marginal) and the complete model (conditional), we used the *MuMIn* R package (Gorsky et al. 2019; Bartoń 2025).

While linear mixed effects models are robust to departures from normality (Schielzeth et al. 2020), traditional ANOVAs and t-tests are less so (Blanca et al. 2017; Havlicek and Peterson 1974). To accommodate extreme outliers and lack of normality, we used nonparametric statistics to test for differences in GHG fluxes among hydrologic treatments and plant presence/absence. We used Wilcoxon Rank Sum tests to assess differences in GHG fluxes between plant and no plant groups, and Kruskal-Wallis tests to compare GHG fluxes among the three hydrologic treatments (dry, interim, and wet). To account for repeated measures, the Wilcoxon Rank Sum and Kruskal-Wallis tests were grouped by sampling date. If the Kruskal-Wallis test for a sampling date revealed differences among treatment groups (*p* ≤ 0.05), we then performed pairwise comparisons using the Dwass, Steel, Critchlow, and Fligner (SDCFlig) test with a Monte Carlo distribution (Schneider et al. 2023).

To assess microbial community structure and function relationships, we examined the relationship between GHG fluxes and microbial community/functional Bray-Curtis dissimilarity matrices using distance-based partial least squares regression (DBPLSR) in the *dbstats* R package (Boj et al. 2024). Finally, we used the *vegan::mantel* function (Oksanen 2015) to examine the relationship between the soil properties (redox, percent moisture, pH, NH_4_ ^+^, total soil C, total soil N, phosphorus ppm, potassium ppm, magnesium ppm, sulfur ppm, iron ppm, manganese ppm, and percent humic matter), GHG fluxes, and microbial community composition.

## Data Availability

All code and data used in this study are in a public GitHub repository (https://github.com/PeraltaLab/WetlandMesocosm_GHG_Timberlake) with a Zenodo DOI (https://doi.org/10.5281/zenodo.15528281). Raw amplicon sequence files can be found at NCBI SRA <u>BioProject ID PRJNA636184</u>, and metagenome sequence files can be found at NCBI SRA <u>BioProject ID PRJNA641216</u>.

## Results

Soil mesocosms sampled from three distinct hydrologic conditions (dry, interim, wet), were subjected to eight weeks of hydrologic and plant presence/absence treatments (Figure 1). The three levels of hydrologic treatments were: (1) flooded for eight weeks, (2) dry for eight weeks, and (3) interim, i.e., two weeks flooded alternating with two weeks dry for a total of eight weeks. Each box was separated by a stainless mesh divider, with plants allowed to grow on one side and plants removed on the other side (Figure 1).

### Characterization of soil physical-chemical properties

There was little variability in soil properties across main effects (hydrologic history, hydrologic treatment, plant presence), except for notable differences in soil moisture between plant treatments within dry and wet hydrologic treatments (Table 1). In addition, NO ^−^N concentrations were below the detection limits in samples with plants, but measurable in samples without plants (Table 1). Soil redox status as measured by IRIS tubes revealed that soils experiencing dry and interim treatments were not reducing (0% iron oxide paint removed), while soils in wet treatments were reducing (Figure 2). In wet treatments, localized oxidizing conditions were maintained on portions of IRIS tubes in contact with roots (Figure 2, Appendix S1: Figure S1).

**Table 1.** Soil chemical and physical properties measured after eight weeks of hydrologic manipulation.

| Treatment<br>Plant | Dry |  | Interim |  | Wet |  |
| --- | --- | --- | --- | --- | --- | --- |
|  | No Plant | Plant | No Plant | Plant | No Plant | Plant |
| Moisture (%) | 39 ± 0.31 | 14 ± 0.03 | 19 ± 0.16 | 13 ± 0.02 | 29 ± 0.23 | 62 ± 0.07 |
| pH | 5.31 ± 0.11 | 5.31 ± 0.09 | 5.4 ± 0.16 | 5.35 ± 0.1 | 5.55 ± 0.15 | 5.41 ± 0.17 |
| NH<sub>4</sub><sup>+</sup> mg L<sup>-1</sup> | 0.39 ± 0.07 | 0.27 ± 0.04 | 0.26 ± 0.1 | 0.19 ± 0.02 | 0.77 ± 0.76 | 0.39 ± 0.42 |
| NO<sub>3</sub><sup>-</sup> mg L<sup>-1</sup> | 0.08 ± 0.07 | MDL | 0.25 ± 0.38 | MDL | 0.01 ± 0.0005 | MDL |
| Total C (%) | 4.63 ± 0.78 | 4.63 ± 0.81 | 4.48 ± 0.32 | 4.37 ± 0.52 | 4.91 ± 0.92 | 4.68 ± 0.74 |
| Total N (%) | 0.22 ± 0.03 | 0.22 ± 0.03 | 0.22 ± 0.01 | 0.21 ± 0.03 | 0.24 ± 0.04 | 0.23 ± 0.03 |
| P ppm | 21.00 ± 4.14 | 20.41 ± 4.4 | 23.58 ± 8.87 | 22.83 ± 8.37 | 23.16 ± 5.98 | 23 ± 5.79 |
| K ppm | 33.66 ± 5.97 | 30.25 ± 10.31 | 39.5 ± 11.44 | 26.75 ± 3.04 | 46.16 ± 16.02 | 34.25 ± 10.78 |
| Mg ppm | 77.58 ± 6.46 | 74.66 ± 11.43 | 86 ± 22.33 | 78.66 ± 16.47 | 89.25 ± 16.85 | 79.41 ± 18.8 |
| S ppm | 15.75 ± 1.94 | 15.33 ± 2.29 | 13.25 ± 1.25 | 12.16 ± 0.40 | 11.33 ± 1.21 | 11.5 ± 1.00 |
| Fe ppm | 252.66 ± 14.45 | 243.66 ± 17.06 | 275.33 ± 27.07 | 251.16 ± 37.6 | 306.83 ± 22.52 | 280.66 ± 40.71 |
| Mn ppm | 3.91 ± 1.15 | 3.41 ± 0.97 | 4.33 ± 1.16 | 4.58 ± 1.31 | 5.83 ± 1.32 | 5.58 ± 2.05 |
| Humic matter (%) | 2.35 ± 0.19 | 2.17 ± 0.14 | 2.04 ± 0.29 | 2.28 ± 0.22 | 1.98 ± 0.422 | 1.96 ± 0.43 |

Notes: Average (mean ± standard deviation) by hydrologic treatment and plant status are reported to show differences among groups.
Abbreviations: NH<sub>4</sub><sup>+</sup> = ammonium, NO<sub>3</sub><sup>-</sup> = nitrate, MDL = below detection limit, C = carbon, N = nitrogen, P = phosphorus, K = potassium, Mg = magnesium, S = sulfur, Fe = iron, Mn = manganese, ppm = parts per million.

### Plant and soil redox effects on greenhouse gas fluxes

#### Methane (CH_4_)

Wet hydrologic treatment and no plants produced the highest CH_4_ fluxes and the greatest variability between samples (10.31 ± 45.54 mg CH_4_ m^−2^ h^−1^, *x̄* ± *SD*, Appendix S1: Table S2). When comparing plant vs. no plant CH_4_ fluxes at each of the five time points, no differences were detected, except during the second time point on June 28^th^, where CH_4_ concentrations were greater in the presence of plants than in the absence of plants (Appendix S1: Figure S2, Table S3). The wet treatment produced greater CH_4_ fluxes than the dry treatment throughout the middle three time points (June 28^th^ through July 25^th^), and differences in wet vs. interim and dry vs. interim were observed but did not persist throughout the experiment (Figure 3, Appendix S1: Table S4). The linear mixed-effects model with random effects only (sample mesocosm and date) was the most plausible (ΔAICc = 0), with an AICc weight of 0.37 (Appendix S1: Table S5).

**Figure 3.**
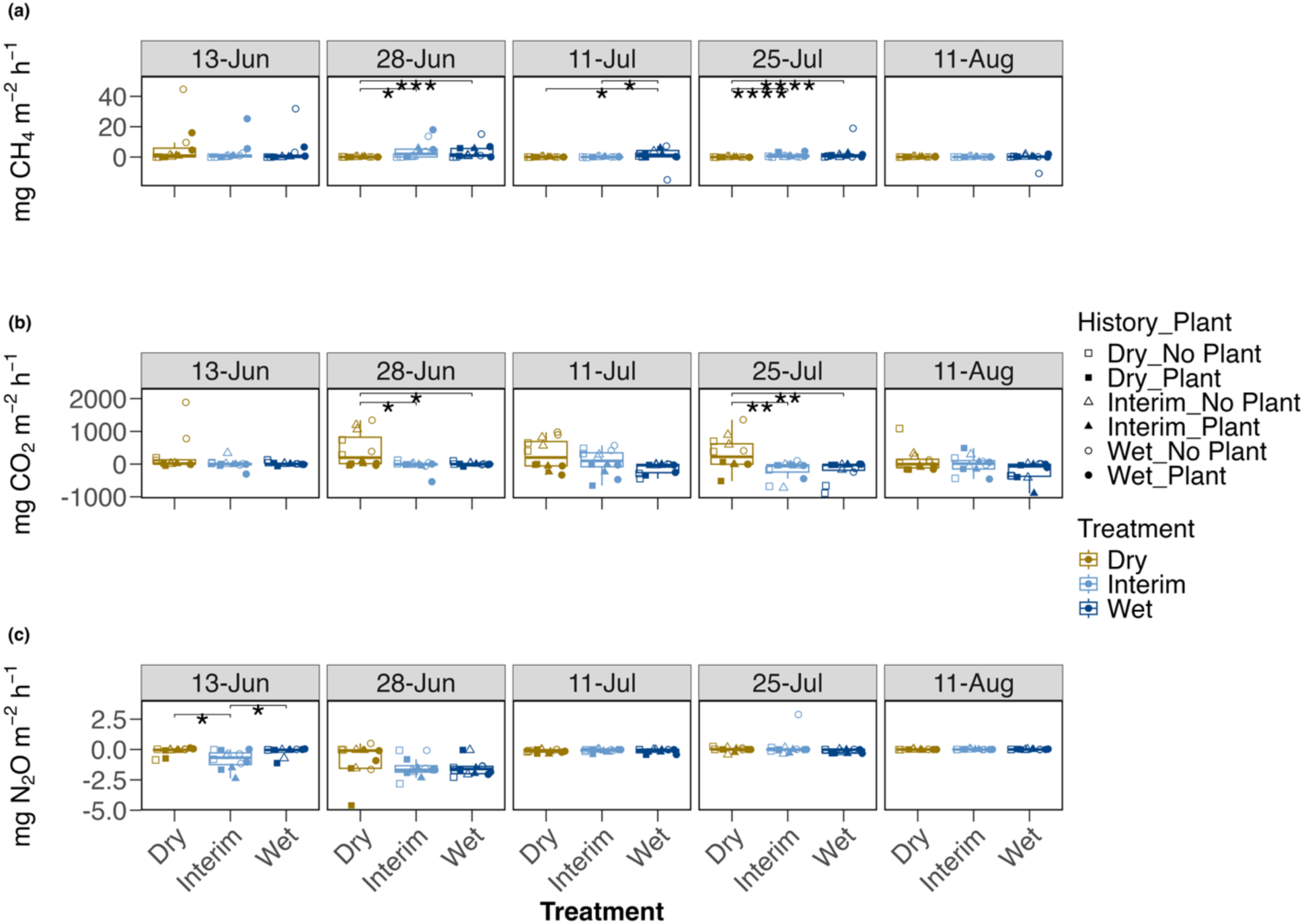
Greenhouse gas fluxes in dry, interim, and wet hydrologic treatments. Fluxes are in milligrams of methane (CH_4_) (a), carbon dioxide (CO_2_) (b), and nitrous oxide (N_2_O) (c) per square meter per hour (mg m^−2^ h^−1^) and are compared for hydrologic treatments during the five time points. Individual samples are plotted as shapes and colored by hydrologic treatment (brown squares = dry, light blue triangles = interim, dark blue circles = wet). Point shape represents hydrologic history (square = dry history, triangle = interim history, circle = wet history). Closed points represent mesocosms with plants, while open points represent mesocosms without plants. Boxplots summarize the median, first and third quartiles, and two whiskers extending |≤| 1.5 * interquartile range. Asterisks represent significance levels of Dwass, Steel, Critchlow, and Fligner (SDCFlig) tests (****: *p* ≤ 0.0001, ***: *p* ≤ 0.001, **: *p* ≤ 0.01, *: *p* ≤ 0.05, no symbol: *p* > 0.05). An extreme outlier of 248 mg CH_4_ m^−2^ h^−1^, measured in a mesocosm from interim soil history and wet treatment with no plants on July 11^th^, was used in statistical calculations but not displayed in the plot to improve visualization of differences across treatments. Month names are abbreviated as follows: Jun = June, Jul = July, Aug = August.

#### Carbon dioxide (CO_2_)

CO_2_ fluxes in the dry hydrologic treatment were greater than CO_2_ fluxes in the interim and wet treatments during two out of five time points (Figure 3, Appendix S1: Table S4). The linear mixed effects model, which included the interactive effects of hydrologic treatment and plant presence/absence, was the most plausible (ΔAICc = 0), and explained 49% of the variance in CO_2_ fluxes (conditional *R*^2^, Appendix S1: Table S5). In comparison, the fixed effect of plant presence/absence explained 12% of the variance in CO_2_ fluxes, and hydrologic treatment and hydrologic history explained 19% and 1%, respectively (Appendix S1: Table S5).

#### Nitrous oxide (N_2_O)

The N_2_O fluxes were near zero in all hydrologic and plant treatments (range: -4.61 – 2.88 mg N_2_O m^−2^ h^−1^) (Figure 3, Appendix S1: Figure S2). On July 11^th^ and July 25^th^, no plant N_2_O fluxes exceeded plant N_2_O fluxes (Appendix S1: Figure S2, Table S3). The N_2_O fluxes were similar among hydrologic treatments, except for lower fluxes in interim hydrology on June 13^th^ (Figure 3, Appendix S1: Table S4). The model with plant and hydrologic treatment additive effects was most plausible (ΔAICc = 0), based on an AICc weight of 0.25 (Appendix S1: Table S5).

### Relationship between soil redox conditions and GHG fluxes

We used linear regression to examine the relationship between soil redox status (represented by the percent of paint removed from IRIS tubes) and GHG fluxes. At the start of the experiment, the CH_4_ fluxes had a weak positive relationship with redox status (*F*_34_ = 3.14, *R*^2^ = 0.06, *p* = 0.09, Appendix S1: Table S6, Figure S3). At the end of the experiment, the CO_2_ fluxes had a negative relationship with redox conditions (*F*_34_ = 6.90, *R*^2^ = 0.14, *p* = 0.01, Appendix S1: Table S6, Figure S3). The N_2_O fluxes were not significantly related to redox conditions throughout the experiment (Appendix S1: Table S6, Figure S3).

### Patterns in microbial community composition

Bacterial and archaeal community composition clustered by hydrologic treatment within hydrologic history (Figure 4). Hydrologic history (PERMANOVA, *F*_2, 35_ = 7.562, *R*^2^ = 0.305, *p* = 0.001) and hydrologic treatment (PERMANOVA, *F*_2, 35_ = 2.605, *R*^2^ = 0.110, *p* = 0.002) explained variance in microbial community patterns (Figure 4, Appendix S1: Table S7). In contrast, microbial community composition was similar in the presence or absence of plant communities (PERMANOVA, *F*_1, 35_ = 0.698, *R*^2^ = 0.102, *p* = 0.824, Appendix S1: Table S7). Overall patterns in microbial composition and soil properties were significantly correlated (Mantel *r* = 0.384, *p* = 1.0 × 10^−4^, Appendix S1: Table S8).

**Figure 4.**
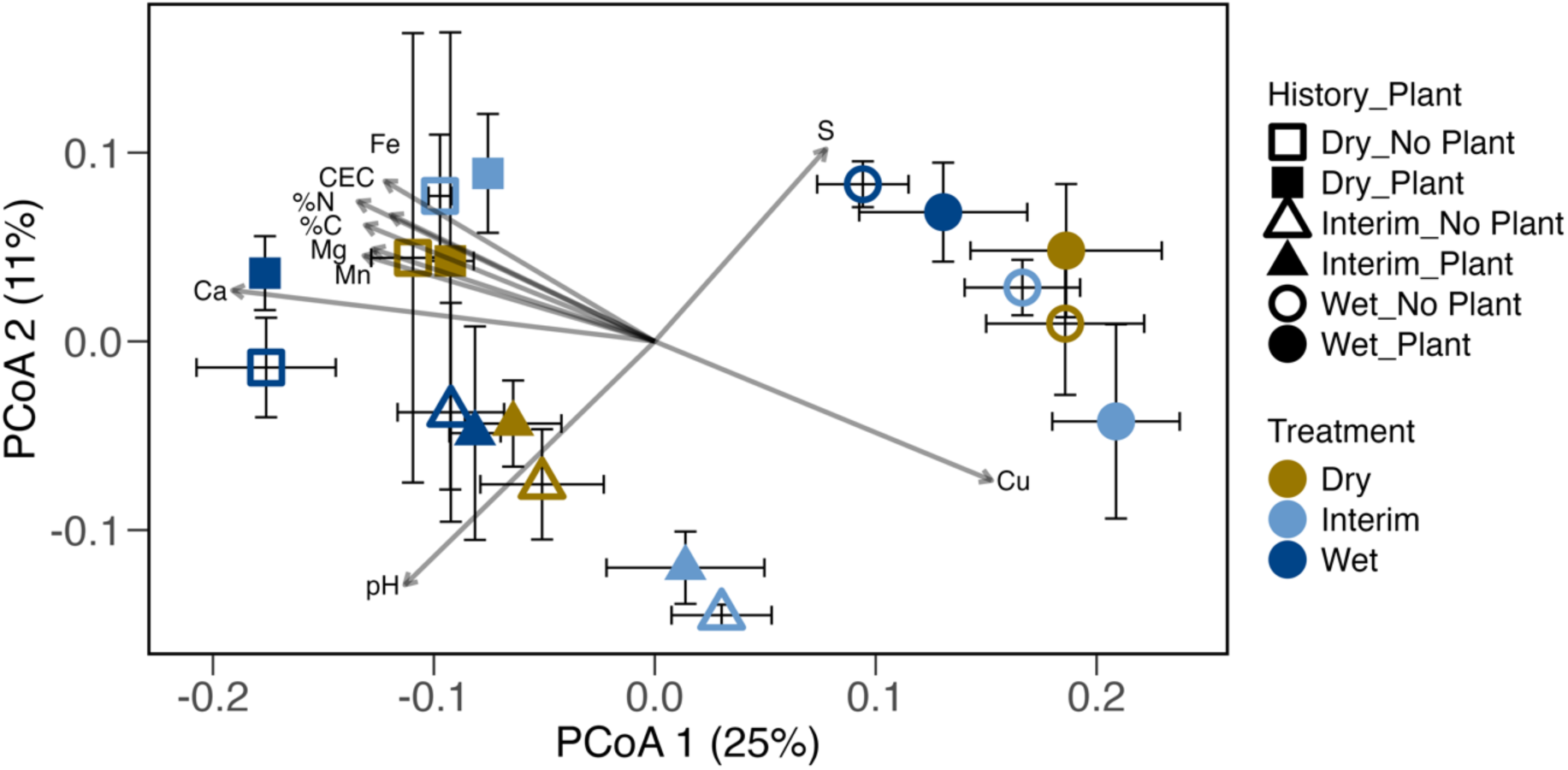
Ordination based on a principal coordinates analysis (PCoA) depicting the composition of the bacterial and archaeal communities based on the 16S rRNA gene. Colors refer to hydrologic treatments: brown indicates dry, light blue indicates interim, and dark blue indicates wet. Shapes refer to the hydrologic history of the sample: a square indicates dry, a triangle indicates interim, and a circle indicates wet. Open shapes refer to samples from mesocosms without plants, while closed shapes refer to samples from mesocosms with plants. Labeled vectors represent relationships among soil physicochemical variables and the microbial community ordination, with *p* ≤ 0.05, scaled by their correlation (using the *envfit* function). Abbreviations: CEC = cation exchange capacity, Fe = iron, S = sulfur, Cu = copper, Ca = calcium, Mn = manganese, Mg = magnesium, %C = percent carbon, %N = percent nitrogen.

Indicator species analysis of the top 2.5% of OTUs (by relative abundance at a significance level of *p* ≤ 0.01) revealed that dry history was represented by aerobic bacterial genera (*Gaiella*, *Reyranella*, and *Terrimonas*). Microbial taxa putatively associated with methane cycling, including the archaeal genus *Methanobacterium* and the bacterial genera *Methylocystis* and *Syntrophobacter,* were unique to microbial communities from wet hydrologic histories (Appendix S1: Table S9). Wet treatment was indicated by nitrogen-fixing taxa, *Anaeromyxobacter* and *Bradyrhizobium*, while *Reyranella* signaled both dry historical conditions and wet hydrologic treatment (Appendix S1: Table S10).

### Patterns in functional gene composition

The composition of functional genes was influenced by hydrologic conditions, but not by the presence or absence of plants. Carbohydrate metabolic composition differed between hydrologic treatments (PERMANOVA, *F*_1, 23_ = 2.735, *R*^2^ = 0.092, *p* = 0.013), while denitrification, methanogenesis, and CcO genes were similar across hydrologic treatments (Appendix S1: Table S7). Hydrologic history influenced the composition of genes associated with methanogenesis, central carbohydrate metabolism, and CcO (Appendix S1: Table S7).

Based on environmental fitting (*vegan::envfit*), denitrification functional gene composition correlated with carbon losses via CO_2_ flux (envfit, *R*^2^ = 0.243, *p* = 0.063, Figure 5). The PCoA of methanogenesis genes correlated with percent soil moisture (envfit, *R*^2^ = 0.201, *p* = 0.094), and the PCoA of central carbohydrate metabolism genes correlated with the percent of paint removed from IRIS tubes (envfit, *R*^2^ = 0.238, *p* = 0.059, Figure 5). The composition of CcO genes did not vary significantly with either GHG fluxes or soil redox conditions (Figure 5).

**Figure 5.**
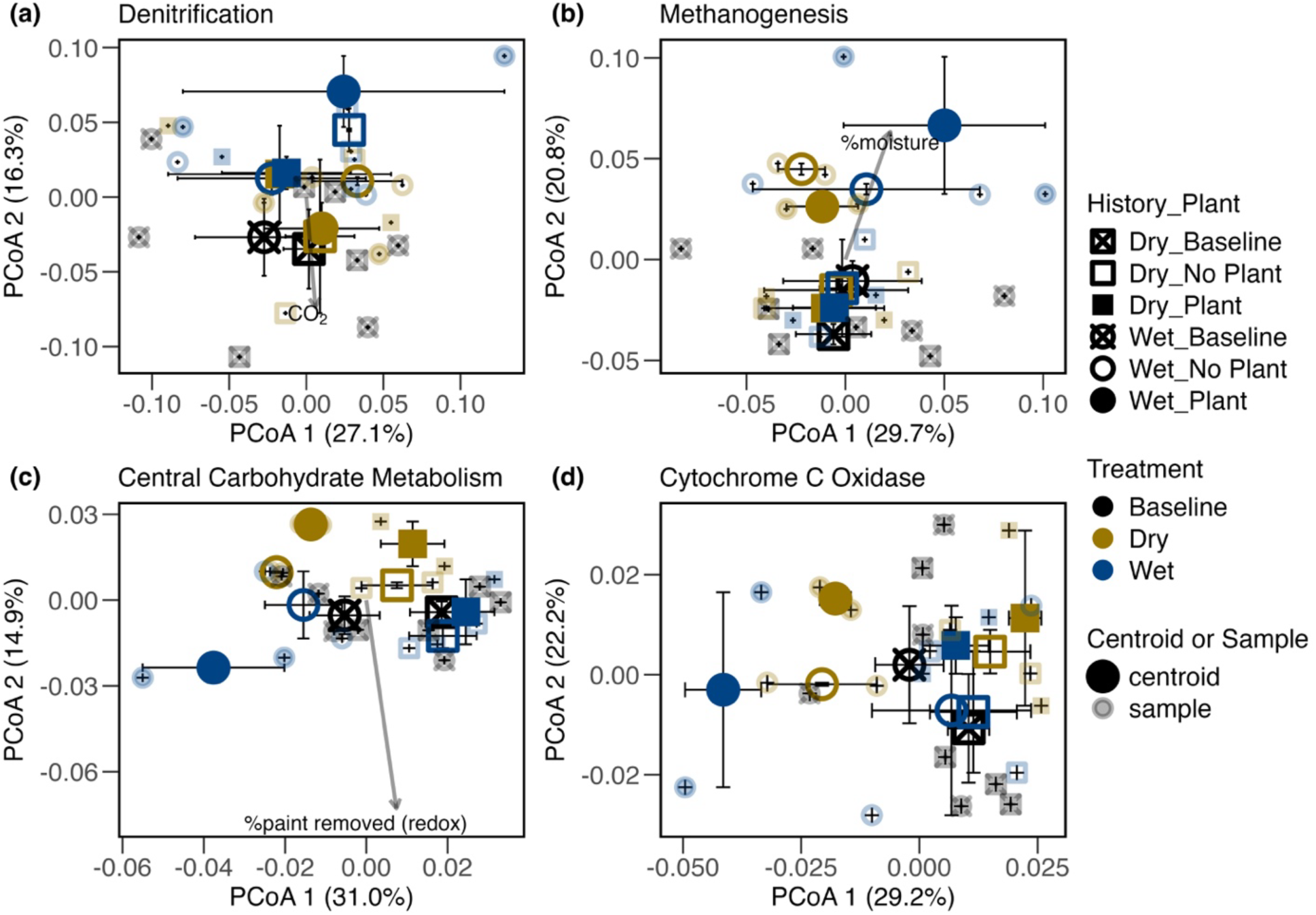
Ordination based on a principal coordinates analysis (PCoA) depicting community composition of the functional gene modules. The PCoA ordination is based on the Bray-Curtis distance of relative abundance of functional genes within respective Kyoto Encyclopedia of Genes and Genomes (KEGG) modules: Panel (a): Denitrification (KEGG Module M00529), (b): Methanogen (M00617), (c): Central Carbohydrate Metabolism (M00001-M00011, M00307-M00309, M00580, and M00633), (d): Prokaryotic Cytochrome C Oxidase (M00155). Colors refer to hydrologic treatments: black represents the baseline, brown represents dry conditions, and dark blue represents wet conditions. Shapes refer to the hydrologic history of the sample: a square indicates dry, a triangle indicates interim, and a circle indicates wet. The shape fill represents plant treatment: cross = baseline, open = no plant, closed = plant. Baseline samples were collected before the initiation of hydrologic and plant treatments. Large, dark, opaque points represent centroids of samples within a group, while smaller transparent points represent individual samples. Vectors represent significant (*p* ≤ 0.1) relationships among greenhouse gas fluxes, soil physicochemical factors, and functional gene composition, with the length of the vectors scaled by the strength of correlation (using the *envfit* function).

### Relationships between GHG fluxes, microbial community composition, and soil properties

Microbial community composition explained more variance in GHG fluxes than did soil properties. DBPLSR revealed that microbial community composition explained 58%, 37%, and 47% (components 1 and 2) of the variance in concentrations of CH_4_, CO_2_, and N_2_O fluxes, respectively (Appendix S1: Table S11). Patterns in soil properties, on the other hand, only explained 19%, 29%, and 26% of the variance in concentrations of CH_4_, CO_2_, and N_2_O fluxes, respectively, based on components 1 and 2 (Appendix S1: Table S11).

We evaluated relationships among the variance in GHG fluxes and functional gene composition at the beginning and end of the experiment using DBPLSR. Denitrification gene composition explained 54% of the variance in N_2_O production at the start of the experiment, and 60% of the variance in N_2_O fluxes at the end of the experiment (Appendix S1: Table S11, components 1 and 2). Methanogenesis functional gene composition explained a higher proportion of variance in CH_4_ fluxes in ‘start’ samples (63%) than ‘final’ samples (33%) (Appendix S1: Table S11, components 1 and 2). ‘Start’ compositions of CcO and carbohydrate metabolism explained 91% and 98% of variance in CO_2_ fluxes, respectively (Appendix S1: Table S11, components 1 and 2). At the end of the experiment, CcO composition only accounted for - 0.007% of the variance in CO_2_ fluxes, while carbohydrate metabolism explained 40% of the variance in CO_2_ fluxes (Appendix S1: Table S11, components 1 and 2).

## Discussion

This study revealed that historical hydrologic conditions influenced microbial taxonomic and functional composition more strongly than eight weeks of hydrologic treatments and plant removal manipulation. Hydrologic treatments impacted CH_4_ and CO_2_ fluxes but had little effect on N_2_O fluxes. Plant presence mitigated flood-induced reducing conditions in the rhizosphere but did not shift microbial community composition or soil GHG fluxes. These findings highlight the resistance of soil microbial community composition to short-term changes in hydrology and plant cover, indicating that a more plastic shift in community function underlies responses in CH_4_ and CO_2_ fluxes to hydrologic treatments.

### Past hydrology dictates bacterial community composition

Hydrologic history was a stronger driver of microbial community composition than our eight-week manipulation of hydrology and plant presence (Figure 4, Appendix S1: Table S7), indicating strong community resistance. This resistance was likely established through a combination of priority effects and protected soil microenvironments. During soil colonization following wetland restoration, priority effects would have allowed early arriving microbes to occupy the most favorable niches within the newly established hydrologic and vegetation regimes (Fukami et al. 2010; van de Voorde et al. 2011). Over time, this process leads to resource partitioning where organisms fill available niches and prevent the establishment of incoming taxa (Northfield et al. 2010). Additionally, hydrologic and plant manipulations may not have sufficiently altered the soil microenvironments (e.g., soil aggregates), which could have buffered the microbial community against environmental disturbance (Zhao et al. 2024; Ran et al. 2020). Soil physicochemical properties and microbial community composition are intertwined, and therefore, the relative impacts of biotic vs. abiotic controls on microbiome stability are difficult to distinguish (Griffiths et al. 2008; Griffiths and Philippot 2013). Collectively, the assembly and establishment by a historically adapted community and the persistence of soil microenvironments would maintain compositional stability, explaining why the short-term press disturbance from our treatments did not overpower the legacy community composition.

Our indicator species analysis provides evidence for observed legacy effects. For instance, the strictly aerobic genus *Gaiella* was an indicator for soils with a dry history (Albuquerque et al. 2011). In contrast, taxa associated with anaerobic processes like methanogenesis and iron reduction were indicators of soils with a wet history (Appendix S1: Table S9). Further evidence for how soil history confers resistance comes from the genus *Reyranella*, which was a significant indicator for soils with a dry history and those subjected to wet treatment. *Reyranella* is an aerobic genus in the family *Rhodospirillaceae* with variable capacity for nitrate reduction (Lee et al. 2017). This suggests *Reyranella* relied on aerobic respiration in the dry-history soils. When the mesocosms were flooded, *Reyranella* likely shifted from aerobic respiration to NO_3_ ^−^ reduction, using this metabolic flexibility to increase in relative abundance. This is an example of the community absorbing the disturbance via taxa already equipped for that specific environmental change, thereby maintaining compositional stability. While hydrologic history was the strongest driver of community composition, hydrologic treatment created a smaller but significant separation in community composition (Figure 4, Appendix S1: Table S7). This shift in composition, combined with changes in community activity, may explain observed differences in CH_4_ and CO_2_ fluxes.

### Dynamic hydrology influences greenhouse gas production more than plant presence versus absence

CH_4_ fluxes were highest, and CO_2_ fluxes were lowest in soils experiencing the wet hydrologic treatment (Figure 3). In the wet treatment, oxidizing conditions were maintained by plant roots and resulted in an organic plant root pattern due to oxidized iron paint on the IRIS tubes (Figure 2, Appendix S1: Figure S1). We expected that O_2_ radiation from plant roots would decrease soil CH_4_ fluxes by providing methanotrophs with O_2_ for CH_4_ consumption and supporting aerobic respiration at the expense of obligate anaerobic metabolisms like methanogenesis. However, methanotrophic functional gene composition was unresponsive to the presence or absence of plants (Finlay and Peralta 2025), and random effects (sample mesocosm and date) predominantly explained CH_4_ fluxes (Appendix S1: Table S5). Localized O_2_ availability adjacent to roots was insufficient to shift the bulk soil community composition or net CH_4_ fluxes. Patterns in net CH_4_ flux are further complicated by aerobic and anaerobic methanotrophs distributed across soil O_2_ gradients (e.g., hydrologic gradients) (Maietta et al. 2020). Despite the prevalence of random effects, the higher CH_4_ fluxes observed in wet versus dry treatments align with extensive literature reporting greater CH_4_ fluxes in saturated soils. (Christiansen et al. 2016; Maietta et al. 2020; Fairbairn et al. 2023). While CH_4_ and N_2_O fluxes were similar in the presence and absence of plants, CO_2_ fluxes displayed a predictable decline in the presence of plants.

The observed decline in CO_2_ fluxes in the presence of plants was unsurprising due to plant photosynthesis and fixation of atmospheric CO_2_ from within the transparent collection chamber (Luan and Wu 2014). The elevated CO_2_ fluxes in the dry hydrologic treatment are presumably due to aerobic respiration. A comparison of O_2_ and soil moisture impacts on CO_2_ fluxes found the greatest CO_2_ fluxes in intermediate soil moisture (Fairbairn et al. 2023). Percent soil moisture was moderate in our dry treatment (x̄ = 26.5%, Table 1). The moderate moisture likely promoted access to O_2_ while maintaining sufficient water content for dissolved C substrate access (Fairbairn et al. 2023). Plants consistently decreased CO_2_ fluxes throughout the experiment, while differences in GHG fluxes between hydrologic treatments were more variable. Therefore, models of wetland GHG emissions should consider plant cover as a stable control of CO_2_ fluxes. Conversely, the impacts of soil moisture on GHG fluxes vary dynamically over time and may require finer temporal resolution soil moisture measurements to model GHG fluxes accurately.

### Functional gene composition predicts greenhouse gas fluxes to varying degrees

#### Cytochrome C oxidase

We analyzed the composition of prokaryotic CcO genes as an indicator of aerobic respiration in oxidizing conditions. While hydrologic history influenced CcO composition, CcO composition did not explain significant variance in GHG fluxes during the eight-week incubation (Figure 5, Appendix S1: Table S7, Table S11). This lack of explanatory power is unsurprising, given the diversity of O_2_-reducing terminal oxidases and the known presence of CcO genes in strict anaerobes (Esposti 2020; Jabłońska and Tawfik 2019). There is no single enzyme that can be used to distinguish aerobic and anaerobic phenotypes because aerobic and anaerobic organisms do not exist as binary groups (Jabłońska and Tawfik 2019). Instead, a spectrum of oxygen usage phenotypes exists (Jabłońska and Tawfik 2019). Evaluating the presence and number of a full suite of O_2_-utilizing enzymes may be a useful indicator of aerobic vs. anaerobic status (Jabłońska and Tawfik 2019). Therefore, bioinformatic methods to facilitate this method ought to be further developed.

#### Central carbohydrate metabolism

While CcO composition was unable to explain differences in CO_2_ production across treatments, the composition of central carbohydrate metabolism genes was strongly related to differences in hydrologic history, hydrologic treatment, and the production of CO_2_ (Figure 5, Appendix S1: Table S7, Table S11). A previous study found a significant positive correlation between the abundance of carbohydrate-active enzymes and cumulative respiration in a tundra soil warming experiment (Johnston et al. 2019). Similarly, carbohydrate metabolism genes predicted differences in soil respiration between paddy and upland soils (Liu et al. 2020). While there is no single gene or defined collection of genes that have been consistently used to explain microbial community respiration responses to redox changes, various collections of carbohydrate metabolism genes often reflect differences in soil redox status and respiration (Johnston et al. 2019; Liu et al. 2020).

#### Denitrification

The composition of denitrification functional genes correlated with CO_2_ concentrations but not N_2_O concentrations (Figure 5). The lack of association with N_2_O is unsurprising since the denitrifier community composition included the suite of genes involved in the reduction of NO_3_ ^−^ to N_2_, with N_2_O as an intermediate product that could be consumed before detection via gas chromatography. The association between denitrification and CO_2_ production is likely the result of heterotrophic denitrification, where organic matter is oxidized to CO_2_ with NO_3_ ^−^ as an electron acceptor (Dlamini et al. 2020). Heterotrophic denitrification is typically a facultative process, found in organisms with the ability to respire aerobically when O_2_ is available, and also use NO_3_^−^to respire anaerobically when O_2_ availability decreases (Tiedje et al. 1989). A denitrification community-level response to soil hydrologic history and experimental drying/wetting has also been observed in a previous incubation experiment (Peralta et al. 2013). The lack of differences in denitrification functional gene composition across histories and treatments observed here may be explained by the diversity of denitrifiers and their O_2_ response phenotypes, and the variability of O_2_ concentrations in soil microenvironments (Rohe et al. 2021; Wallenstein et al. 2006; Ji et al. 2015).

#### Methanogenesis

Our results showed a strong influence of hydrologic history on methanogen functional gene composition (Figure 5, Appendix S1: Table S7), which supports the classical view of methanogenesis as a specialized and strictly anaerobic metabolism, maintained in the historically wet (i.e., low O_2_) field conditions (Lyu et al. 2018). The structure-function analysis (based on DBPLSR) provides another line of evidence that hydrologic history influenced CH_4_ production, where methanogen functional gene composition explained a greater proportion of variance in CH_4_ production at the start of the experiment than at the end of the experiment (Appendix S1: Table S11). Despite the lack of treatment effects on methanogen functional gene composition, CH_4_ fluxes did respond to hydrologic treatments, indicating changes in methanogen activity were used to respond to the short-term hydrologic press disturbance.

### Mismatch between microbial community composition and greenhouse gas function

The DNA-based functional gene composition explained more variance in GHG fluxes measured at the beginning than at the end of the experiment. Hydrologic history influenced wetland soil microbial community composition more than contemporary treatments. The decrease in the strength of structure-function associations from the beginning to the end of the experiment could be due to the DNA-based molecular measurement providing insight into historical and integrated environmental conditions, rather than contemporary changes to redox conditions. For example, DNA from dormant and dead cells contributes to DNA-based community composition, and these relic signatures are more indicative of past than present conditions (Lennon et al. 2018). Changes to functional gene composition over the eight-week experiment may have been obscured by the stable pool of dormant and relic DNA, thereby weakening structure-function relationships, as observed here for CH_4_ ∼ methanogenesis, CO_2_ ∼ central carbohydrate metabolism, and CO_2_ ∼ CcO (Appendix S1: Table S11).

Another explanation for the decreased structure-function associations could be the differences in temporal resolution between DNA-based molecular methods and *in situ* GHG measurements. Differential gene expression can substantially alter CH_4_ yields (Shi et al. 2014). Protein expression and activity may be altered in response to O_2_ concentrations (e.g., Lynch and Lin 1996, reviewed in Saleh-Lakha et al. 2005). These changes to gene expression and protein activity could alter GHG production rapidly, while changes to DNA abundance could alter GHG production over multiple microbial generations. Mesocosm microbial communities in the present study likely responded to experimental treatments (hydrology and plant presence or absence) by altering gene expression and protein activity. Changes to functional gene abundance were likely difficult to detect after eight weeks using DNA-based methods, hence the weakening of structure-function relationships in CH_4_ and CO_2_ production at the end of the experiment.

### Limitations

While this study provides valuable insights into the interplay between hydrology, plant presence, and microbial communities, several limitations should be considered when interpreting the results. First, the mesocosm setup, while necessary for experimental treatments, differs from wetland conditions. The experiment was conducted in a covered hoop house that prevented natural precipitation and altered light availability. Soil sampling may have altered soil structure, and mesocosms were disconnected from groundwater and deeper soil layers that may contribute to GHG emissions (Ball 2013; Sadat-Noori et al. 2016; Nikolenko et al. 2019). These factors could cause microbial responses to similar disturbances to differ in the field.

Finally, this study manipulated soils from a single restored coastal plain wetland. There are important differences between this restored wetland and remnant wetlands that restrict the generalization of results. The top 30 cm of soil were homogenized by decades of tillage (Hopfensperger et al. 2014). This likely reduced soil carbon storage, and 12 years post-restoration (at time of sampling), soil carbon storage likely lagged behind comparable remnant wetlands (Moreno-Mateos et al. 2012). Similarly, nitrogen storage and plant community structure in restored wetlands often recover but plateau below pre-degradation levels, representing an alternative stable state (Moreno-Mateos et al. 2012). Therefore, responses to hydrologic manipulation and loss of vegetation observed in the present study may vary from remnant wetland responses to similar disturbances. Increased restoration area (≥ 100 ha) and hydrologic connectivity are known to increase wetland restoration success (Moreno-Mateos et al. 2012). Both these features are present at the TOWeR site in the present study, and GHG flux responses to mesocosm hydrologic manipulation further underscore the importance of hydrology in wetland GHG function.

### Conclusions

The long-term hydrologic history of a wetland shapes the resident microbial community, fostering a high degree of resistance to short-term hydrologic disturbances and loss of vegetation. While microbial community composition remained largely stable, we observed shifts in microbial function in response to short-term hydrologic manipulations. Dry treatment favored CO_2_ production, and wet treatment favored CH_4_ emissions. Our findings highlight the need to incorporate historical hydrologic conditions into models predicting wetland greenhouse gas emissions. Further research focusing on the interplay of soil history, short-term soil redox shifts, and plant-microbe interactions will be crucial for developing effective wetland management and climate change mitigation strategies.

## Acknowledgments

We thank J. LeCrone, L. Armstrong, M. Stillwagon, G. Gunderson, C. Eakins, J. Basco, and C. Bledsoe for field and laboratory assistance. We thank J. Gill and the ECU grounds crew for their efforts in maintaining the grounds surrounding the shade house. We also thank M. Muscarella for contributions to microbiome analyses and anonymous reviewers for feedback on earlier versions of this manuscript. This work was supported by the National Science Foundation (Graduate Research Fellowship Program [GRFP] grant to R.B.B. and grant DEB #1845845 and CNH2 #2009185 to A.L.P.). This material is based upon work supported by the National Science Foundation under grant no. DGE-2125684. Any opinions, findings, and conclusions or recommendations expressed in this material are those of the author(s) and do not necessarily reflect the views of the National Science Foundation. The metagenomes were produced by the DOE JGI under the Community Science Program (CSP) (JGI CSP grant 503952). The work conducted by the DOE JGI, a DOE Office of Science User Facility, is supported under contract DE-AC02-05CH11231.

