## Appendix_S1 for "Wetland soil history shapes microbial community composition, while hydrologic disturbance alters greenhouse gas fluxes"

*ECOSPHERE*

**Appendix S1**

### ***Wetland soil history shapes microbial community composition, while hydrologic disturbance alters greenhouse gas fluxes***

Regina B. Bledsoe, Colin G. Finlay*, Ariane L. Peralta

Regina B. Bledsoe and Colin G. Finlay are co-first authors.

#### TABLES

**Table S1.** Calibration standards for gas chromatography.

| Dates of calibration | Methane (ppm) | Carbon dioxide (ppm) | Nitrous oxide (ppm) |
|---|---|---|---|
| June 13th and 28th, 2016 | 0, 1, 2, 3, 4, 5 | 0, 120, 240, 360, 480 | 0, 0.2, 0.4, 0.6, 0.8, 1 |
| July 11th, July 25th, and August 11th, 2016 | 0, 1, 4, 20, 40, 50 | 0, 120, 480, 1200, 2400, 3000 | 0, 0.2, 0.8, 20, 40, 50 |
| August 2021 – December 2022 (monthly) | 0, 1, 2, 3, 4, 5, 10, 20, 30, 40, 50 | 0, 120, 240, 360, 480, 600, 1200, 1800, 2400, 3000 | 0, 0.2, 0.4, 0.6, 0.8, 1, 10, 20, 30, 40, 50 |

Abbreviations: ppm: parts per million.

**Table S2.** Summary of greenhouse gas fluxes means and standard deviations, grouped by hydrologic treatment and plant presence/absence, to show variation between treatment groups.

| Treatment | Plant/No plant | $CH_4$ flux ($\bar{x} \pm SD$) | $CO_2$ flux ($\bar{x} \pm SD$) | $N_2O$ flux ($\bar{x} \pm SD$) |
|---|---|---|---|---|
| Dry | No plant | 1.85 ± 8.27 | 615.88 ± 470.55 | -0.13 ± 0.45 |
| Dry | Plant | 0.77 ± 3 | -59.98 ± 125.02 | -0.32 ± 0.88 |
| Interim | No plant | 0.75 ± 2.57 | 42.84 ± 287.2 | -0.3 ± 0.93 |
| Interim | Plant | 2.56 ± 5.57 | -106.15 ± 225.33 | -0.59 ± 0.82 |
| Wet | No plant | 10.31 ± 45.54 | -122.56 ± 236.67 | -0.33 ± 0.66 |
| Wet | Plant | 1.6 ± 2.2 | -78.8 ± 184.21 | -0.43 ± 0.68 |

Notes: Greenhouse gas flux was analyzed in terms of milligrams of gas per square meter per hour.

Abbreviations: $\bar{x}$: mean, *SD*: standard deviation.

**Table S3.** Wilcoxon Rank-Sum Test of greenhouse gas fluxes in no plant vs. plant treatments.

| Date | GHG (flux) | Group 1 | Group 2 | *V* | *p* |
|---|---|---|---|---|---|
| 13-Jun | $CH_4$ | No Plant | Plant | 48 | 0.108 |
| **28-Jun** | **$CH_4$** | **No Plant** | **Plant** | **38** | **0.038** |
| 11-Jul | $CH_4$ | No Plant | Plant | 74 | 0.640 |
| 25-Jul | $CH_4$ | No Plant | Plant | 55 | 0.196 |
| 11-Aug | $CH_4$ | No Plant | Plant | 58 | 0.246 |
| 13-Jun | $N_2O$ | No Plant | Plant | 99 | 0.580 |
| 28-Jun | $N_2O$ | No Plant | Plant | 108 | 0.347 |
| **11-Jul** | **$N_2O$** | **No Plant** | **Plant** | **144** | **0.009** |
| **25-Jul** | **$N_2O$** | **No Plant** | **Plant** | **140** | **0.016** |
| 11-Aug | $N_2O$ | No Plant | Plant | 56 | 0.212 |

Notes: Bold text indicates significant differences ($p \leq 0.05$). Wilcoxon Rank-Sum tests are not presented for $CO_2$ fluxes because plant photosynthesis was the driver of lower $CO_2$ fluxes in mesocosms with plants.

Abbreviations: GHG: greenhouse gas, *V*: Wilcoxon Rank Sum Test *V* statistic.

**Table S4.** Differences in greenhouse gas fluxes among hydrologic treatments.

| Date | GHG (flux) | $\chi^2$ | Kruskal-Wallis *p* | Group 1 | Group 2 | *W* | SDCFlig *p* |
|---|---|---|---|---|---|---|---|
| 13-Jun | $CH_4$ | 1.556 | 0.459 | NS | NS | NS | NS |
| | | | | **D** | **I** | **3.756** | **0.019** |
| **28-Jun** | **$CH_4$** | **13.351** | **0.001** | **D** | **W** | **5.062** | **$8.0 \times 10^{-4}$** |
| | | | | I | W | 0.327 | 0.985 |
| | | | | D | I | 1.470 | 0.580 |
| **11-Jul** | **$CH_4$** | **10.492** | **0.005** | **D** | **W** | **3.919** | **0.013** |
| | | | | **I** | **W** | **3.756** | **0.020** |
| | | | | **D** | **I** | **5.5522** | **0.000** |
| **25-Jul** | **$CH_4$** | **22.456** | **$1.3 \times 10^{-5}$** | **D** | **W** | **5.879** | **0.000** |
| | | | | I | W | 1.143 | 0.733 |
| 11-Aug | $CH_4$ | 4.821 | 0.090 | NS | NS | NS | NS |
| 13-Jun | $CO_2$ | 2.761 | 0.251 | NS | NS | NS | NS |
| | | | | **D** | **I** | **-3.756** | **0.021** |
| **28-Jun** | **$CO_2$** | **9.194** | **0.010** | **D** | **W** | **-3.511** | **0.034** |
| | | | | I | W | 0.980 | 0.799 |
| 11-Jul | $CO_2$ | 4.330 | 0.115 | NS | NS | NS | NS |
| | | | | **D** | **I** | **-4.491** | **0.002** |
| **25-Jul** | **$CO_2$** | **13.515** | **0.001** | **D** | **W** | **-4.409** | **0.003** |
| | | | | I | W | 0.735 | 0.887 |
| 11-Aug | $CO_2$ | 4.005 | 0.135 | NS | NS | NS | NS |
| | | | | **D** | **I** | **-3.674** | **0.024** |
| **13-Jun** | **$N_2O$** | **8.569** | **0.014** | D | W | -0.327 | 0.986 |
| | | | | **I** | **W** | **3.429** | **0.039** |
| 28-Jun | $N_2O$ | 5.722 | 0.057 | NS | NS | NS | NS |
| 11-Jul | $N_2O$ | 0.662 | 0.718 | NS | NS | NS | NS |
| 25-Jul | $N_2O$ | 3.743 | 0.154 | NS | NS | NS | NS |
| 11-Aug | $N_2O$ | 1.797 | 0.407 | NS | NS | NS | NS |

Notes: Kruskal-Wallis test p-values revealed differences between groups. If differences were identified ($p < 0.05$), then Dwass, Steel, and Critchlow-Fligner tests were used for pairwise comparisons. If the Kruskal-Wallis test did not reveal differences, NS (nonsignificant) is marked for pairwise comparisons. Rows with *p* values less than or equal to 0.05 are in bold. All Kruskal-Wallis tests (three groups) had two degrees of freedom. For all pairwise comparisons, there were 12 samples in each group. An SDCFlig *p* of 0.0000 indicates a *p* value smaller than $1 \times 10^{-5}$.

Abbreviations: GHG: greenhouse gas, $\chi^2$: chi-squared, SDCFlig P: Dwass, Steel, and Critchlow-Fligner P-value, NS: nonsignificant, D: dry, I: interim, W: wet.

**Table S5.** Summary of mixed effects models to explain variation in greenhouse gas fluxes ($CH_4$, $CO_2$, $N_2O$) due to fixed effects of hydrologic history, hydrologic treatment, and plant presence, and random effects of sample mesocosm and date.

| GHG | Model | K | AICc | ΔAICc | AICc Wt | $R^2$ Marg | $R^2$ Cond |
|---|---|---|---|---|---|---|---|
| $CH_4$ | null | 4 | 1580.84 | 0.00 | 0.37 | 0.000 | 0.000 |
| $CH_4$ | P | 5 | 1582.08 | 1.24 | 0.19 | 0.005 | 0.005 |
| $CH_4$ | T | 6 | 1582.88 | 2.04 | 0.13 | 0.012 | 0.012 |
| $CH_4$ | H | 6 | 1583.27 | 2.43 | 0.11 | 0.010 | 0.010 |
| $CH_4$ | P+T | 7 | 1584.16 | 3.32 | 0.07 | 0.017 | 0.017 |
| $CH_4$ | H+P | 7 | 1584.55 | 3.71 | 0.06 | 0.015 | 0.015 |
| $CH_4$ | T+H | 8 | 1585.38 | 4.54 | 0.04 | 0.022 | 0.022 |
| $CH_4$ | T*P | 9 | 1586.08 | 5.24 | 0.03 | 0.031 | 0.031 |
| $CH_4$ | P+T+H | 9 | 1586.71 | 5.87 | 0.02 | 0.027 | 0.027 |
| $CH_4$ | P*H | 9 | 1587.42 | 6.58 | 0.01 | 0.023 | 0.023 |
| $CO_2$ | T*P | 9 | 2546.69 | 0.00 | 1.00 | 0.477 | 0.494 |
| $CO_2$ | P+T | 7 | 2576.43 | 29.74 | $3.48 \times 10^{-7}$ | 0.312 | 0.496 |
| $CO_2$ | P+T+H | 9 | 2579.04 | 32.35 | $9.43 \times 10^{-8}$ | 0.325 | 0.495 |
| $CO_2$ | T | 6 | 2587.58 | 40.90 | $1.32 \times 10^{-9}$ | 0.192 | 0.496 |
| $CO_2$ | T+H | 8 | 2590.71 | 44.02 | $2.76 \times 10^{-10}$ | 0.205 | 0.496 |
| $CO_2$ | P | 5 | 2591.56 | 44.88 | $1.80 \times 10^{-10}$ | 0.120 | 0.496 |
| $CO_2$ | H+P | 7 | 2594.84 | 48.15 | $3.51 \times 10^{-11}$ | 0.133 | 0.496 |
| $CO_2$ | P*H | 9 | 2597.36 | 50.68 | $9.90 \times 10^{-12}$ | 0.156 | 0.496 |
| $CO_2$ | null | 4 | 2597.79 | 51.10 | $8.02 \times 10^{-12}$ | 0.000 | 0.496 |
| $CO_2$ | H | 6 | 2601.23 | 54.54 | $1.44 \times 10^{-12}$ | 0.013 | 0.496 |
| $N_2O$ | P+T | 7 | 340.49 | 0.00 | 0.25 | 0.031 | 0.426 |
| $N_2O$ | P | 5 | 340.73 | 0.24 | 0.22 | 0.016 | 0.411 |
| $N_2O$ | P+T+H | 9 | 341.51 | 1.03 | 0.15 | 0.042 | 0.437 |
| $N_2O$ | H+P | 7 | 341.74 | 1.26 | 0.13 | 0.027 | 0.422 |
| $N_2O$ | T | 6 | 343.28 | 2.79 | 0.06 | 0.015 | 0.410 |

| | | | | | | | |
|---|---|---|---|---|---|---|---|
| $N_2O$ | null | 4 | 343.40 | 2.92 | 0.06 | 0.000 | 0.401 |
| $N_2O$ | T*P | 9 | 344.09 | 3.61 | 0.04 | 0.033 | 0.429 |
| $N_2O$ | T+H | 8 | 344.35 | 3.87 | 0.04 | 0.025 | 0.421 |
| $N_2O$ | H | 6 | 344.50 | 4.02 | 0.03 | 0.011 | 0.406 |
| $N_2O$ | P*H | 9 | 345.77 | 5.28 | 0.02 | 0.028 | 0.423 |

Notes: Fixed effects: H = hydrologic history (dry, interim, wet), T = hydrologic treatment (dry, interim, wet), and P = plant (plant or no plant). The null model only considers the random effects (sample mesocosm and date).

Abbreviations: GHG: greenhouse gas, K: number of estimated parameters, AICc: Akaike's Information Corrected Criterion, ΔAICc: change in AICc, Wt: weight, $R^2$ Marg: marginal coefficient of determination, $R^2$ Cond: conditional coefficient of determination, H: hydrologic history, T: hydrologic treatment, P: plant, $CH_4$: methane, $CO_2$: carbon dioxide, and $N_2O$: nitrous oxide.

**Table S6**. Summary of linear regression: greenhouse gas fluxes as a function of redox status.

| Formula | Timepoint | Residual Standard Error | Degrees of Freedom | *F* | Adjusted $R^2$ | *p* |
|---|---|---|---|---|---|---|
| $CH_4$ ~ redox status | Start | 9.61 | 34 | 3.14 | 0.06 | 0.09 |
| $CH_4$ ~ redox status | Finish | 1.87 | 34 | 1.37 | 0.01 | 0.25 |
| $CO_2$ ~ redox status | Start | 350 | 34 | 0.12 | -0.03 | 0.73 |
| **$CO_2$ ~ redox status** | **Finish** | **297** | **34** | **6.90** | **0.14** | **0.01** |
| $N_2O$ ~ redox status | Start | 0.60 | 34 | 0.20 | -0.02 | 0.66 |
| $N_2O$ ~ redox status | Finish | 0.02 | 34 | $4.16 \times 10^{-3}$ | -0.03 | 0.95 |

Notes: Greenhouse gas fluxes were measured in milligrams of greenhouse gas per square meter per hour. Redox status is the percentage of paint removed from the IRIS (indicator of reduction in soils) tubes. Bold text indicates significant differences ($p \leq 0.05$).

Abbreviations: redox: oxidation-reduction.

**Table S7**. Summary of PERMANOVA comparing microbial community composition due to main effects (plant, hydrologic history, hydrologic treatment) and interaction between plant × history and plant × treatment.

| Gene Module | Main Effect | DF | SumSq | *F* | $R^2$ | *p* |
|---|---|---|---|---|---|---|
| Denitrification (KEGG M00529) | Plant | 2 | 0.010 | 0.840 | 0.086 | 0.576 |
| | History | 1 | $-2.6 \times 10^{-5}$ | -0.005 | $-2.4 \times 10^{-5}$ | 0.992 |
| | Treatment | 1 | 0.004 | 0.667 | 0.034 | 0.622 |
| | Plant × History | 2 | 0.004 | 0.386 | 0.040 | 0.932 |
| | Plant × Treatment | 1 | 0.002 | 0.309 | 0.016 | 0.891 |
| Methanogenesis (KEGG M00617) | Plant | 2 | 0.006 | 1.045 | 0.085 | 0.406 |
| | **History** | 1 | **0.011** | **3.795** | **0.154** | **0.007** |
| | Treatment | 1 | 0.004 | 1.184 | 0.048 | 0.314 |
| | Plant × History | 2 | 0.003 | 0.515 | 0.042 | 0.868 |
| | Plant × Treatment | 1 | 0.002 | 0.586 | 0.024 | 0.695 |
| Central Carbohydrate Metabolism (KEGG M00001-M00011, M00307-M00309, M00580, M00633) | Plant | 2 | 0.002 | 0.998 | 0.067 | 0.429 |
| | **History** | 1 | **0.006** | **6.262** | **0.211** | $\mathbf{1.0 \times 10^{-5}}$ |
| | **Treatment** | 1 | **0.003** | **2.735** | **0.092** | **0.013** |
| | Plant × History | 2 | 0.002 | 0.803 | 0.054 | 0.681 |
| | Plant × Treatment | 1 | 0.001 | 1.065 | 0.036 | 0.347 |
| Cytochrome C Oxidase (M00155) | Plant | 2 | 0.001 | 1.211 | 0.076 | 0.327 |
| | **History** | 1 | **0.002** | **7.224** | **0.227** | **0.001** |
| | Treatment | 1 | $3.1 \times 10^{-4}$ | 0.977 | 0.031 | 0.413 |
| | Plant × History | 2 | 0.001 | 1.388 | 0.087 | 0.255 |
| | Plant × Treatment | 1 | 0.001 | 2.491 | 0.078 | 0.081 |
| 16S rRNA gene | Plant | 1 | 0.034 | 0.698 | 0.102 | 0.824 |
| | **History** | **2** | **0.743** | **7.562** | **0.305** | **0.001** |
| | **Treatment** | **2** | **0.256** | **2.605** | **0.110** | **0.002** |
| | Plant × History | 2 | 0.066 | 0.667 | 0.027 | 0.942 |
| | Plant × Treatment | 2 | 0.059 | 0.598 | 0.024 | 0.986 |

Notes: Bold text indicates significant differences ($p \leq 0.05$). For each functional gene module, the total degrees of freedom are 23. For the 16S rRNA gene module, the total degrees of freedom are 35.

Abbreviations: PERMANOVA: permutational multivariate analysis of variance, DF: degrees of freedom, SumSq: sum of squares, KEGG: Kyoto Encyclopedia of Genes and Genomes.

**Table S8.** Summary of Mantel tests for correlation between soil properties, greenhouse gases, and microbial community composition.

| X distance matrix | Y distance matrix | Mantel statistic *r* | *p* |
|---|---|---|---|
| **Soil properties** | **Microbial community (16S rRNA gene)** | **0.384** | **1.0 x $10^{-4}$** |
| **Soil properties and greenhouse gases** | **Microbial community (16S rRNA gene)** | **0.398** | **1.0 x $10^{-4}$** |
| Greenhouse gases | Microbial community (16S rRNA gene) | 0.150 | 0.076 |

Notes: Soil properties include pH, $NH_4^+$ (mg/L), %C, %N, ppm P, ppm K, ppm Mg, ppm S, ppm Fe, ppm Mn, humic matter, and redox status. Greenhouse gases include fluxes of $CH_4$, $CO_2$, and $N_2O$. X distance matrices are based on Euclidean distance, while Y distance is the Bray-Curtis distance matrix of microbial community composition based on the 16S rRNA gene. Bold text indicates significant differences ($p \leq 0.05$).

**Table S9**. Summary of bacterial and archaeal Operational Taxonomic Units (OTUs) representative of hydrologic history based on indicator species analysis.

| Hst | IndVal | *p* | Phylum | Class | Family | Genus |
|---|---|---|---|---|---|---|
| D | 0.38 | 0.01 | *Proteobacteria* | *Alphaproteobacteria* | *Rhizobiales* | *Rhizobiales* |
| D | 0.43 | $1.2 \times 10^{-3}$ | *Acidobacteria* | *Acidobacteria Gp6* | *Gp6* | *Gp6* |
| D | 0.44 | $1.6 \times 10^{-3}$ | *Acidobacteria* | *Acidobacteria Gp6* | *Gp6* | *Gp6* |
| D | 0.51 | $2.0 \times 10^{-4}$ | *Bacteria* | *Bacteria* | *Bacteria* | *Bacteria* |
| D | 0.42 | $1.6 \times 10^{-3}$ | *Proteobacteria* | *Alphaproteobacteria* | *Rhizobiales* | *Rhizobiales* |
| D | 0.53 | $3.0 \times 10^{-4}$ | *Proteobacteria* | *Betaproteobacteria* | *Betaproteobacteria* | *Betaproteobacteria* |
| D | 0.40 | $4.2 \times 10^{-3}$ | *Proteobacteria* | *Betaproteobacteria* | *Betaproteobacteria* | *Betaproteobacteria* |
| D | 0.53 | $1.0 \times 10^{-4}$ | *Proteobacteria* | *Betaproteobacteria* | *Betaproteobacteria* | *Betaproteobacteria* |
| D | 0.41 | $2.4 \times 10^{-3}$ | *Acidobacteria* | *Acidobacteria Gp6* | *Gp6* | *Gp6* |
| D | 0.45 | $5.0 \times 10^{-4}$ | *Bacteroidetes* | *Sphingobacteriia* | *Chitinophagaceae* | *Terrimonas* |
| D | 0.51 | $1.0 \times 10^{-4}$ | *Acidobacteria* | *Acidobacteria Gp3* | *Gp3* | *Gp3* |
| D | 0.68 | $6.0 \times 10^{-4}$ | *Thaumarchaeota* | *Nitrososphaerales* | *Nitrososphaera* | *Nitrososphaera* |
| D | 0.48 | $1.0 \times 10^{-3}$ | *Actinobacteria* | *Actinobacteria* | *Gaiellaceae* | *Gaiella* |
| D | 0.53 | $2.0 \times 10^{-3}$ | *Actinobacteria* | *Actinobacteria* | *Actinobacteria* | *Actinobacteria* |
| D | 0.41 | $3.1 \times 10^{-3}$ | *Actinobacteria* | *Actinobacteria* | *Gaiellaceae* | *Gaiella* |
| D | 0.53 | $2.0 \times 10^{-4}$ | *Proteobacteria* | *Alphaproteobacteria* | *Sphingomonadaceae* | *Sphingomonadaceae* |
| D | 0.54 | $3.0 \times 10^{-4}$ | *Acidobacteria* | *Acidobacteria Gp6* | *Gp6* | *Gp6* |
| D | 0.53 | $1.0 \times 10^{-4}$ | *Bacteroidetes* | *Sphingobacteriia* | *Chitinophagaceae* | *Chitinophagaceae* |
| D | 0.48 | $3.0 \times 10^{-4}$ | *Bacteria* | *Bacteria* | *Bacteria* | *Bacteria* |
| D | 0.52 | 0.01 | *Proteobacteria* | *Betaproteobacteria* | *Nitrosomonadaceae* | *Nitrosospira* |
| D | 0.44 | 0.01 | *Proteobacteria* | *Alphaproteobacteria* | *Reyranella* | *Reyranella* |
| D | 0.53 | $2.2 \times 10^{-3}$ | *Acidobacteria* | *Acidobacteria Gp6* | *Gp6* | *Gp6* |
| D | 0.79 | $1.0 \times 10^{-4}$ | *Proteobacteria* | *Gammaproteobacteria* | *Gammaproteobacteria* | *Gammaproteobacteria* |
| D | 0.47 | $1.0 \times 10^{-4}$ | *Proteobacteria* | *Alphaproteobacteria* | *Rhodospirillales* | *Rhodospirillales* |
| D | 0.42 | $3.7 \times 10^{-3}$ | *Bacteria* | *Bacteria* | *Bacteria* | *Bacteria* |

| | | | | | | |
|---|---|---|---|---|---|---|
| D | 0.51 | 1.1 x $10^{-3}$ | *Actinobacteria* | *Actinobacteria* | *Actinomycetales* | *Actinomycetales* |
| D | 0.66 | 1.0 x $10^{-4}$ | *Proteobacteria* | *Gammaproteobacteria* | *Gammaproteobacteria* | *Gammaproteobacteria* |
| D | 0.47 | 2.0 x $10^{-4}$ | *Bacteria* | *Bacteria* | *Bacteria* | *Bacteria* |
| D | 0.54 | 2.1 x $10^{-3}$ | *Verrucomicrobia* | *Spartobacteria* | *Spartobacteria* | *Spartobacteria* |
| D | 0.47 | 1.0 x $10^{-3}$ | *Verrucomicrobia* | *Subdivision3* | *Subdivision3* | *Subdivision3* |
| D | 0.50 | 5.0 x $10^{-4}$ | *Planctomycetes* | *Planctomycetia* | *Planctomycetaceae* | *Planctomycetaceae* |
| D | 0.51 | 3.8 x $10^{-3}$ | *Bacteria* | *Bacteria* | *Bacteria* | *Bacteria* |
| D | 0.41 | 0.01 | *Bacteria* | *Bacteria* | *Bacteria* | *Bacteria* |
| I | 0.39 | 2.7 x $10^{-3}$ | *Proteobacteria* | *Alphaproteobacteria* | *Rhizobiales* | *Rhizobiales* |
| I | 0.48 | 2.0 x $10^{-4}$ | *Acidobacteria* | *Acidobacteria Gp1* | *Gp1* | *Gp1* |
| I | 0.47 | 7.0 x $10^{-4}$ | *Bacteria* | *Bacteria* | *Bacteria* | *Bacteria* |
| I | 0.46 | 1.1 x $10^{-3}$ | *Acidobacteria* | *Acidobacteria Gp3* | *Gp3* | *Gp3* |
| I | 0.48 | 4.3 x $10^{-3}$ | *Bacteria* | *Bacteria* | *Bacteria* | *Bacteria* |
| I | 0.47 | 1.5 x $10^{-3}$ | *Planctomycetes* | *Planctomycetia* | *Planctomycetaceae* | *Planctomycetaceae* |
| I | 0.48 | 5.0 x $10^{-4}$ | *Proteobacteria* | *Deltaproteobacteria* | *Deltaproteobacteria* | *Deltaproteobacteria* |
| I | 0.46 | 0.01 | *Proteobacteria* | *Betaproteobacteria* | *Betaproteobacteria* | *Betaproteobacteria* |
| I | 0.44 | 0.01 | *Proteobacteria* | *Alphaproteobacteria* | *Hyphomicrobiaceae* | *Pedomicrobium* |
| I | 0.43 | 1.1 x $10^{-3}$ | *Planctomycetes* | *Planctomycetia* | *Planctomycetaceae* | *Planctomycetaceae* |
| I | 0.51 | 1.0 x $10^{-4}$ | *Acidobacteria* | *Acidobacteria Gp3* | *Gp3* | *Gp3* |
| I | 0.63 | 1.0 x $10^{-4}$ | *Acidobacteria* | *Acidobacteria Gp10* | *Gp10* | *Gp10* |
| I | 0.45 | 9.0 x $10^{-4}$ | *Acidobacteria* | *Acidobacteria Gp7* | *Gp7* | *Gp7* |
| I | 0.47 | 1.0 x $10^{-3}$ | *Chloroflexi* | *Ktedonobacteria* | *Ktedonobacteria* | *Ktedonobacteria* |
| I | 0.47 | 1.0 x $10^{-4}$ | *Acidobacteria* | *Acidobacteria Gp3* | *Gp3* | *Gp3* |
| I | 0.47 | 7.0 x $10^{-4}$ | *Proteobacteria* | *Alphaproteobacteria* | *Rhizobiales* | *Rhizobiales* |
| I | 0.44 | 0.01 | *Bacteria* | *Bacteria* | *Bacteria* | *Bacteria* |
| I | 0.56 | 4.7 x $10^{-3}$ | *Proteobacteria* | *Betaproteobacteria* | *Betaproteobacteria* | *Betaproteobacteria* |
| I | 0.47 | 9.0 x $10^{-4}$ | *Proteobacteria* | *Betaproteobacteria* | *Betaproteobacteria* | *Betaproteobacteria* |
| I | 0.47 | 2.5 x $10^{-3}$ | *Planctomycetes* | *Planctomycetia* | *Planctomycetaceae* | *Planctomycetaceae* |
| I | 0.49 | 1.1 x $10^{-3}$ | *Chloroflexi* | *Ktedonobacteria* | *Ktedonobacterales* | *Ktedonobacterales* |

| | | | | | | |
|---|---|---|---|---|---|---|
| I | 0.44 | 1.1 x $10^{-3}$ | *Bacteria* | *Bacteria* | *Bacteria* | *Bacteria* |
| I | 0.65 | 4.0 x $10^{-4}$ | *Bacteria* | *Bacteria* | *Bacteria* | *Bacteria* |
| W | 0.56 | 1.0 x $10^{-4}$ | *Acidobacteria* | *Acidobacteria Gp1* | *Gp1* | *Gp1* |
| W | 0.49 | 1.0 x $10^{-4}$ | *Acidobacteria* | *Acidobacteria Gp1* | *Gp1* | *Gp1* |
| W | 0.67 | 1.0 x $10^{-4}$ | *Acidobacteria* | *Acidobacteria Gp1* | *Gp1* | *Gp1* |
| W | 0.48 | 2.0 x $10^{-4}$ | *Actinobacteria* | *Actinobacteria* | *Solirubrobacterales* | *Solirubrobacterales* |
| W | 0.43 | 4.6 x $10^{-3}$ | *Proteobacteria* | *Alphaproteobacteria* | *Roseiarcaceae* | *Roseiarcus* |
| W | 0.43 | 2.0 x $10^{-4}$ | *Actinobacteria* | *Actinobacteria* | *Thermomonosporaceae* | *Actinoallomurus* |
| W | 0.43 | 1.4 x $10^{-3}$ | *Proteobacteria* | *Alphaproteobacteria* | *Alphaproteobacteria* | *Alphaproteobacteria* |
| W | 0.54 | 1.0 x $10^{-4}$ | *Acidobacteria* | *Acidobacteria Gp1* | *Gp1* | *Gp1* |
| W | 0.71 | 2.0 x $10^{-4}$ | *Acidobacteria* | *Acidobacteria Gp1* | *Acidobacteria Gp1* | *Acidobacteria Gp1* |
| W | 0.46 | 1.0 x $10^{-4}$ | *Proteobacteria* | *Alphaproteobacteria* | *Rhizomicrobium* | *Rhizomicrobium* |
| W | 0.62 | 1.0 x $10^{-4}$ | *Bacteria* | *Bacteria* | *Bacteria* | *Bacteria* |
| W | 0.68 | 1.0 x $10^{-4}$ | *Proteobacteria* | *Deltaproteobacteria* | *Geobacteraceae* | *Geobacter* |
| W | 0.59 | 5.0 x $10^{-4}$ | *Chloroflexi* | *Chloroflexi* | *Chloroflexi* | *Chloroflexi* |
| W | 0.58 | 1.0 x $10^{-4}$ | *Bacteria* | *Bacteria* | *Bacteria* | *Bacteria* |
| W | 0.74 | 1.0 x $10^{-4}$ | *Bacteria* | *Bacteria* | *Bacteria* | *Bacteria* |
| W | 0.65 | 1.0 x $10^{-4}$ | *Proteobacteria* | *Alphaproteobacteria* | *Methylocystaceae* | *Methylocystis* |
| W | 0.74 | 1.0 x $10^{-4}$ | *Acidobacteria* | *Acidobacteria Gp1* | *Gp1* | *Gp1* |
| W | 0.63 | 1.0 x $10^{-4}$ | *Acidobacteria* | *Acidobacteria Gp1* | *Gp1* | *Gp1* |
| W | 0.63 | 1.0 x $10^{-4}$ | *Verrucomicrobia* | *Subdivision3* | *Subdivision3* | *Subdivision3* |
| W | 0.61 | 6.0 x $10^{-4}$ | *Bacteria* | *Bacteria* | *Bacteria* | *Bacteria* |
| W | 0.55 | 2.0 x $10^{-3}$ | *Bacteria* | *Bacteria* | *Bacteria* | *Bacteria* |
| W | 0.56 | 2.0 x $10^{-4}$ | *Acidobacteria* | *Acidobacteria Gp2* | *Gp2* | *Gp2* |
| W | 0.60 | 4.0 x $10^{-4}$ | *Proteobacteria* | *Gammaproteobacteria* | *Gammaproteobacteria* | *Gammaproteobacteria* |
| W | 0.85 | 1.0 x $10^{-4}$ | *Verrucomicrobia* | *Subdivision3* | *Subdivision3* | *Subdivision3* |
| W | 0.84 | 1.0 x $10^{-4}$ | *Acidobacteria* | *Acidobacteria Gp1* | *Gp1* | *Gp1* |
| W | 0.64 | 1.0 x $10^{-4}$ | *Proteobacteria* | *Deltaproteobacteria* | *Syntrophobacteraceae* | *Syntrophobacter* |

| | | | | | | |
|---|---|---|---|---|---|---|
| W | 0.41 | $4.7 \times 10^{-3}$ | *Actinobacteria* | *Actinobacteria* | *Actinomycetales* | *Actinomycetales* |
| W | 0.52 | $1.3 \times 10^{-3}$ | *Proteobacteria* | *Deltaproteobacteria* | *Geobacteraceae* | *Geobacter* |
| W | 0.84 | $1.0 \times 10^{-4}$ | *Bacteria* | *Bacteria* | *Bacteria* | *Bacteria* |
| W | 0.49 | 0.01 | *Acidobacteria* | *Acidobacteria Gp1* | *Acidobacteria Gp1* | *Acidobacteria Gp1* |
| W | 0.68 | $6.0 \times 10^{-4}$ | *Acidobacteria* | *Acidobacteria Gp1* | *Acidobacteria Gp1* | *Acidobacteria Gp1* |
| W | 0.93 | $1.0 \times 10^{-4}$ | *Euryarchaeota* | *Methanobacteria* | *Methanobacteriaceae* | *Methanobacterium* |
| W | 0.47 | $1.0 \times 10^{-4}$ | *Acidobacteria* | *Acidobacteria Gp1* | *Gp1* | *Gp1* |

Notes: These are the top OTUs (>2.5% relative abundance) that are significantly ($p \leq 0.01$) associated with dry, interim, and wet hydrologic histories. Text files of indicator species can be accessed online: https://doi.org/10.5281/zenodo.15528281.

Abbreviations: Hst: hydrologic history, IndVal: indicator value, D: dry history, I: interim history, W: wet history.

**Table S10**. Summary of bacterial and archaeal (OTUs) representative of hydrologic treatments based on indicator species analysis.

| Trt | IndVal | *p* | Phylum | Class | Family | Genus |
|---|---|---|---|---|---|---|
| I | 0.46 | 0.01 | *Acidobacteria* | *Acidobacteria Gp2* | *Gp2* | *Gp2* |
| I | 0.46 | $4.6 \times 10^{-3}$ | *Proteobacteria* | *Deltaproteobacteria* | *Geobacteraceae* | *Geobacter* |
| I | 0.47 | $5.4 \times 10^{-3}$ | *Acidobacteria* | *Acidobacteria Gp2* | *Gp2* | *Gp2* |
| I | 0.44 | $2.0 \times 10^{-3}$ | *Verrucomicrobia* | *Subdivision3* | *Subdivision3* | *Subdivision3* |
| I | 0.51 | $3.0 \times 10^{-3}$ | *Verrucomicrobia* | *Subdivision3* | *Subdivision3* | *Subdivision3* |
| I | 0.62 | 0.01 | *Acidobacteria* | *Acidobacteria Gp2* | *Gp2* | *Gp2* |
| W | 0.40 | $9.0 \times 10^{-4}$ | *Proteobacteria* | *Alphaproteobacteria* | *Bradyrhizobiaceae* | *Bradyrhizobium* |
| W | 0.42 | $1.8 \times 10^{-3}$ | *Proteobacteria* | *Alphaproteobacteria* | *Rhizobiales* | *Rhizobiales* |
| W | 0.40 | $1.8 \times 10^{-3}$ | *Proteobacteria* | *Alphaproteobacteria* | *Hyphomicrobiaceae* | *Hyphomicrobium* |
| W | 0.45 | 0.01 | *Proteobacteria* | *Deltaproteobacteria* | *Cystobacteraceae* | *Anaeromyxobacter* |
| W | 0.61 | $1.0 \times 10^{-4}$ | *Bacteria* | *Bacteria* | *Bacteria* | *Bacteria* |
| W | 0.47 | $1.0 \times 10^{-4}$ | *Proteobacteria* | *Alphaproteobacteria* | *Rhizobiales* | *Rhizobiales* |
| W | 0.45 | $2.3 \times 10^{-3}$ | *Proteobacteria* | *Alphaproteobacteria* | *Reyranella* | *Reyranella* |
| W | 0.41 | $2.0 \times 10^{-3}$ | *Proteobacteria* | *Alphaproteobacteria* | *Rhodospirillaceae* | *Rhodospirillaceae* |
| W | 0.54 | $1.6 \times 10^{-3}$ | *Proteobacteria* | *Deltaproteobacteria* | *Cystobacteraceae* | *Anaeromyxobacter* |

Notes: These are the top OTUs (>2.5% relative abundance) that are significantly ($p \leq 0.01$) associated with dry, interim, and wet hydrologic treatments. No significant indicator species were identified for the dry hydrologic treatment. Text files of indicator species can be accessed online: https://doi.org/10.5281/zenodo.15528281.

Abbreviations: Trt: hydrologic treatement, IndVal: indicator value, I: interim treatment, W: wet treatment

**Table S11.** Summary of distance-based partial least squares regression representing the percent of variance in greenhouse gas fluxes explained by the first two components of models derived from functional gene composition, 16S rRNA gene composition, or soil properties distance matrix.

| GHG ~ Gene module or 16S rRNA or soils | Timepoint | Component | Comp 1 | Comp 2 |
|---|---|---|---|---|
| $CH_4$ ~ Methanogenesis | Start | $R^2$ | 0.52 | 0.74 |
| | | adjusted $R^2$ | 0.44 | 0.63 |
| | | gvar | 26.0 | 62.0 |
| | | crit | 6.10 | 4.56 |
| $CH_4$ ~ Methanogenesis | Final | $R^2$ | 0.19 | 0.42 |
| | | adjusted $R^2$ | 0.13 | 0.33 |
| | | gvar | 35.8 | 58.0 |
| | | crit | 0.43 | 0.35 |
| $CO_2$ ~ Central Carbohydrate Metabolism | Start | $R^2$ | 0.69 | 0.98 |
| | | adjusted $R^2$ | 0.64 | 0.98 |
| | | gvar | 43.6 | 59.9 |
| | | crit | 4671 | 329 |
| $CO_2$ ~ Central Carbohydrate Metabolism | Final | $R^2$ | 0.25 | 0.48 |
| | | adjusted $R^2$ | 0.20 | 0.40 |
| | | gvar | 27.1 | 54.4 |
| | | crit | 5649 | 4464 |
| $CO_2$ ~ Cytochrome C Oxidase | Start | $R^2$ | 0.41 | 0.93 |
| | | adjusted $R^2$ | 0.31 | 0.91 |
| | | gvar | 70.9 | 88.6 |
| | | crit | 8996 | 1388 |
| $CO_2$ ~ Cytochrome C Oxidase | Final | $R^2$ | 0.09 | 0.13 |
| | | adjusted $R^2$ | 0.03 | $-6.8 \times 10^{-3}$ |
| | | gvar | 27.4 | 76.2 |
| | | crit | 6858 | 7552 |
| $CO_2$ ~ Denitrification | Start | $R^2$ | 0.31 | 0.79 |
| | | adjusted $R^2$ | 0.19 | 0.71 |
| | | gvar | 45.0 | 60.2 |
| | | crit | 10585 | 4282 |
| $CO_2$ ~ Denitrification | Final | $R^2$ | 0.61 | 0.80 |
| | | adjusted $R^2$ | 0.58 | 0.76 |
| | | gvar | 19.3 | 46.3 |
| | | crit | 2955 | 1768 |
| $N_2O$ ~ Denitrification | Start | $R^2$ | 0.28 | 0.67 |
| | | adjusted $R^2$ | 0.16 | 0.54 |
| | | gvar | 50.1 | 62.0 |
| | | crit | 0.0071 | 0.0044 |
| $N_2O$ ~ Denitrification | Final | $R^2$ | 0.36 | 0.65 |
| | | adjusted $R^2$ | 0.32 | 0.60 |
| | | gvar | 38.46 | 47.4 |

| | | | | |
|---|---|---|---|---|
| | | crit | $1.0 \times 10^{-5}$ | $6.5 \times 10^{-6}$ |
| $CH_4$ ~ 16S rRNA gene | Final | $R^2$ | 0.17 | 0.60 |
| | | adjusted $R^2$ | 0.14 | 0.58 |
| | | gvar | 31.5 | 42.3 |
| | | crit | 0.08 | 0.04 |
| $CO_2$ ~ 16S rRNA gene | Final | $R^2$ | 0.29 | 0.40 |
| | | adjusted $R^2$ | 0.27 | 0.37 |
| | | gvar | 16.1 | 43.9 |
| | | crit | 2082 | 1868 |
| $N_2O$ ~ 16S rRNA gene | Final | $R^2$ | 0.39 | 0.50 |
| | | adjusted $R^2$ | 0.38 | 0.47 |
| | | gvar | 13.8 | 43.1 |
| | | crit | $6.4 \times 10^{-6}$ | $5.5 \times 10^{-6}$ |
| $CH_4$ ~ soil properties | Final | $R^2$ | 0.14 | 0.24 |
| | | adjusted $R^2$ | 0.11 | 0.19 |
| | | gvar | 31.1 | 63.4 |
| | | crit | 0.09 | 0.08 |
| $CO_2$ ~ soil properties | Final | $R^2$ | 0.12 | 0.33 |
| | | adjusted $R^2$ | 0.10 | 0.29 |
| | | gvar | 44.5 | 54.5 |
| | | crit | 2591 | 2092 |
| $N_2O$ ~ soil properties | Final | $R^2$ | 0.10 | 0.30 |
| | | adjusted $R^2$ | 0.76 | 0.26 |
| | | gvar | 41.6 | 54.3 |
| | | crit | $9.4 \times 10^{-6}$ | $7.8 \times 10^{-6}$ |

Notes: Functional gene distance matrices are based on the Bray-Curtis dissimilarities of gene relative abundances within respective KEGG modules (see main text for KEGG module numbers). The 16S rRNA gene matrix is based on the Bray-Curtis distance. Euclidean distance matrices represent soil parameters and concentrations of greenhouse gases.

Abbreviations: GHG: greenhouse gas, comp: component, gvar: total weighted geometric variability, crit: value of criterion defined in method.

#### FIGURES

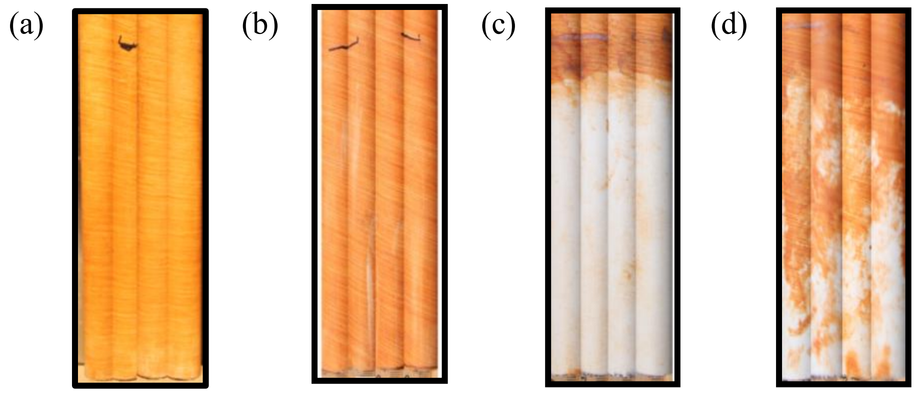


**Figure S1.** Indicator of Reduction in Soils (IRIS) tubes. Representative IRIS tubes collected from the dry hydrologic treatment without plants (a) and with plants (b), wet treatment without plants (c), and wet treatment with plants (d). Photo credit: R. Bledsoe.

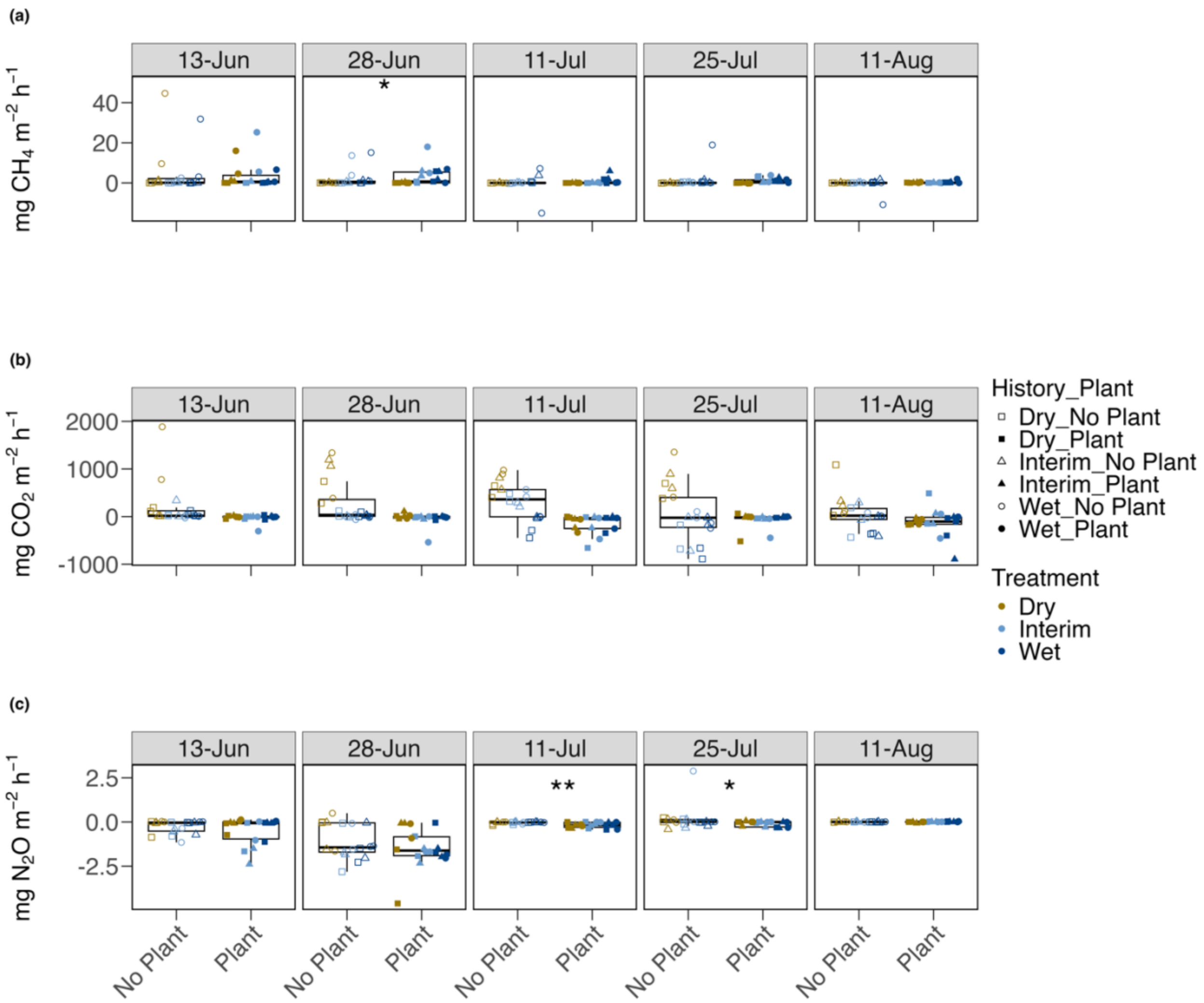


**Figure S2.** Greenhouse gas fluxes comparing no plant vs. plant treatments. Fluxes are in milligrams of $CH_4$ (a), $CO_2$ (b), and $N_2O$ (c) per square meter per hour, and are compared for plant presence/absence treatments during the five time points. Individual samples are plotted as shapes and colored by hydrologic treatment (brown squares = dry, light blue triangles = interim, dark blue circles = wet). Point shape represents hydrologic history (square = dry history, triangle = interim history, circle = wet history). Closed points represent mesocosms with plants, while open points represent mesocosms without plants. Boxplots summarize the median, first and third quartiles, and two whiskers extending |≤| 1.5 * interquartile range. Asterisks represent significance levels of Wilcoxon Rank-Sum tests (****: $p \leq 0.0001$, ***: $p \leq 0.001$, **: $p \leq 0.01$,

*: $p \leq 0.05$, no symbol: $p > 0.05$). An extreme outlier of 248 mg $CH_4$ $m^{-2}$ $h^{-1}$, measured in a wet treatment with no plants on July 11th, was used in statistical calculations but not displayed in the plot to improve visualization of differences across treatments. Wilcoxon Rank-Sum tests are not presented for $CO_2$ fluxes because plant photosynthesis was the driver of lower $CO_2$ fluxes in mesocosms with plants.

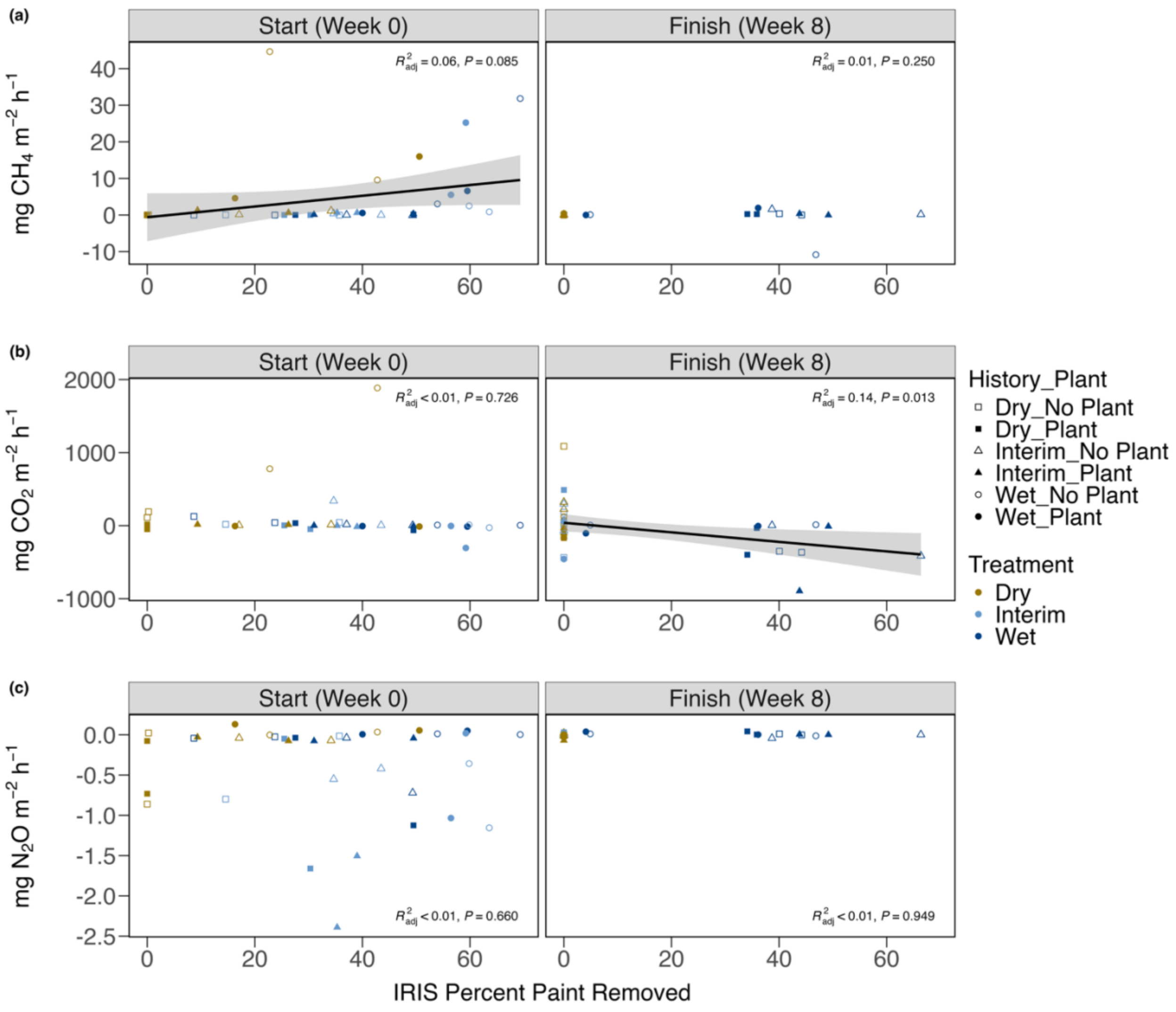


**Figure S3.** Greenhouse gas fluxes as a function of redox status. IRIS (indicator of reduction in soils) percent is the percent of paint removed from IRIS tubes, with a higher percent of paint removed corresponding to more reducing conditions. Individual samples are plotted as points and colored by hydrologic treatment (brown = dry, light blue = interim, dark blue = wet). Grey shading surrounding the line of best fit represents the standard error. Linear regression lines (greenhouse gas ~ IRIS percent) are shown in black, and associated statistics ($R^2_{adj}$: adjusted coefficient of determination and $P$: $p$ value) are printed on each plot.

**Supplemental Methods**

*Amplicon library preparation*

For each sample, three 50 µL PCR libraries were prepared by combining 35.75 µL molecular grade water, 5 µL Amplitaq Gold 360 10x buffer, 5 µL $MgCl_2$ (25 mM), 1 µL dNTPs (40mM total, 10mM each), 0.25 µL Amplitaq Gold 360 polymerase, 1 µL 515 forward barcoded primer (10 µM), 1 µL 806 reverse primer (10 µM), and 1 µL DNA template (10 ng $\mu L^{-1}$). Thermocycler conditions for PCR reactions were as follows: initial denaturation (94 °C, 3 minutes); 30 cycles of 94 °C for 45 seconds, 50 °C for 30 seconds, 72 °C for 90 seconds; final elongation (72 °C, 10 minutes). The triplicate 50 µL PCR libraries were combined and then cleaned using the Axygen® AxyPrep Magnetic (MAG) Bead Purification Kit (Axygen, Union City, California, USA). Cleaned PCR products were quantified using the QuantIT dsDNA BR assay (Thermo Scientific, Waltham, Massachusetts, USA) and diluted to a concentration of 10 ng/µL before pooling the libraries at an equimolar concentration of 5 ng/µL.

*Sequencing quality control and rarefaction*

Amplicon sequencing efforts of the 16S rRNA gene returned 1,570,135 reads and 35,897 OTUs before removing low-abundance sequences occurring less than 10 times or 0.01% in all samples. After removing low-abundance sequences, 1,505,797 reads representing 7,026 OTUs were retained after filtering. All samples were rarefied to 23,528 reads since this was the lowest read count among all samples.

Following quality control and filtering of raw metagenomic sequencing data, annotation and gene calling resulted in 600,507 ± 172,049 ($\bar{x} \pm SD$) annotated contigs per sample. An average of 998,604 ± 307,431 gene features per sample were identified. Of the identified gene features, an average of 232,809 ± 70,486 KEGG features per sample were identified. Among the four curated functional gene sets, central carbohydrate metabolism had the highest average KEGG features per sample (13,145 ± 3,848), followed by methanogenesis (1,434 ± 423), CcO (1,325 ± 398.26), and denitrification (350 ± 111).
