## Appendix_S2 for "Wetland soil history shapes microbial community composition, while hydrologic disturbance alters greenhouse gas fluxes"

*ECOSPHERE*

**Appendix S2**

### ***Wetland soil history shapes microbial community composition, while hydrologic disturbance alters greenhouse gas fluxes***

Regina B. Bledsoe, Colin G. Finlay*, Ariane L. Peralta

Regina B. Bledsoe and Colin G. Finlay are co-first authors.

HMR and robust linear are displayed if compatible with the flux data. If an HMR fit is displayed, then HMR was used for the final flux calculation. If no HMR fit is present then robust linear was used for the final flux calculation. If neither HMR nor robust linear fit was appropriate for the time series, then a linear fit was used for the final flux calculation. Time is in hours. N = No plant, P = Plant.

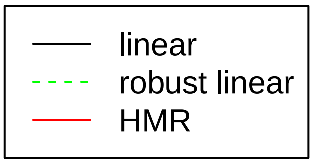


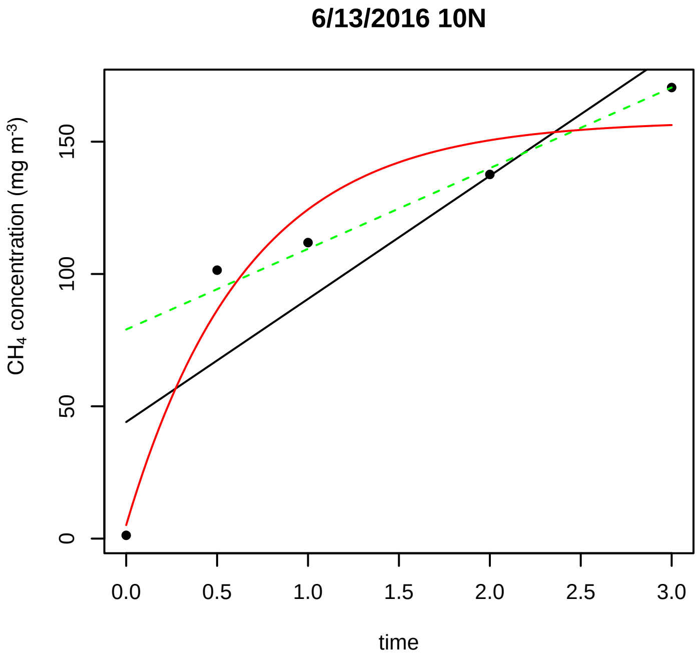


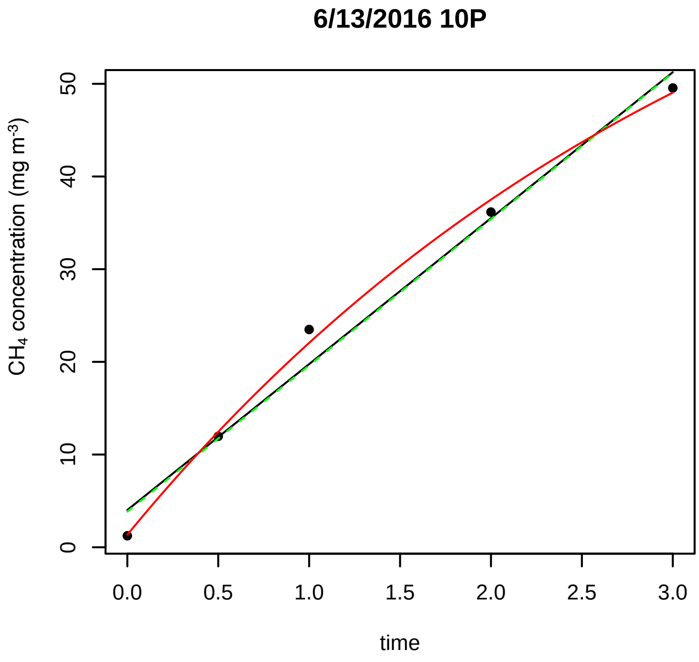


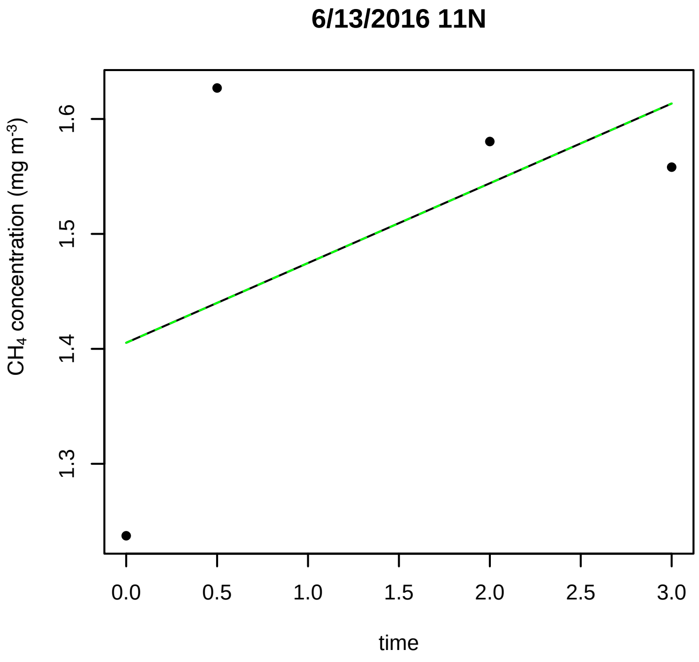


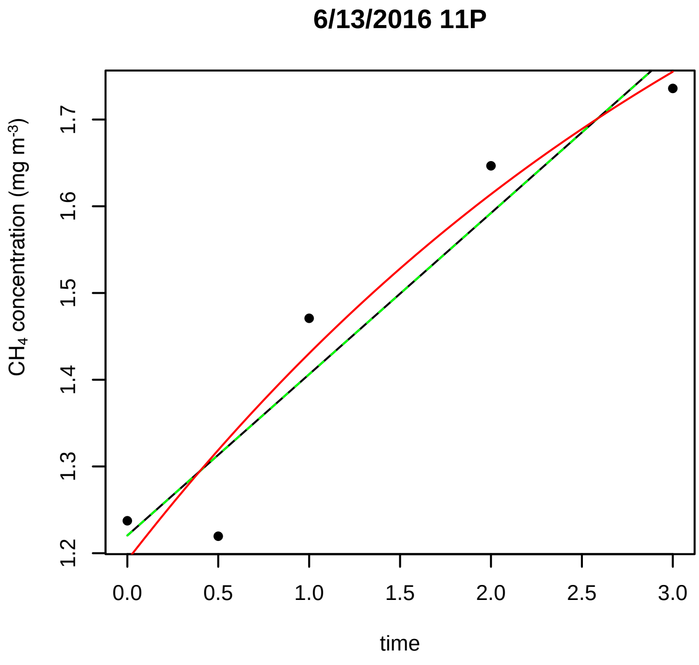


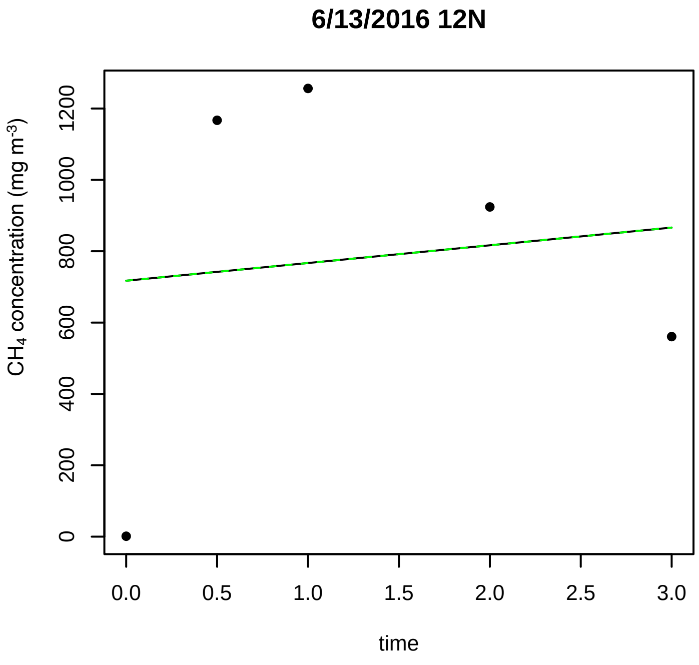

6/13/2016 12P

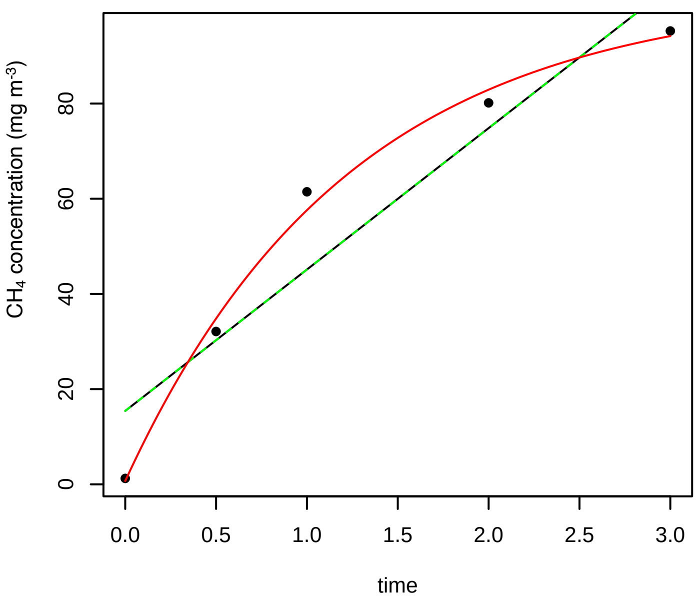


6/13/2016 13N

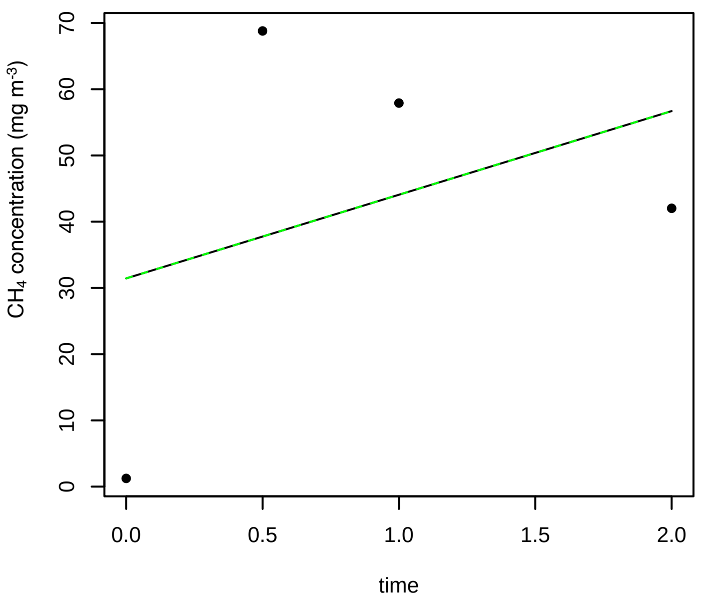


6/13/2016 13P

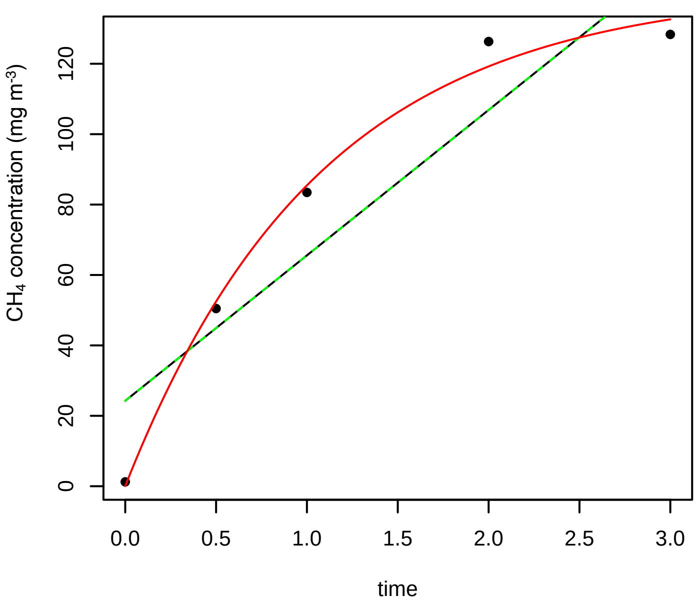


6/13/2016 14N

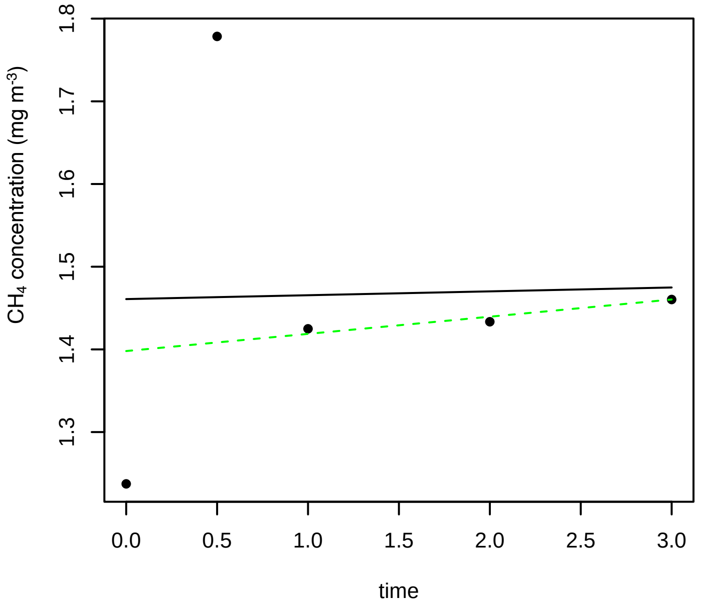


6/13/2016 14P

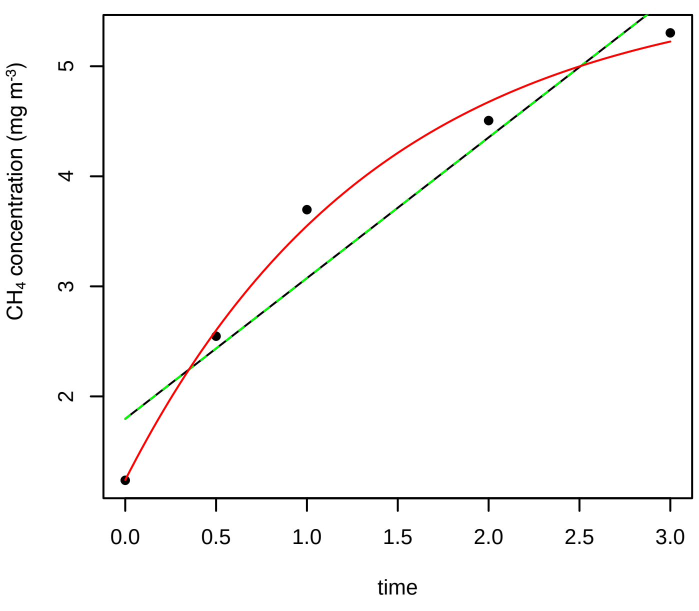


6/13/2016 15N

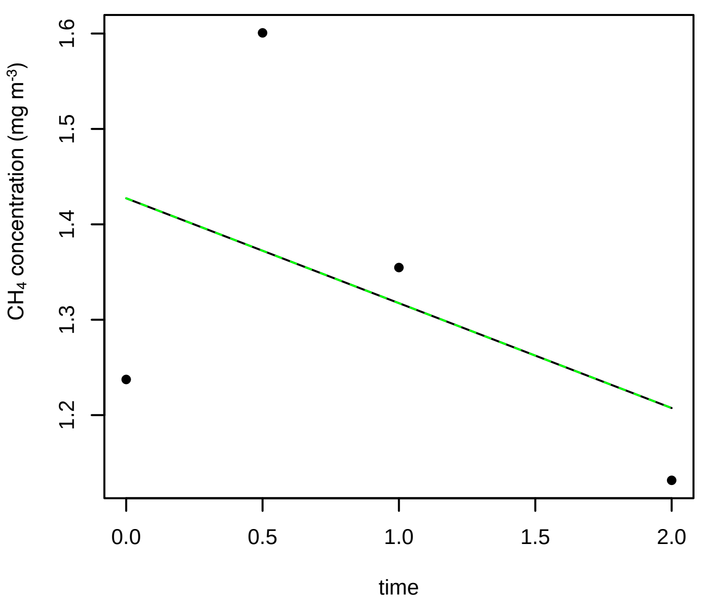

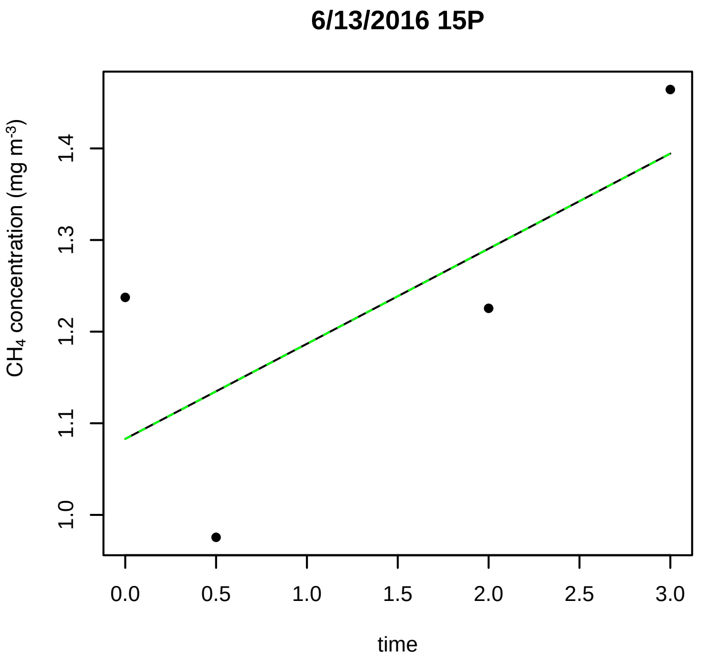
6/13/2016 15P
CH$_4$ concentration (mg m$^{-3}$)
1.0
1.1
1.2
1.3
1.4
0.0
0.5
1.0
1.5
2.0
2.5
3.0
time

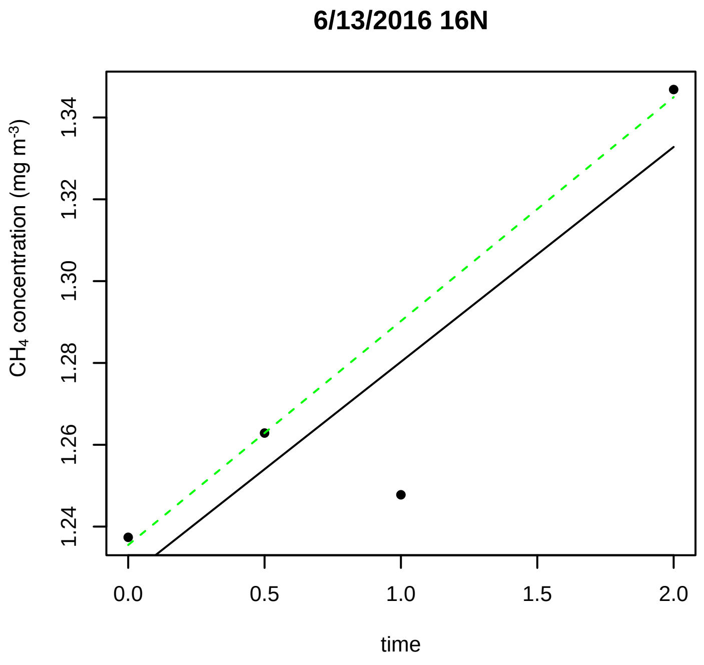
6/13/2016 16N
CH$_4$ concentration (mg m$^{-3}$)
1.24
1.26
1.28
1.30
1.32
1.34
0.0
0.5
1.0
1.5
2.0
time

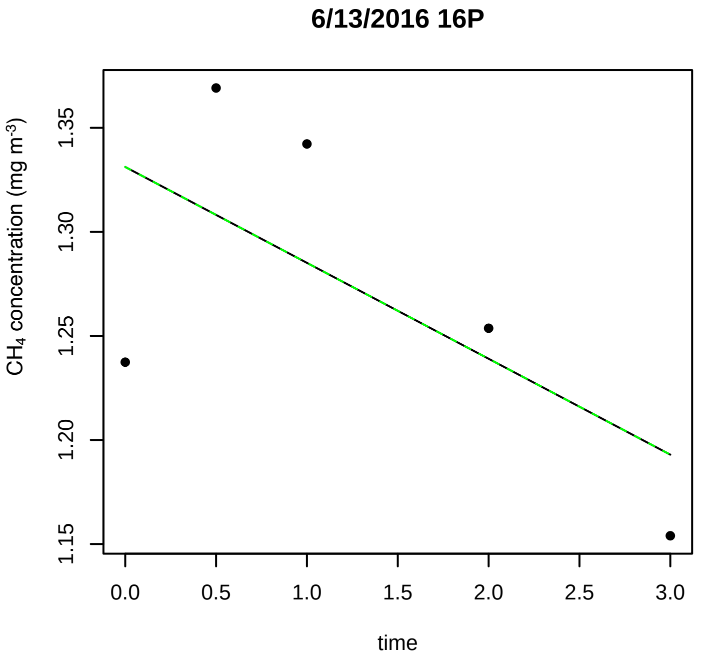
6/13/2016 16P
CH$_4$ concentration (mg m$^{-3}$)
1.15
1.20
1.25
1.30
1.35
0.0
0.5
1.0
1.5
2.0
2.5
3.0
time

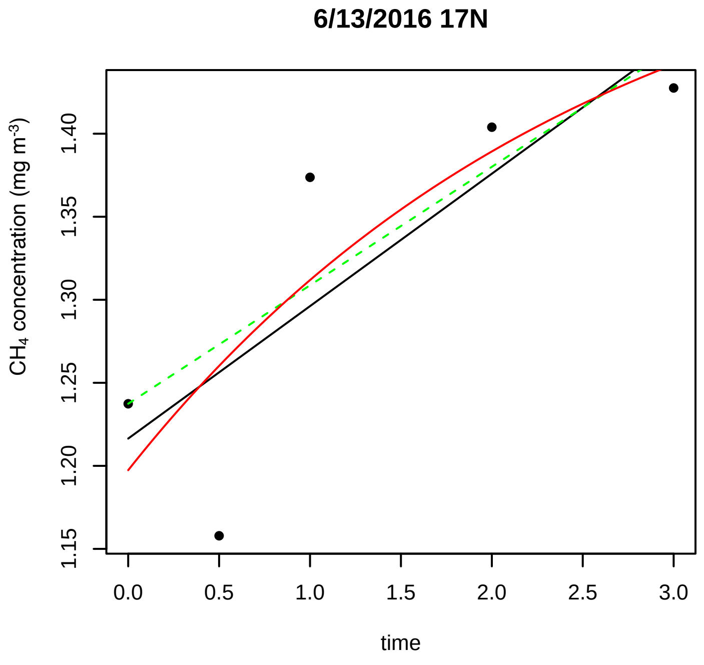
6/13/2016 17N
CH$_4$ concentration (mg m$^{-3}$)
1.15
1.20
1.25
1.30
1.35
1.40
0.0
0.5
1.0
1.5
2.0
2.5
3.0
time

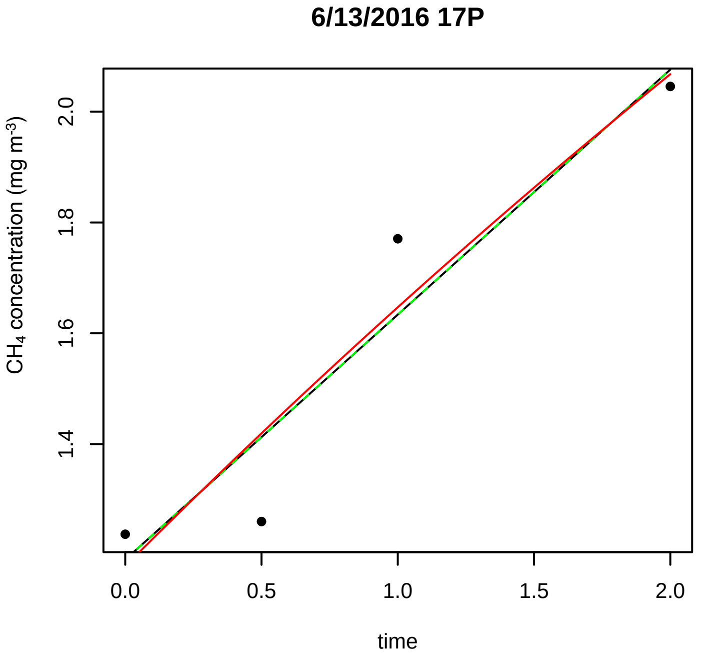
6/13/2016 17P
CH$_4$ concentration (mg m$^{-3}$)
1.4
1.6
1.8
2.0
0.0
0.5
1.0
1.5
2.0
time

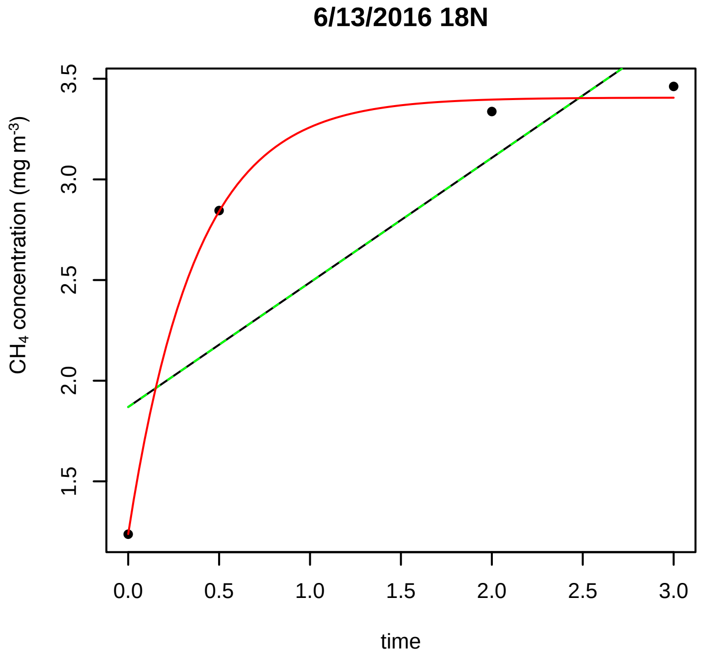
6/13/2016 18N
CH$_4$ concentration (mg m$^{-3}$)
1.5
2.0
2.5
3.0
3.5
0.0
0.5
1.0
1.5
2.0
2.5
3.0
time

6/13/2016 18P

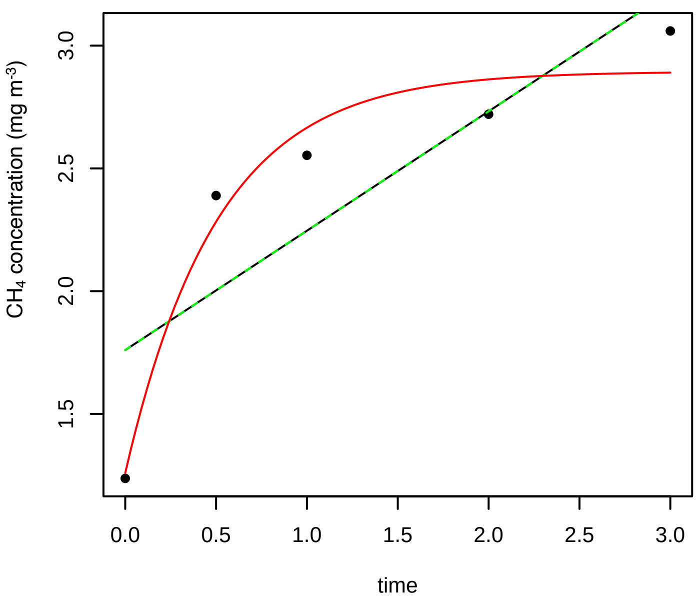


6/13/2016 1N

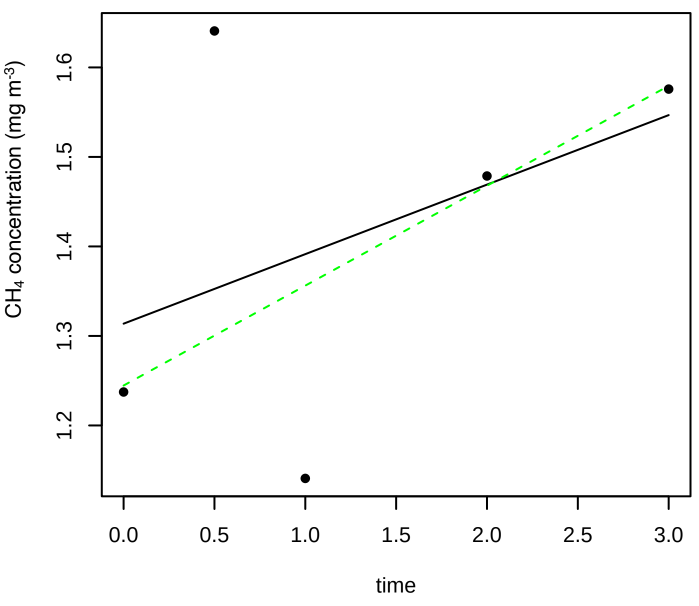


6/13/2016 1P

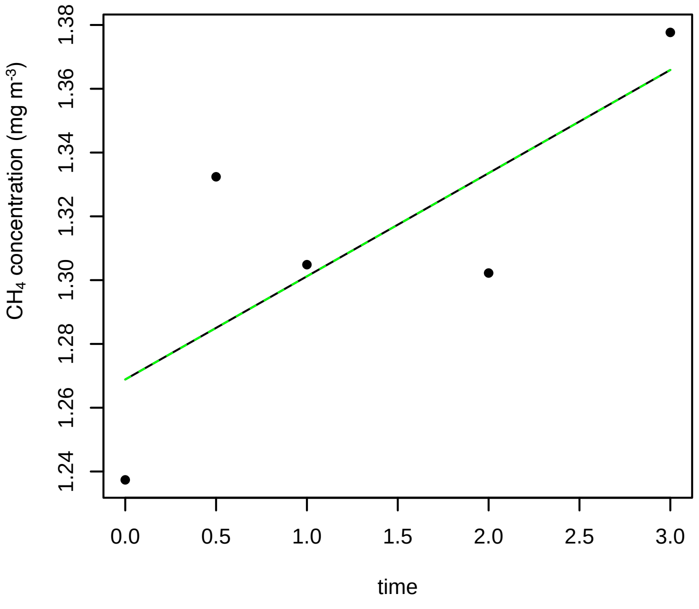


6/13/2016 2N

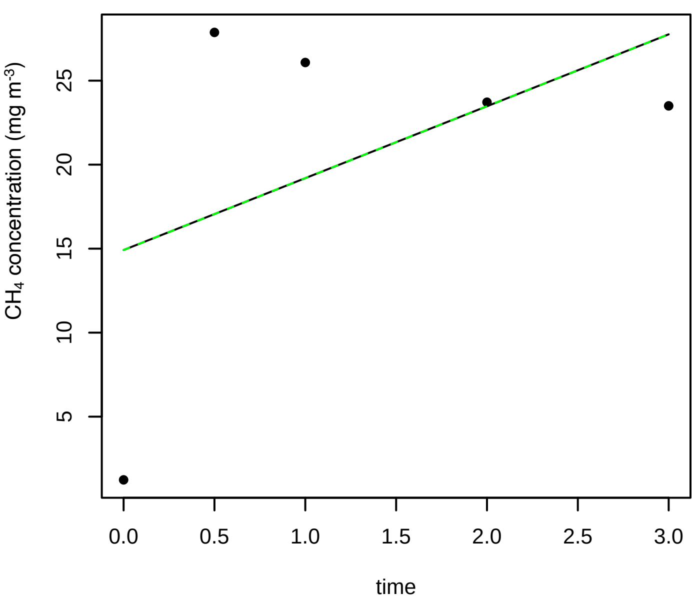


6/13/2016 2P

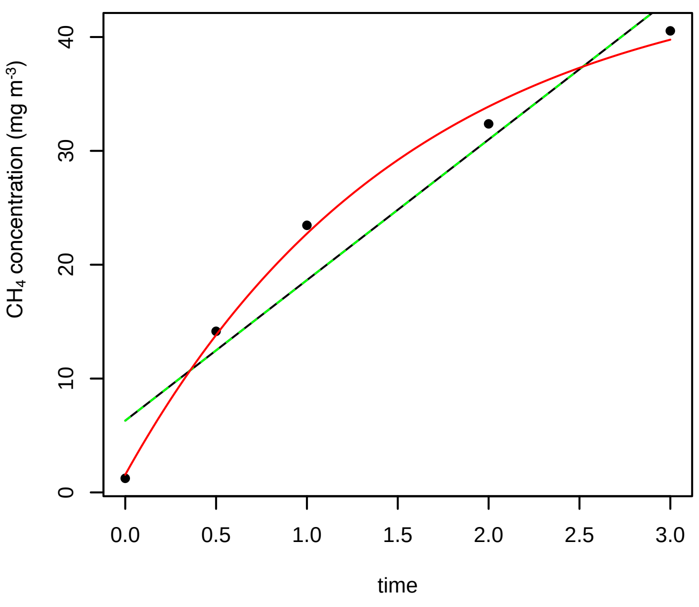


6/13/2016 3N

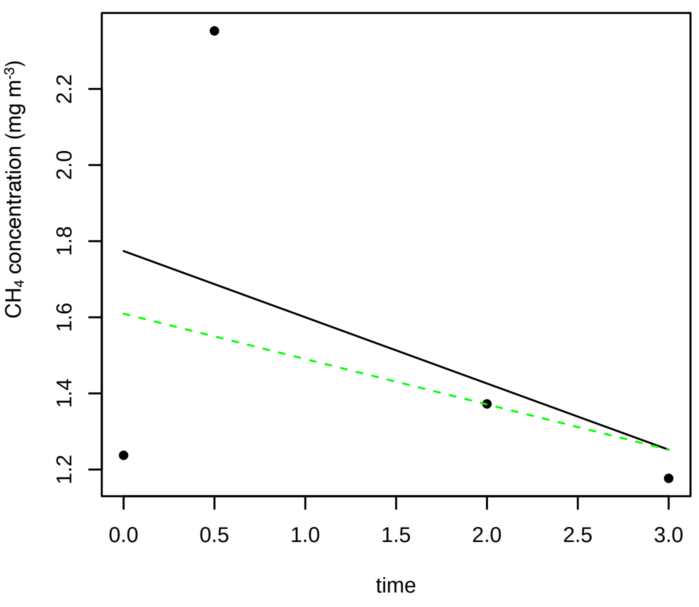

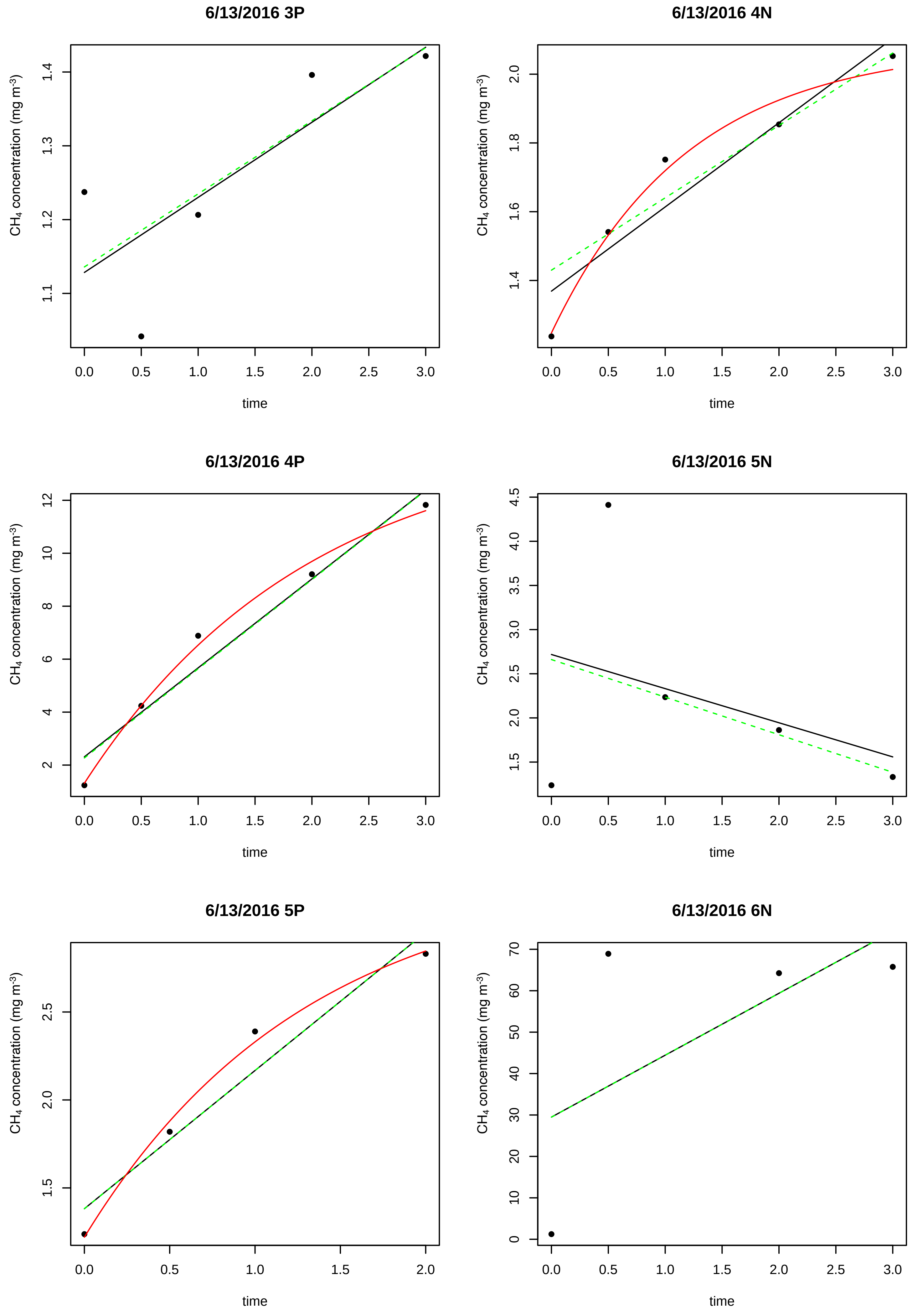
6/13/2016 3P
6/13/2016 4N
6/13/2016 4P
6/13/2016 5N
6/13/2016 5P
6/13/2016 6N
CH4 concentration (mg m-3)
time
0.0
0.5
1.0
1.5
2.0
2.5
3.0

### 6/13/2016 6P

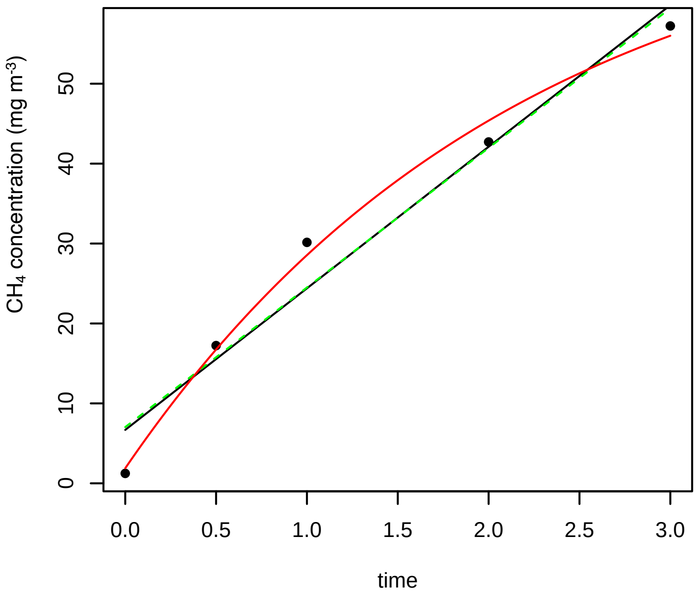


### 6/13/2016 7N

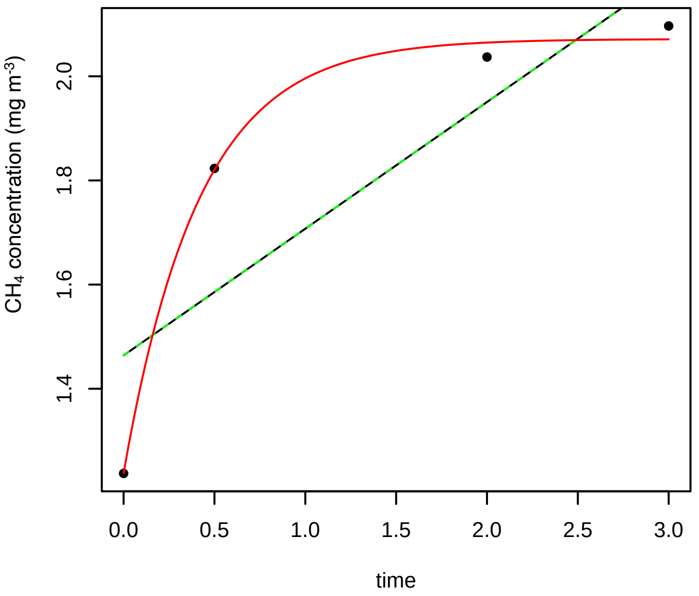


### 6/13/2016 7P

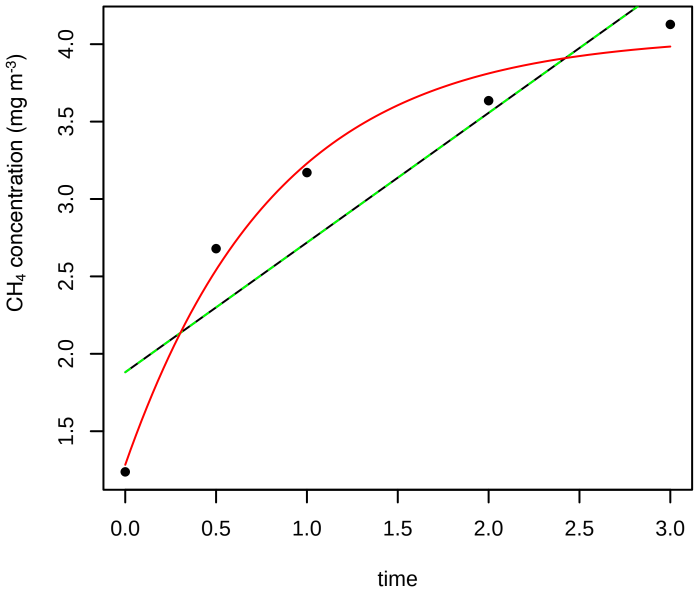


### 6/13/2016 8N

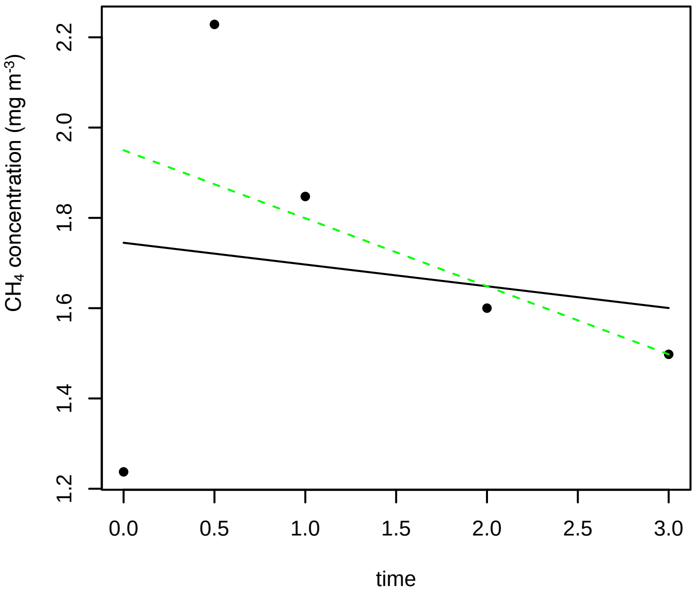


### 6/13/2016 8P

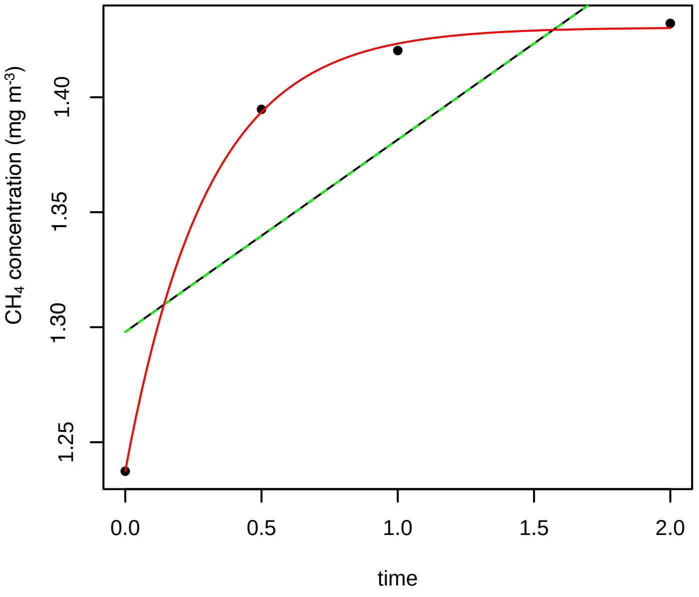


### 6/13/2016 9N

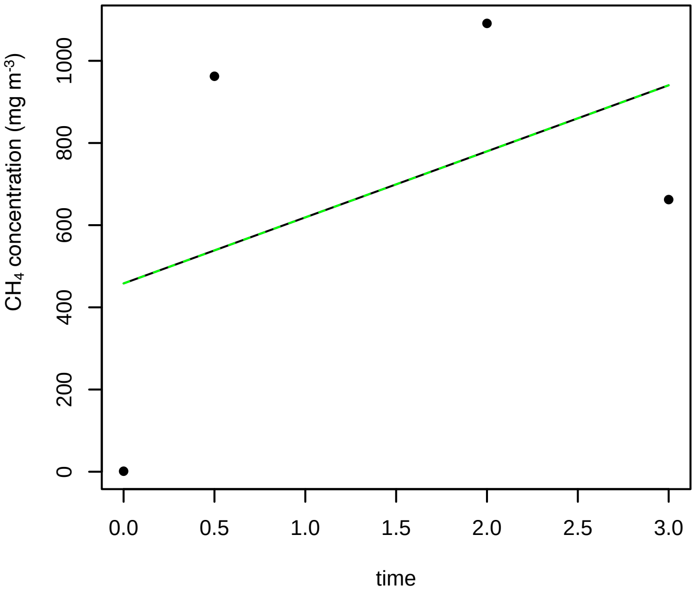

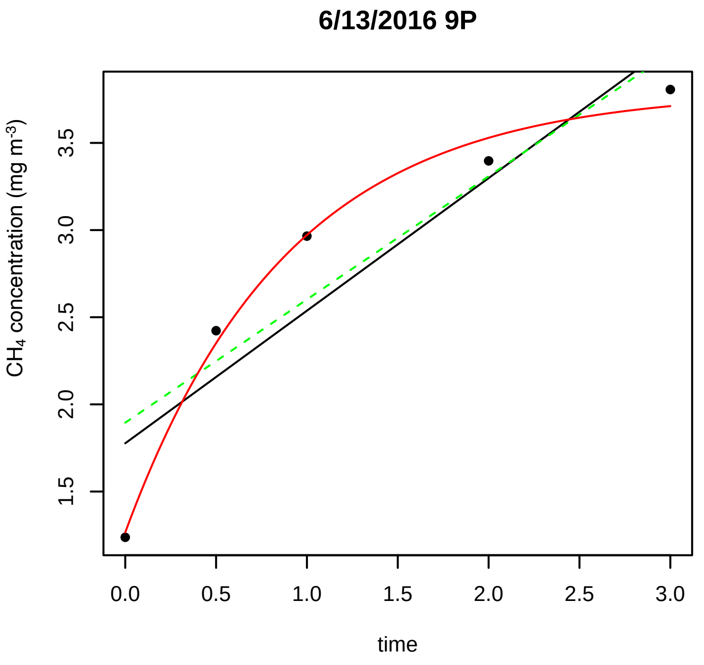
6/13/2016 9P
CH4 concentration (mg m-3)
1.5
2.0
2.5
3.0
3.5
0.0 0.5 1.0 1.5 2.0 2.5 3.0
time

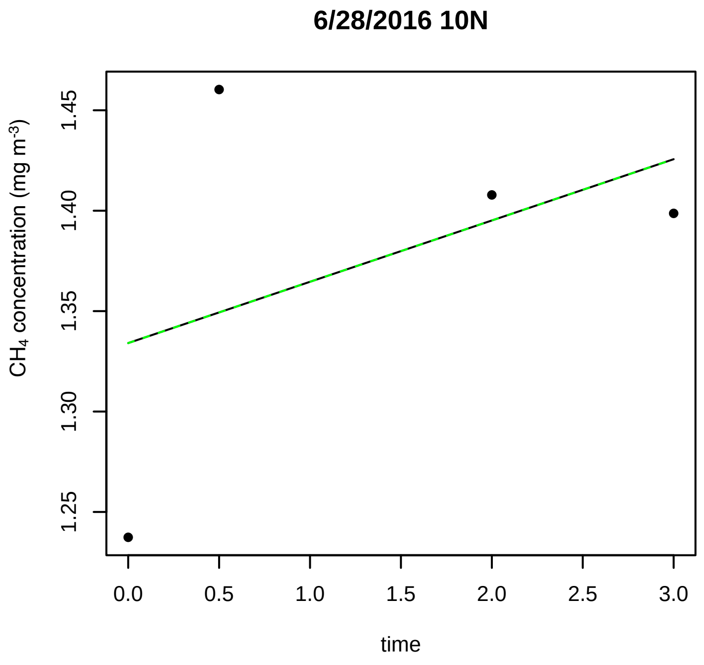
6/28/2016 10N
CH4 concentration (mg m-3)
1.25
1.30
1.35
1.40
1.45
0.0 0.5 1.0 1.5 2.0 2.5 3.0
time

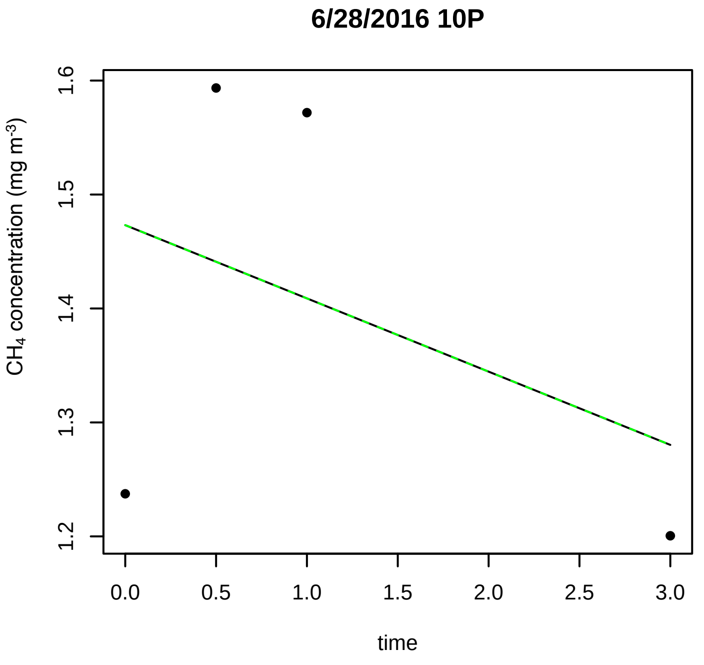
6/28/2016 10P
CH4 concentration (mg m-3)
1.2
1.3
1.4
1.5
1.6
0.0 0.5 1.0 1.5 2.0 2.5 3.0
time

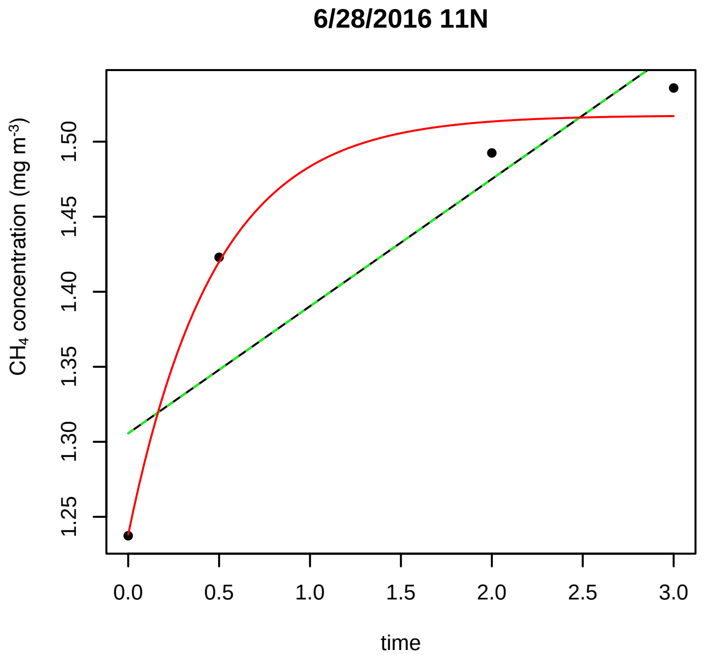
6/28/2016 11N
CH4 concentration (mg m-3)
1.25
1.30
1.35
1.40
1.45
1.50
0.0 0.5 1.0 1.5 2.0 2.5 3.0
time

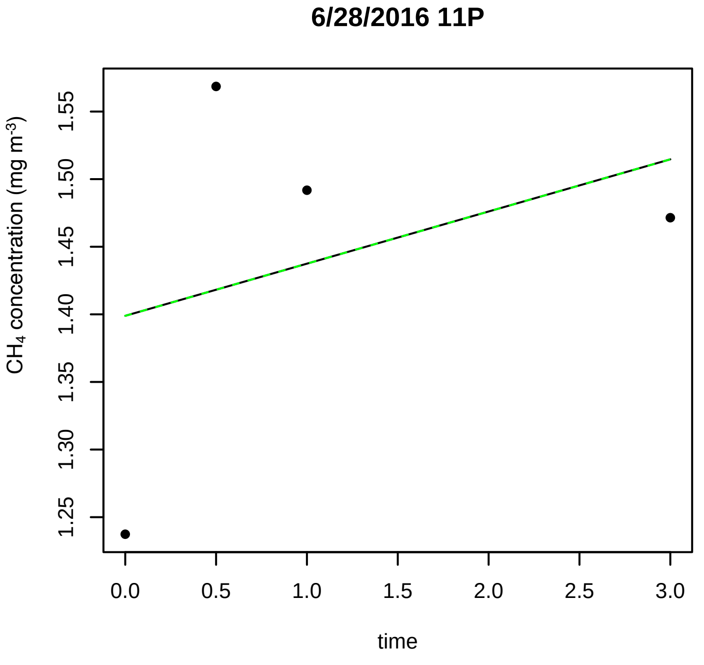
6/28/2016 11P
CH4 concentration (mg m-3)
1.25
1.30
1.35
1.40
1.45
1.50
1.55
0.0 0.5 1.0 1.5 2.0 2.5 3.0
time

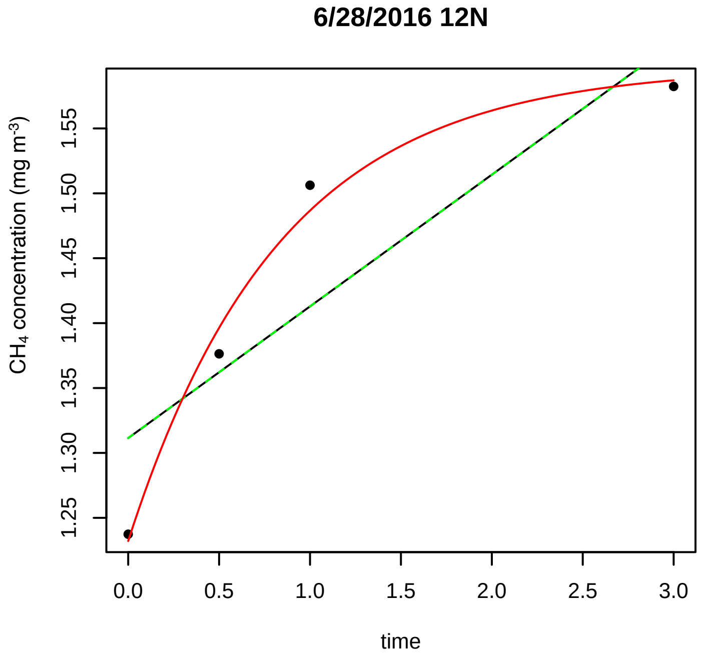
6/28/2016 12N
CH4 concentration (mg m-3)
1.25
1.30
1.35
1.40
1.45
1.50
1.55
0.0 0.5 1.0 1.5 2.0 2.5 3.0
time

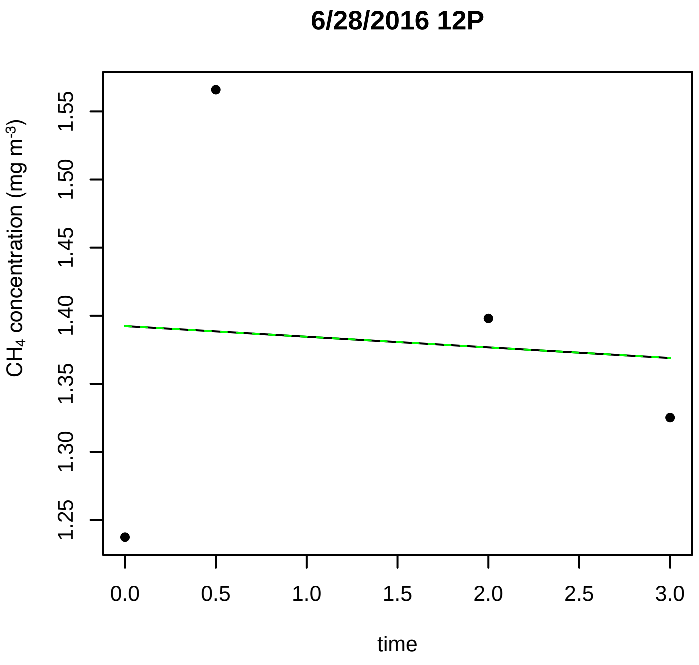
6/28/2016 12P
CH4 concentration (mg m-3)
1.25 1.30 1.35 1.40 1.45 1.50 1.55
0.0 0.5 1.0 1.5 2.0 2.5 3.0
time

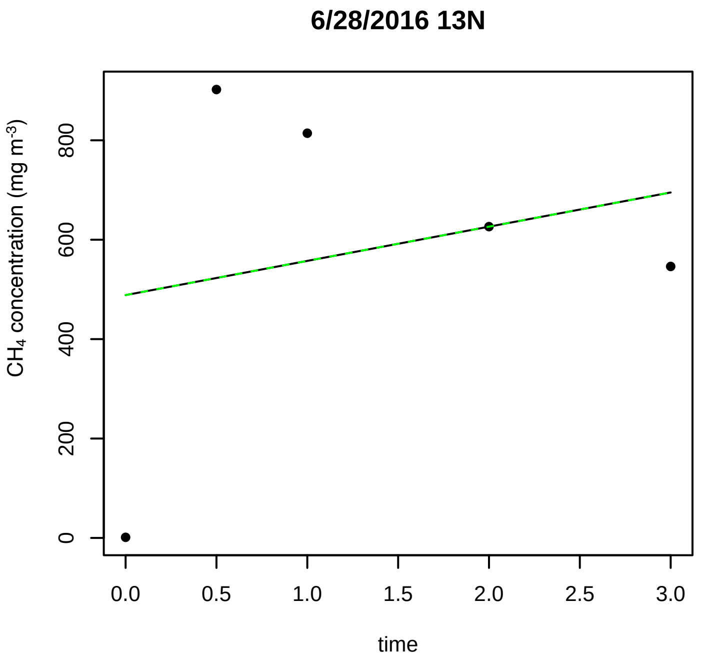
6/28/2016 13N
CH4 concentration (mg m-3)
0 200 400 600 800
0.0 0.5 1.0 1.5 2.0 2.5 3.0
time

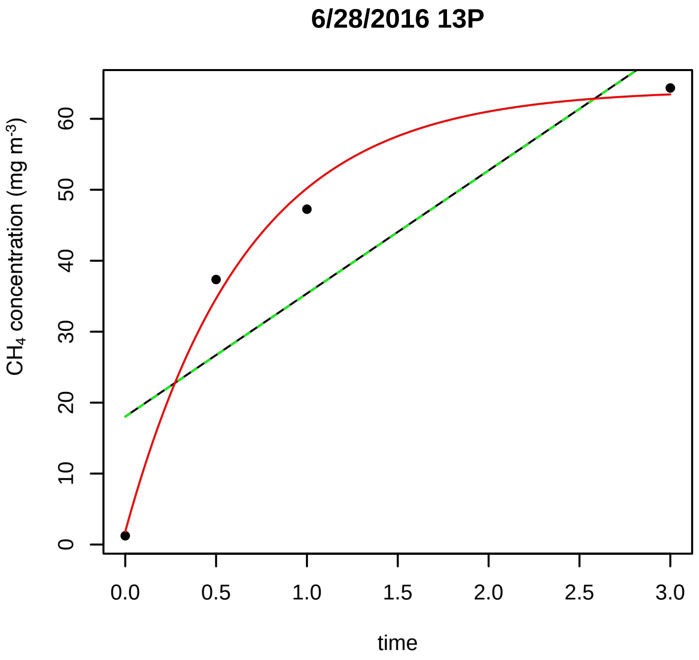
6/28/2016 13P
CH4 concentration (mg m-3)
0 10 20 30 40 50 60
0.0 0.5 1.0 1.5 2.0 2.5 3.0
time

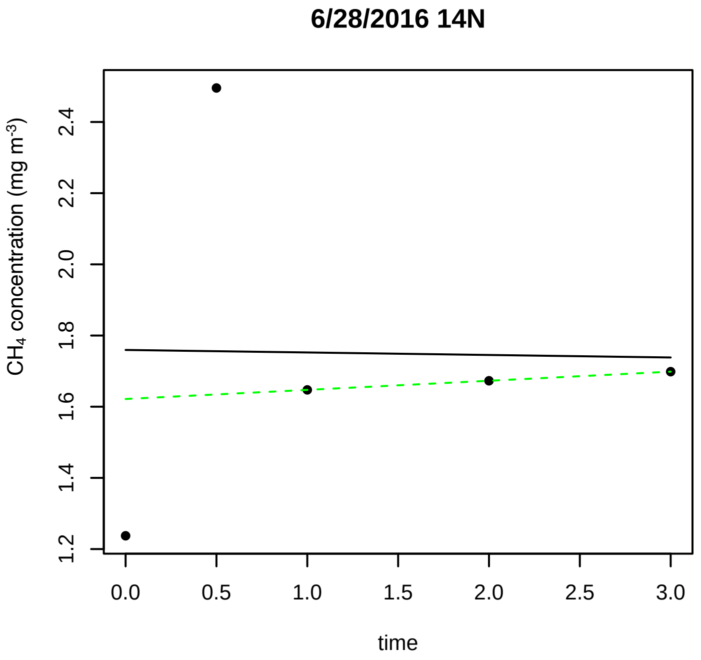
6/28/2016 14N
CH4 concentration (mg m-3)
1.2 1.4 1.6 1.8 2.0 2.2 2.4
0.0 0.5 1.0 1.5 2.0 2.5 3.0
time

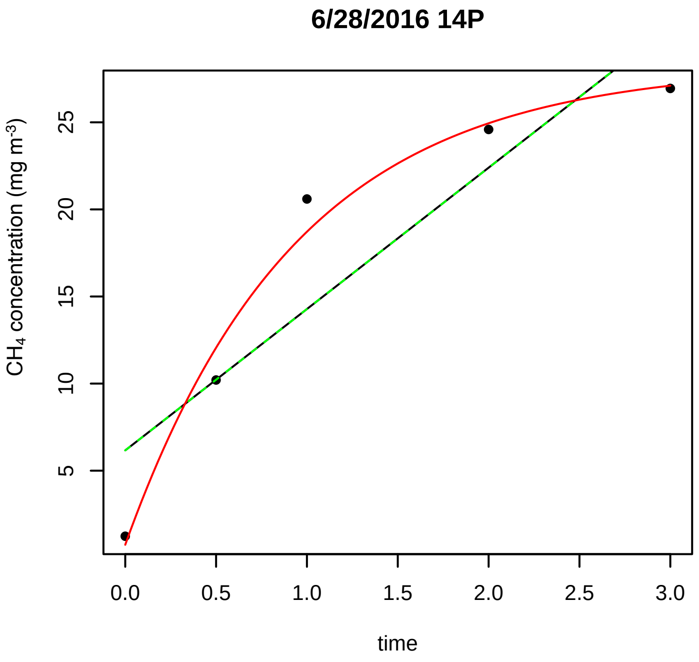
6/28/2016 14P
CH4 concentration (mg m-3)
5 10 15 20 25
0.0 0.5 1.0 1.5 2.0 2.5 3.0
time

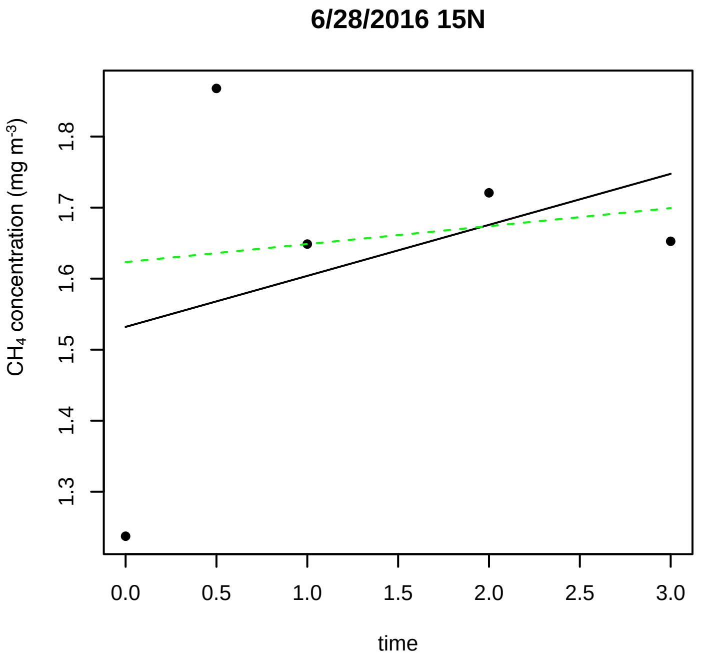
6/28/2016 15N
CH4 concentration (mg m-3)
1.3 1.4 1.5 1.6 1.7 1.8
0.0 0.5 1.0 1.5 2.0 2.5 3.0
time

**6/28/2016 15P**

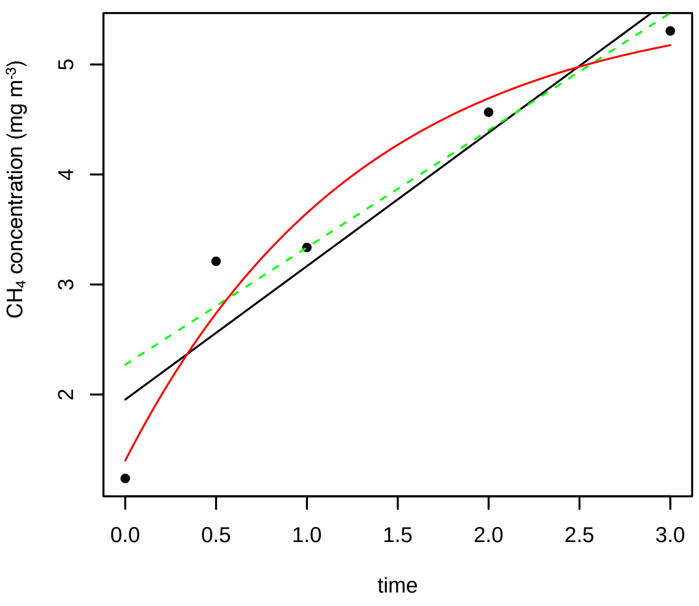


**6/28/2016 16N**

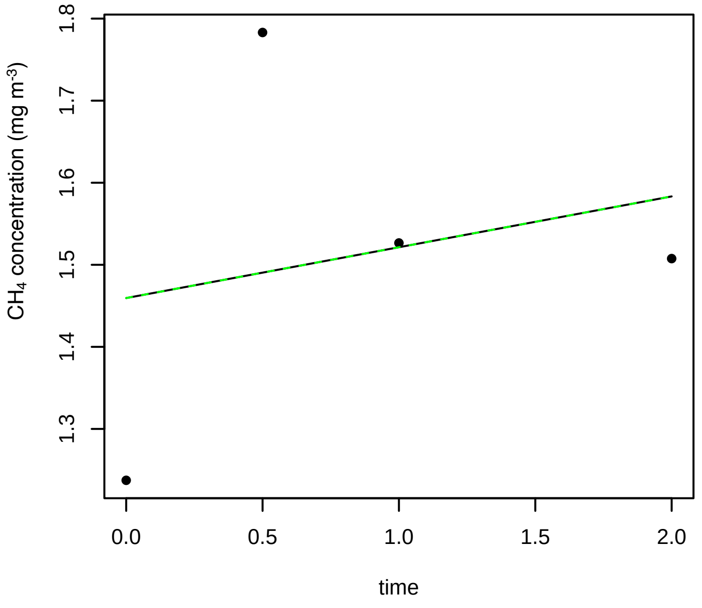


**6/28/2016 16P**

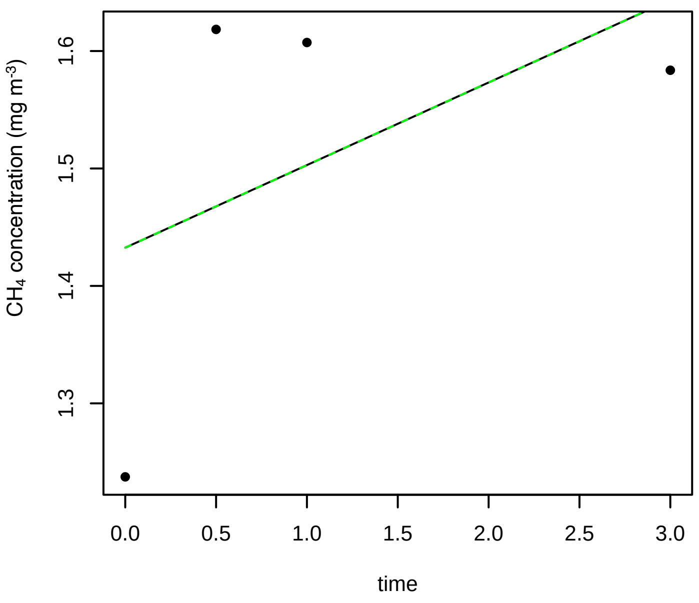


**6/28/2016 17N**

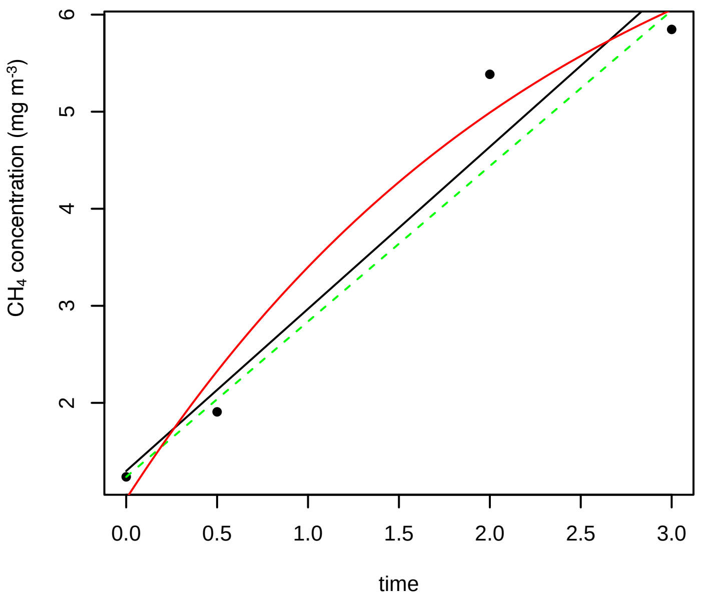


**6/28/2016 17P**

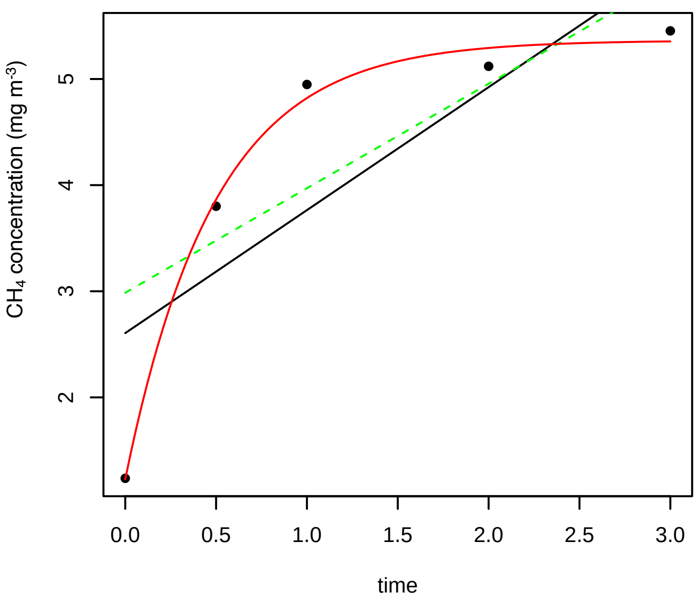


**6/28/2016 18N**

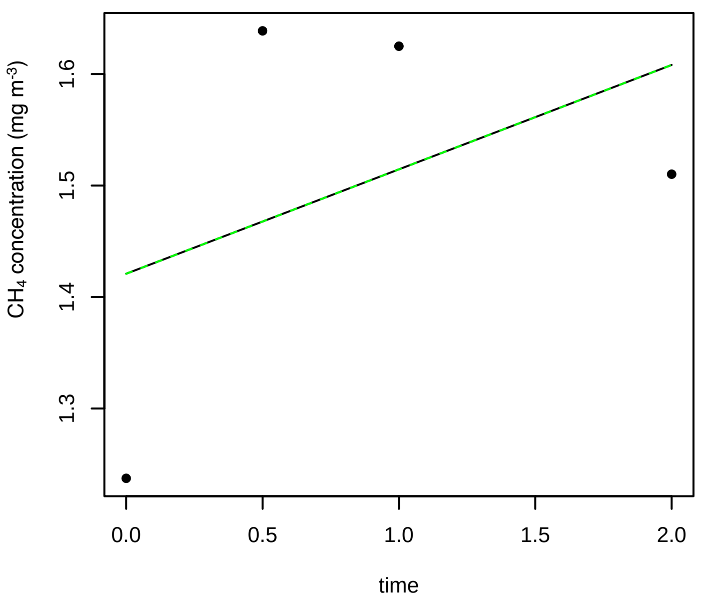

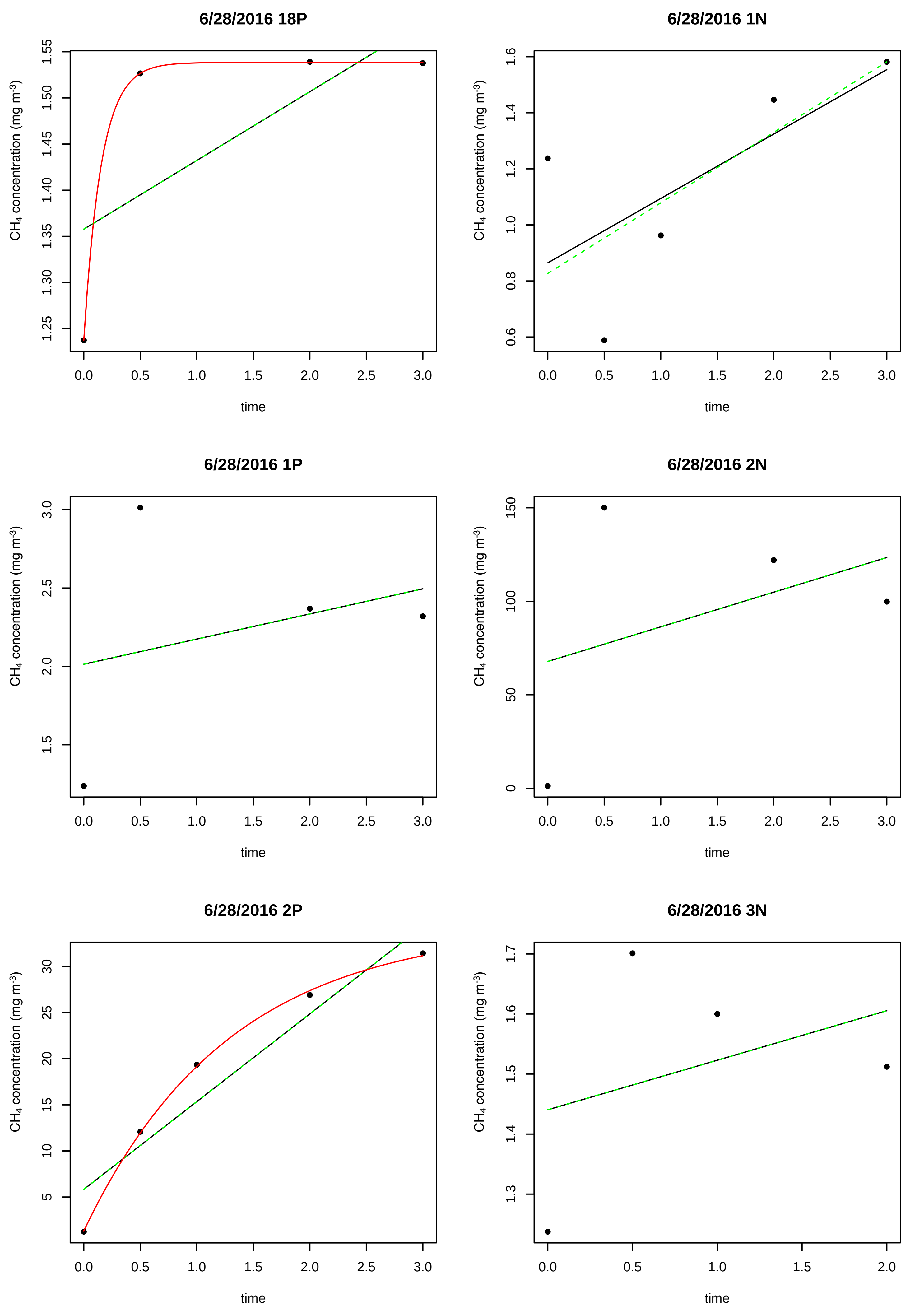

6/28/2016 18P
6/28/2016 1N
6/28/2016 1P
6/28/2016 2N
6/28/2016 2P
6/28/2016 3N
CH4 concentration (mg m-3)
time

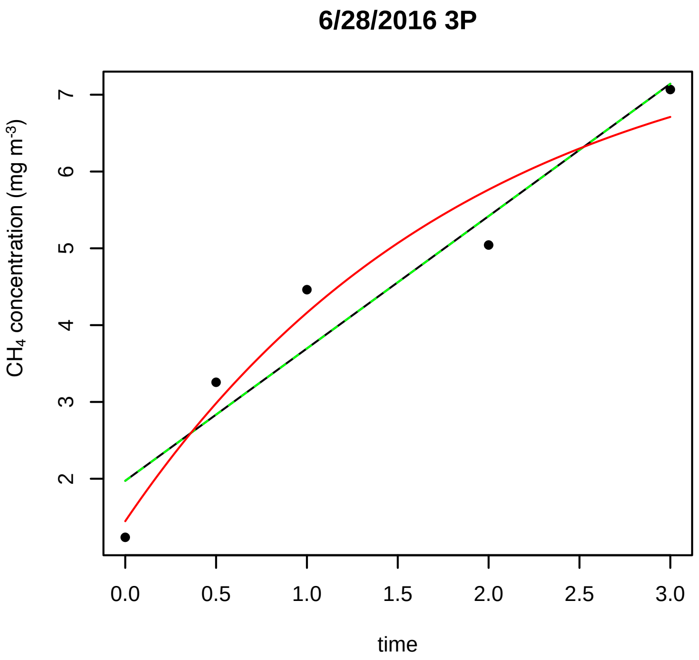
6/28/2016 3P
CH4 concentration (mg m-3)
time

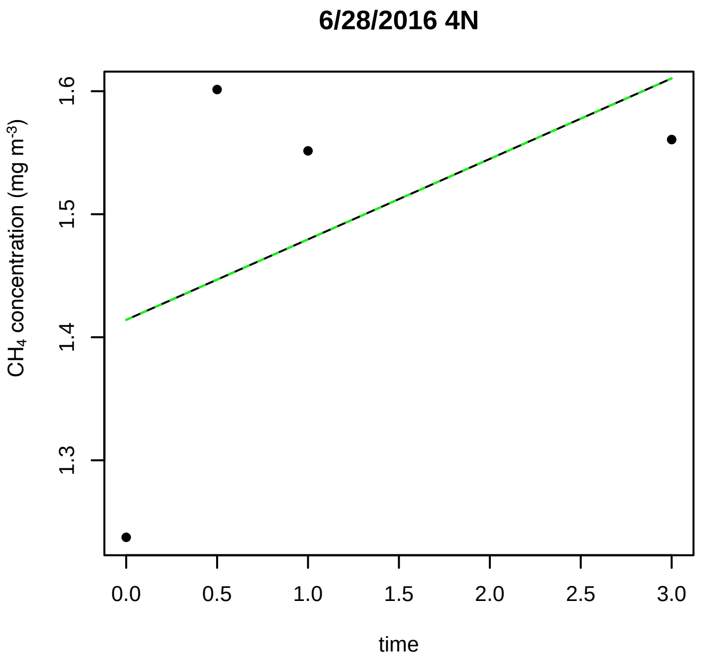
6/28/2016 4N
CH4 concentration (mg m-3)
time

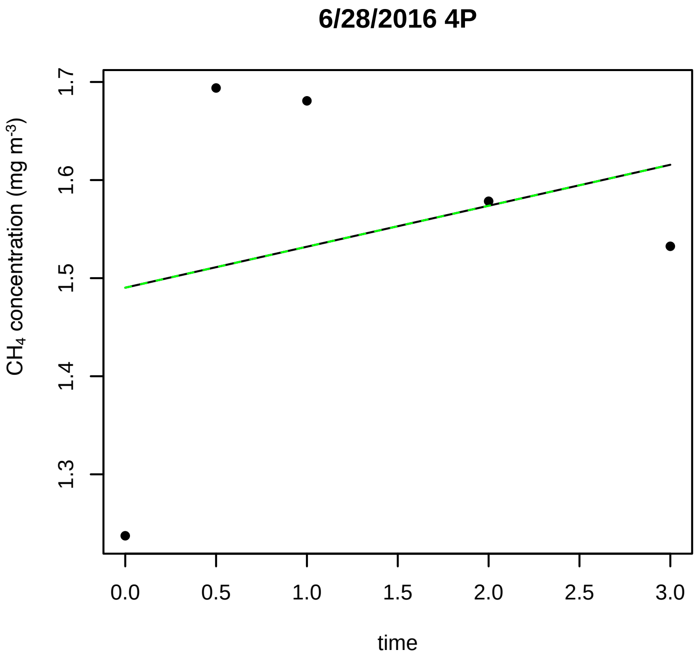
6/28/2016 4P
CH4 concentration (mg m-3)
time

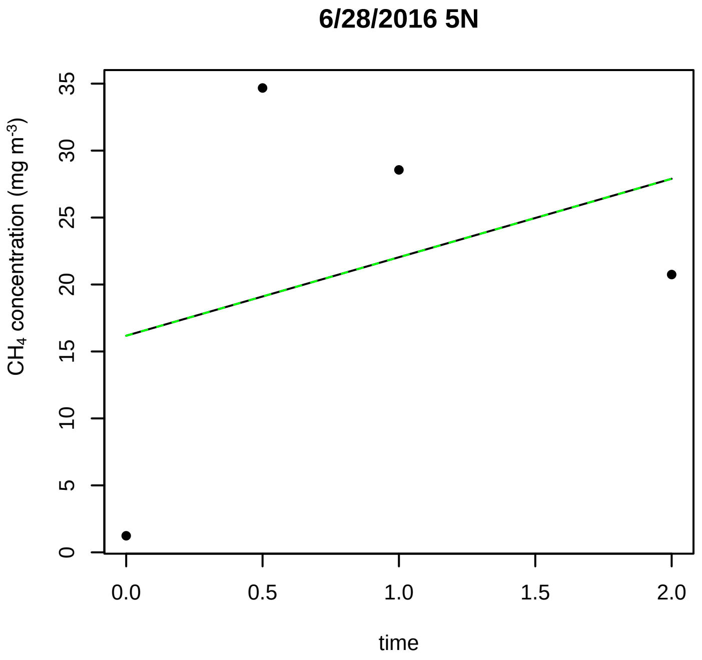
6/28/2016 5N
CH4 concentration (mg m-3)
time

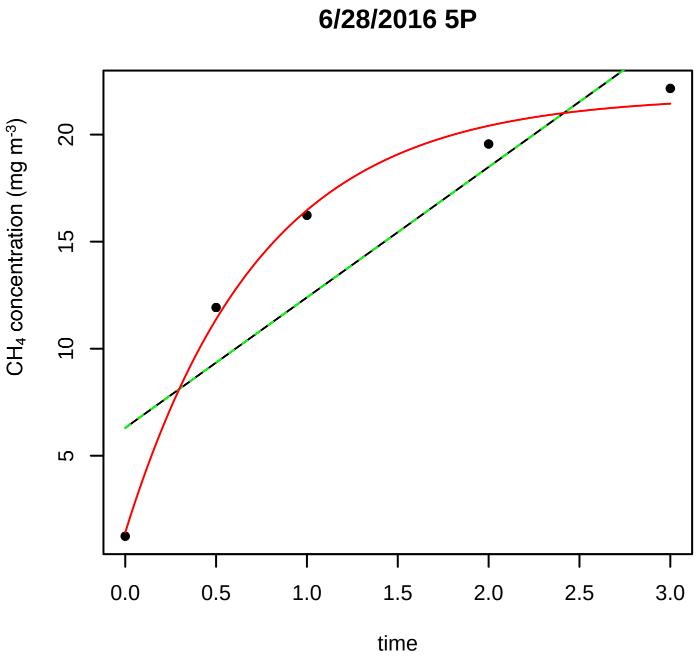
6/28/2016 5P
CH4 concentration (mg m-3)
time

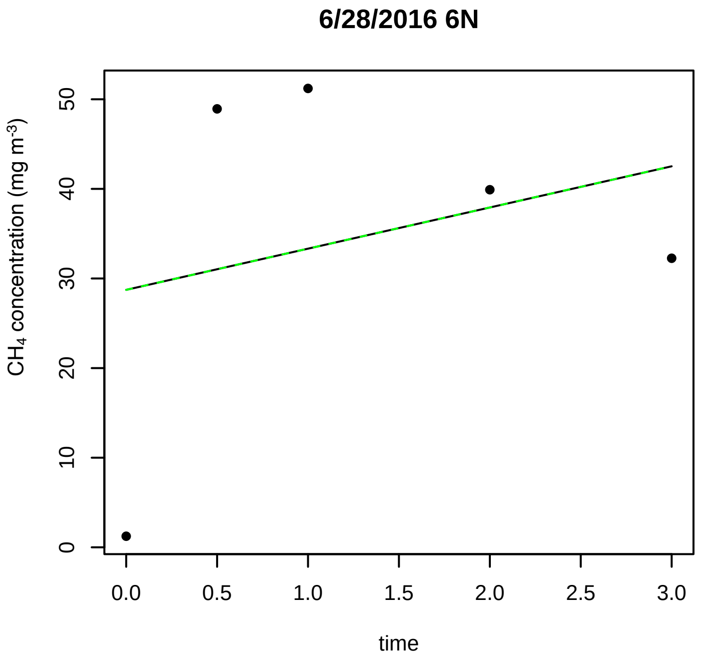
6/28/2016 6N
CH4 concentration (mg m-3)
time

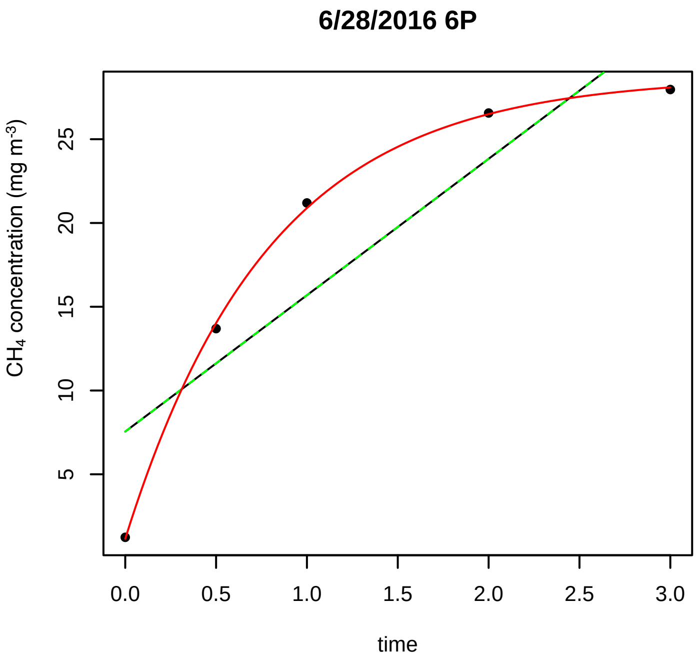
6/28/2016 6P
CH4 concentration (mg m-3)
5
10
15
20
25
0.0
0.5
1.0
1.5
2.0
2.5
3.0
time

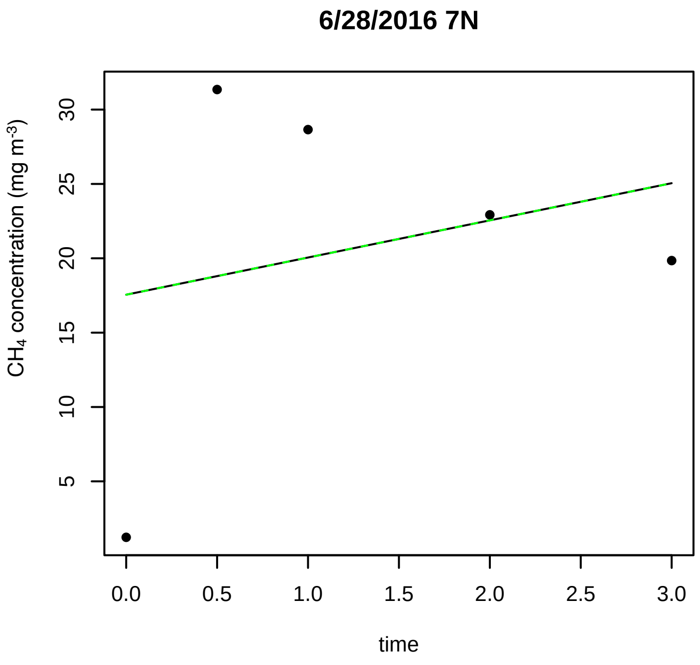
6/28/2016 7N
CH4 concentration (mg m-3)
5
10
15
20
25
30
0.0
0.5
1.0
1.5
2.0
2.5
3.0
time

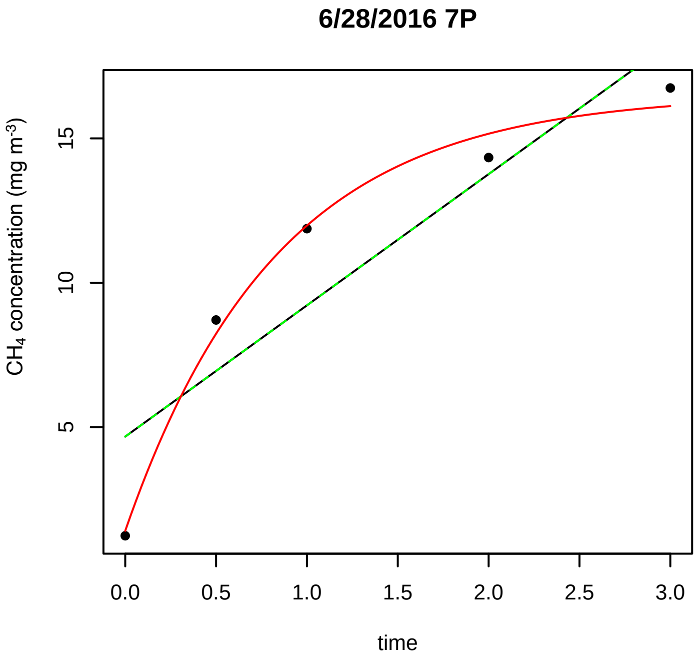
6/28/2016 7P
CH4 concentration (mg m-3)
5
10
15
0.0
0.5
1.0
1.5
2.0
2.5
3.0
time

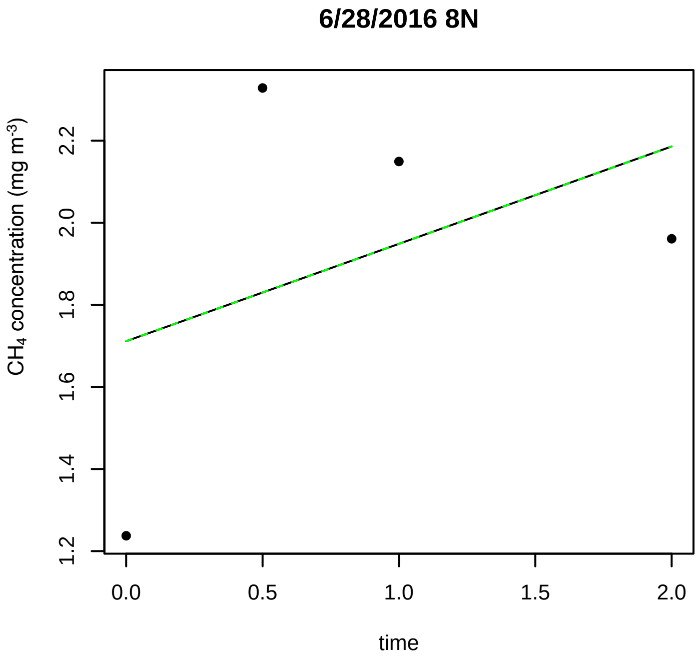
6/28/2016 8N
CH4 concentration (mg m-3)
1.2
1.4
1.6
1.8
2.0
2.2
0.0
0.5
1.0
1.5
2.0
time

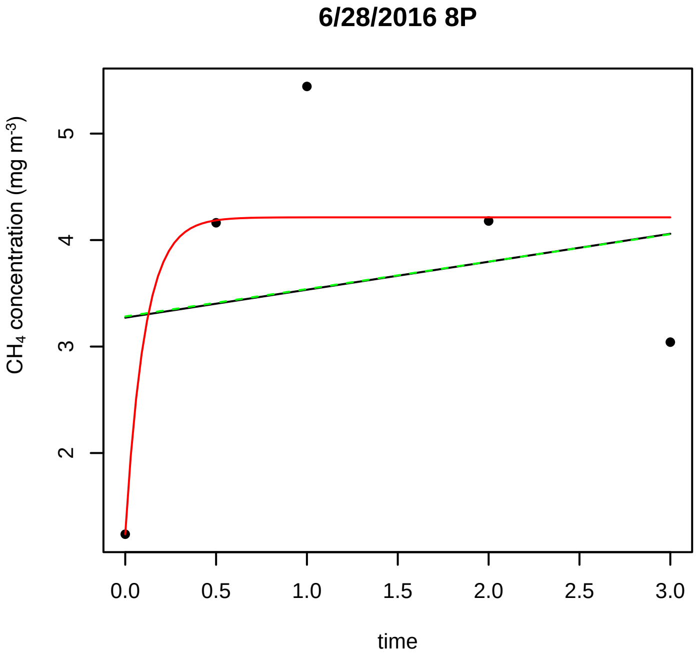
6/28/2016 8P
CH4 concentration (mg m-3)
2
3
4
5
0.0
0.5
1.0
1.5
2.0
2.5
3.0
time

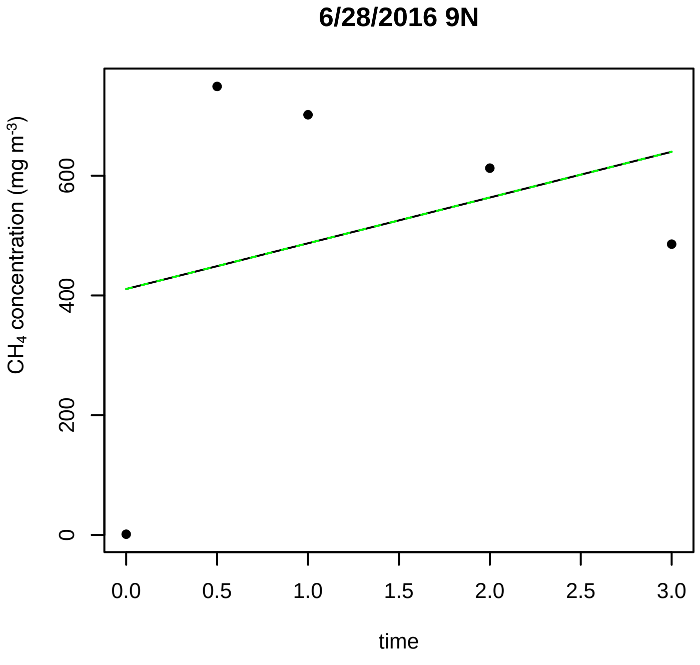
6/28/2016 9N
CH4 concentration (mg m-3)
0
200
400
600
0.0
0.5
1.0
1.5
2.0
2.5
3.0
time

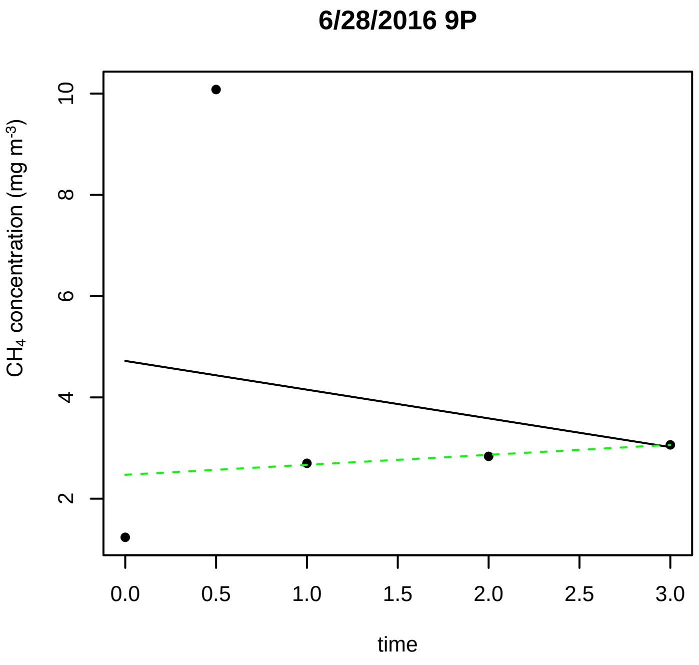
6/28/2016 9P
$CH_4$ concentration (mg m$^{-3}$)
2 4 6 8 10
0.0 0.5 1.0 1.5 2.0 2.5 3.0
time

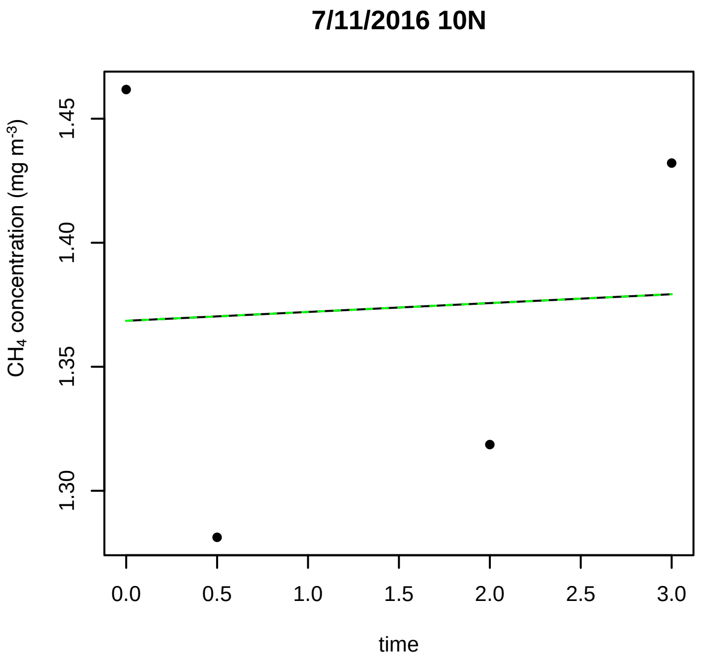
7/11/2016 10N
$CH_4$ concentration (mg m$^{-3}$)
1.30 1.35 1.40 1.45
0.0 0.5 1.0 1.5 2.0 2.5 3.0
time

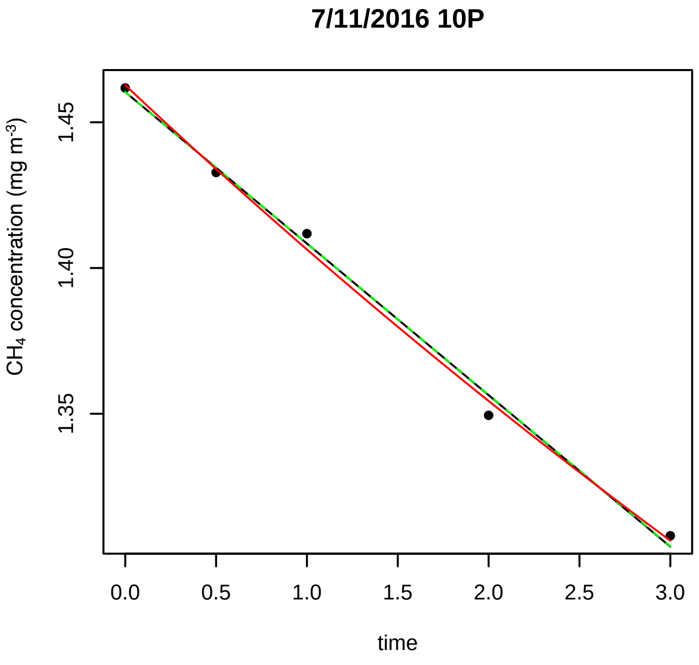
7/11/2016 10P
$CH_4$ concentration (mg m$^{-3}$)
1.35 1.40 1.45
0.0 0.5 1.0 1.5 2.0 2.5 3.0
time

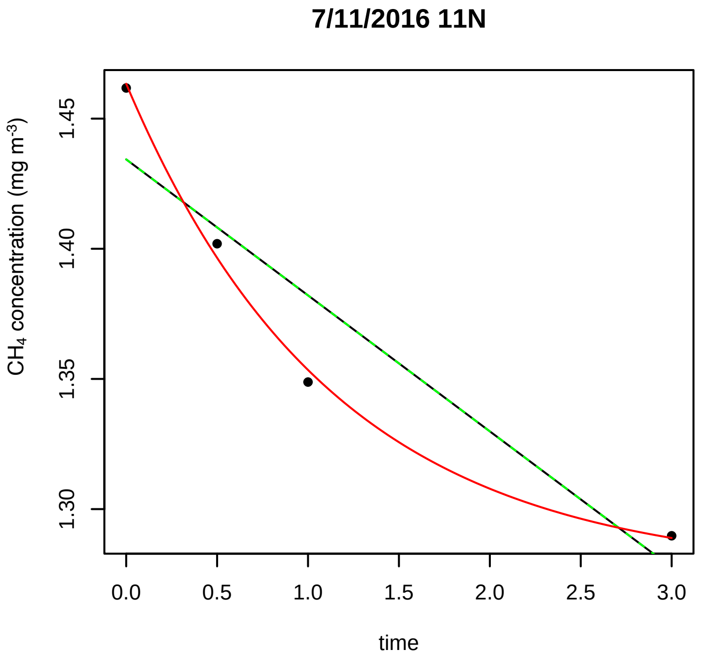
7/11/2016 11N
$CH_4$ concentration (mg m$^{-3}$)
1.30 1.35 1.40 1.45
0.0 0.5 1.0 1.5 2.0 2.5 3.0
time

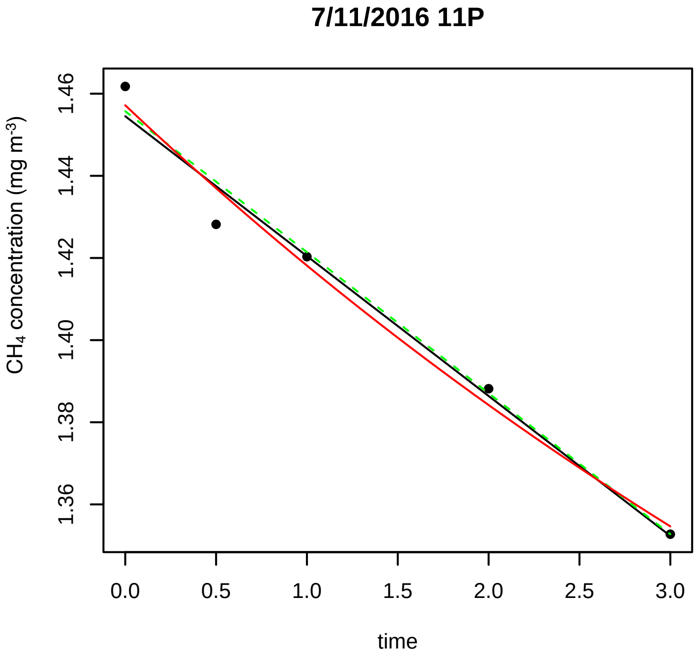
7/11/2016 11P
$CH_4$ concentration (mg m$^{-3}$)
1.36 1.38 1.40 1.42 1.44 1.46
0.0 0.5 1.0 1.5 2.0 2.5 3.0
time

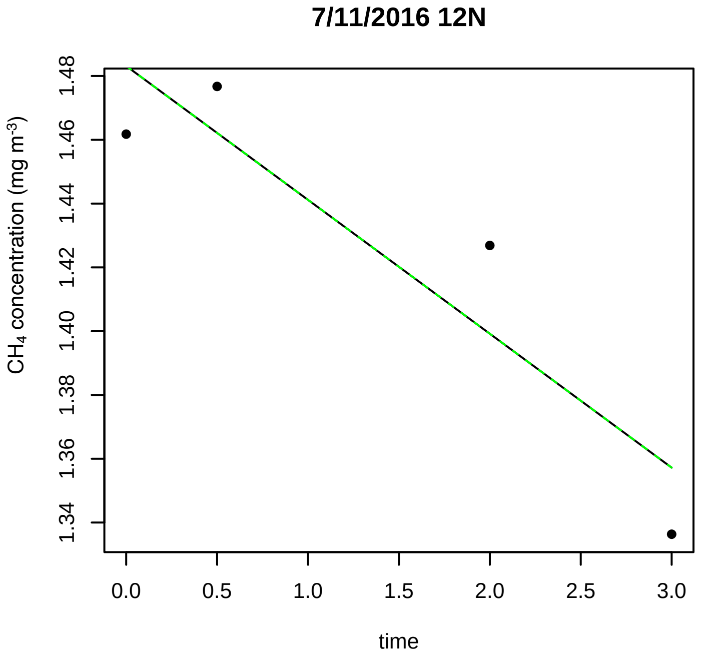
7/11/2016 12N
$CH_4$ concentration (mg m$^{-3}$)
1.34 1.36 1.38 1.40 1.42 1.44 1.46 1.48
0.0 0.5 1.0 1.5 2.0 2.5 3.0
time

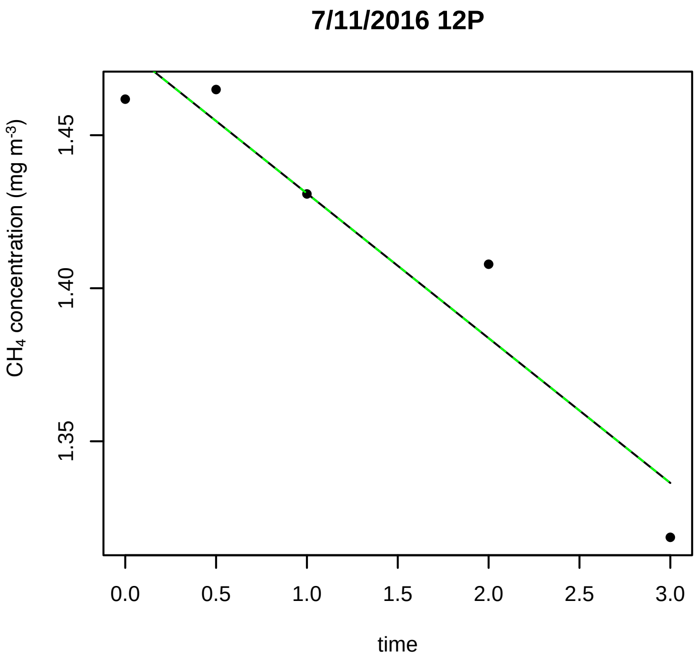
7/11/2016 12P
CH4 concentration (mg m-3)
1.35
1.40
1.45
0.0 0.5 1.0 1.5 2.0 2.5 3.0
time

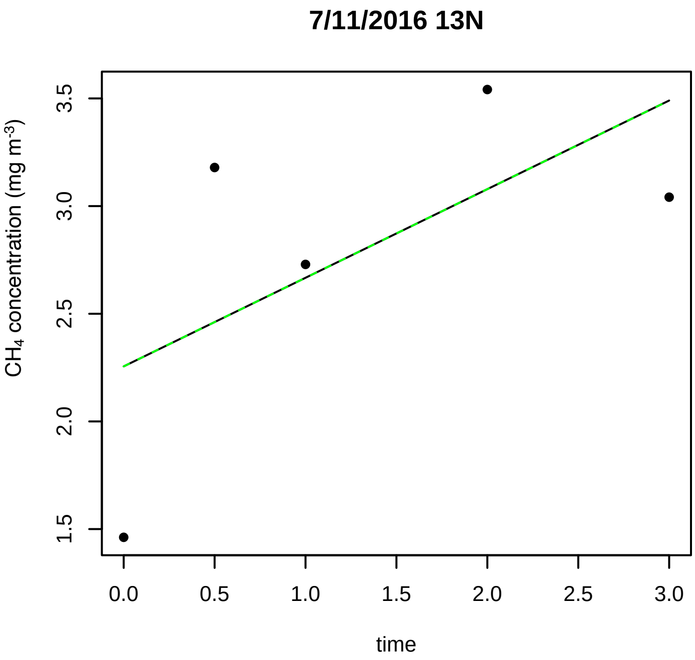
7/11/2016 13N
CH4 concentration (mg m-3)
1.5
2.0
2.5
3.0
3.5
0.0 0.5 1.0 1.5 2.0 2.5 3.0
time

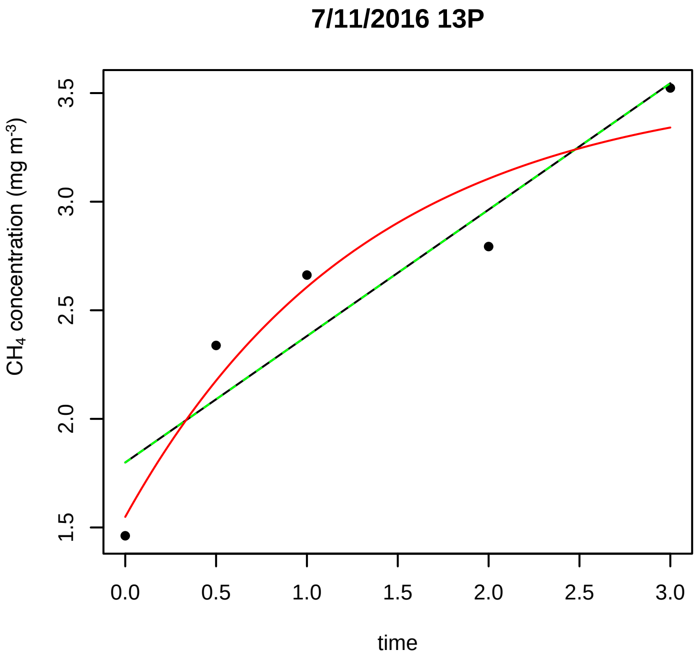
7/11/2016 13P
CH4 concentration (mg m-3)
1.5
2.0
2.5
3.0
3.5
0.0 0.5 1.0 1.5 2.0 2.5 3.0
time

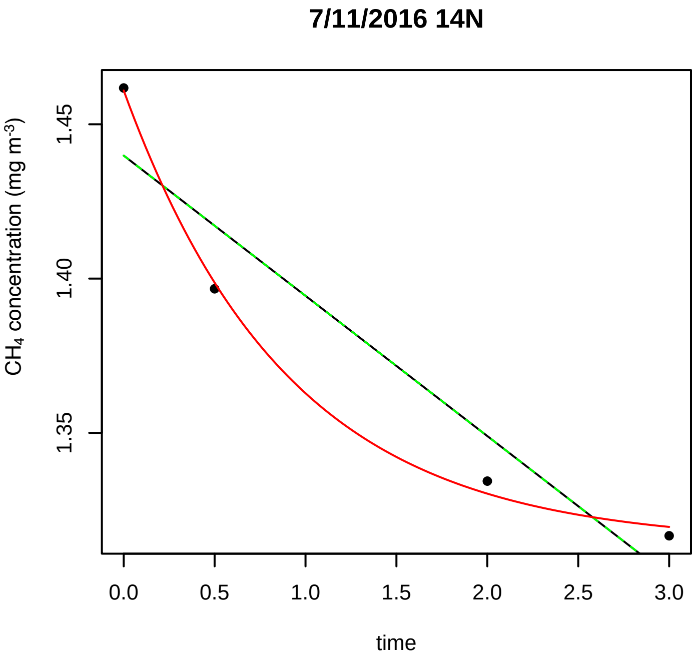
7/11/2016 14N
CH4 concentration (mg m-3)
1.35
1.40
1.45
0.0 0.5 1.0 1.5 2.0 2.5 3.0
time

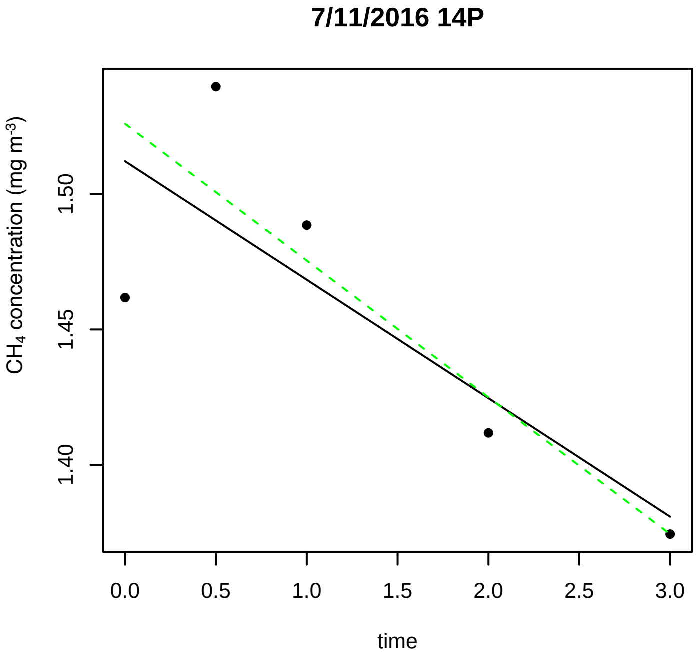
7/11/2016 14P
CH4 concentration (mg m-3)
1.40
1.45
1.50
0.0 0.5 1.0 1.5 2.0 2.5 3.0
time

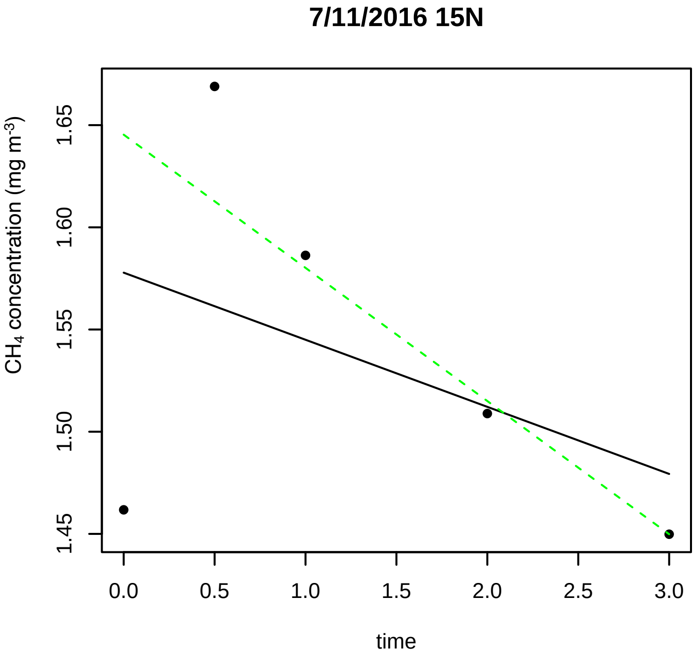
7/11/2016 15N
CH4 concentration (mg m-3)
1.45
1.50
1.55
1.60
1.65
0.0 0.5 1.0 1.5 2.0 2.5 3.0
time

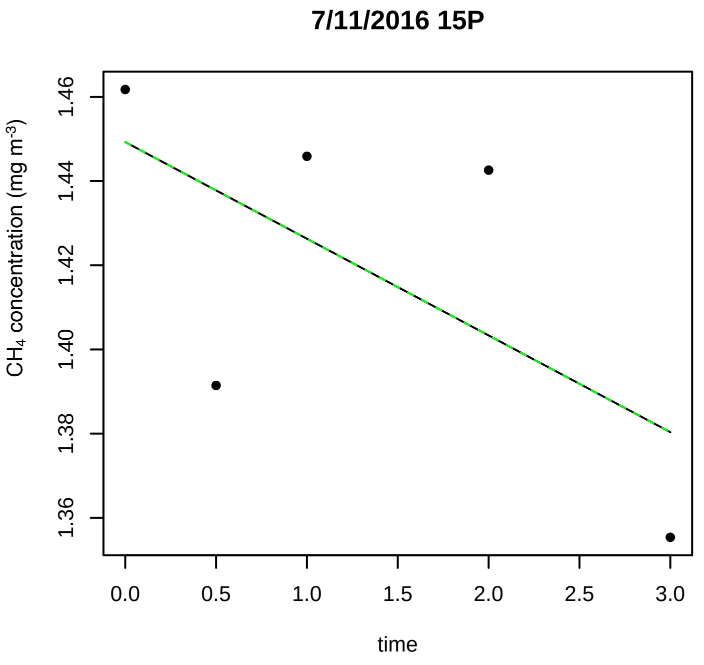
7/11/2016 15P
CH4 concentration (mg m-3)
1.36 1.38 1.40 1.42 1.44 1.46
0.0 0.5 1.0 1.5 2.0 2.5 3.0
time

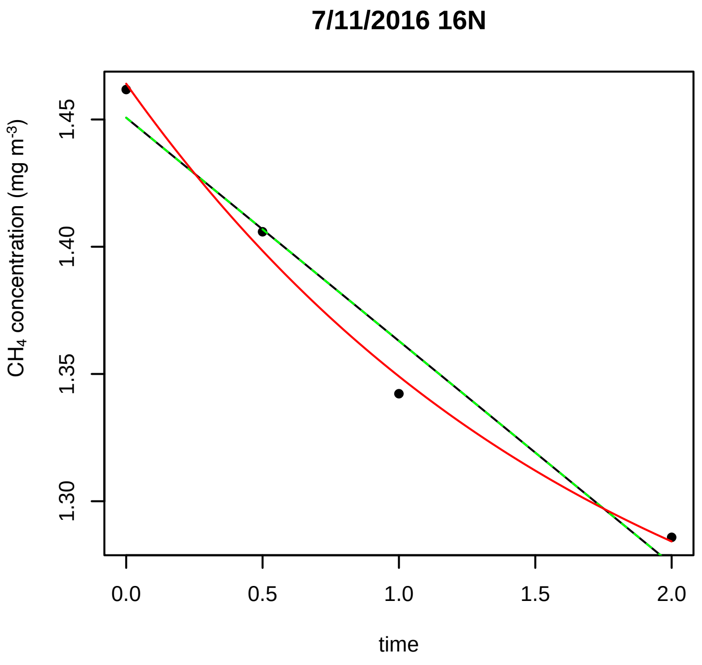
7/11/2016 16N
CH4 concentration (mg m-3)
1.30 1.35 1.40 1.45
0.0 0.5 1.0 1.5 2.0
time

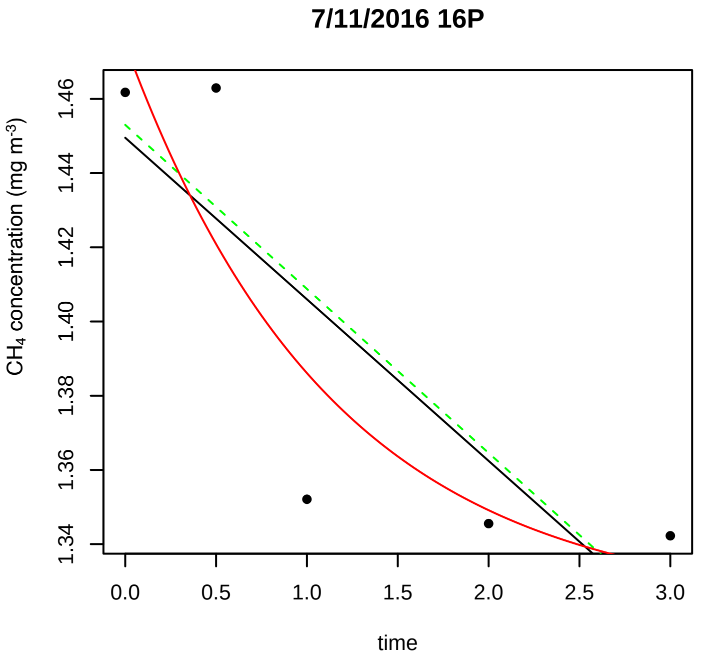
7/11/2016 16P
CH4 concentration (mg m-3)
1.34 1.36 1.38 1.40 1.42 1.44 1.46
0.0 0.5 1.0 1.5 2.0 2.5 3.0
time

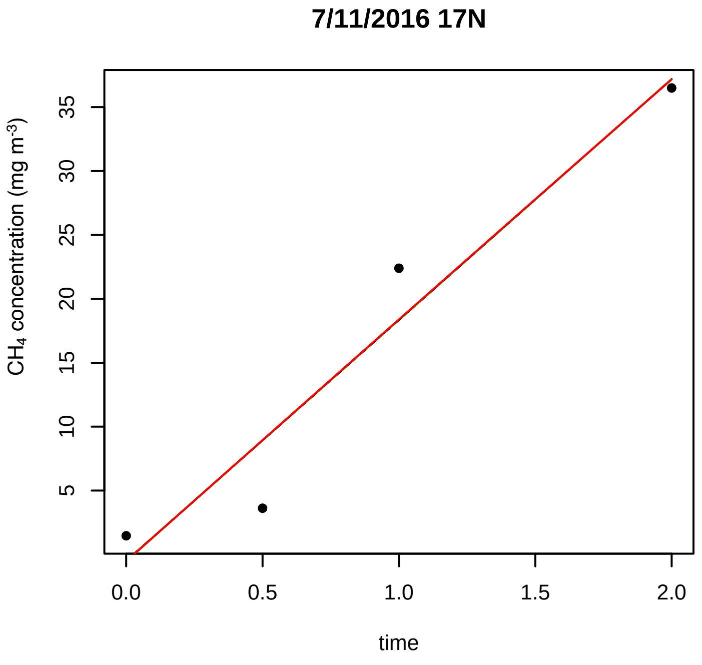
7/11/2016 17N
CH4 concentration (mg m-3)
5 10 15 20 25 30 35
0.0 0.5 1.0 1.5 2.0
time

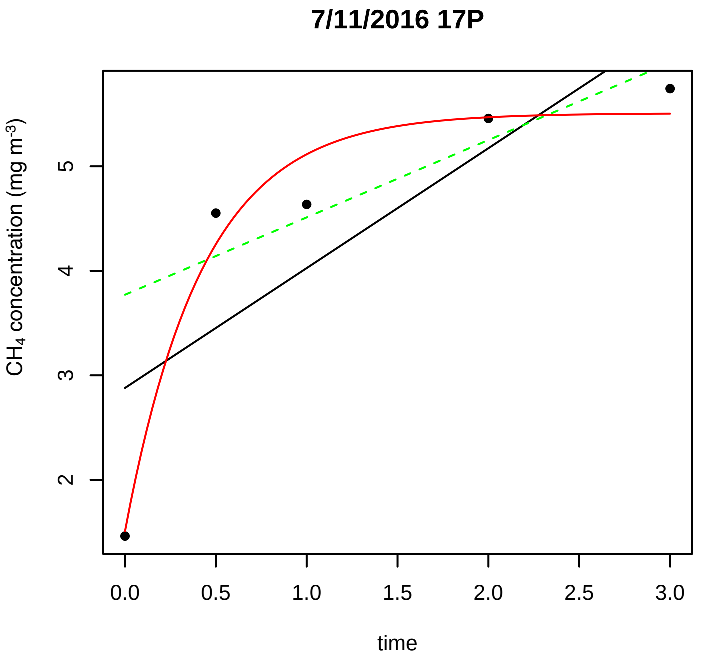
7/11/2016 17P
CH4 concentration (mg m-3)
2 3 4 5
0.0 0.5 1.0 1.5 2.0 2.5 3.0
time

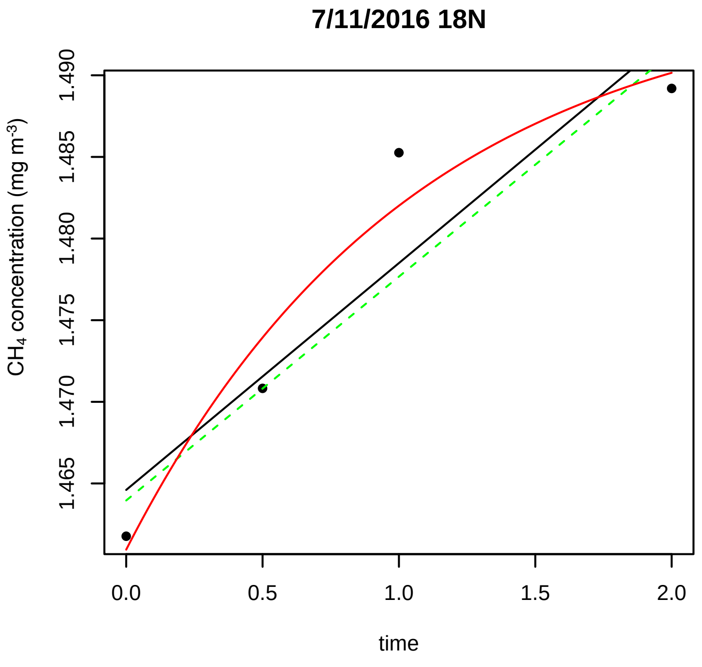
7/11/2016 18N
CH4 concentration (mg m-3)
1.465 1.470 1.475 1.480 1.485 1.490
0.0 0.5 1.0 1.5 2.0
time

7/11/2016 18P

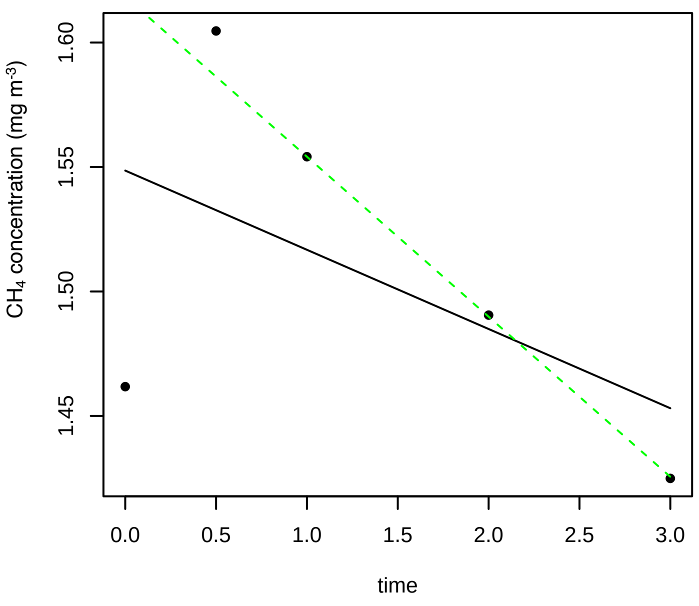


7/11/2016 1N

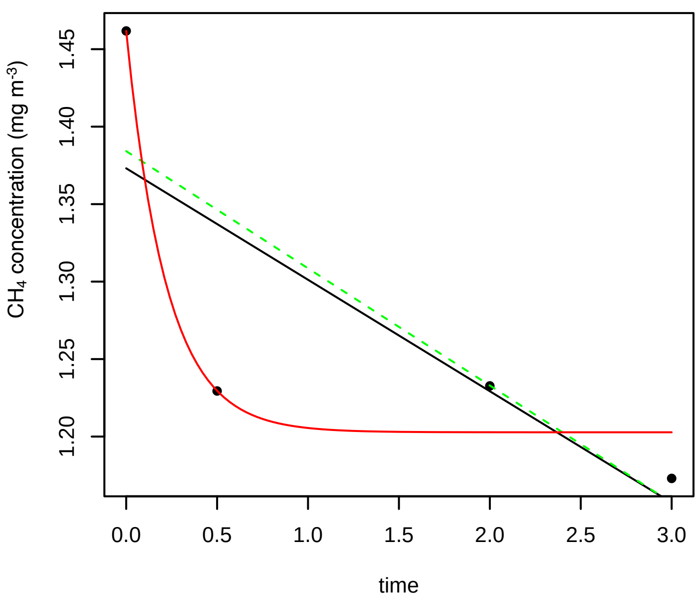


7/11/2016 1P

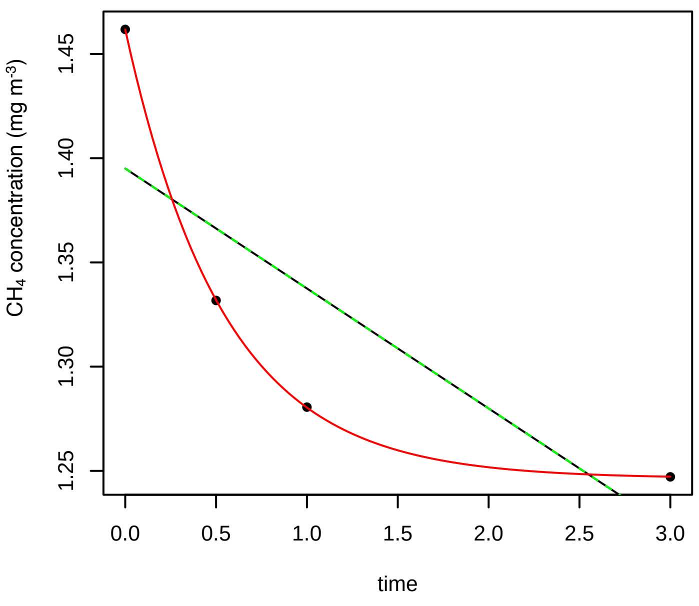


7/11/2016 2N

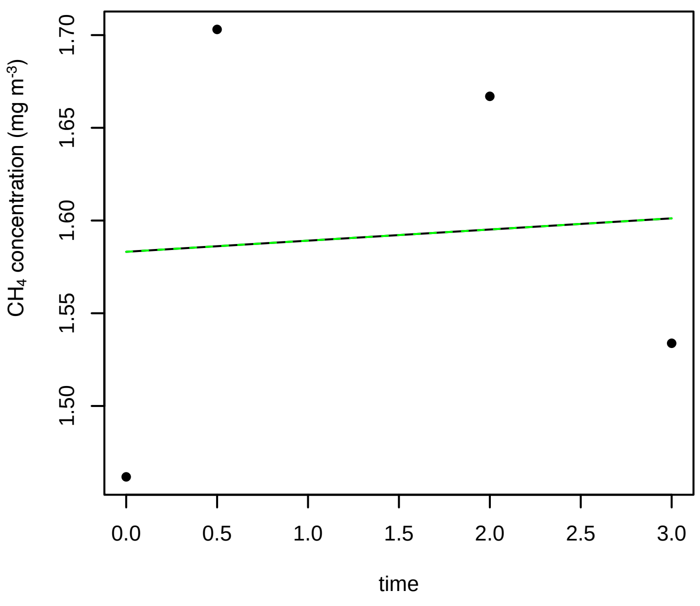


7/11/2016 2P

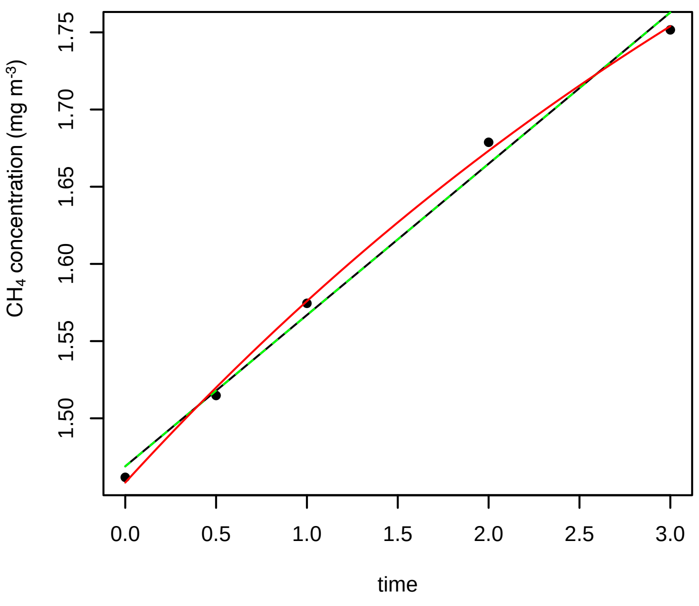


7/11/2016 3N

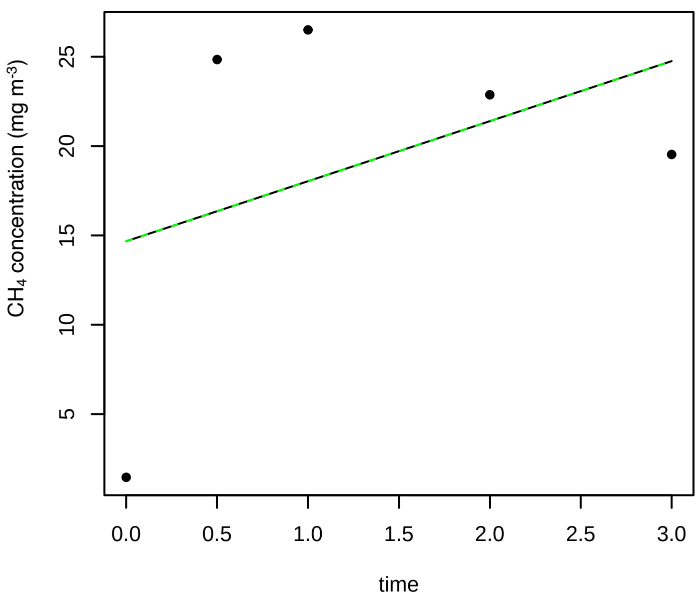

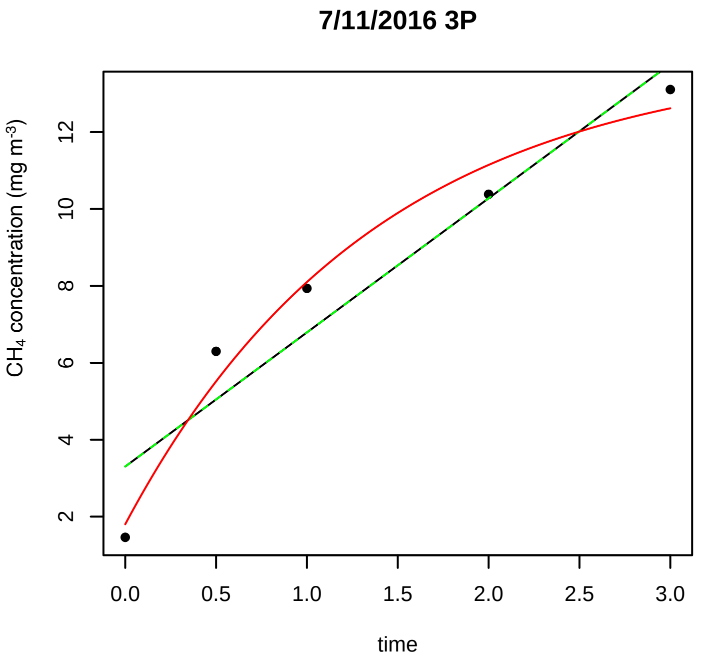
7/11/2016 3P
CH4 concentration (mg m-3)
time

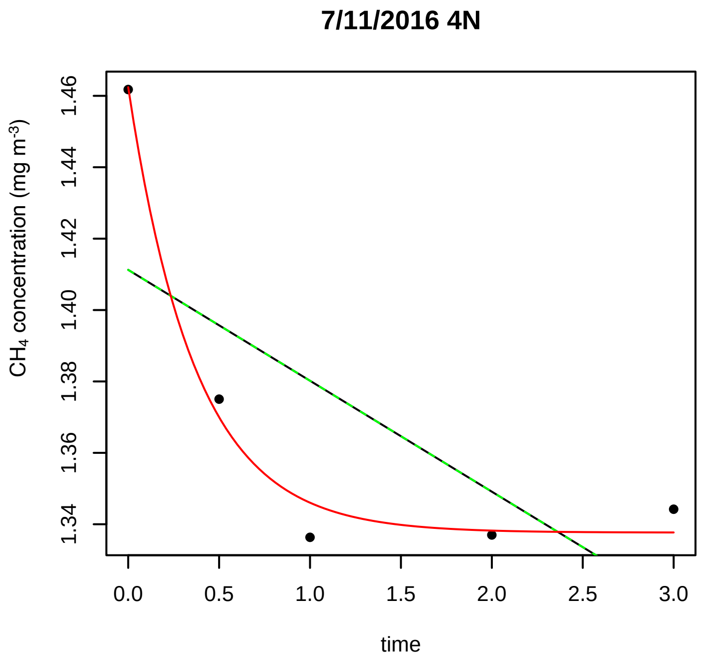
7/11/2016 4N
CH4 concentration (mg m-3)
time

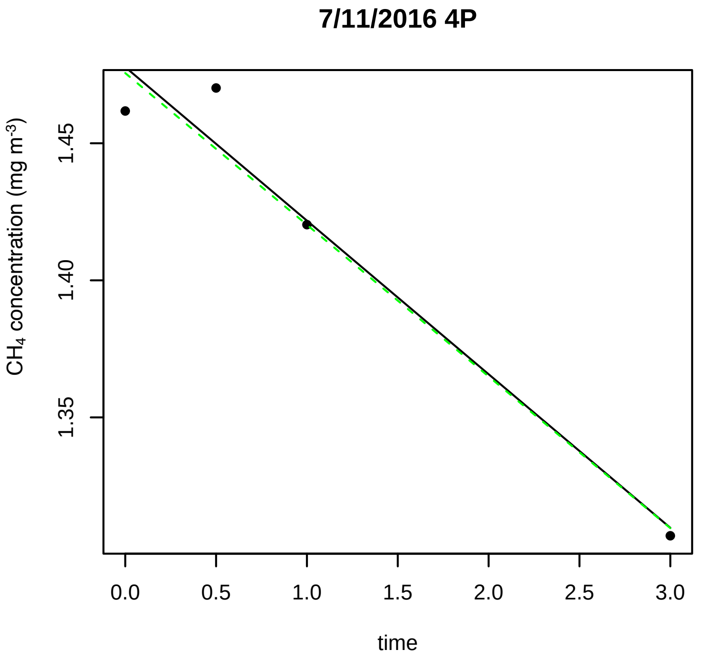
7/11/2016 4P
CH4 concentration (mg m-3)
time

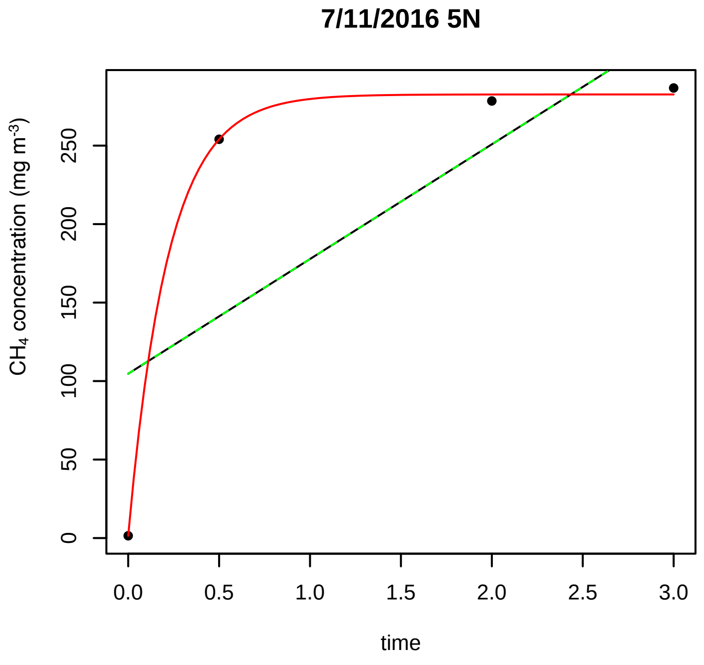
7/11/2016 5N
CH4 concentration (mg m-3)
time

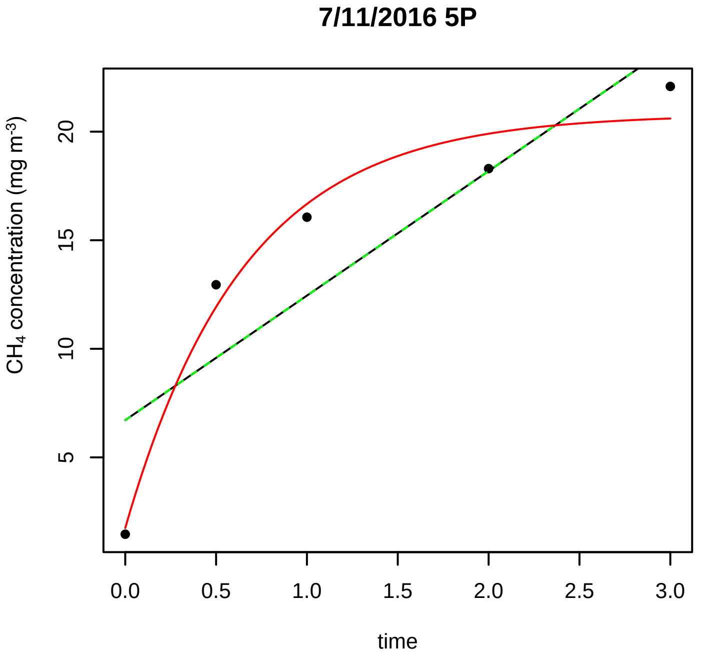
7/11/2016 5P
CH4 concentration (mg m-3)
time

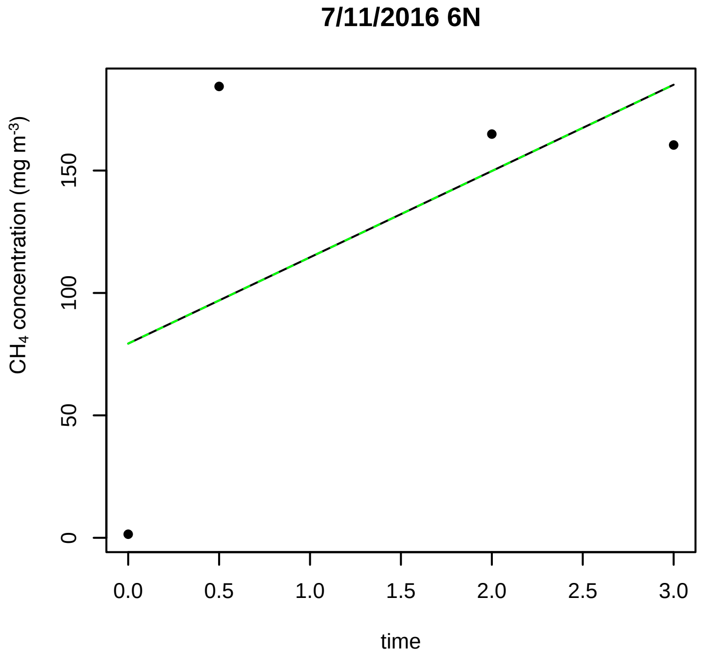
7/11/2016 6N
CH4 concentration (mg m-3)
time

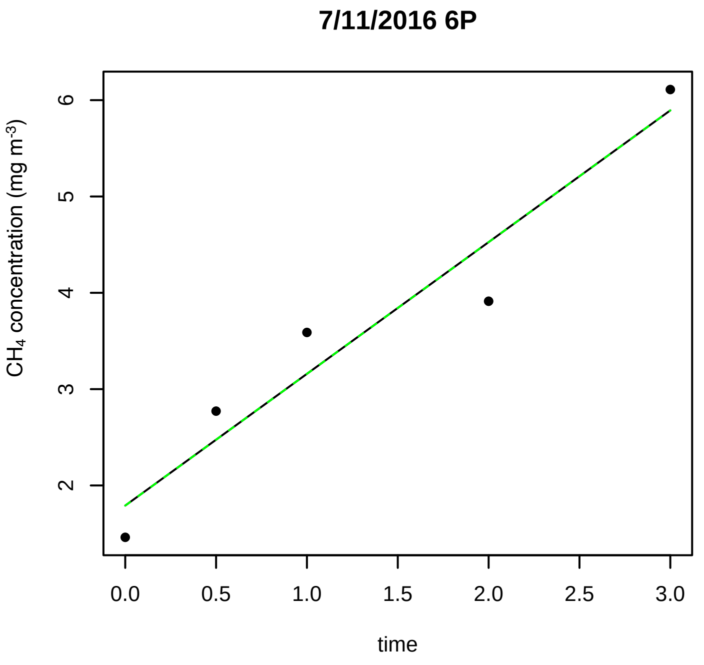
7/11/2016 6P
CH4 concentration (mg m-3)
time

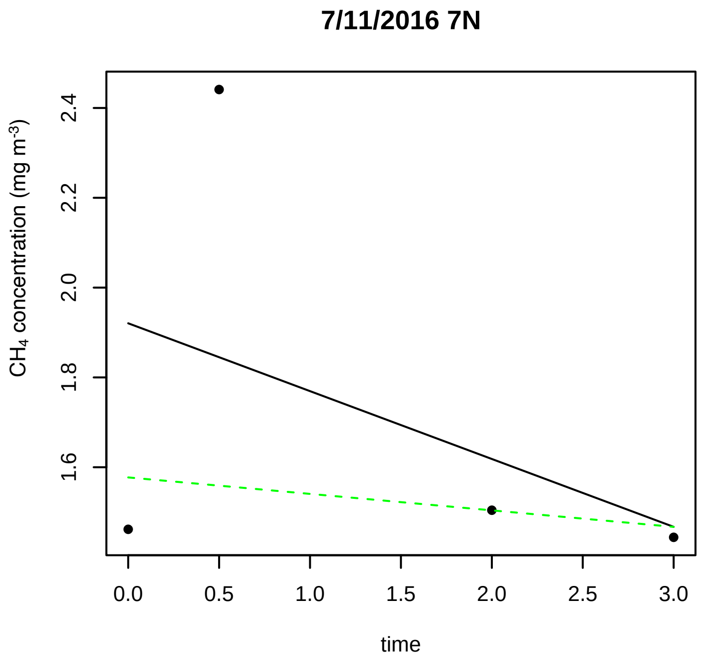
7/11/2016 7N
CH4 concentration (mg m-3)
time

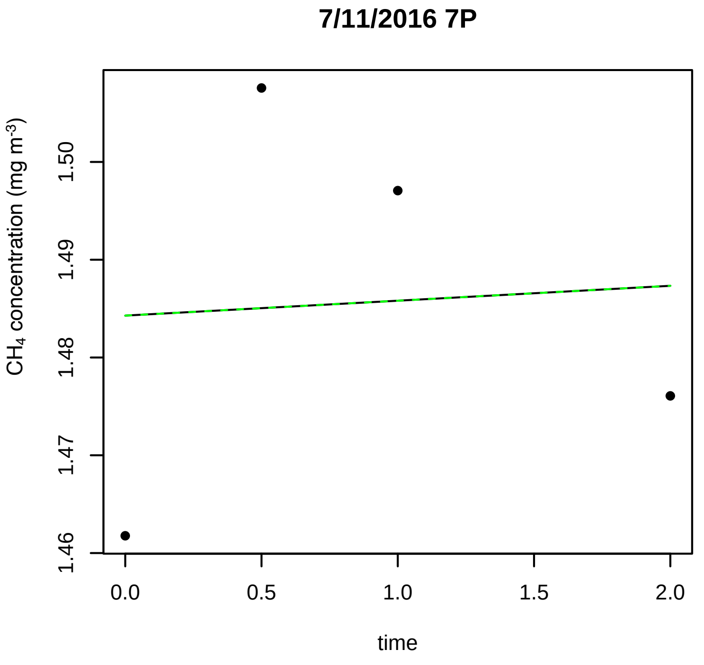
7/11/2016 7P
CH4 concentration (mg m-3)
time

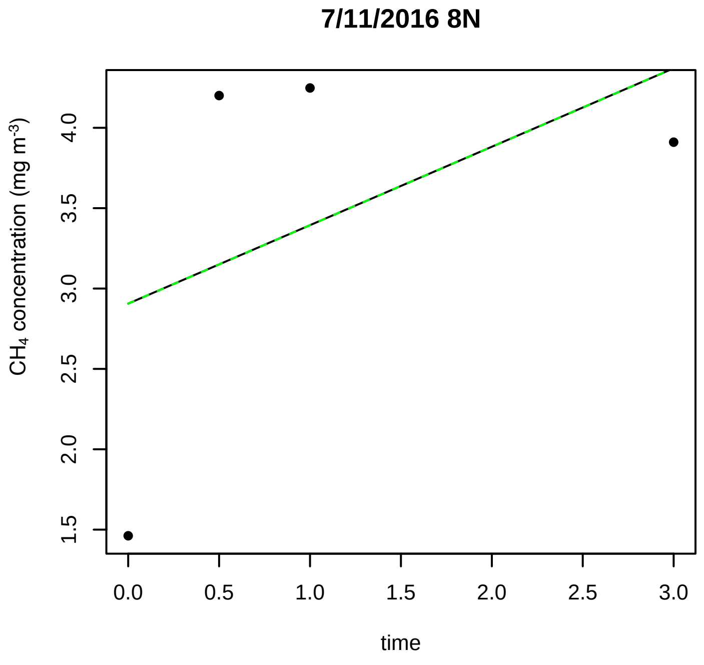
7/11/2016 8N
CH4 concentration (mg m-3)
time

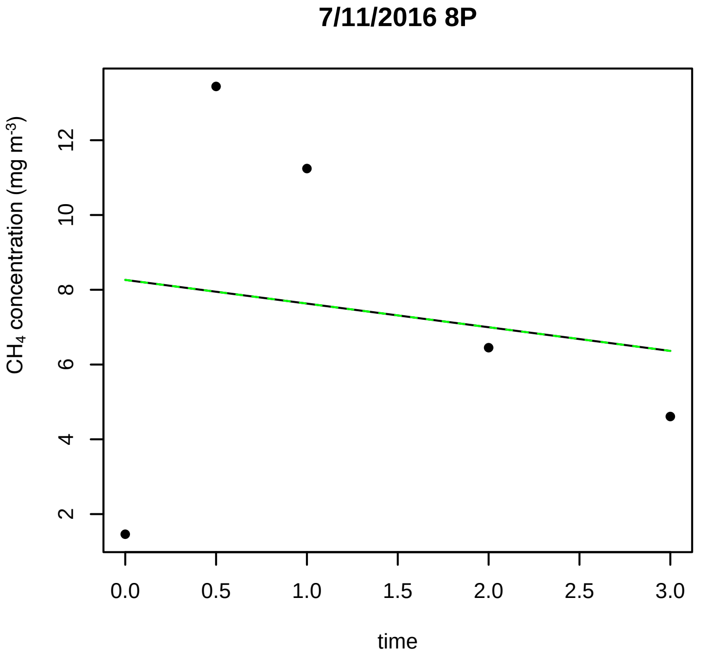
7/11/2016 8P
CH4 concentration (mg m-3)
time

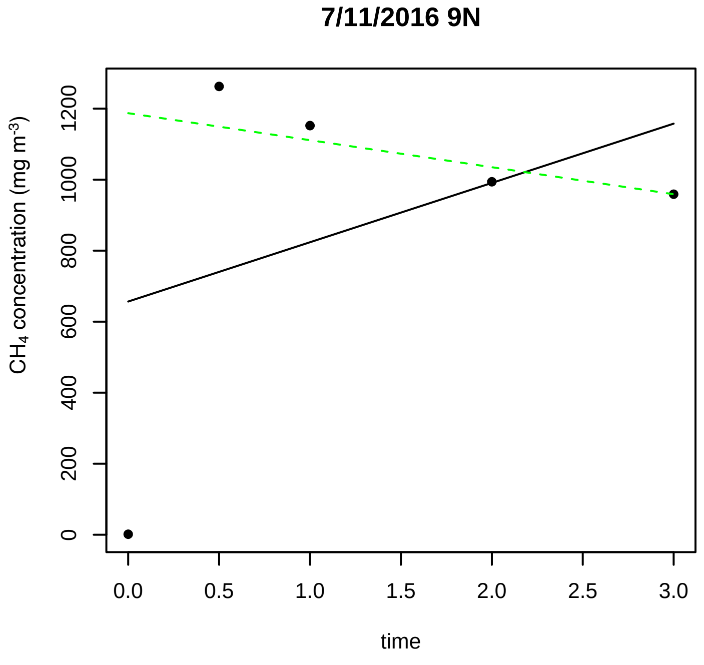
7/11/2016 9N
CH4 concentration (mg m-3)
time

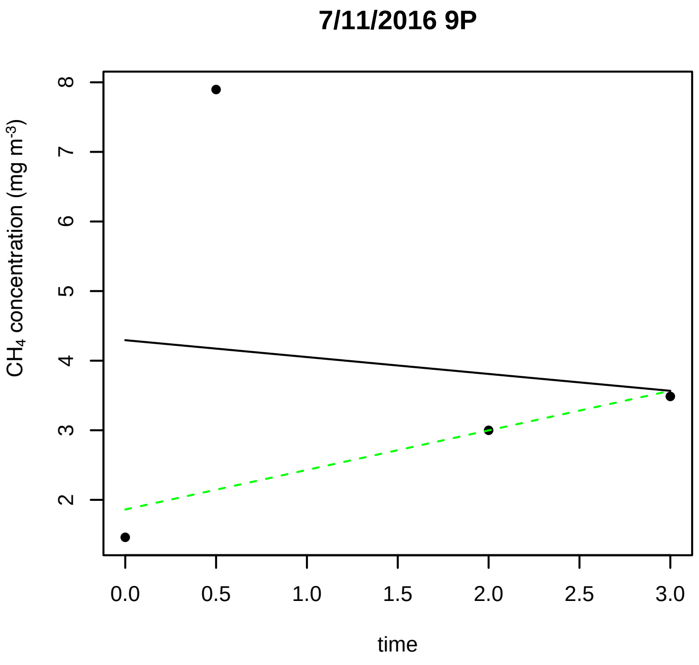
7/11/2016 9P
$CH_4$ concentration (mg $m^{-3}$)
2 3 4 5 6 7 8
0.0 0.5 1.0 1.5 2.0 2.5 3.0
time

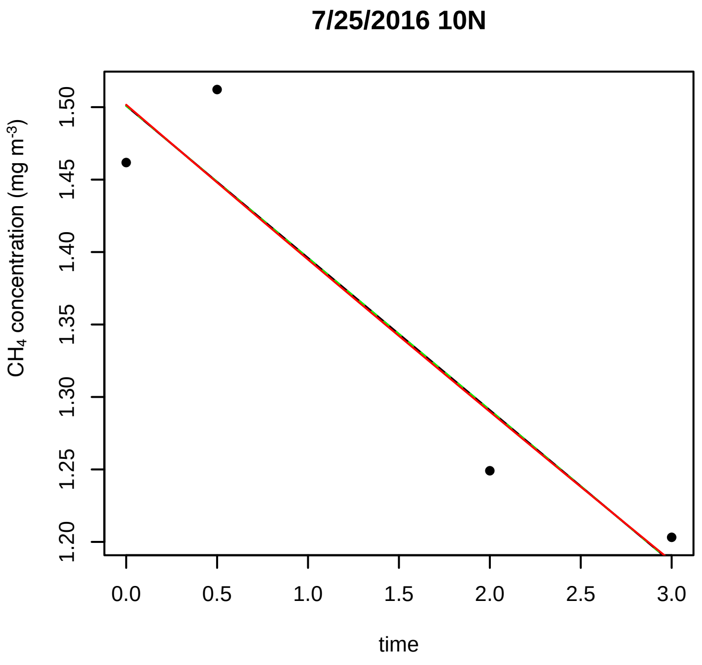
7/25/2016 10N
$CH_4$ concentration (mg $m^{-3}$)
1.20 1.25 1.30 1.35 1.40 1.45 1.50
0.0 0.5 1.0 1.5 2.0 2.5 3.0
time

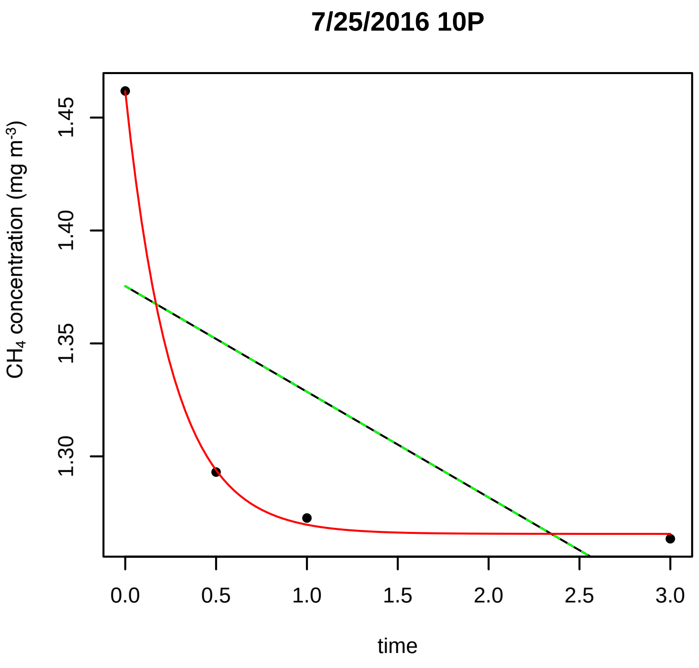
7/25/2016 10P
$CH_4$ concentration (mg $m^{-3}$)
1.30 1.35 1.40 1.45
0.0 0.5 1.0 1.5 2.0 2.5 3.0
time

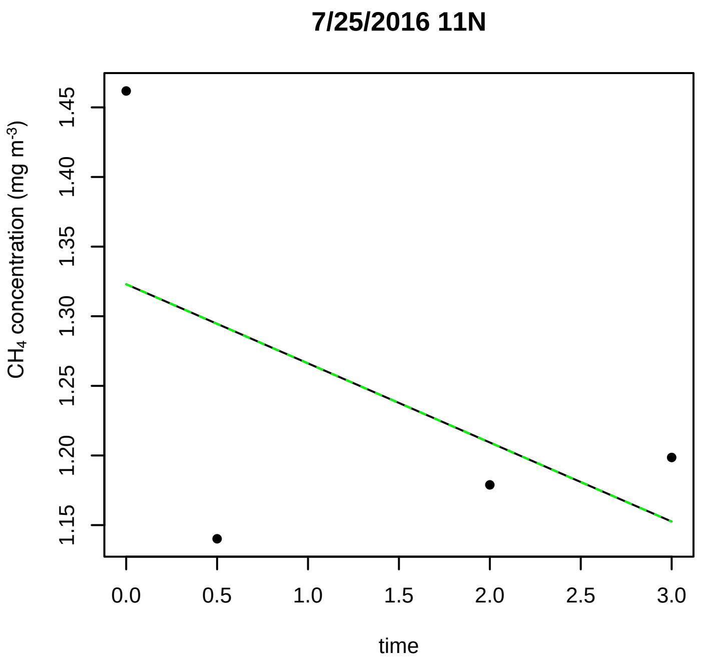
7/25/2016 11N
$CH_4$ concentration (mg $m^{-3}$)
1.15 1.20 1.25 1.30 1.35 1.40 1.45
0.0 0.5 1.0 1.5 2.0 2.5 3.0
time

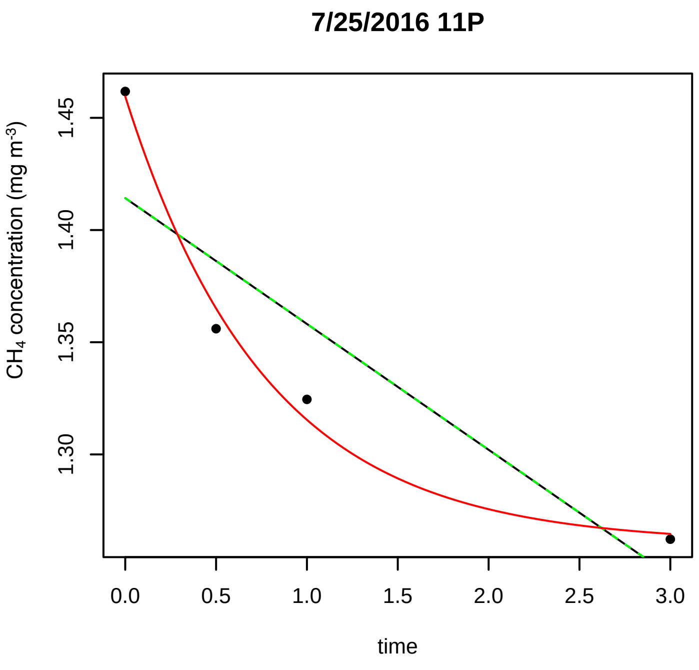
7/25/2016 11P
$CH_4$ concentration (mg $m^{-3}$)
1.30 1.35 1.40 1.45
0.0 0.5 1.0 1.5 2.0 2.5 3.0
time

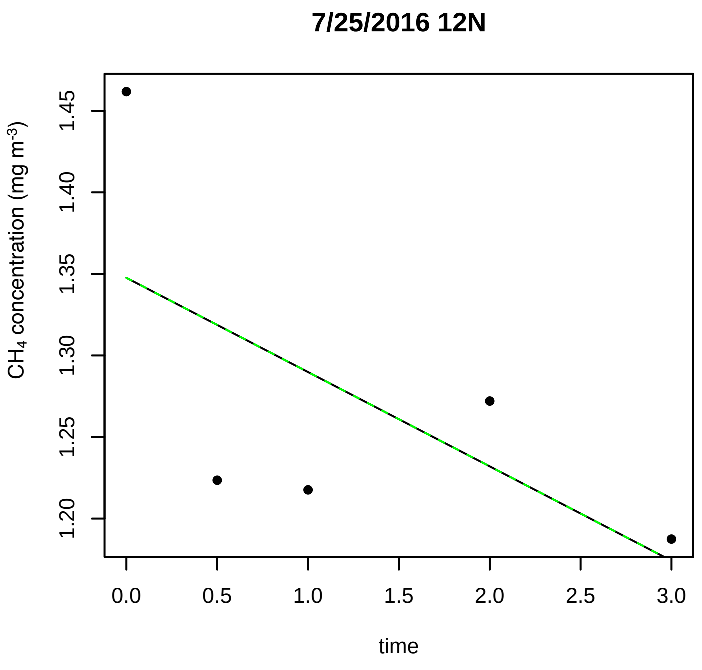
7/25/2016 12N
$CH_4$ concentration (mg $m^{-3}$)
1.20 1.25 1.30 1.35 1.40 1.45
0.0 0.5 1.0 1.5 2.0 2.5 3.0
time

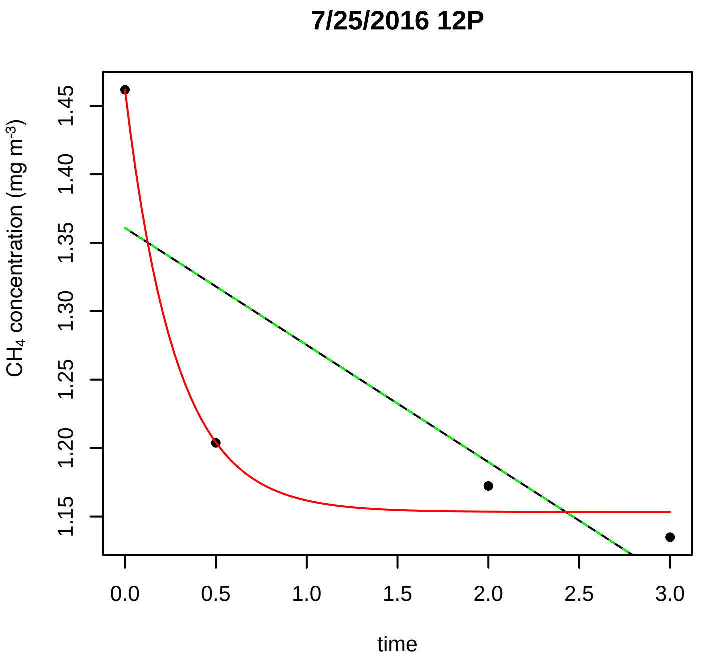
7/25/2016 12P
CH4 concentration (mg m-3)
time

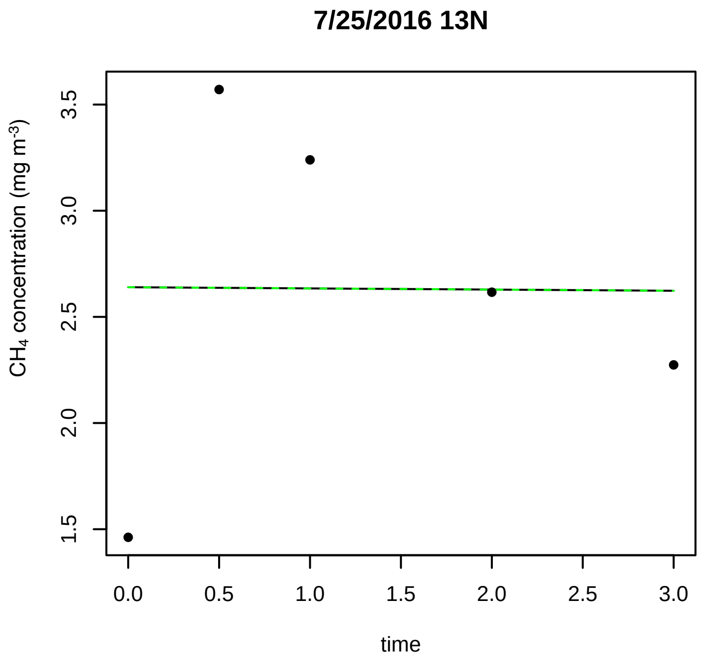
7/25/2016 13N
CH4 concentration (mg m-3)
time

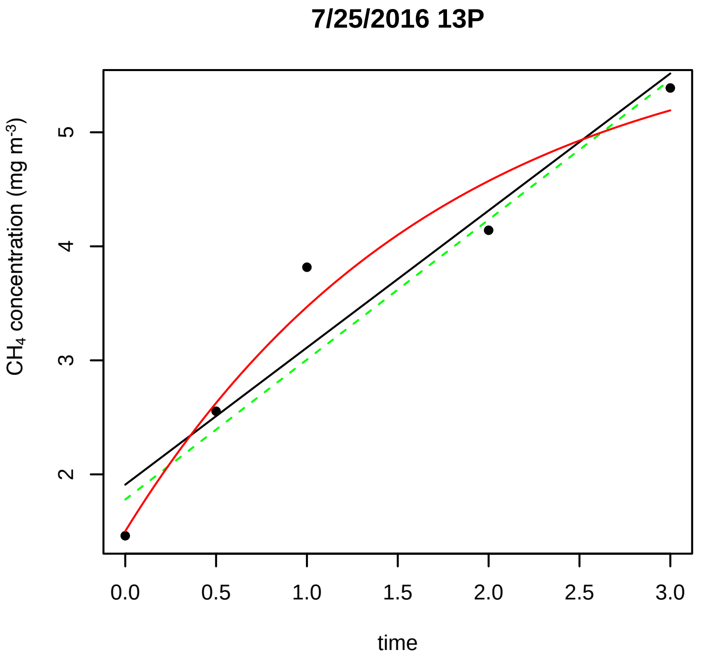
7/25/2016 13P
CH4 concentration (mg m-3)
time

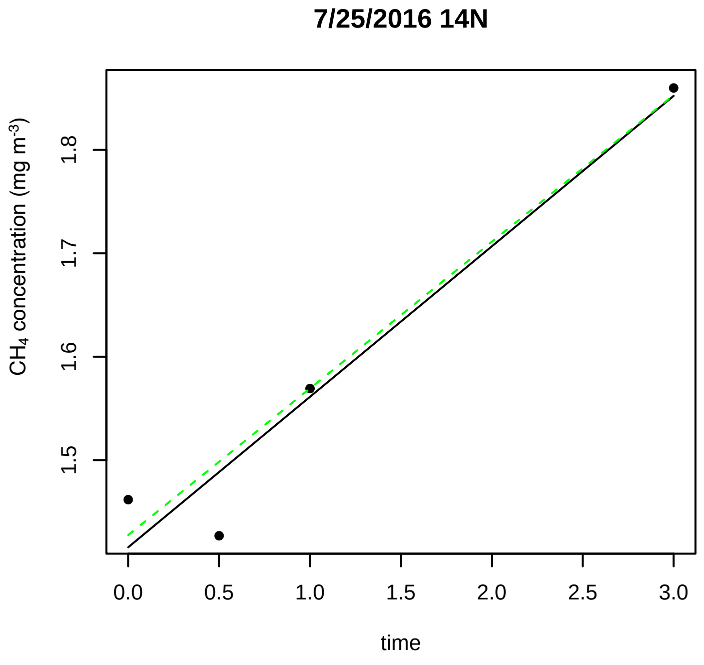
7/25/2016 14N
CH4 concentration (mg m-3)
time

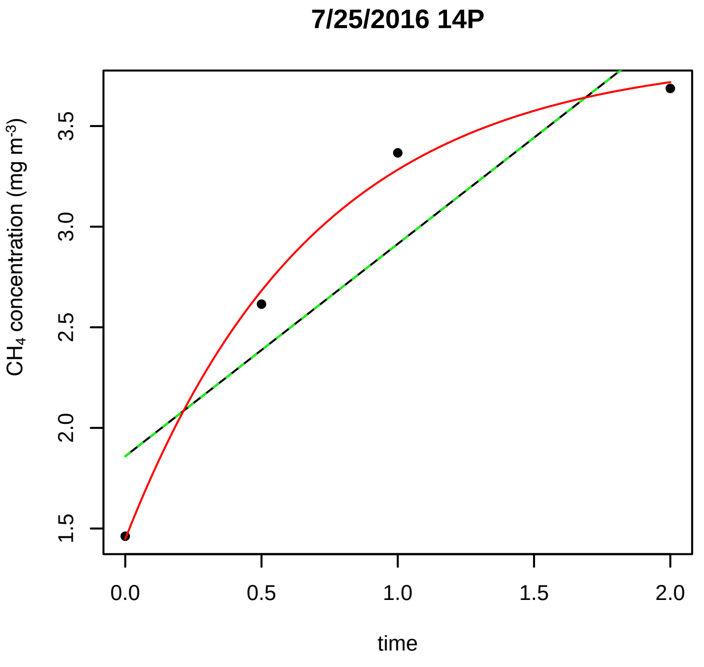
7/25/2016 14P
CH4 concentration (mg m-3)
time

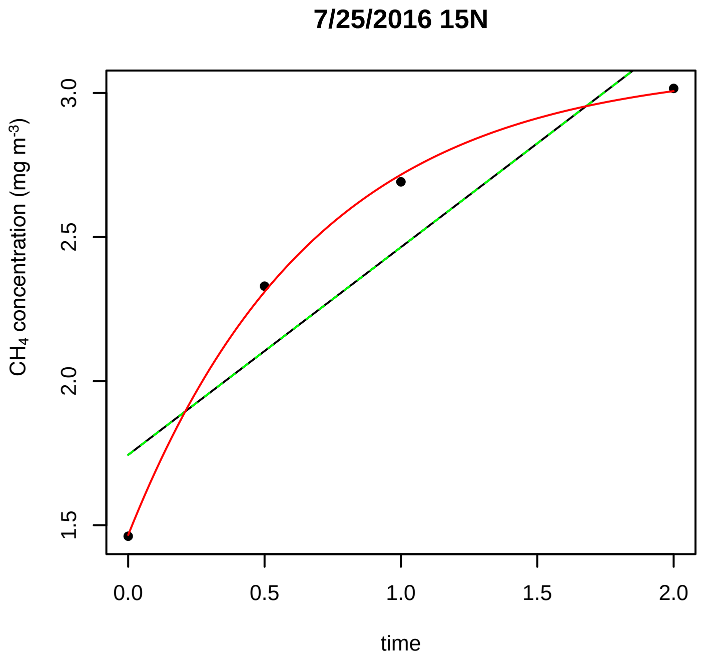
7/25/2016 15N
CH4 concentration (mg m-3)
time

7/25/2016 15P

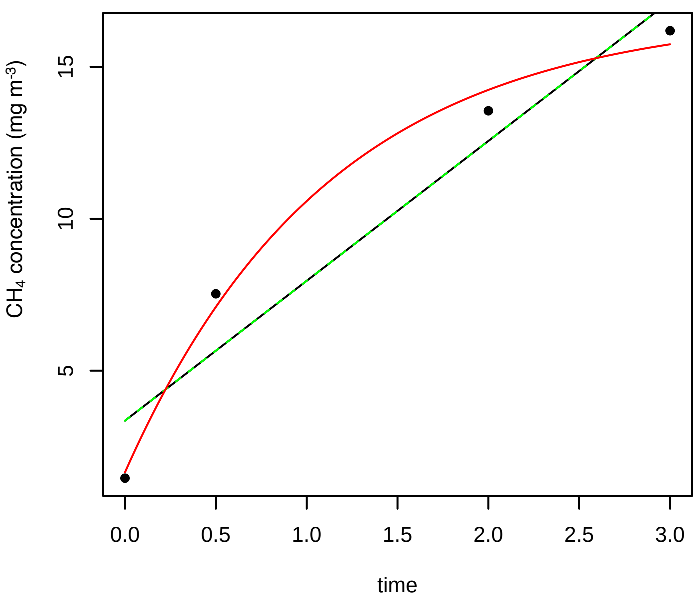


7/25/2016 16N

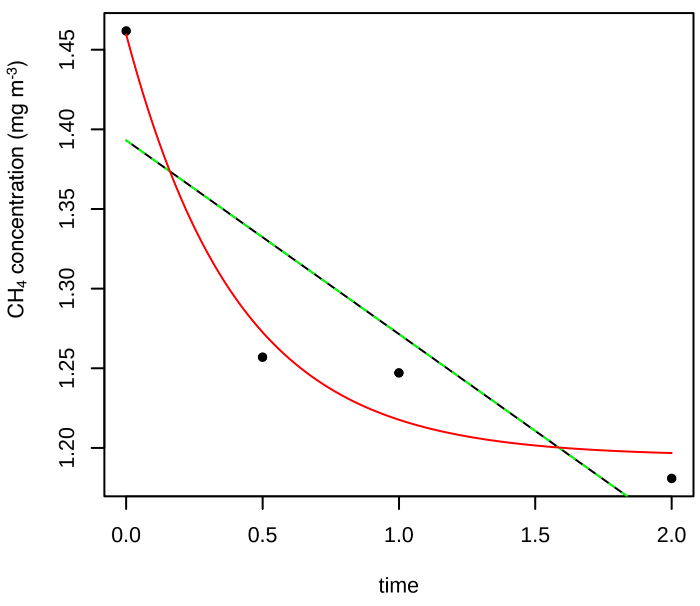


7/25/2016 16P

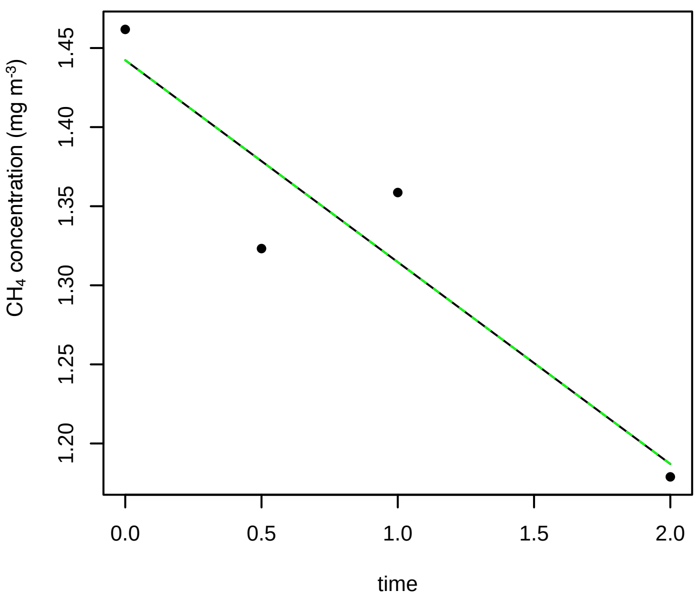


7/25/2016 17N

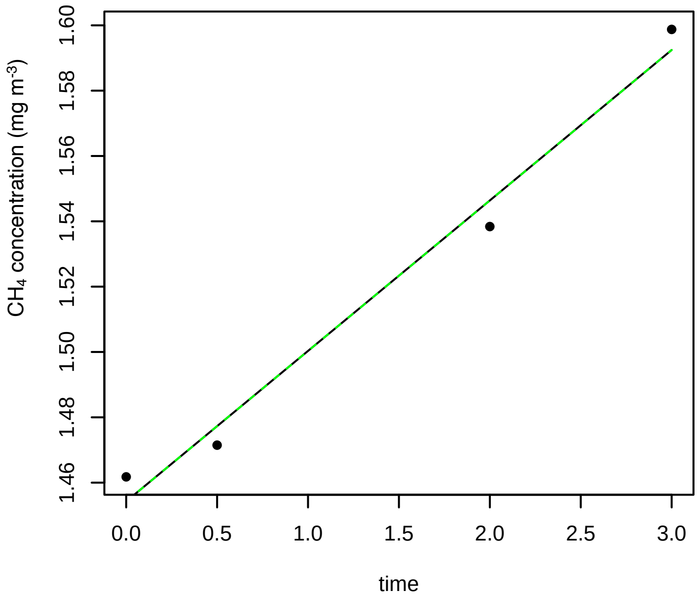


7/25/2016 17P

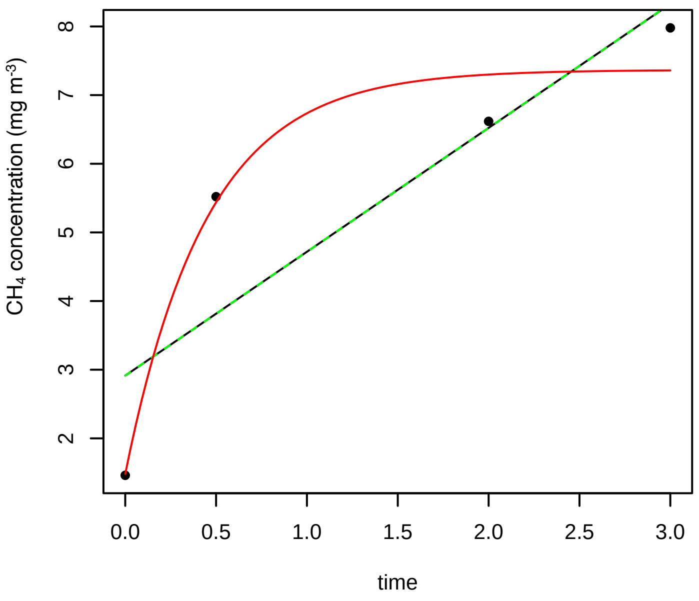


7/25/2016 18N

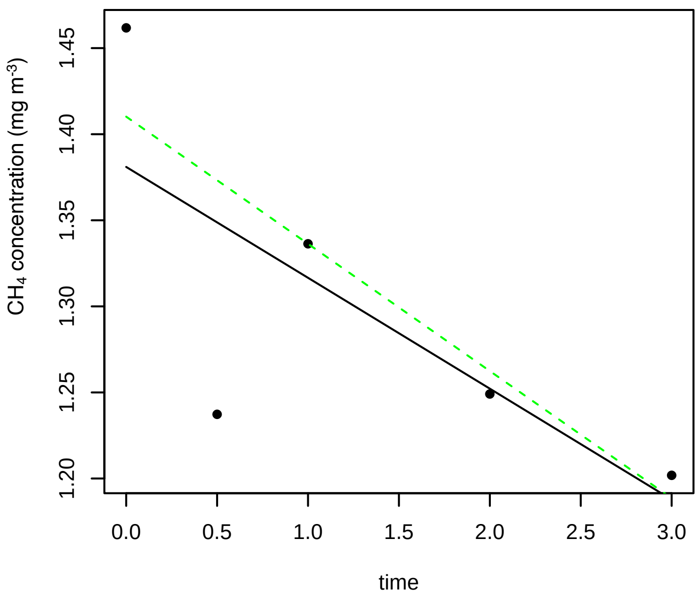

**7/25/2016 18P**

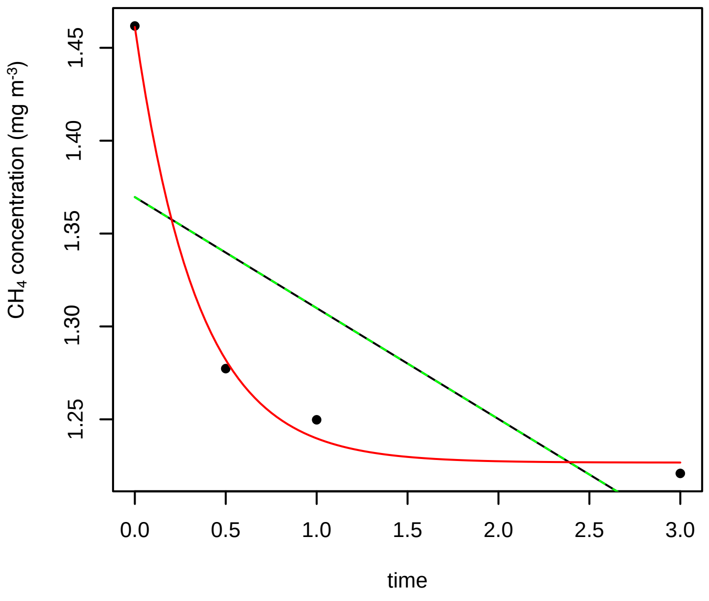


**7/25/2016 1N**

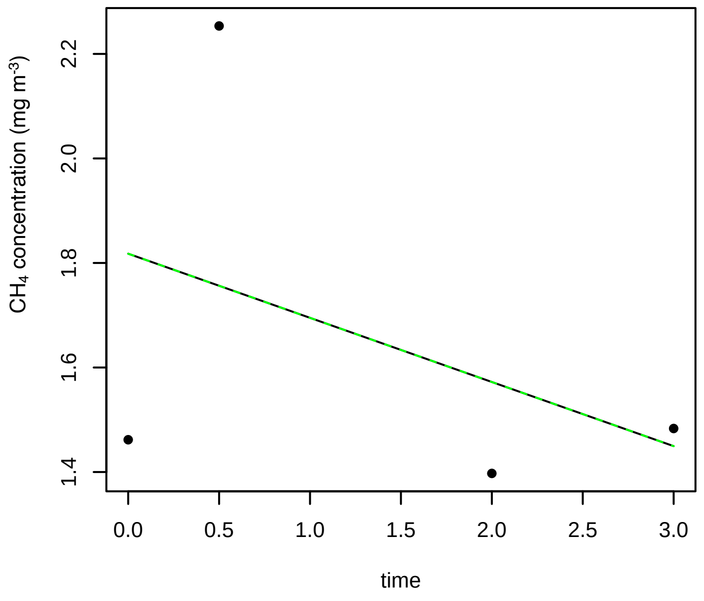


**7/25/2016 1P**

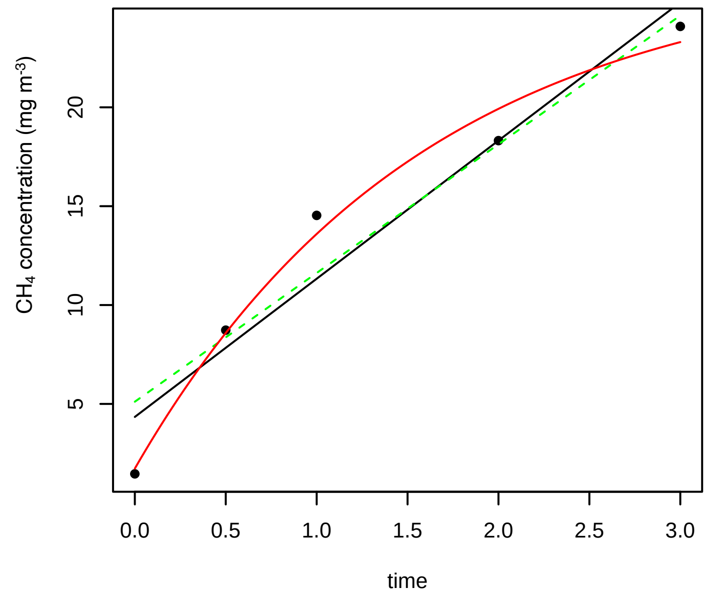


**7/25/2016 2N**

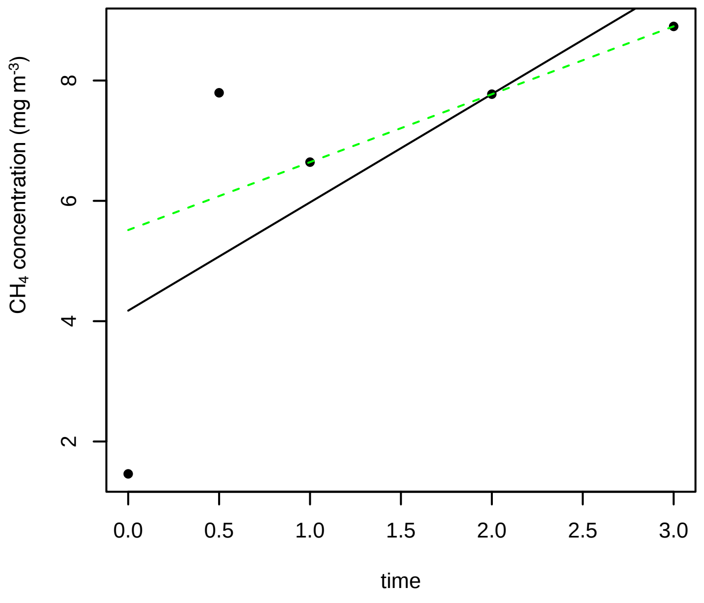


**7/25/2016 2P**

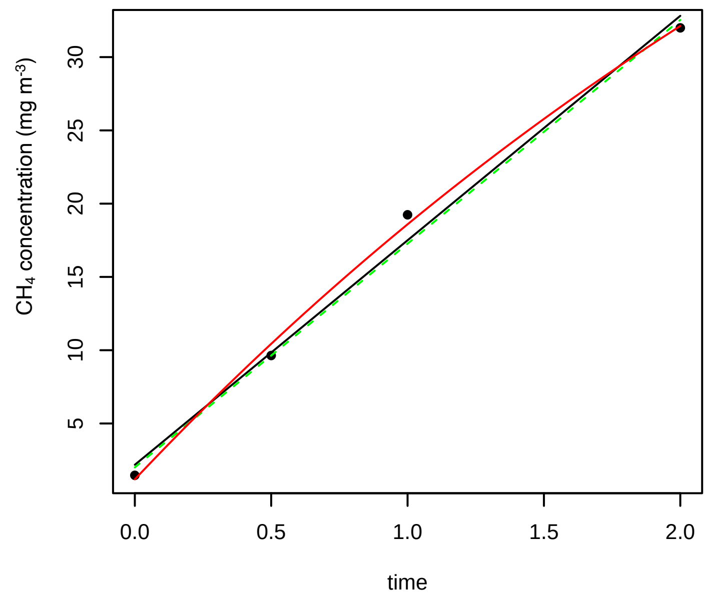


**7/25/2016 3N**

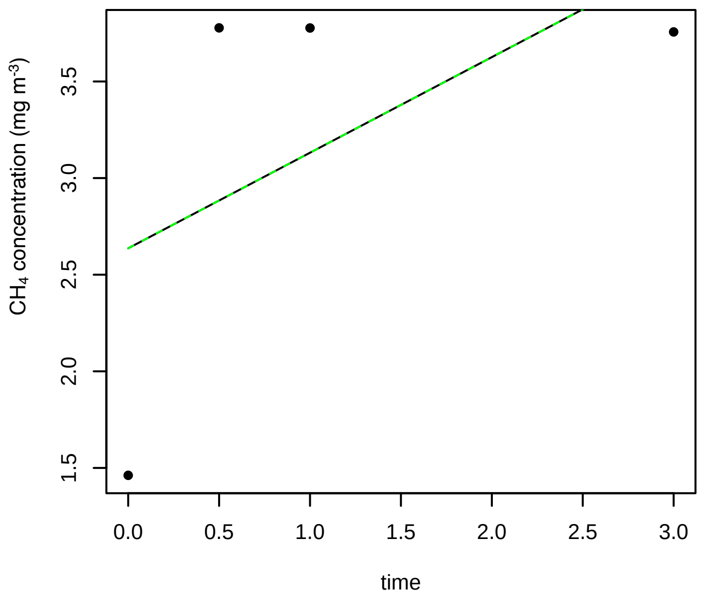

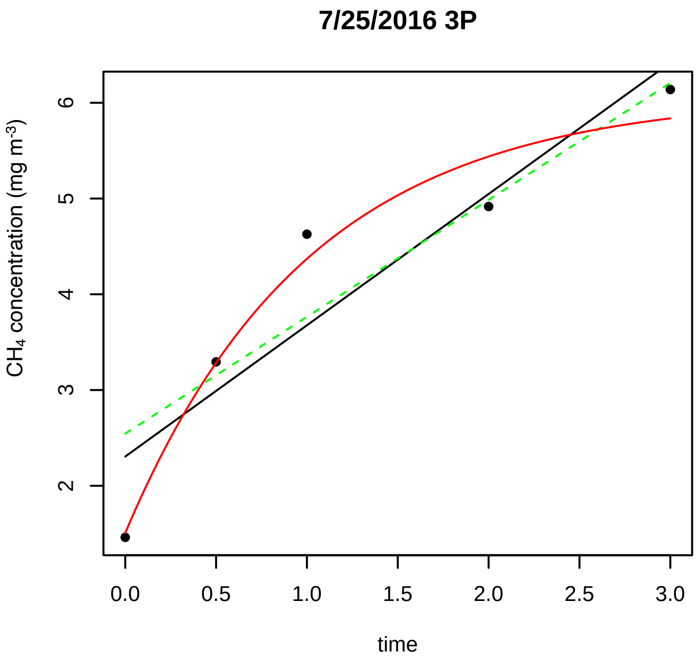
7/25/2016 3P
CH4 concentration (mg m-3)
time
0.0 0.5 1.0 1.5 2.0 2.5 3.0
2 3 4 5 6

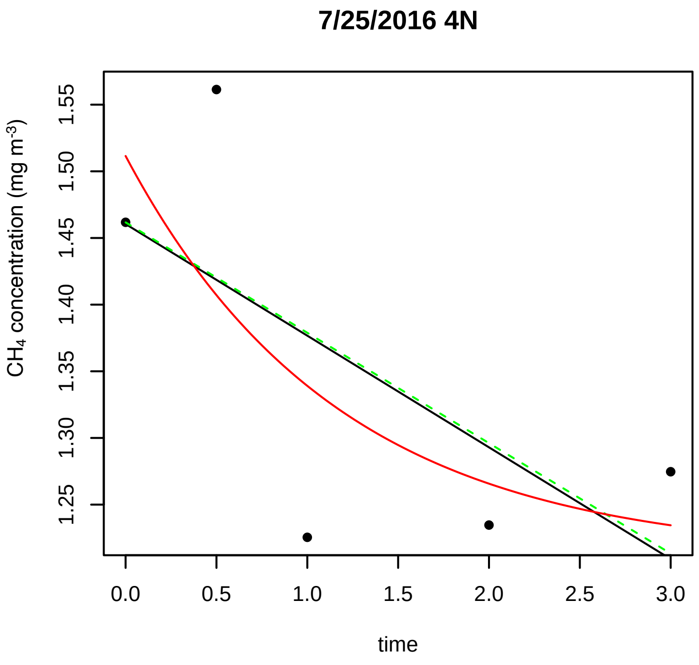
7/25/2016 4N
CH4 concentration (mg m-3)
time
0.0 0.5 1.0 1.5 2.0 2.5 3.0
1.25 1.30 1.35 1.40 1.45 1.50 1.55

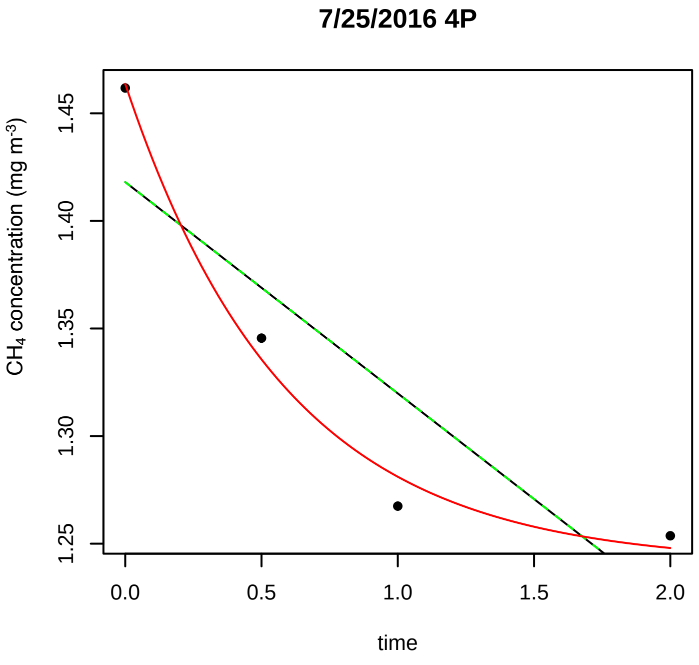
7/25/2016 4P
CH4 concentration (mg m-3)
time
0.0 0.5 1.0 1.5 2.0
1.25 1.30 1.35 1.40 1.45

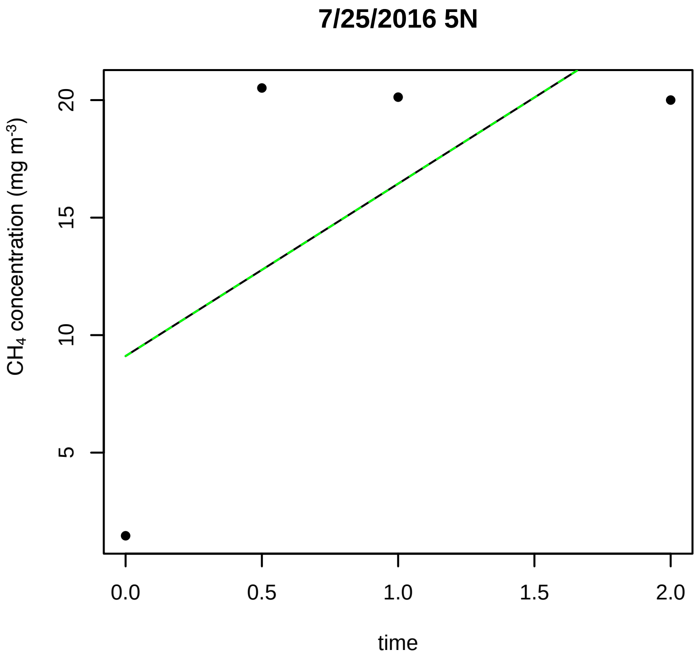
7/25/2016 5N
CH4 concentration (mg m-3)
time
0.0 0.5 1.0 1.5 2.0
5 10 15 20

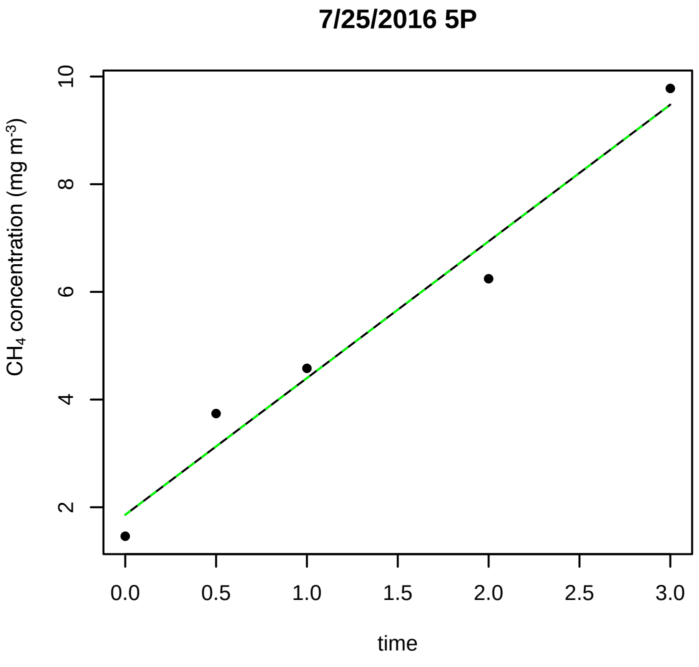
7/25/2016 5P
CH4 concentration (mg m-3)
time
0.0 0.5 1.0 1.5 2.0 2.5 3.0
2 4 6 8 10

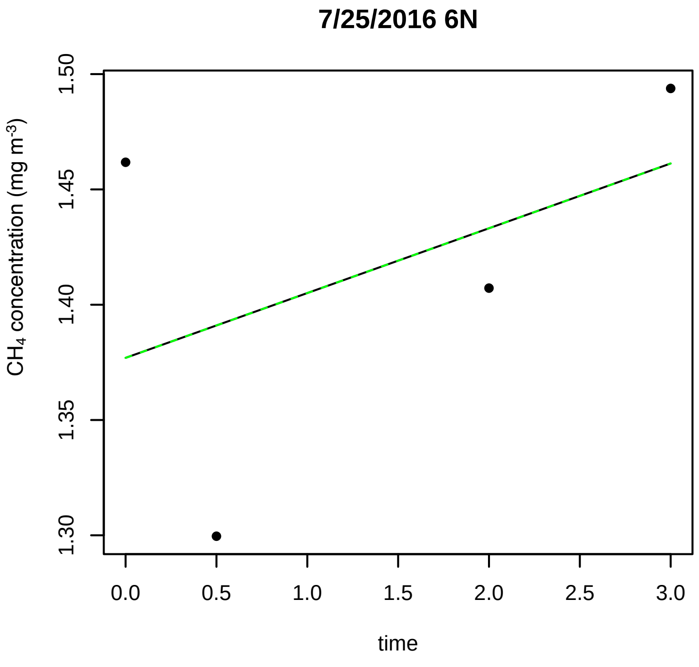
7/25/2016 6N
CH4 concentration (mg m-3)
time
0.0 0.5 1.0 1.5 2.0 2.5 3.0
1.30 1.35 1.40 1.45 1.50

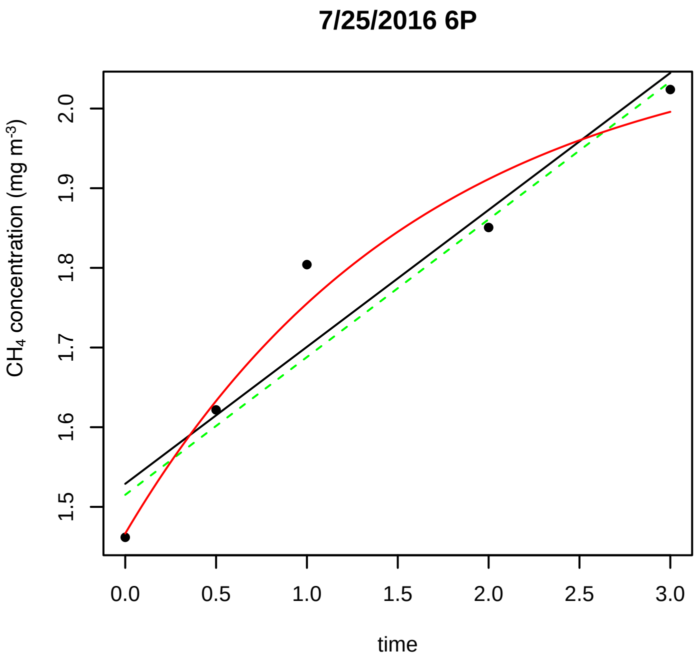
7/25/2016 6P
CH4 concentration (mg m-3)
time

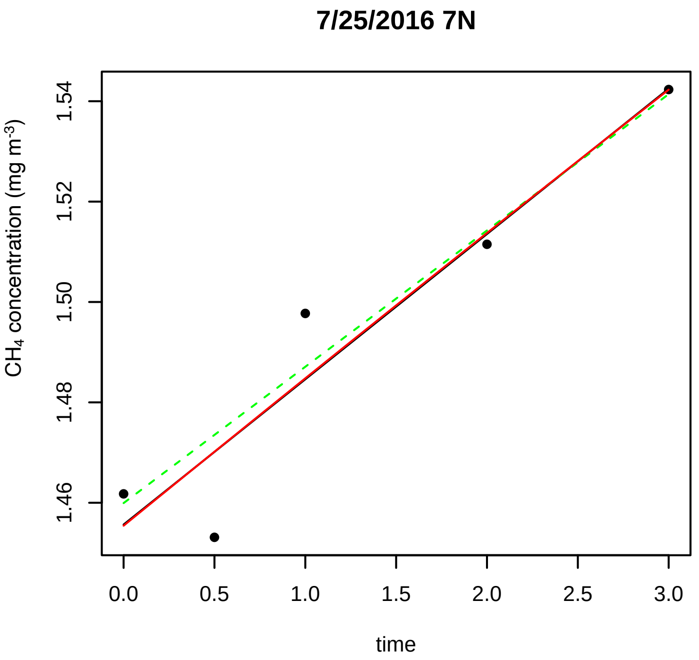
7/25/2016 7N
CH4 concentration (mg m-3)
time

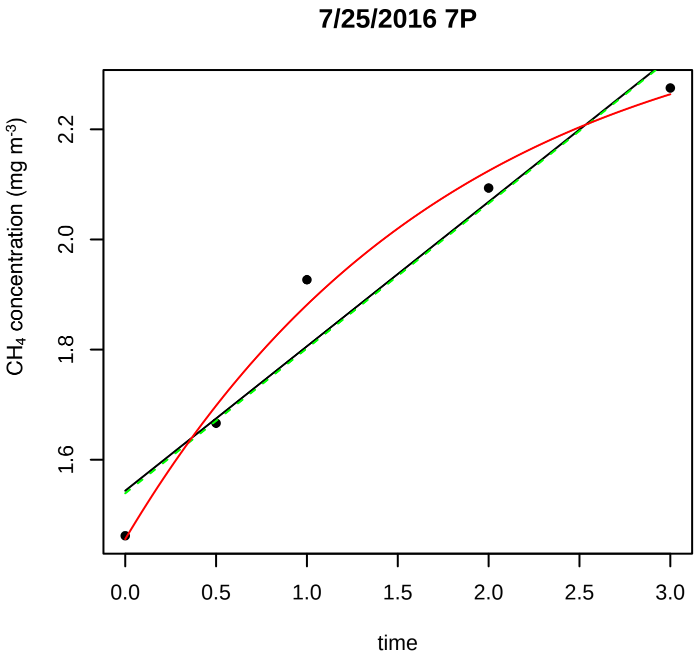
7/25/2016 7P
CH4 concentration (mg m-3)
time

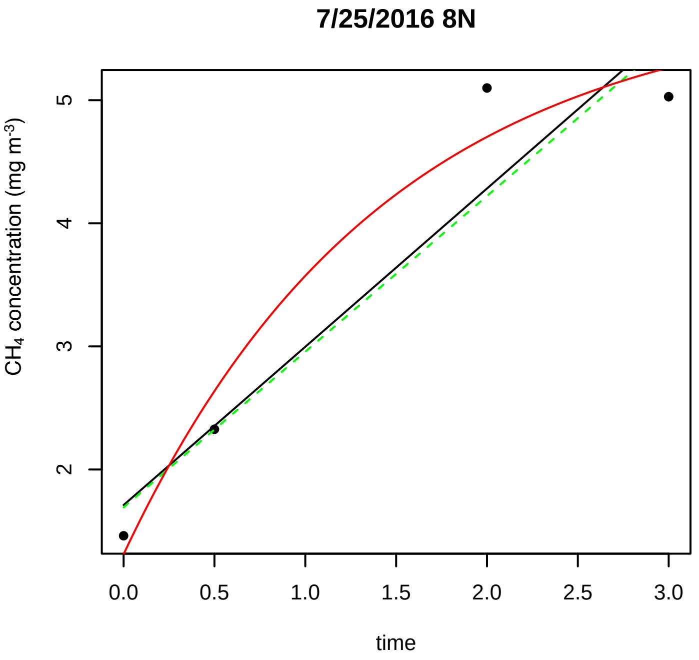
7/25/2016 8N
CH4 concentration (mg m-3)
time

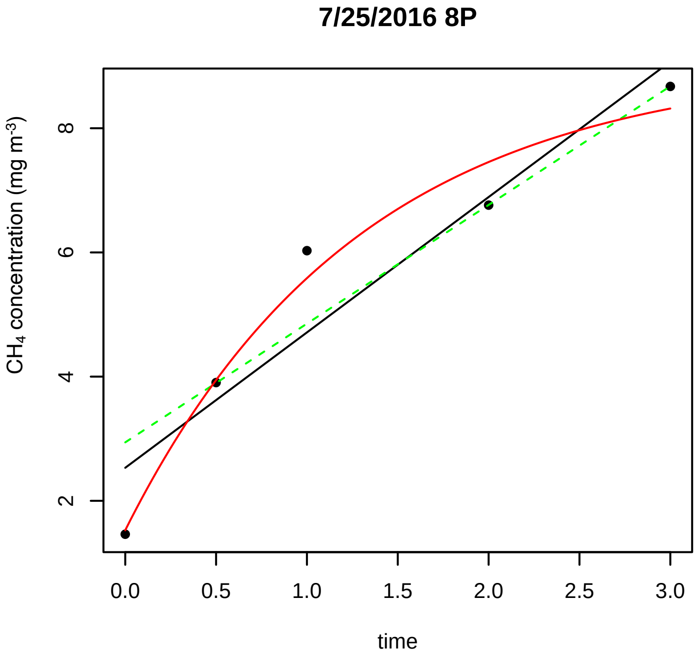
7/25/2016 8P
CH4 concentration (mg m-3)
time

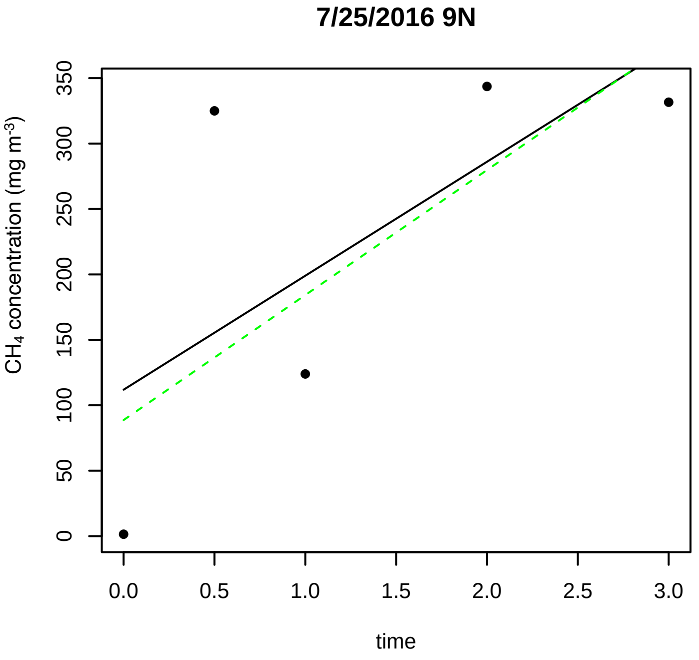
7/25/2016 9N
CH4 concentration (mg m-3)
time

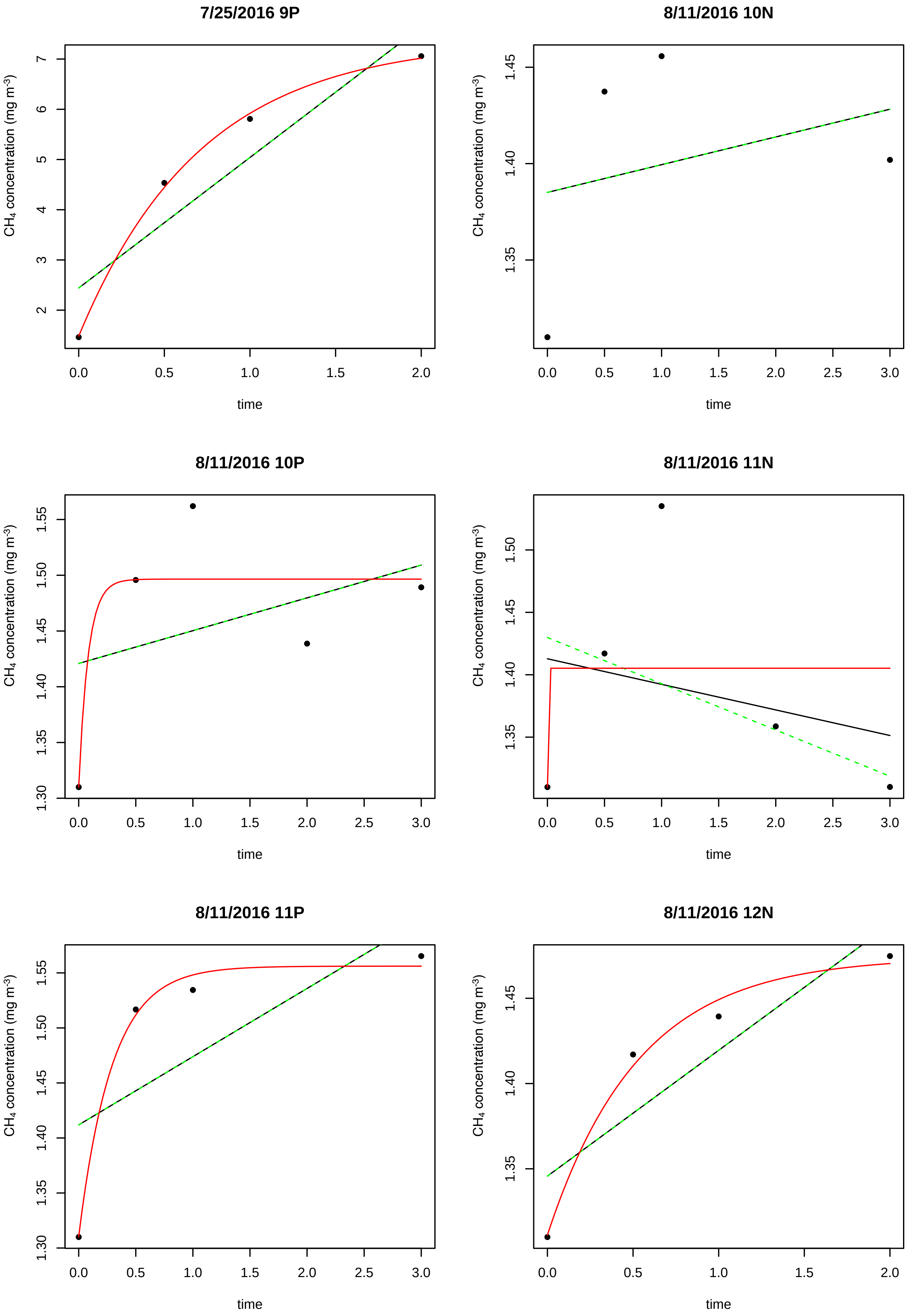

7/25/2016 9P
8/11/2016 10N
8/11/2016 10P
8/11/2016 11N
8/11/2016 11P
8/11/2016 12N
CH4 concentration (mg m-3)
time

**8/11/2016 12P**

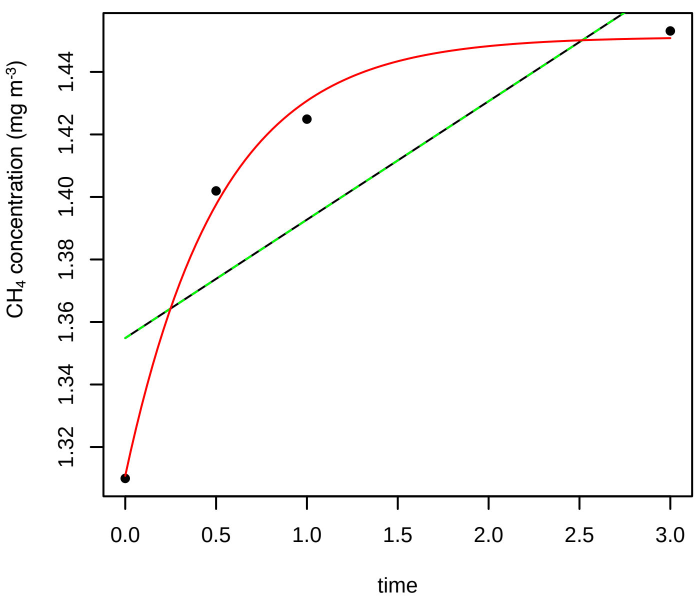


**8/11/2016 13N**

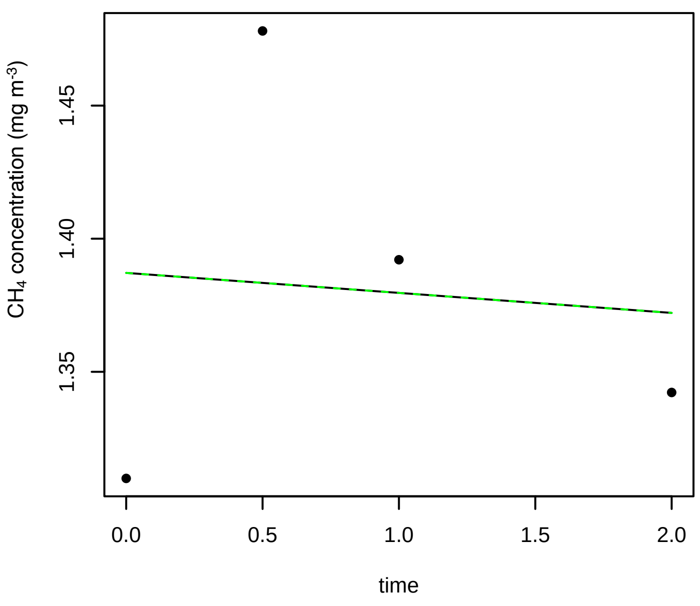


**8/11/2016 13P**

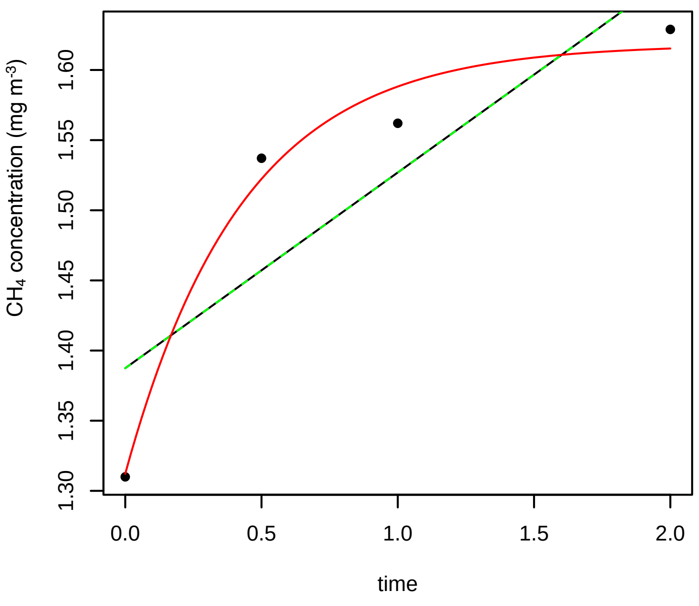


**8/11/2016 14N**

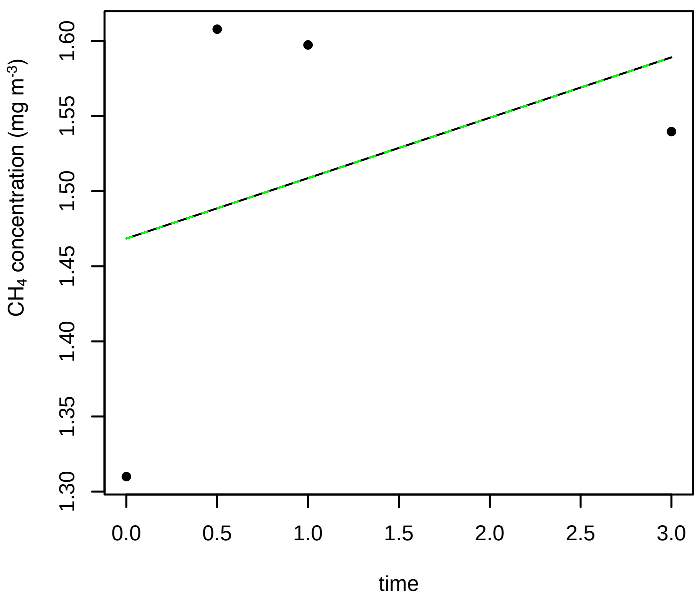


**8/11/2016 14P**

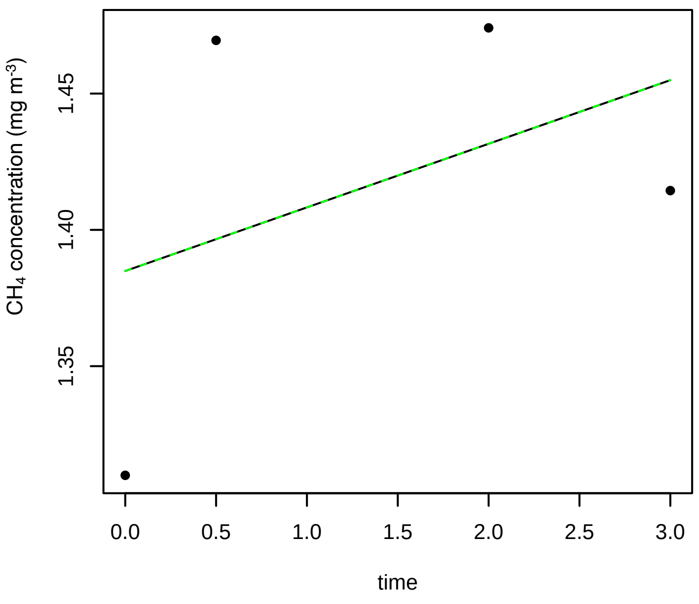


**8/11/2016 15N**

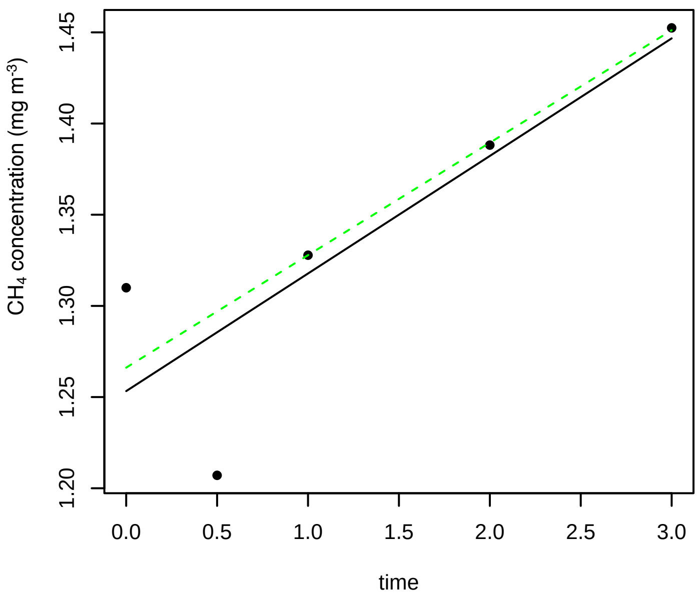

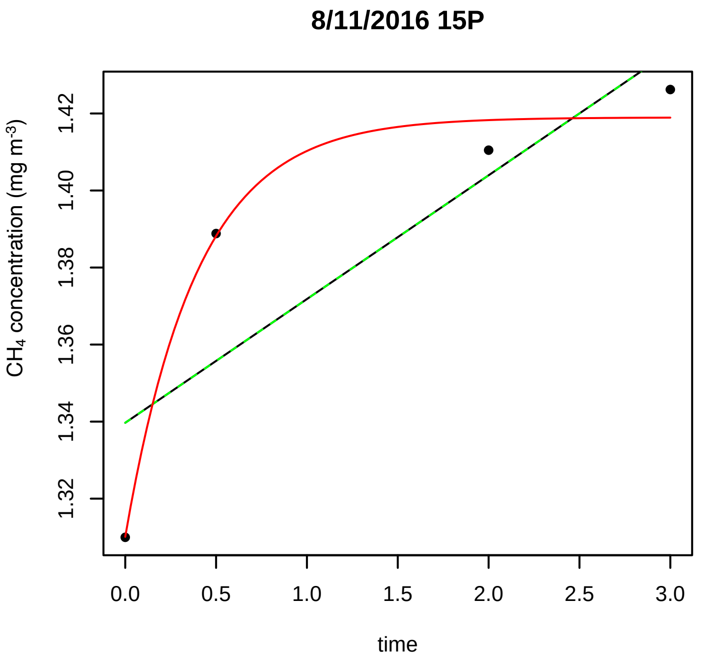
8/11/2016 15P
CH4 concentration (mg m-3)
1.32
1.34
1.36
1.38
1.40
1.42
0.0
0.5
1.0
1.5
2.0
2.5
3.0
time

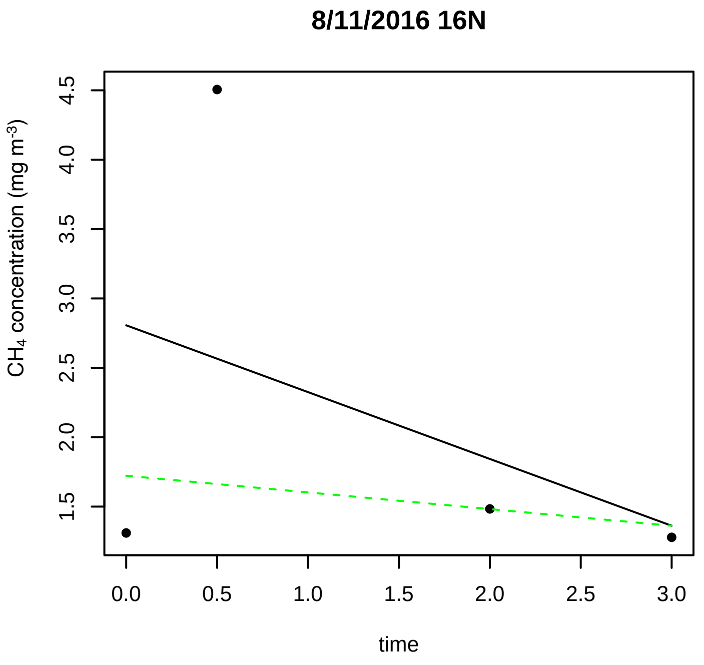
8/11/2016 16N
CH4 concentration (mg m-3)
1.5
2.0
2.5
3.0
3.5
4.0
4.5
0.0
0.5
1.0
1.5
2.0
2.5
3.0
time

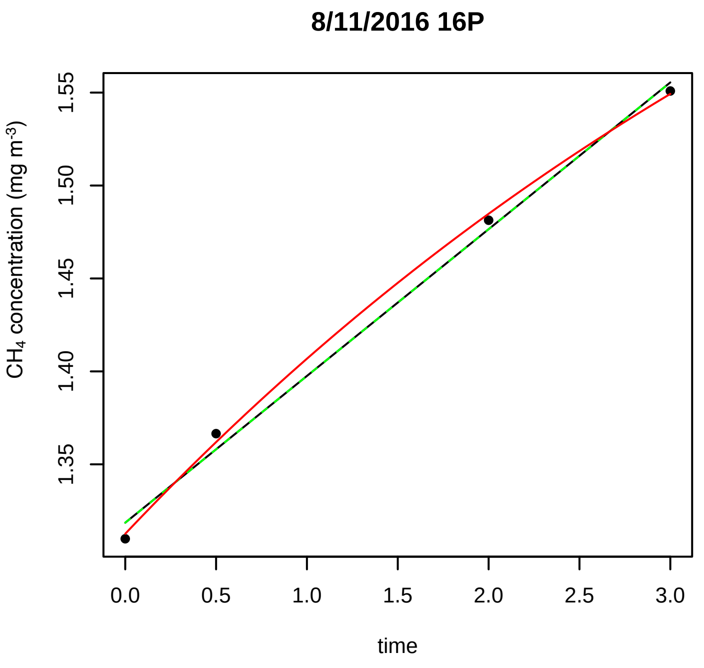
8/11/2016 16P
CH4 concentration (mg m-3)
1.35
1.40
1.45
1.50
1.55
0.0
0.5
1.0
1.5
2.0
2.5
3.0
time

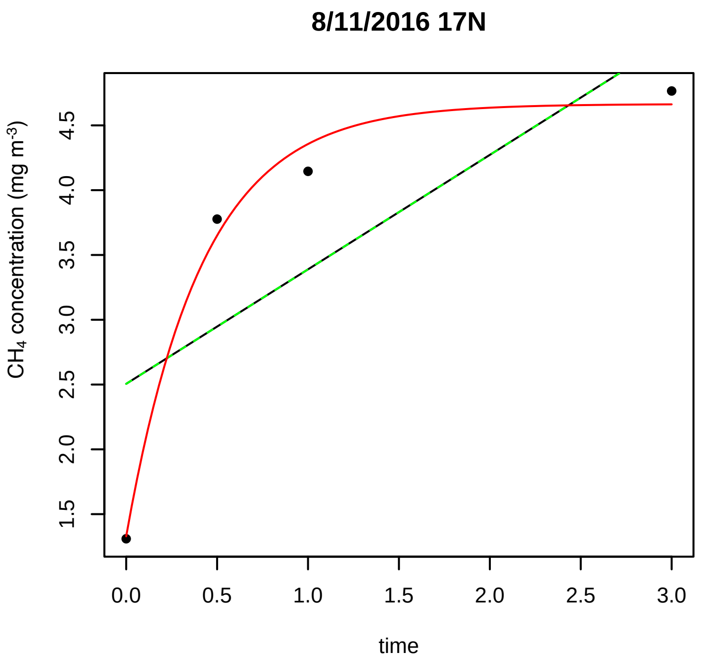
8/11/2016 17N
CH4 concentration (mg m-3)
1.5
2.0
2.5
3.0
3.5
4.0
4.5
0.0
0.5
1.0
1.5
2.0
2.5
3.0
time

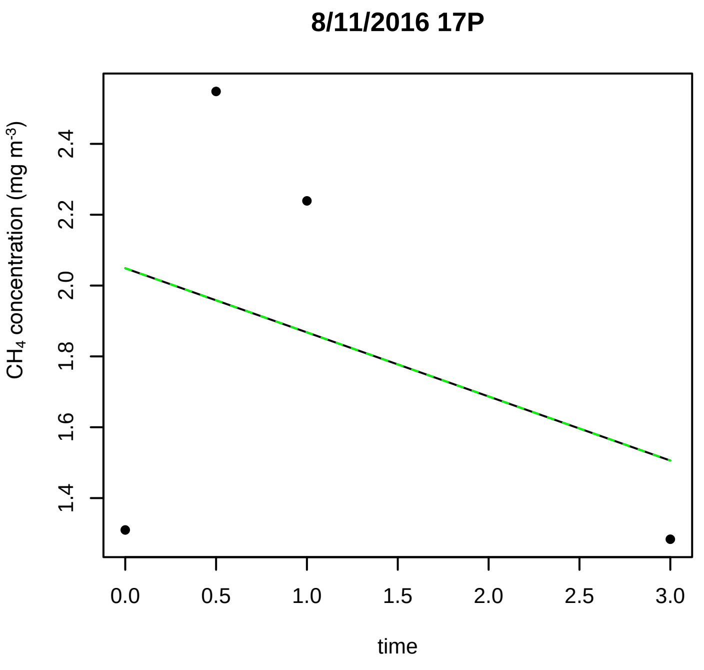
8/11/2016 17P
CH4 concentration (mg m-3)
1.4
1.6
1.8
2.0
2.2
2.4
0.0
0.5
1.0
1.5
2.0
2.5
3.0
time

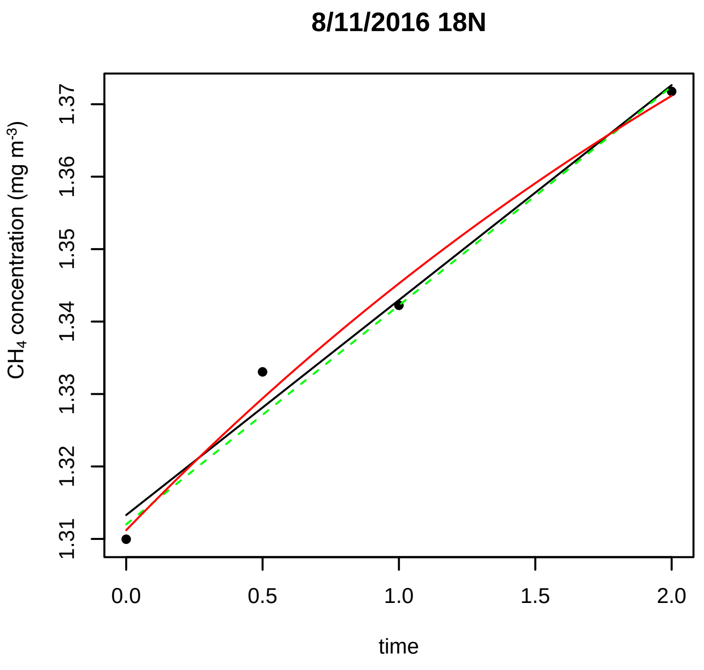
8/11/2016 18N
CH4 concentration (mg m-3)
1.31
1.32
1.33
1.34
1.35
1.36
1.37
0.0
0.5
1.0
1.5
2.0
time

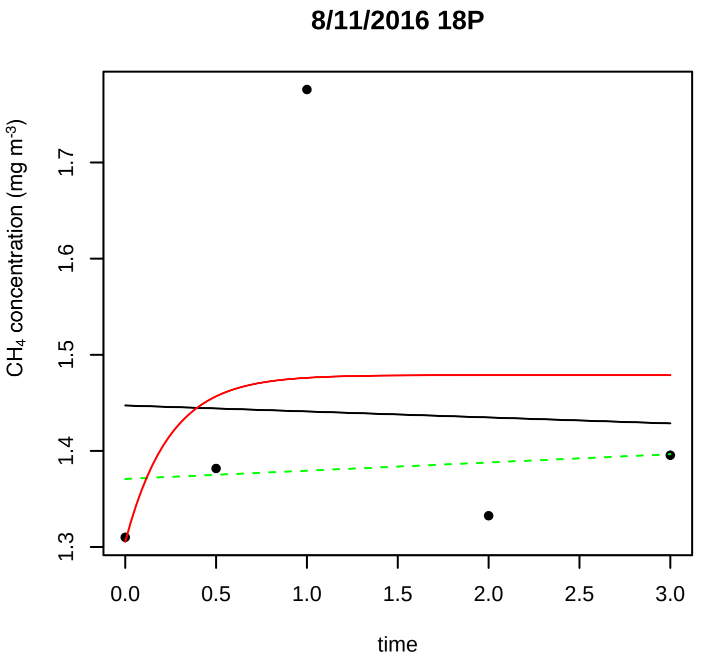
8/11/2016 18P
CH4 concentration (mg m-3)
1.3
1.4
1.5
1.6
1.7
0.0
0.5
1.0
1.5
2.0
2.5
3.0
time

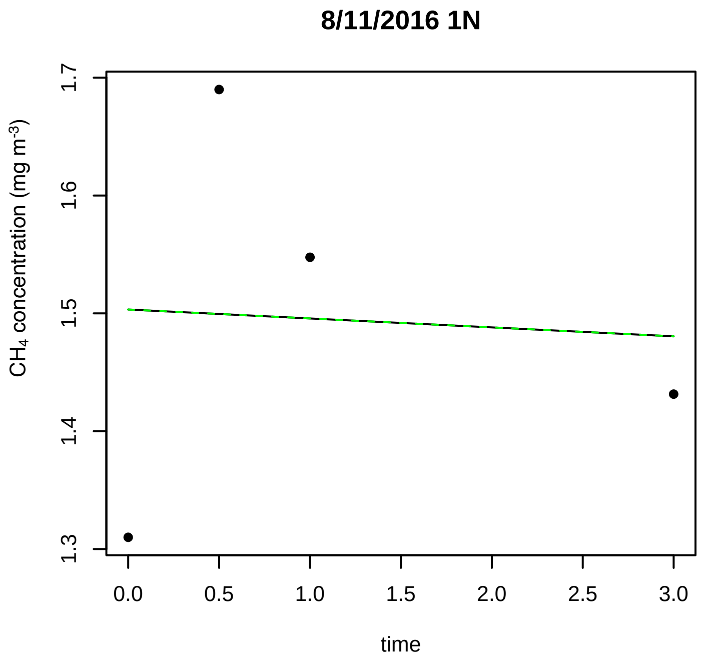
8/11/2016 1N
CH4 concentration (mg m-3)
1.3
1.4
1.5
1.6
1.7
0.0
0.5
1.0
1.5
2.0
2.5
3.0
time

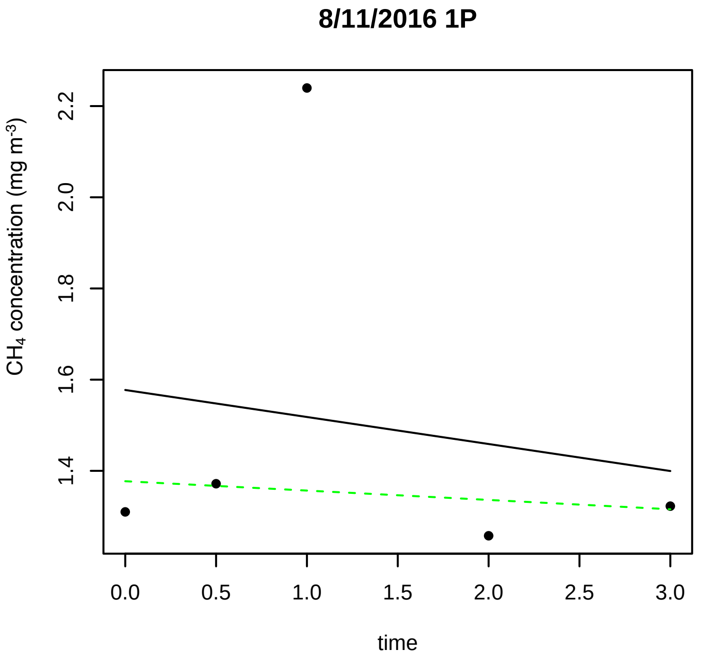
8/11/2016 1P
CH4 concentration (mg m-3)
1.4
1.6
1.8
2.0
2.2
0.0
0.5
1.0
1.5
2.0
2.5
3.0
time

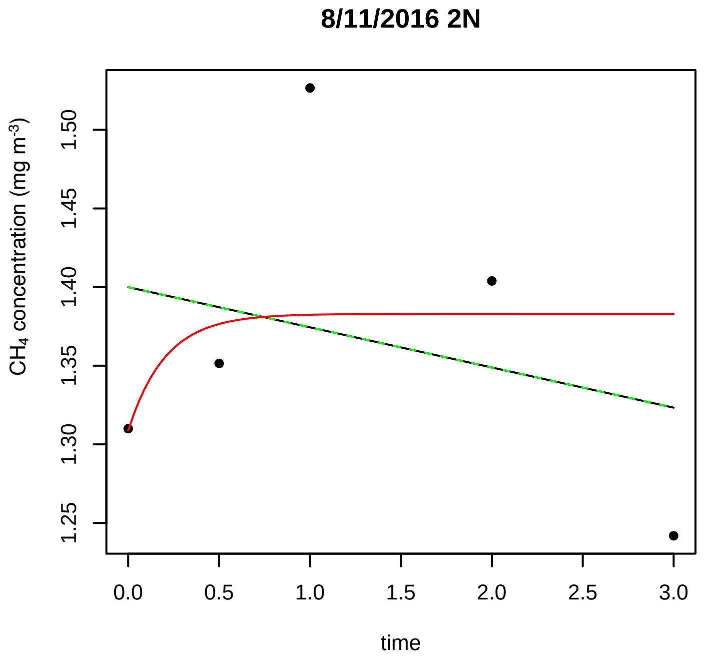
8/11/2016 2N
CH4 concentration (mg m-3)
1.25
1.30
1.35
1.40
1.45
1.50
0.0
0.5
1.0
1.5
2.0
2.5
3.0
time

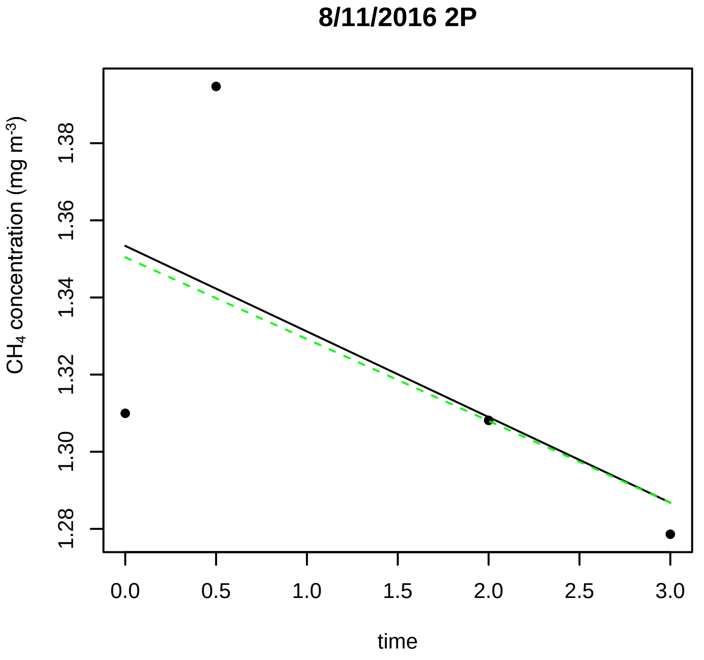
8/11/2016 2P
CH4 concentration (mg m-3)
1.28
1.30
1.32
1.34
1.36
1.38
0.0
0.5
1.0
1.5
2.0
2.5
3.0
time

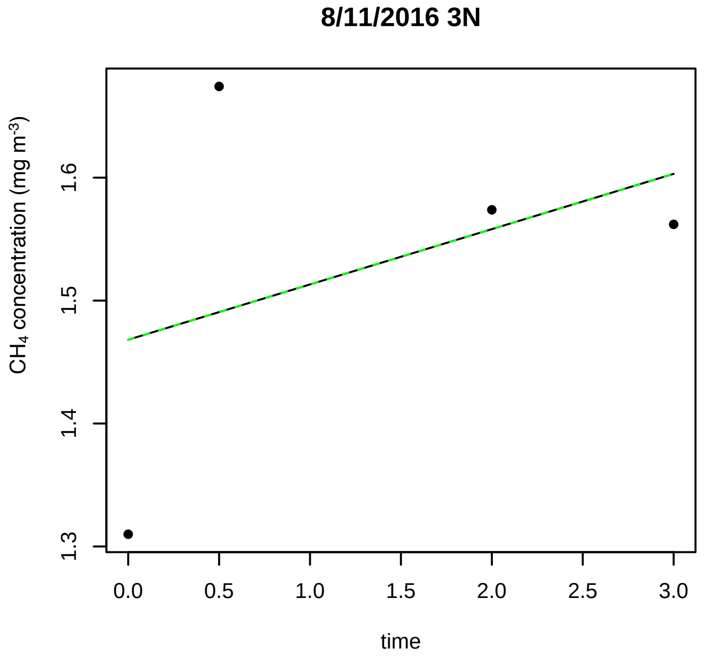
8/11/2016 3N
CH4 concentration (mg m-3)
1.3
1.4
1.5
1.6
0.0
0.5
1.0
1.5
2.0
2.5
3.0
time

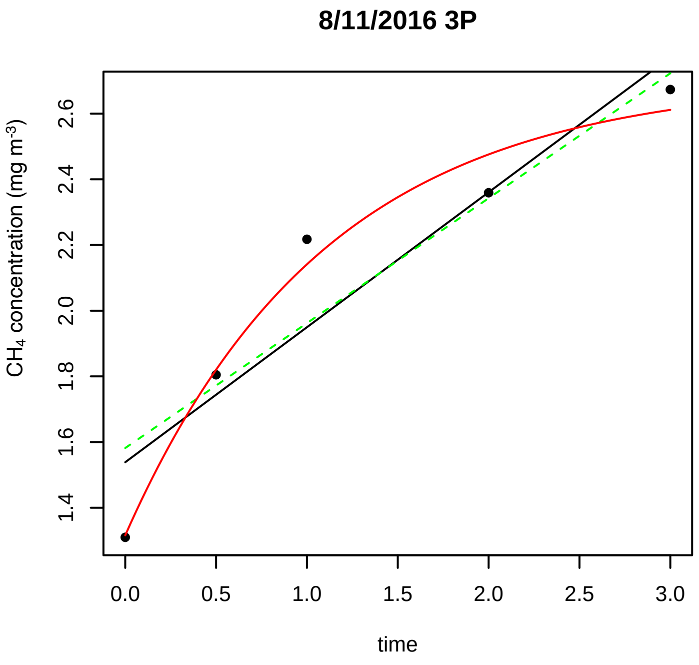
8/11/2016 3P
CH4 concentration (mg m-3)
1.4 1.6 1.8 2.0 2.2 2.4 2.6
0.0 0.5 1.0 1.5 2.0 2.5 3.0
time

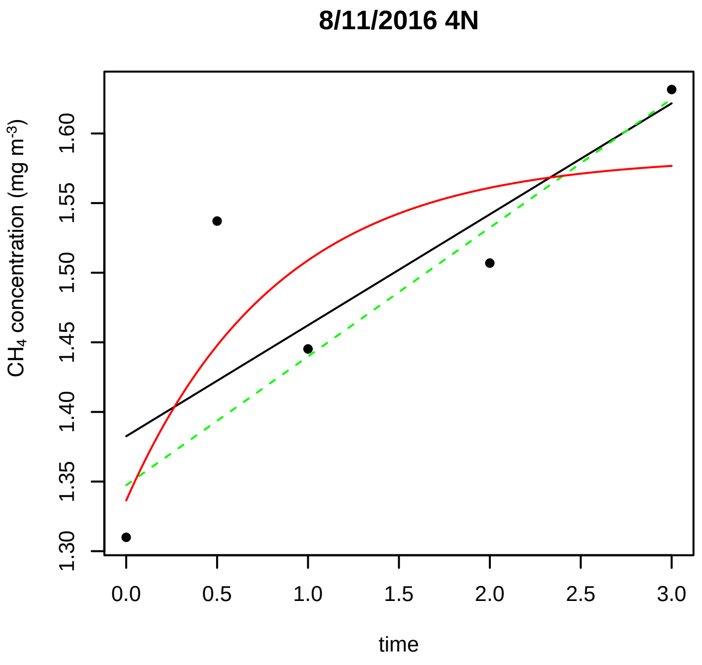
8/11/2016 4N
CH4 concentration (mg m-3)
1.30 1.35 1.40 1.45 1.50 1.55 1.60
0.0 0.5 1.0 1.5 2.0 2.5 3.0
time

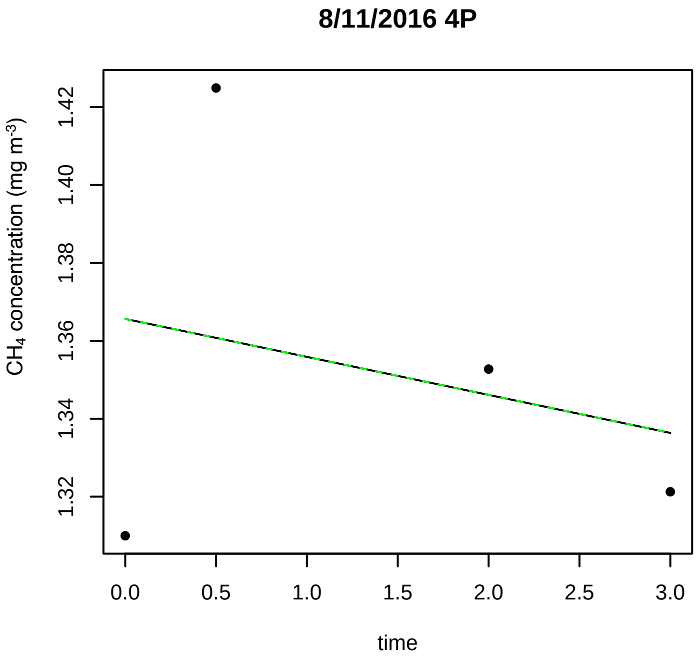
8/11/2016 4P
CH4 concentration (mg m-3)
1.32 1.34 1.36 1.38 1.40 1.42
0.0 0.5 1.0 1.5 2.0 2.5 3.0
time

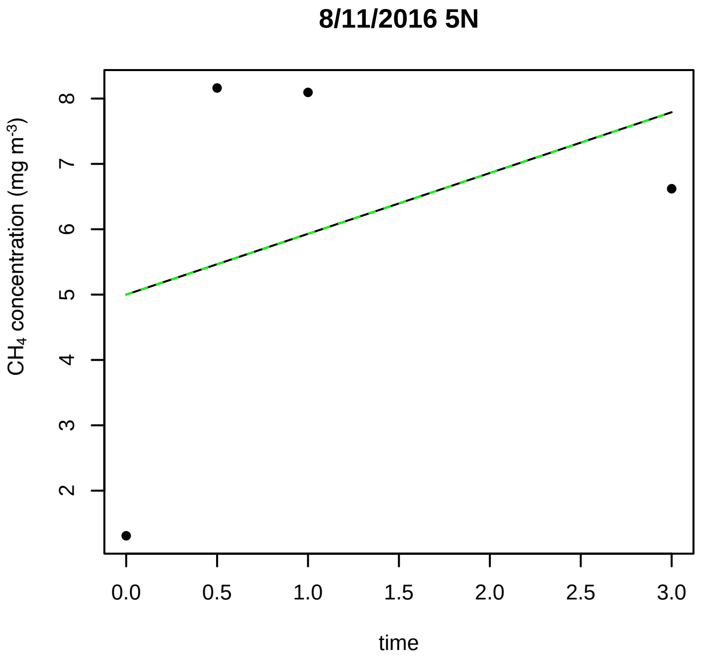
8/11/2016 5N
CH4 concentration (mg m-3)
2 3 4 5 6 7 8
0.0 0.5 1.0 1.5 2.0 2.5 3.0
time

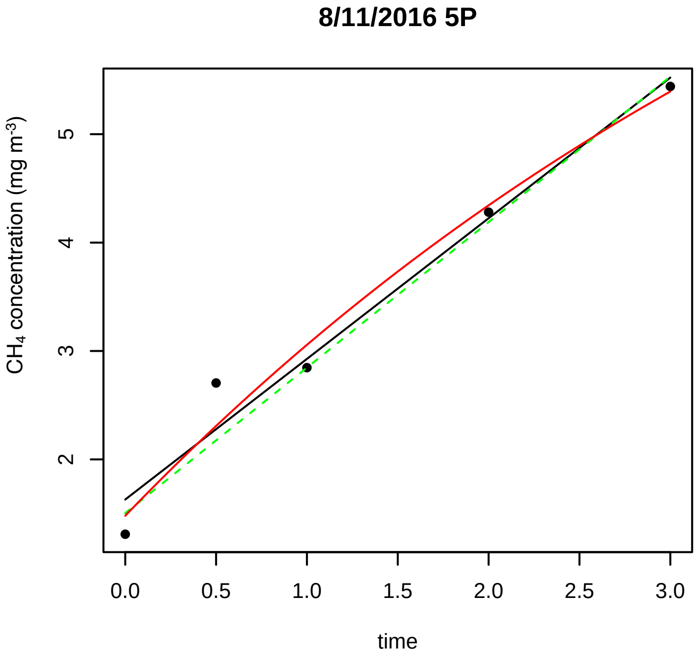
8/11/2016 5P
CH4 concentration (mg m-3)
2 3 4 5
0.0 0.5 1.0 1.5 2.0 2.5 3.0
time

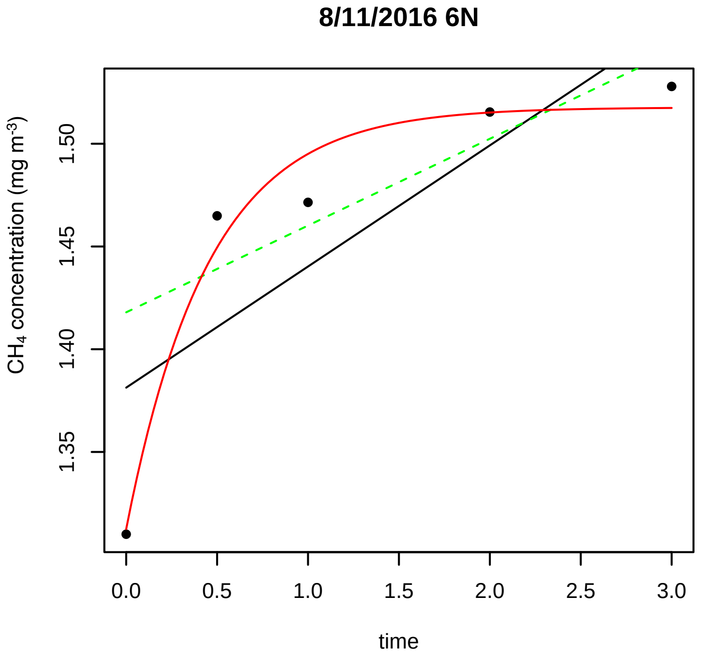
8/11/2016 6N
CH4 concentration (mg m-3)
1.35 1.40 1.45 1.50
0.0 0.5 1.0 1.5 2.0 2.5 3.0
time

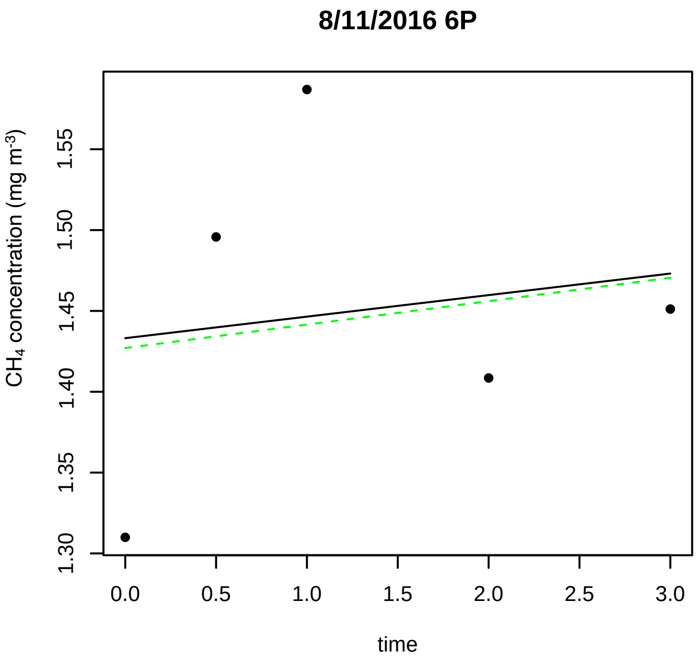
8/11/2016 6P
CH4 concentration (mg m-3)
1.30
1.35
1.40
1.45
1.50
1.55
0.0
0.5
1.0
1.5
2.0
2.5
3.0
time

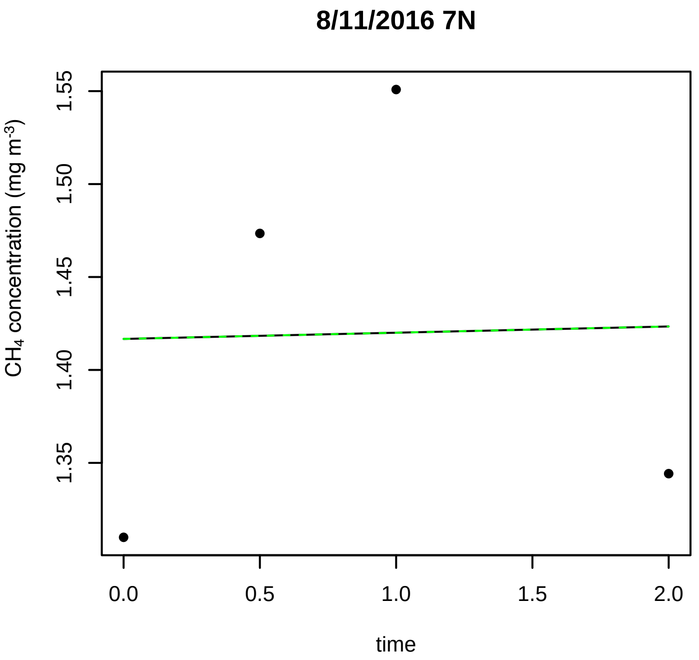
8/11/2016 7N
CH4 concentration (mg m-3)
1.35
1.40
1.45
1.50
1.55
0.0
0.5
1.0
1.5
2.0
time

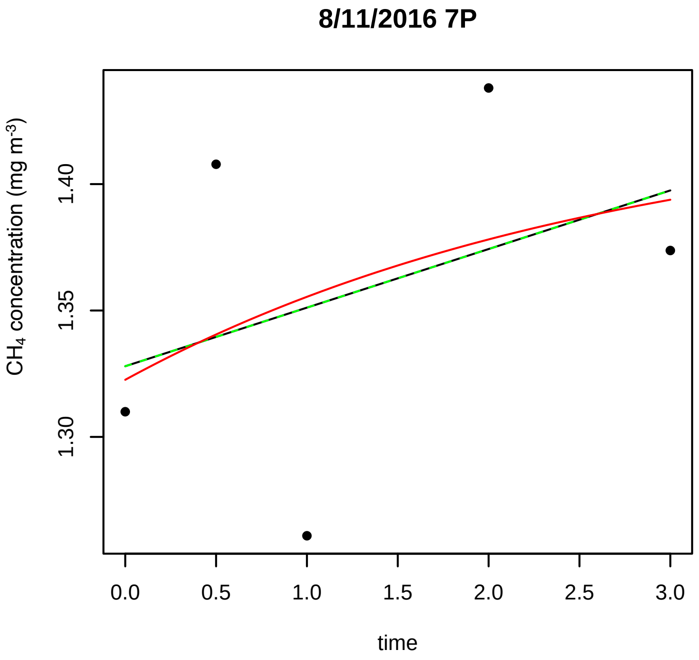
8/11/2016 7P
CH4 concentration (mg m-3)
1.30
1.35
1.40
0.0
0.5
1.0
1.5
2.0
2.5
3.0
time

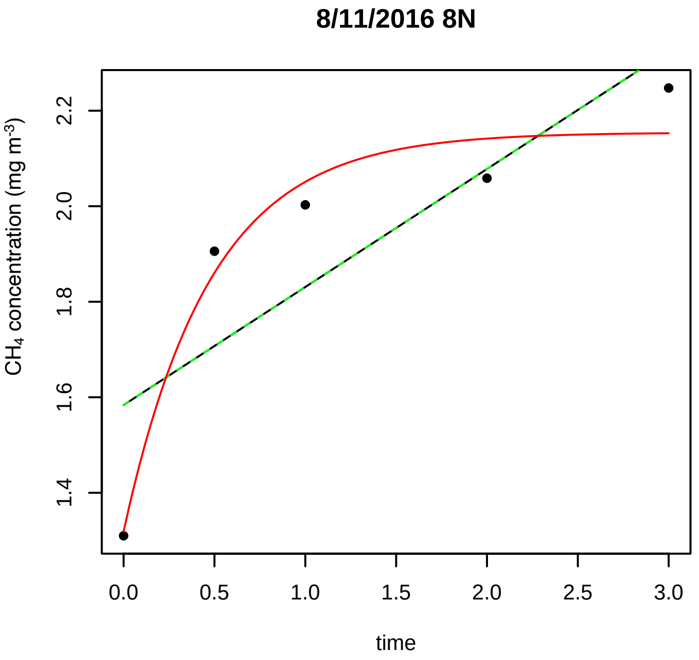
8/11/2016 8N
CH4 concentration (mg m-3)
1.4
1.6
1.8
2.0
2.2
0.0
0.5
1.0
1.5
2.0
2.5
3.0
time

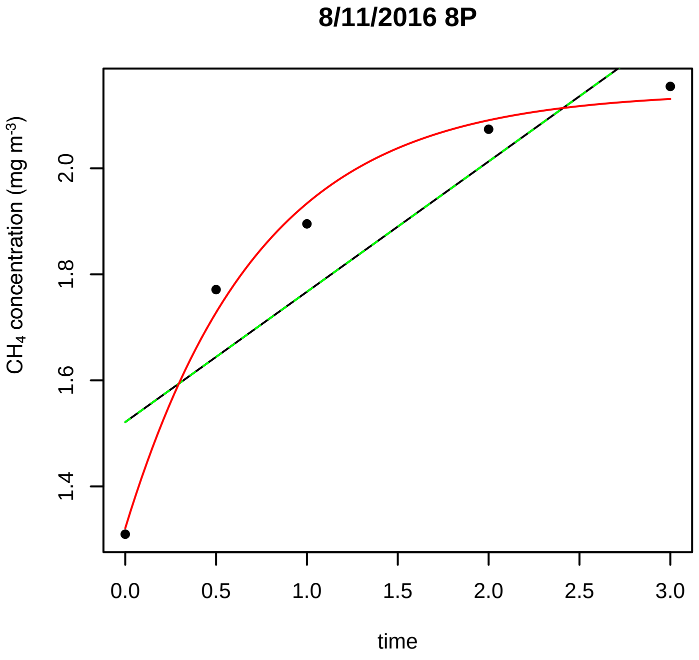
8/11/2016 8P
CH4 concentration (mg m-3)
1.4
1.6
1.8
2.0
0.0
0.5
1.0
1.5
2.0
2.5
3.0
time

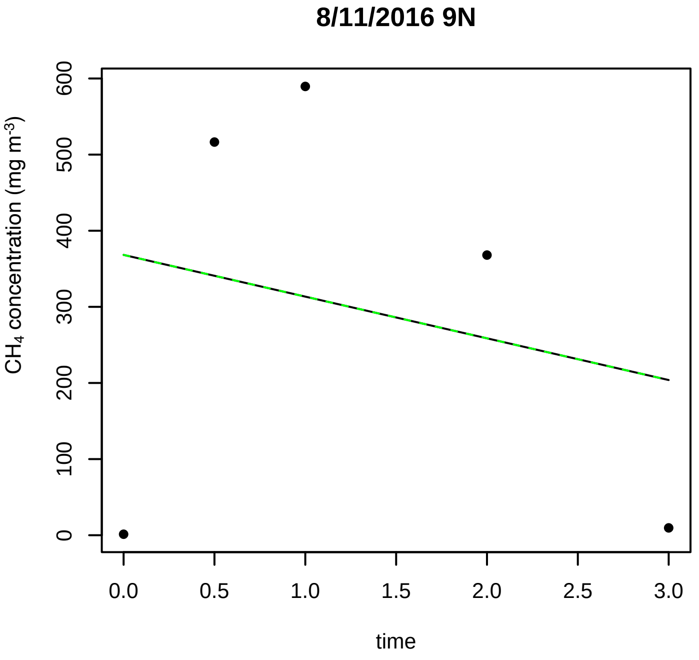
8/11/2016 9N
CH4 concentration (mg m-3)
0
100
200
300
400
500
600
0.0
0.5
1.0
1.5
2.0
2.5
3.0
time

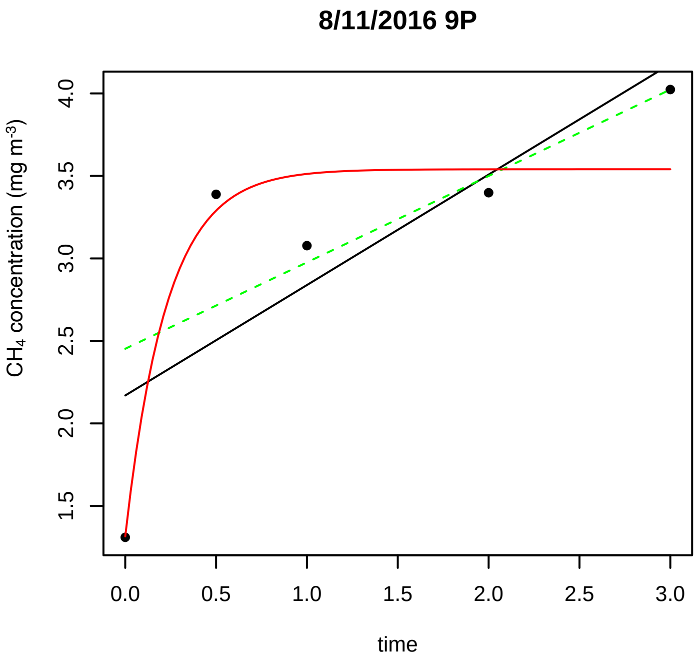

8/11/2016 9P
CH4 concentration (mg m-3)
4.0
3.5
3.0
2.5
2.0
1.5
0.0
0.5
1.0
1.5
2.0
2.5
3.0
time

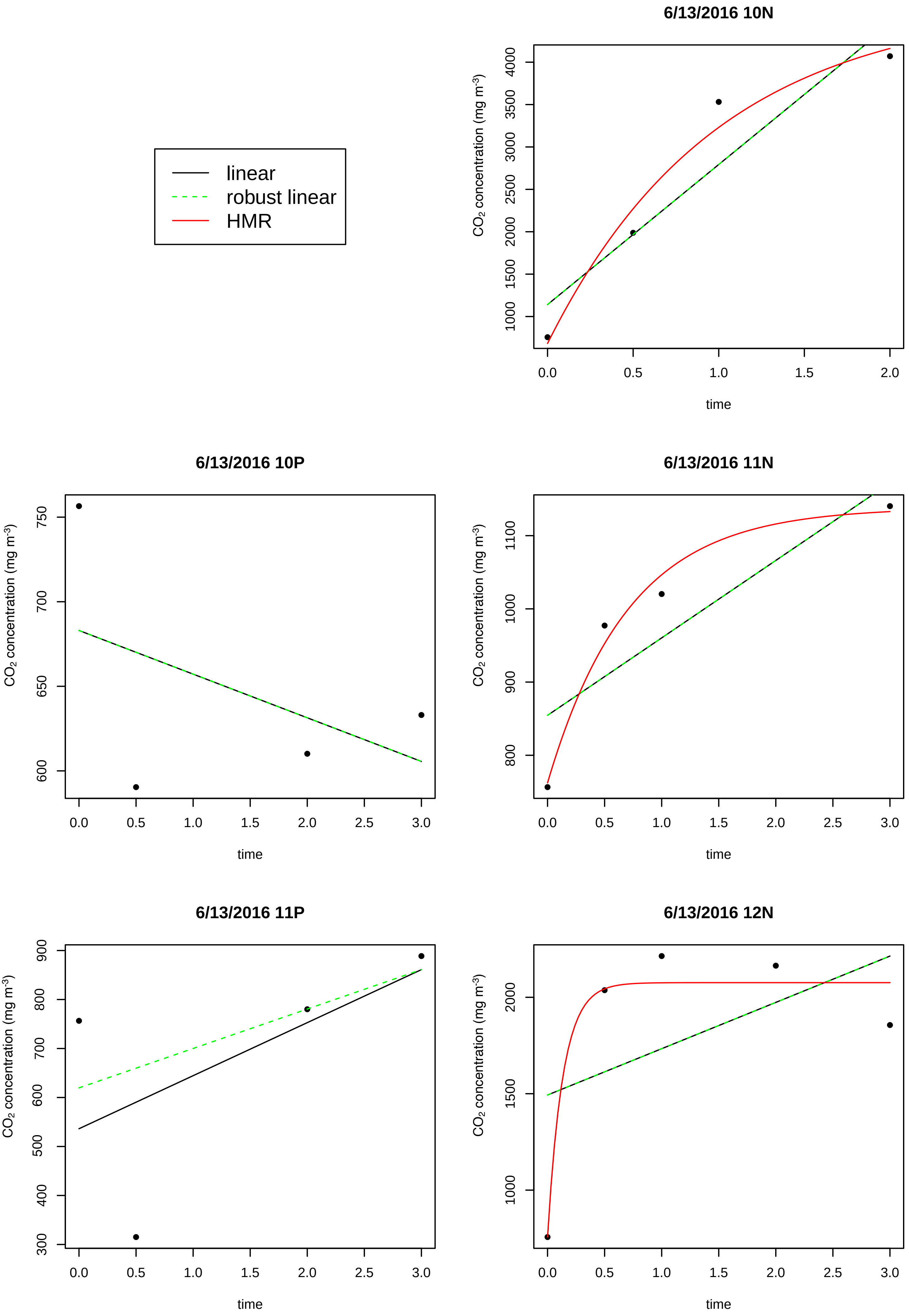

linear
robust linear
HMR
6/13/2016 10N
CO2 concentration (mg m-3)
1000
1500
2000
2500
3000
3500
4000
0.0
0.5
1.0
1.5
2.0
time
6/13/2016 10P
CO2 concentration (mg m-3)
600
650
700
750
0.0
0.5
1.0
1.5
2.0
2.5
3.0
time
6/13/2016 11N
CO2 concentration (mg m-3)
800
900
1000
1100
0.0
0.5
1.0
1.5
2.0
2.5
3.0
time
6/13/2016 11P
CO2 concentration (mg m-3)
300
400
500
600
700
800
900
0.0
0.5
1.0
1.5
2.0
2.5
3.0
time
6/13/2016 12N
CO2 concentration (mg m-3)
1000
1500
2000
0.0
0.5
1.0
1.5
2.0
2.5
3.0
time

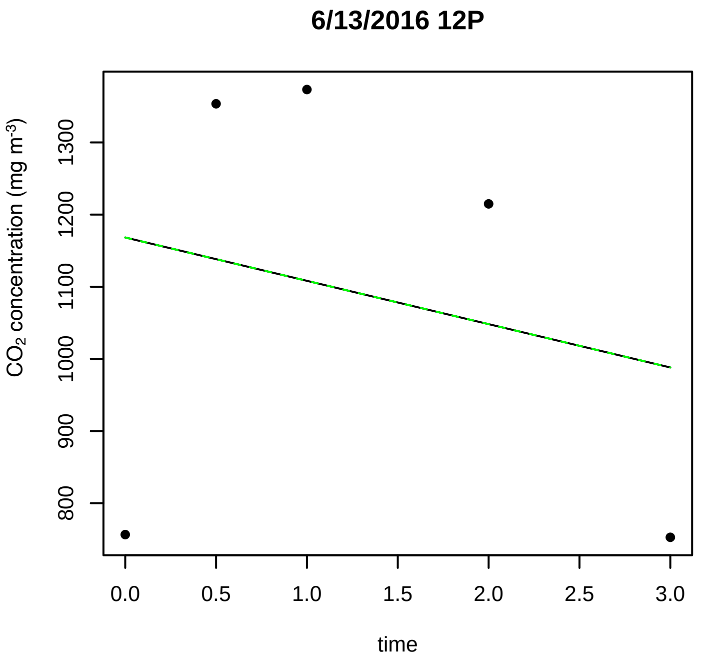
6/13/2016 12P
CO2 concentration (mg m-3)
800
900
1000
1100
1200
1300
0.0
0.5
1.0
1.5
2.0
2.5
3.0
time

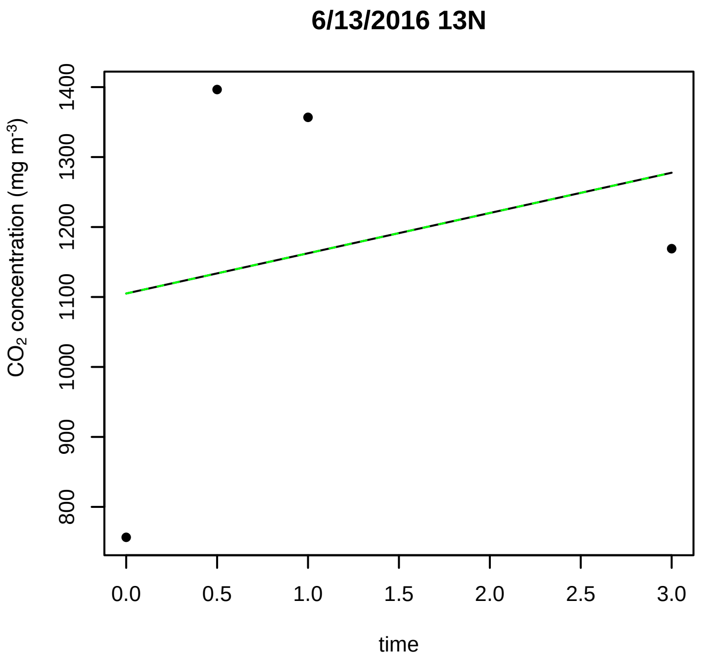
6/13/2016 13N
CO2 concentration (mg m-3)
800
900
1000
1100
1200
1300
1400
0.0
0.5
1.0
1.5
2.0
2.5
3.0
time

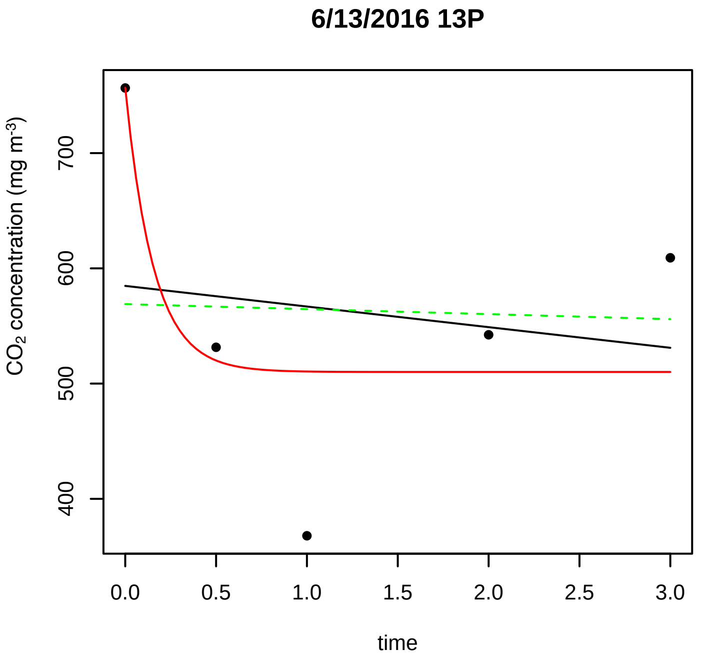
6/13/2016 13P
CO2 concentration (mg m-3)
400
500
600
700
0.0
0.5
1.0
1.5
2.0
2.5
3.0
time

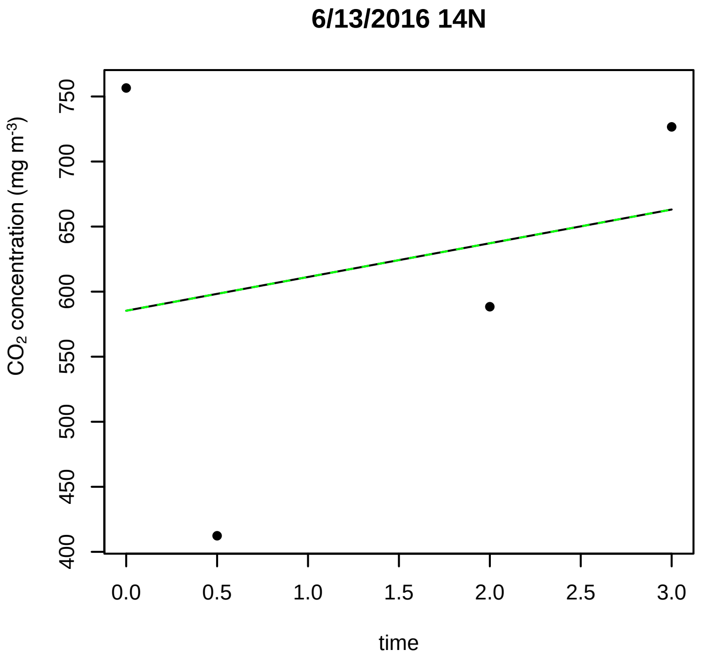
6/13/2016 14N
CO2 concentration (mg m-3)
400
450
500
550
600
650
700
750
0.0
0.5
1.0
1.5
2.0
2.5
3.0
time

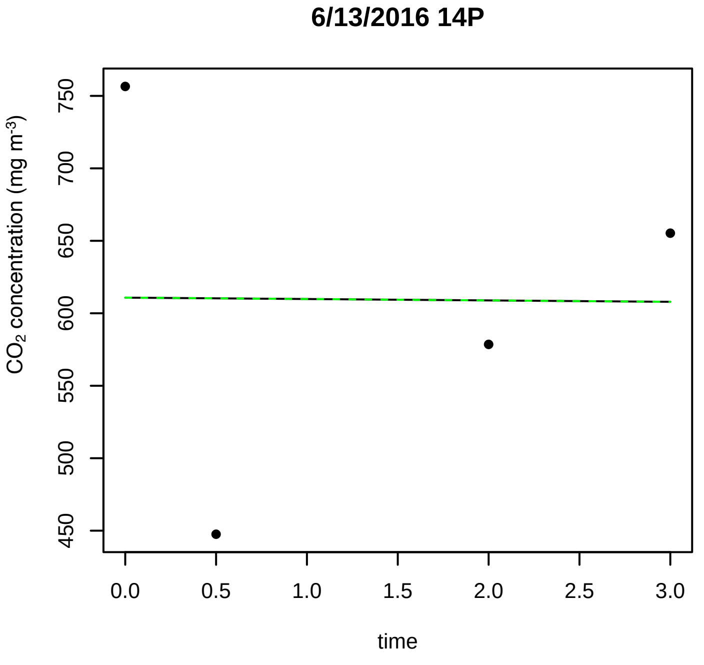
6/13/2016 14P
CO2 concentration (mg m-3)
450
500
550
600
650
700
750
0.0
0.5
1.0
1.5
2.0
2.5
3.0
time

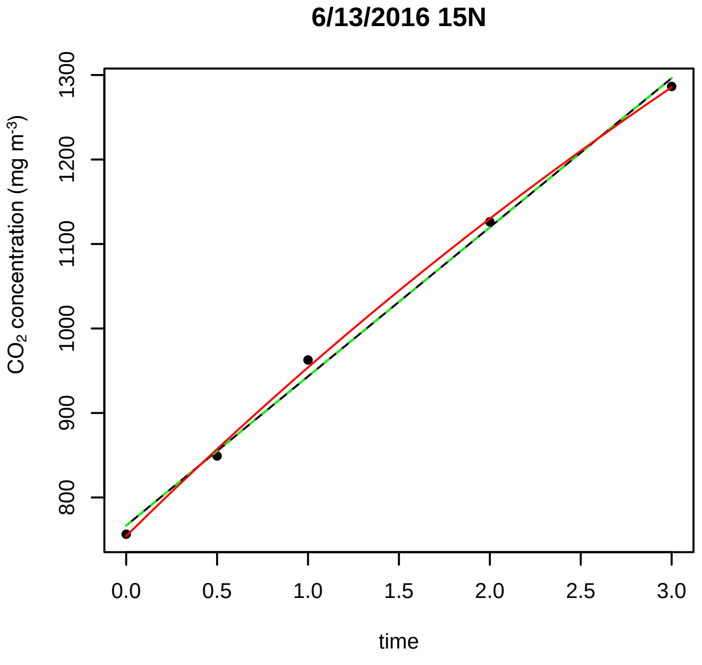
6/13/2016 15N
CO2 concentration (mg m-3)
800
900
1000
1100
1200
1300
0.0
0.5
1.0
1.5
2.0
2.5
3.0
time

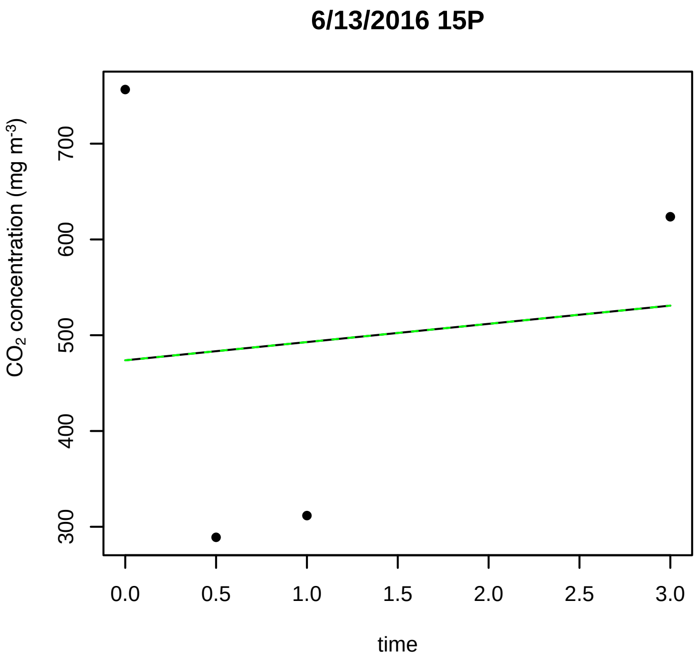
6/13/2016 15P
CO2 concentration (mg m-3)
300
400
500
600
700
0.0
0.5
1.0
1.5
2.0
2.5
3.0
time

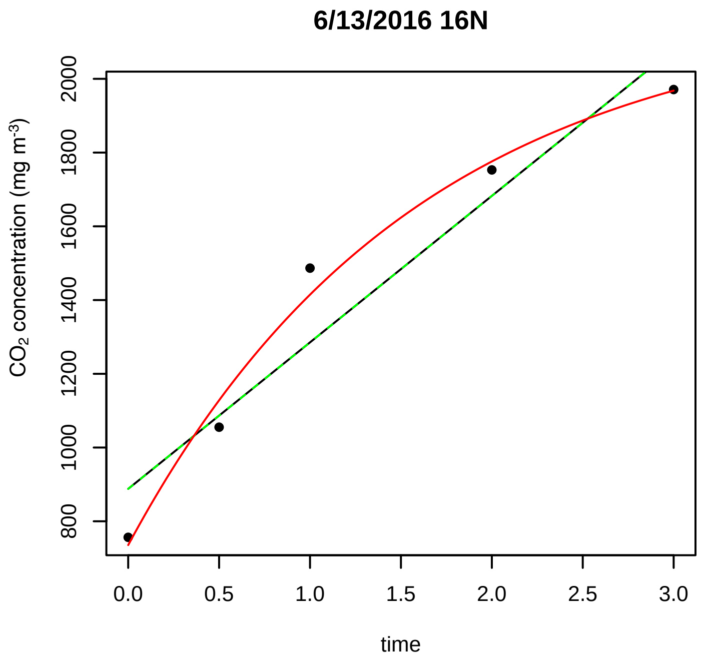
6/13/2016 16N
CO2 concentration (mg m-3)
800
1000
1200
1400
1600
1800
2000
0.0
0.5
1.0
1.5
2.0
2.5
3.0
time

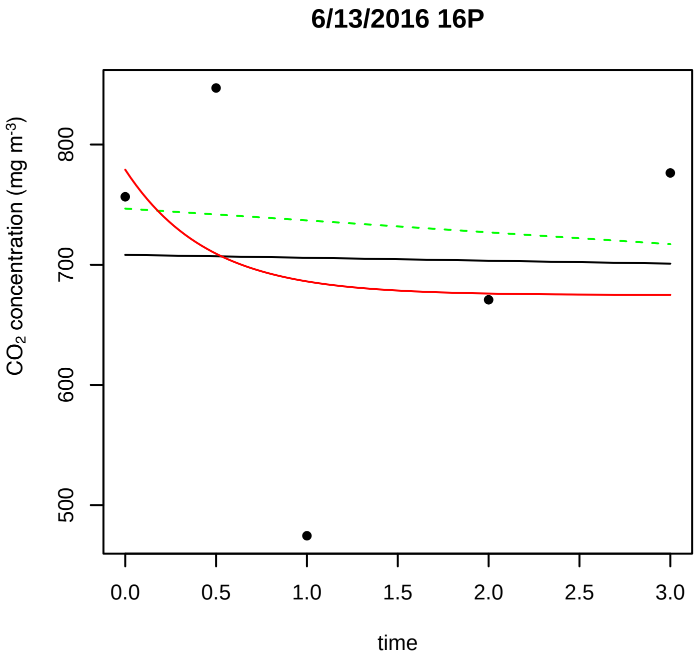
6/13/2016 16P
CO2 concentration (mg m-3)
500
600
700
800
0.0
0.5
1.0
1.5
2.0
2.5
3.0
time

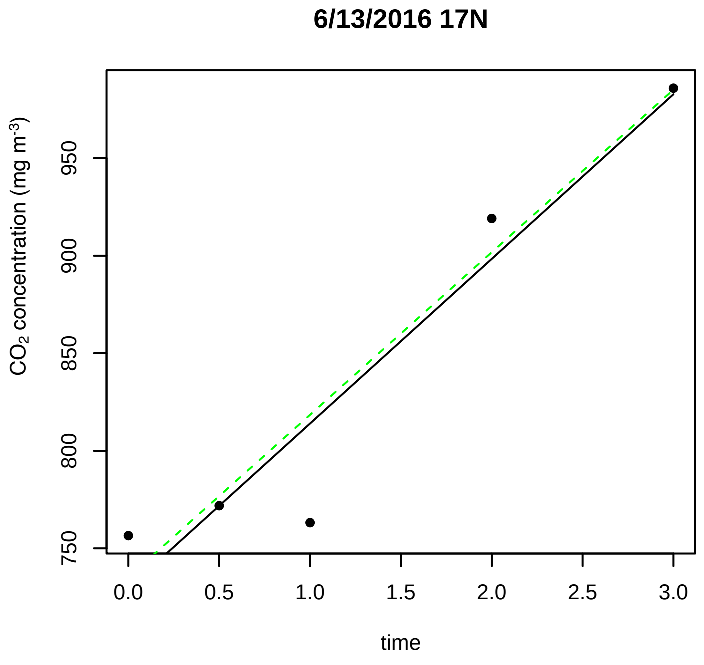
6/13/2016 17N
CO2 concentration (mg m-3)
750
800
850
900
950
0.0
0.5
1.0
1.5
2.0
2.5
3.0
time

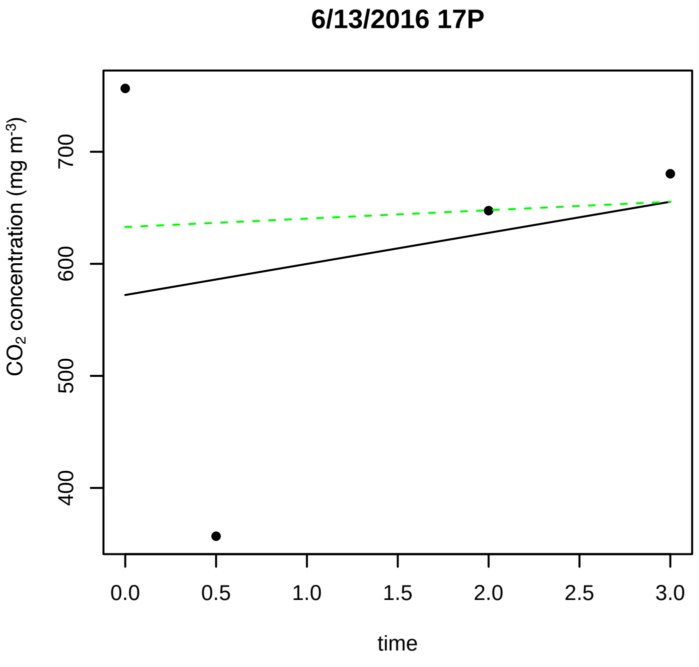
6/13/2016 17P
CO2 concentration (mg m-3)
400
500
600
700
0.0
0.5
1.0
1.5
2.0
2.5
3.0
time

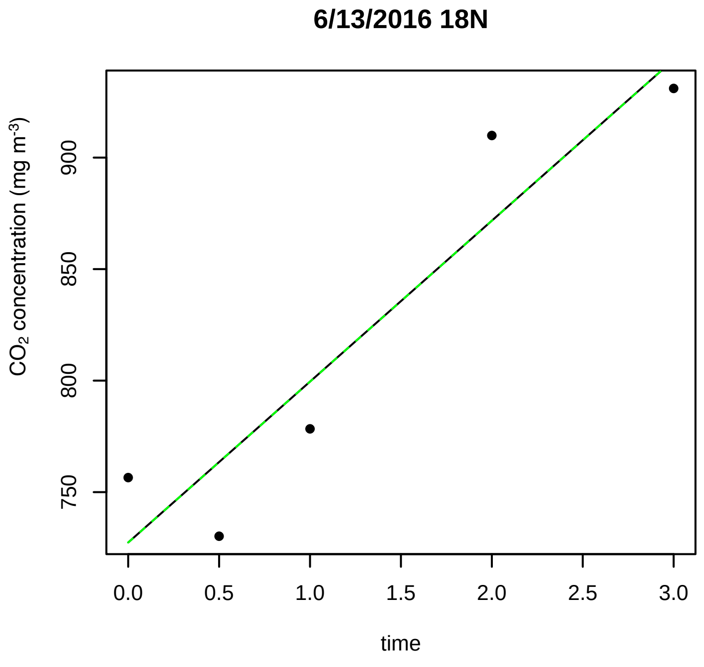
6/13/2016 18N
CO2 concentration (mg m-3)
750
800
850
900
0.0
0.5
1.0
1.5
2.0
2.5
3.0
time

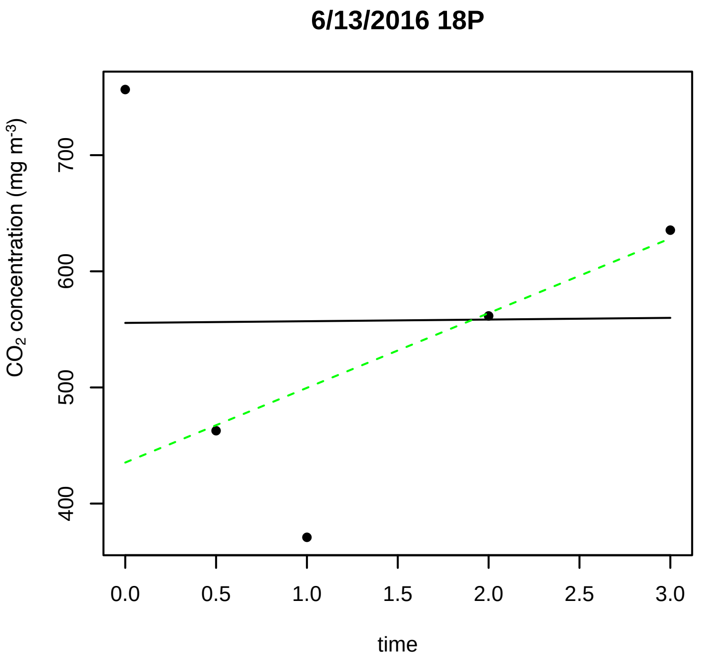
6/13/2016 18P
CO2 concentration (mg m-3)
400
500
600
700
0.0
0.5
1.0
1.5
2.0
2.5
3.0
time

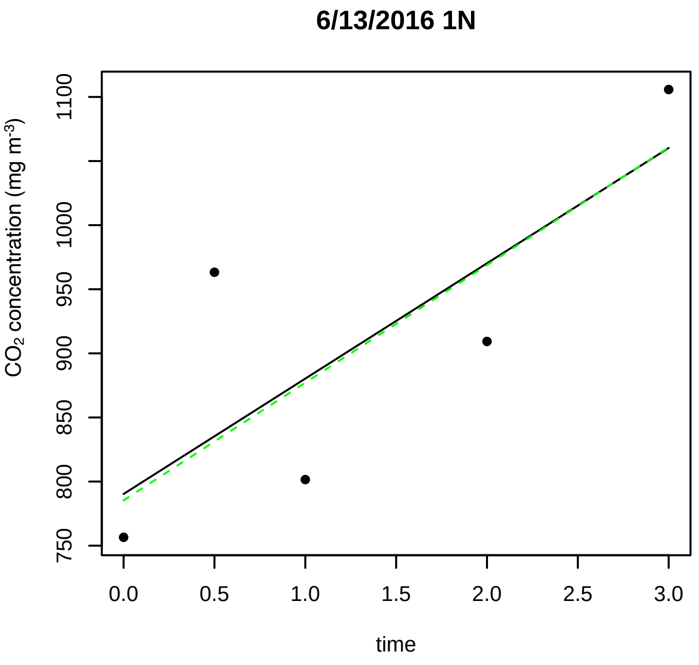
6/13/2016 1N
CO2 concentration (mg m-3)
750
800
850
900
950
1000
1100
0.0
0.5
1.0
1.5
2.0
2.5
3.0
time

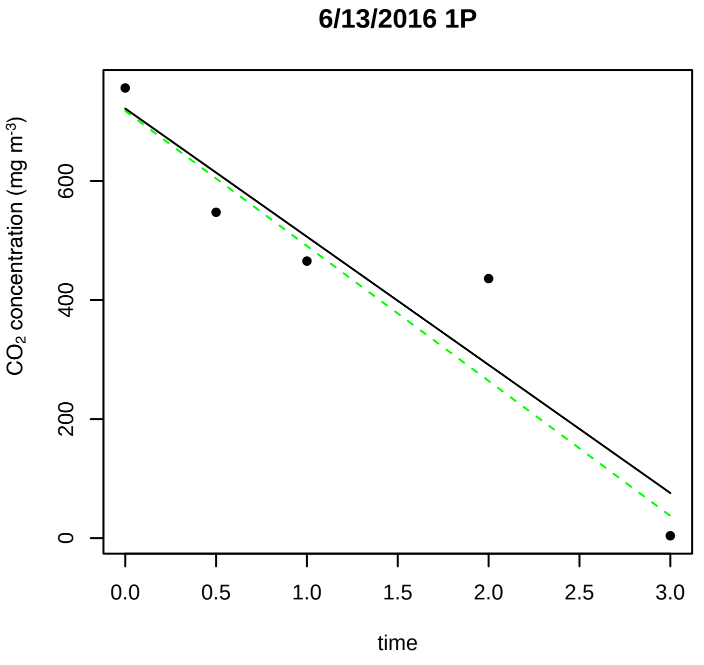
6/13/2016 1P
CO2 concentration (mg m-3)
0
200
400
600
0.0
0.5
1.0
1.5
2.0
2.5
3.0
time

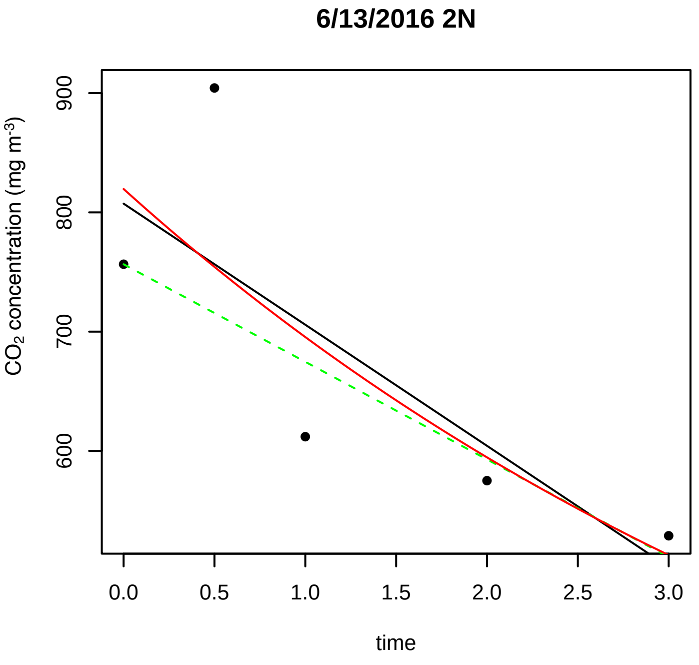
6/13/2016 2N
CO2 concentration (mg m-3)
600
700
800
900
0.0
0.5
1.0
1.5
2.0
2.5
3.0
time

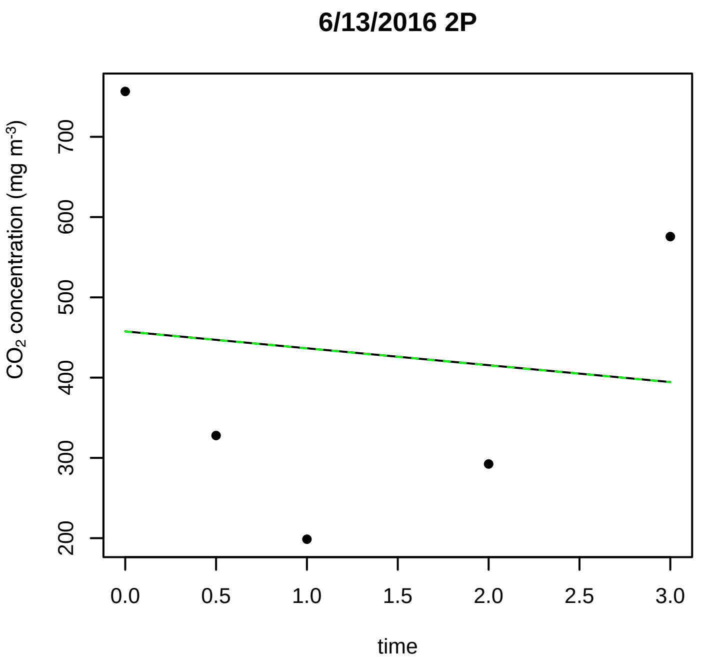
6/13/2016 2P
CO2 concentration (mg m-3)
200
300
400
500
600
700
0.0
0.5
1.0
1.5
2.0
2.5
3.0
time

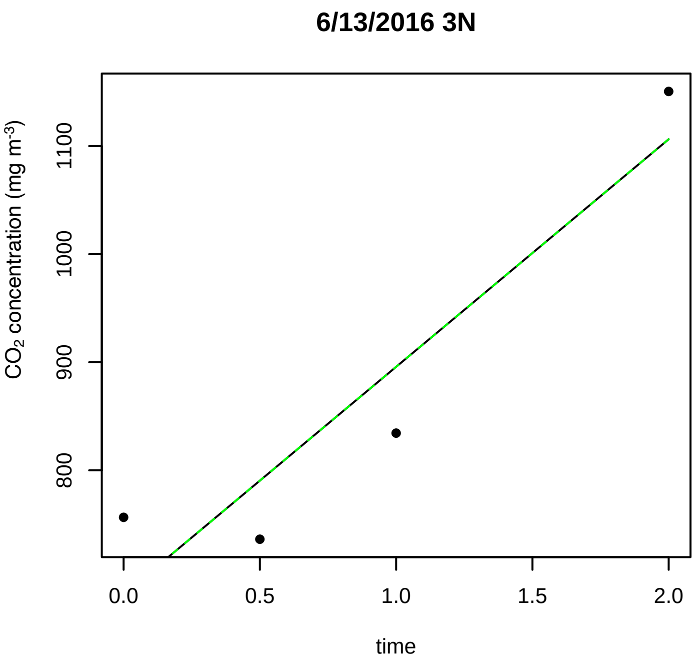
6/13/2016 3N
CO2 concentration (mg m-3)
800
900
1000
1100
0.0
0.5
1.0
1.5
2.0
time

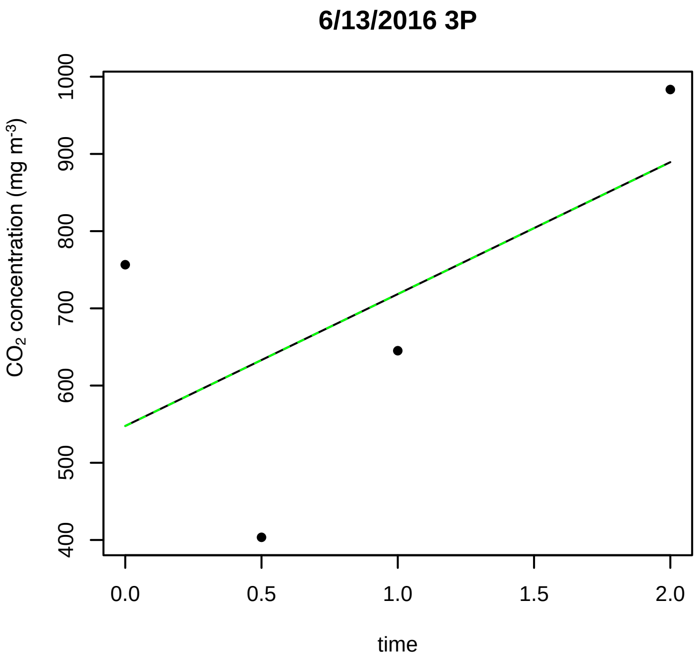
6/13/2016 3P
CO2 concentration (mg m-3)
400 500 600 700 800 900 1000
time
0.0 0.5 1.0 1.5 2.0

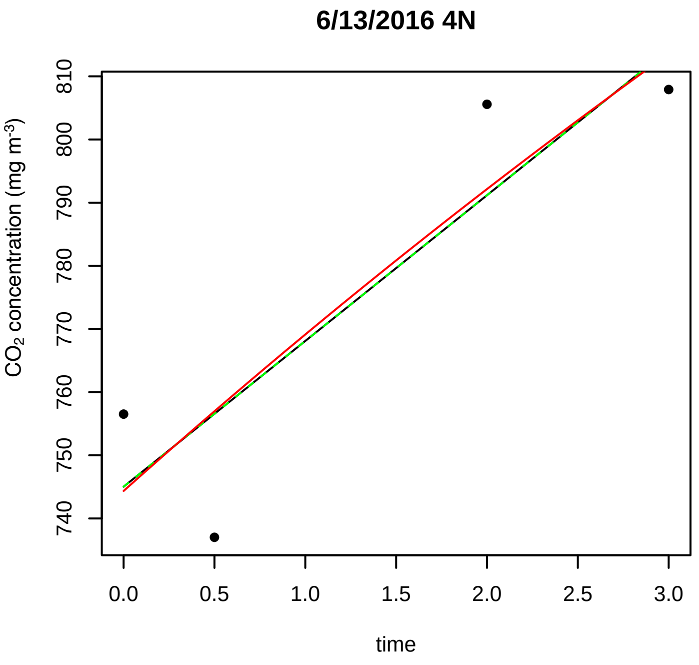
6/13/2016 4N
CO2 concentration (mg m-3)
740 750 760 770 780 790 800 810
time
0.0 0.5 1.0 1.5 2.0 2.5 3.0

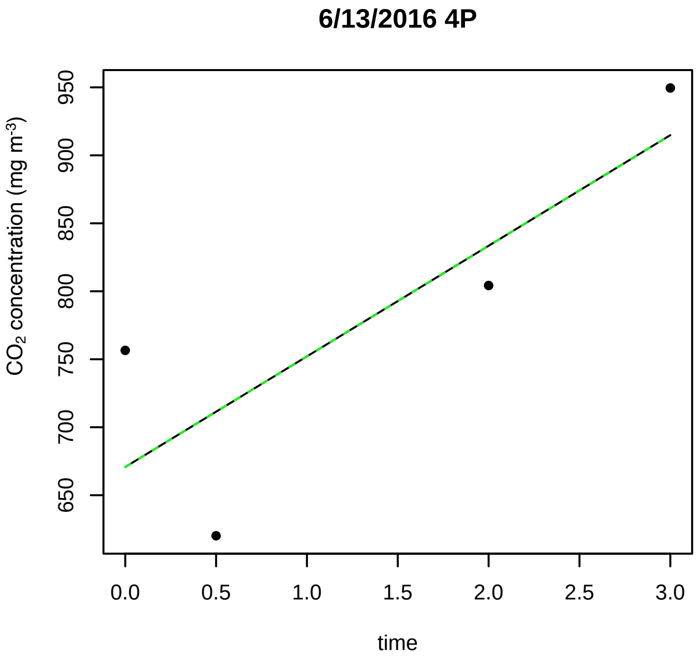
6/13/2016 4P
CO2 concentration (mg m-3)
650 700 750 800 850 900 950
time
0.0 0.5 1.0 1.5 2.0 2.5 3.0

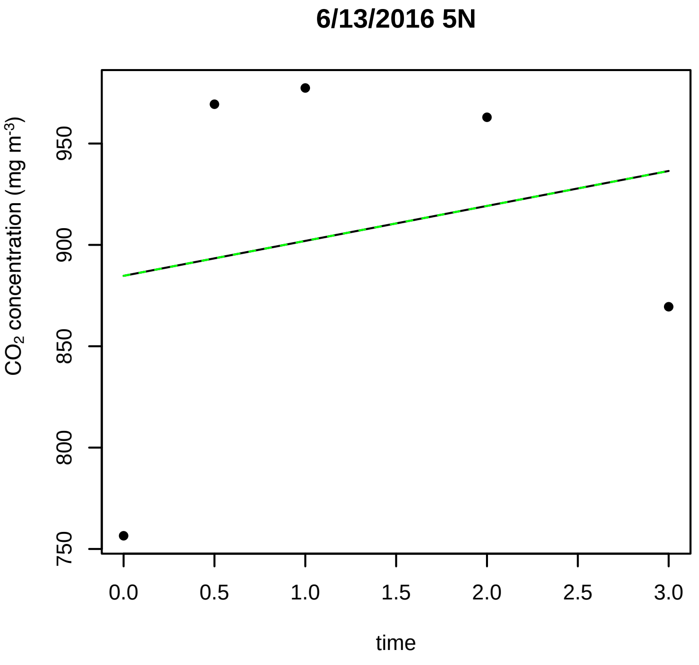
6/13/2016 5N
CO2 concentration (mg m-3)
750 800 850 900 950
time
0.0 0.5 1.0 1.5 2.0 2.5 3.0

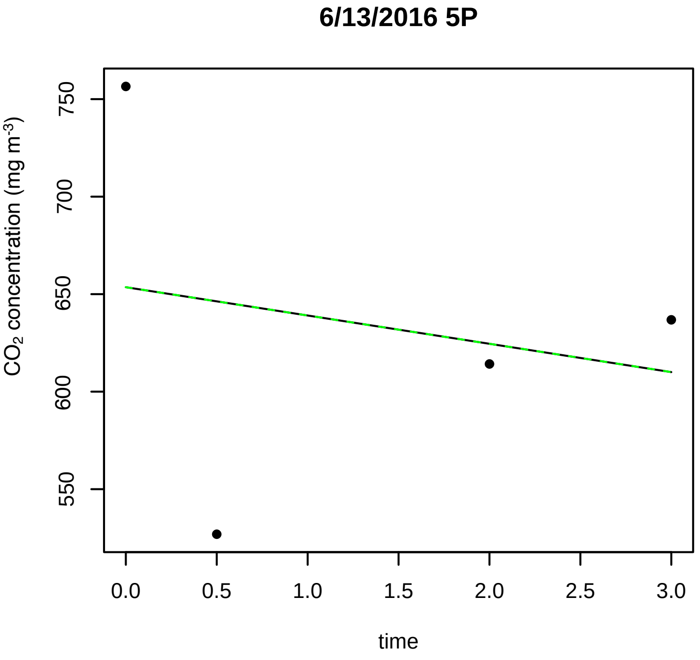
6/13/2016 5P
CO2 concentration (mg m-3)
550 600 650 700 750
time
0.0 0.5 1.0 1.5 2.0 2.5 3.0

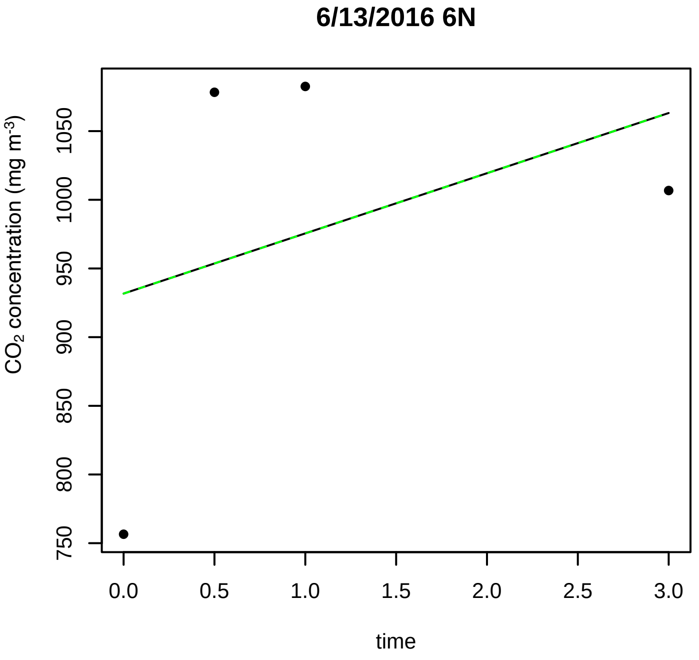
6/13/2016 6N
CO2 concentration (mg m-3)
750 800 850 900 950 1000 1050
time
0.0 0.5 1.0 1.5 2.0 2.5 3.0

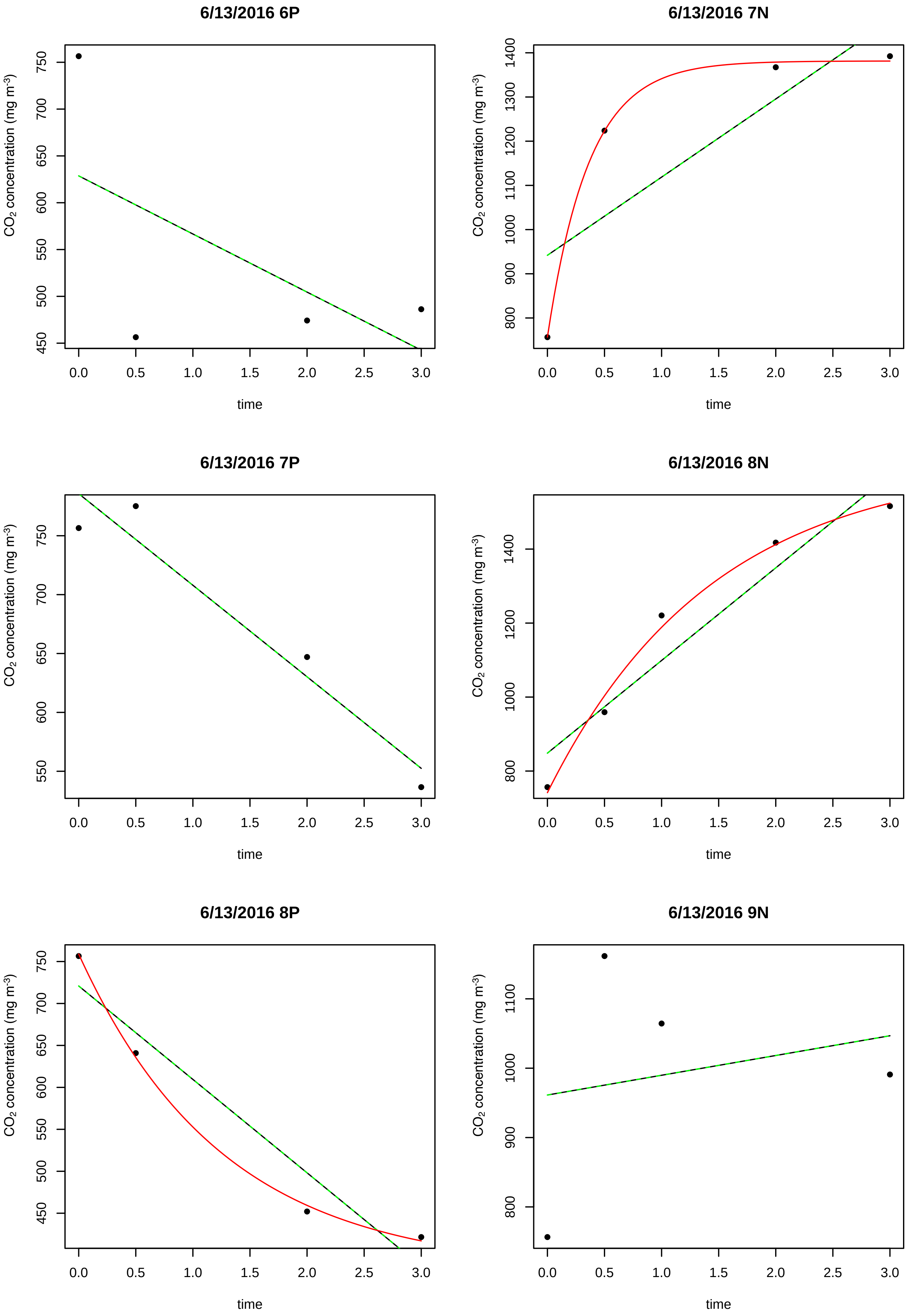

6/13/2016 6P
6/13/2016 7N
6/13/2016 7P
6/13/2016 8N
6/13/2016 8P
6/13/2016 9N
$CO_2$ concentration (mg $m^{-3}$)
time

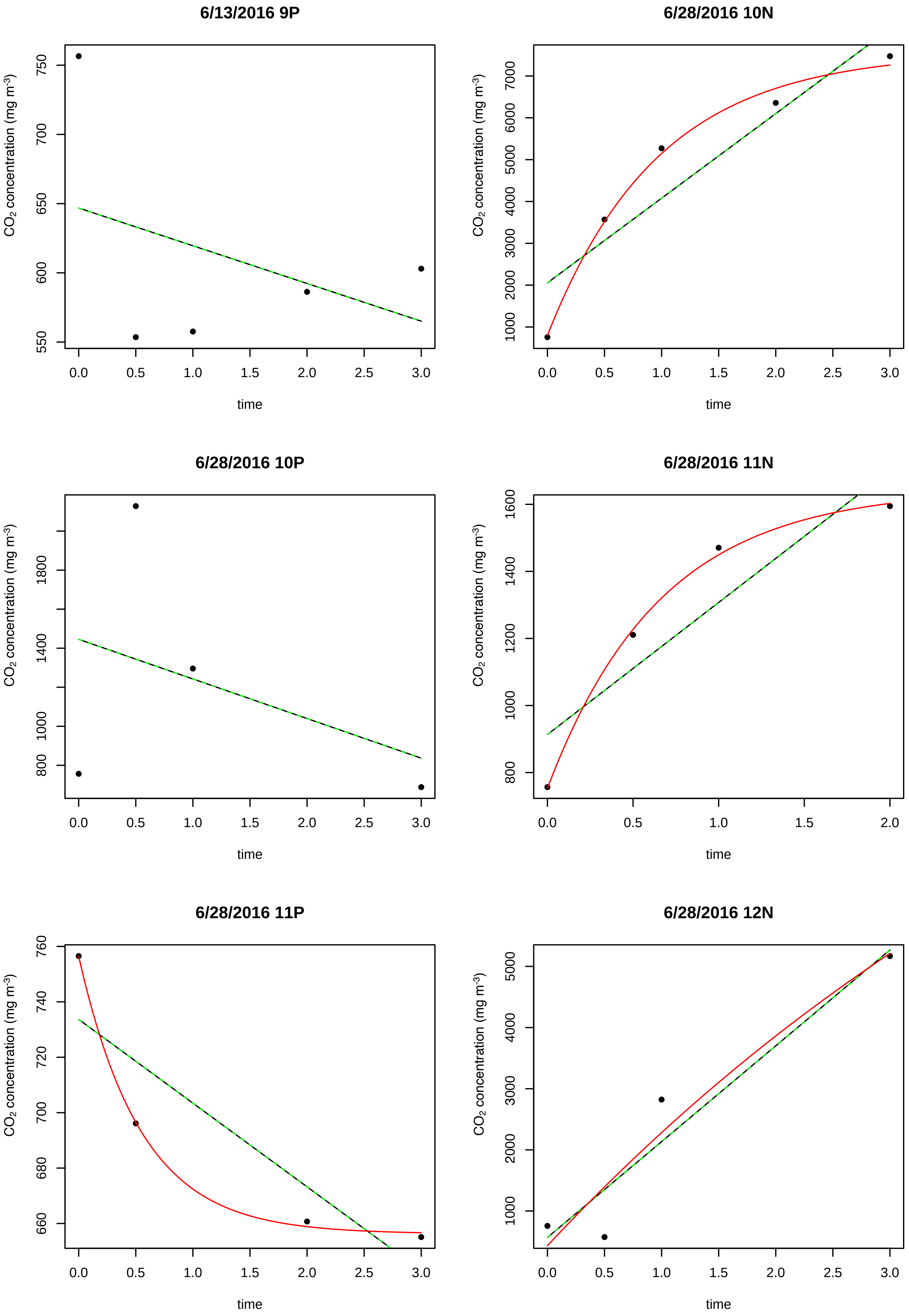
6/13/2016 9P
6/28/2016 10N
6/28/2016 10P
6/28/2016 11N
6/28/2016 11P
6/28/2016 12N
CO2 concentration (mg m-3)
time
0.0
0.5
1.0
1.5
2.0
2.5
3.0
550
600
650
700
750
1000
2000
3000
4000
5000
6000
7000
800
1400
1800
1200
1600
660
680
700
720
740
760

**6/28/2016 12P**

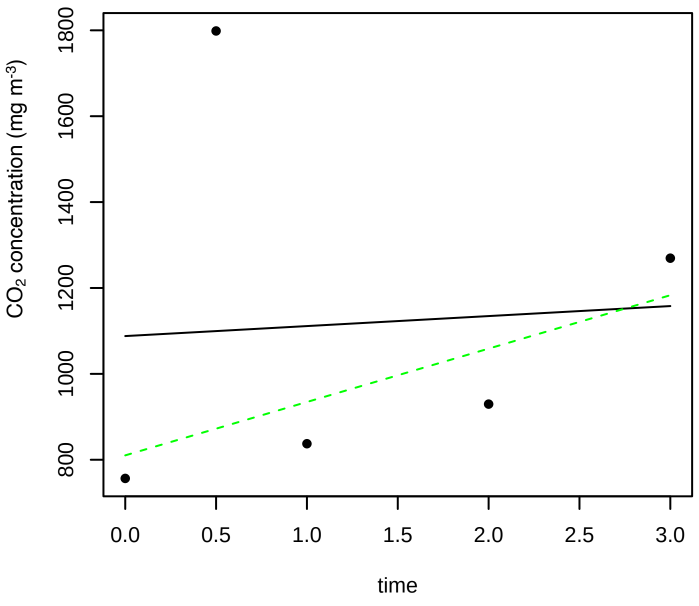


**6/28/2016 13N**

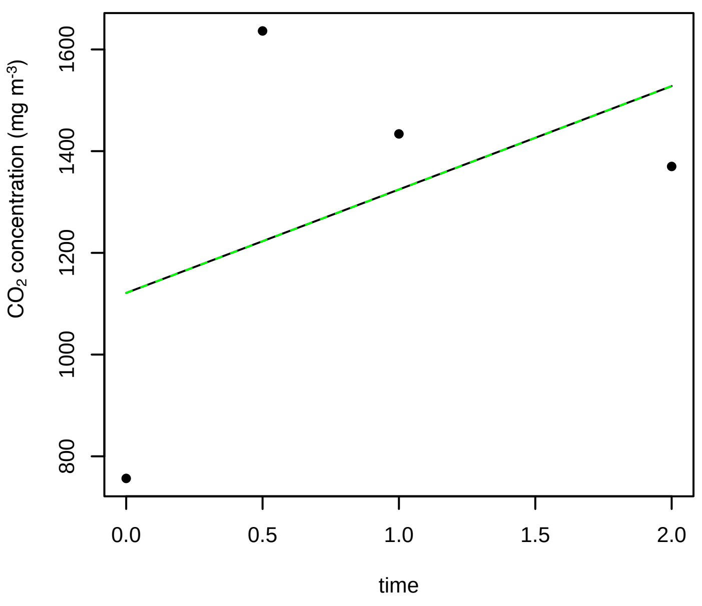


**6/28/2016 13P**

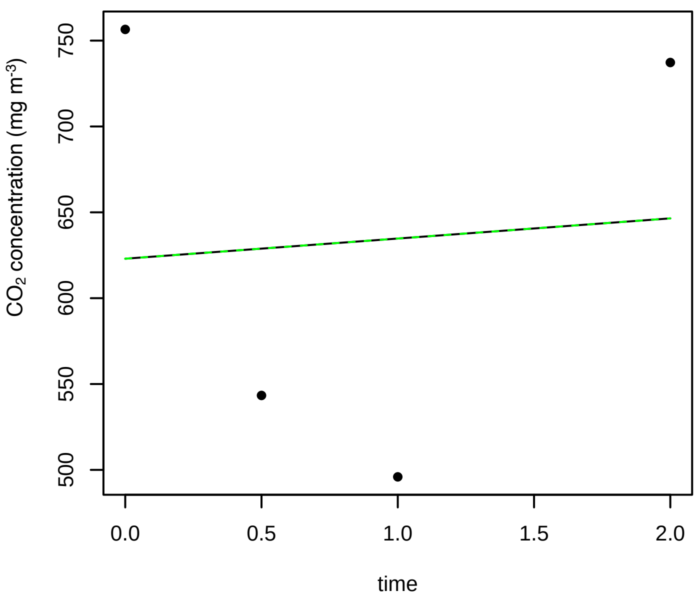


**6/28/2016 14N**

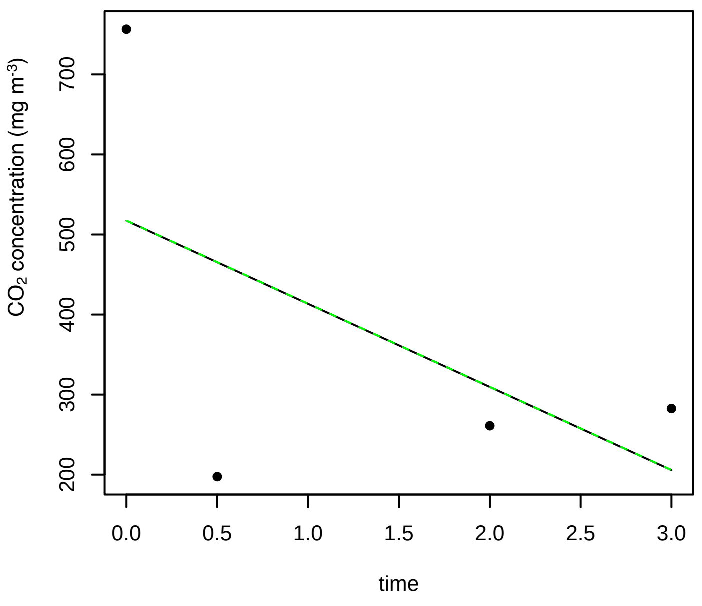


**6/28/2016 14P**

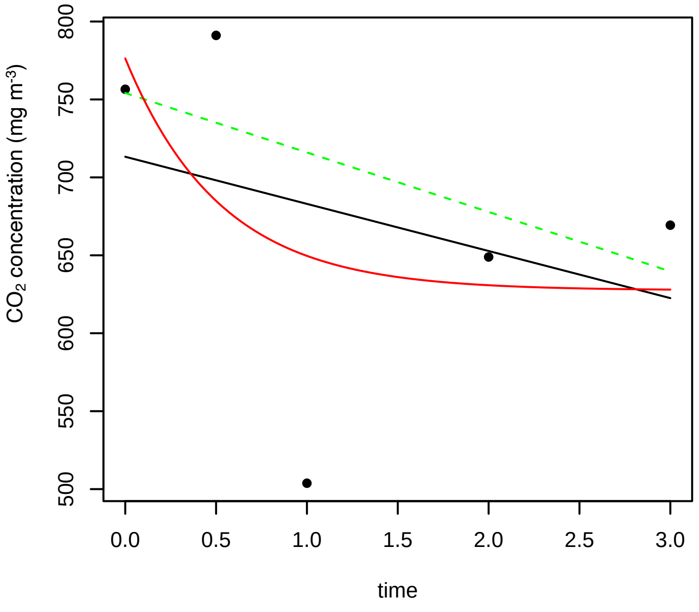


**6/28/2016 15N**

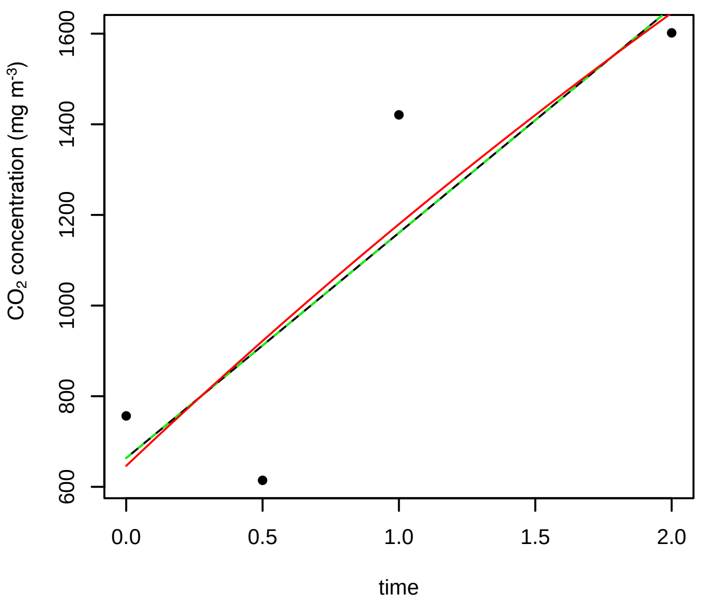

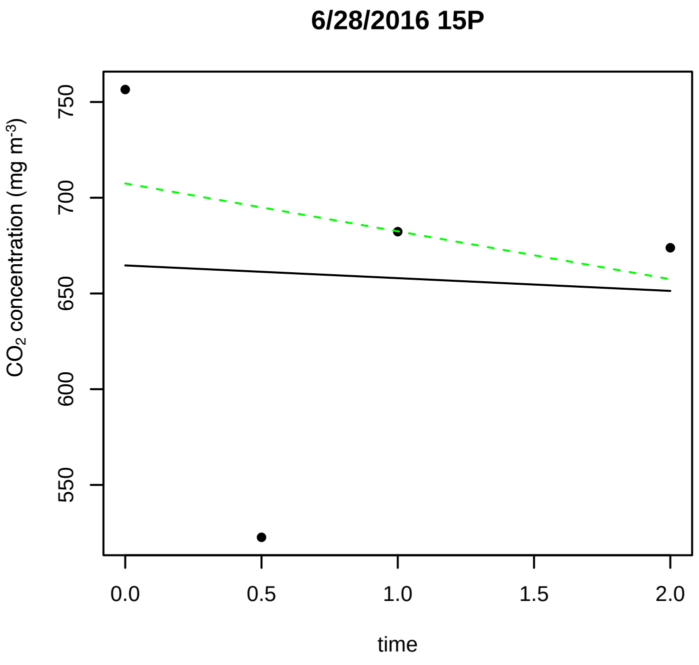
6/28/2016 15P
CO2 concentration (mg m-3)
750
700
650
600
550
0.0
0.5
1.0
1.5
2.0
time

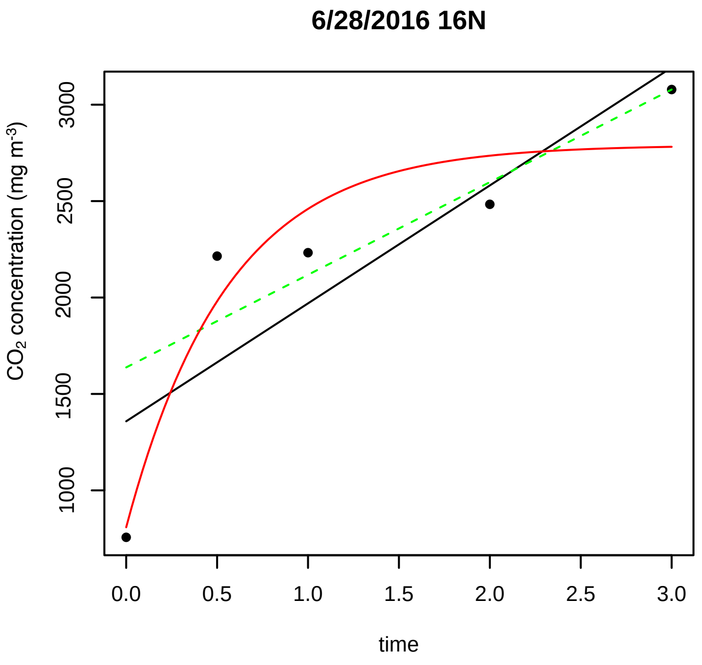
6/28/2016 16N
CO2 concentration (mg m-3)
3000
2500
2000
1500
1000
0.0
0.5
1.0
1.5
2.0
2.5
3.0
time

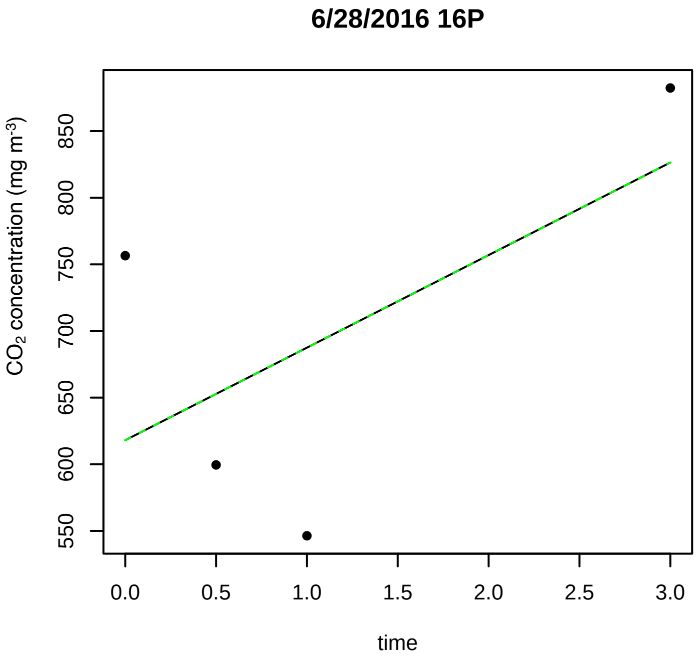
6/28/2016 16P
CO2 concentration (mg m-3)
850
800
750
700
650
600
550
0.0
0.5
1.0
1.5
2.0
2.5
3.0
time

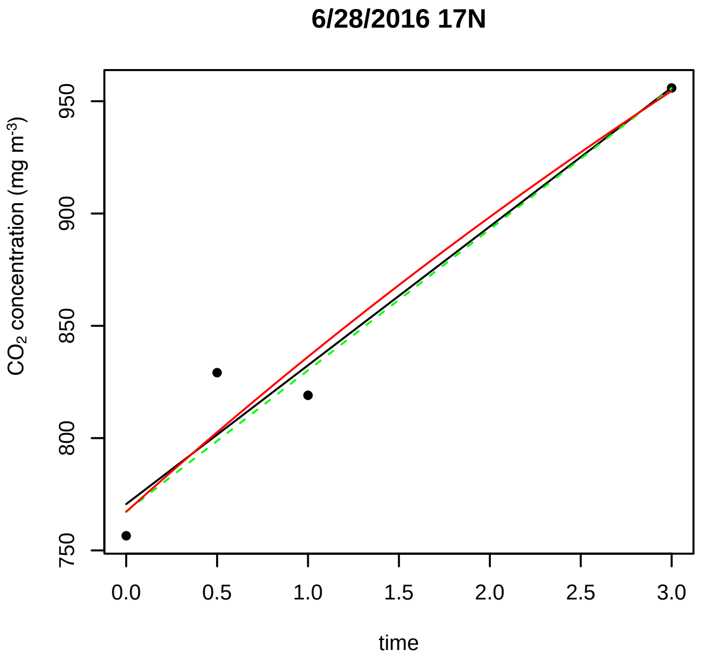
6/28/2016 17N
CO2 concentration (mg m-3)
950
900
850
800
750
0.0
0.5
1.0
1.5
2.0
2.5
3.0
time

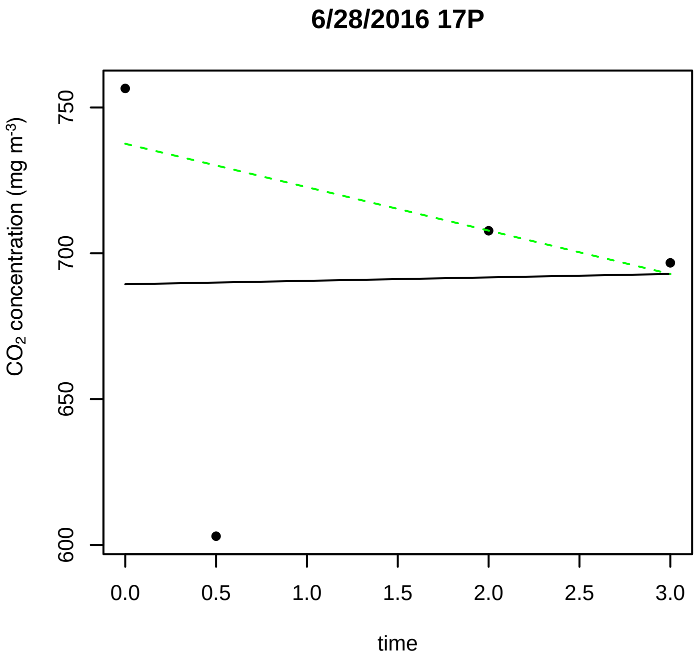
6/28/2016 17P
CO2 concentration (mg m-3)
750
700
650
600
0.0
0.5
1.0
1.5
2.0
2.5
3.0
time

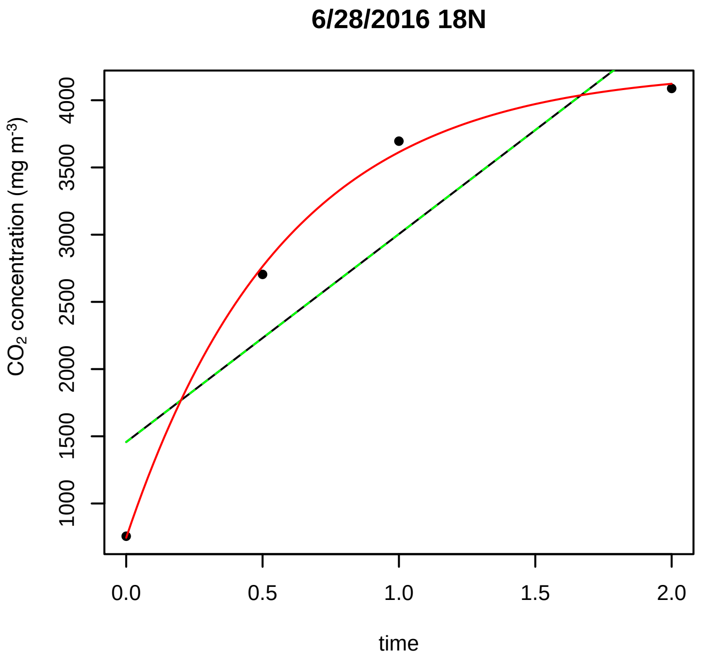
6/28/2016 18N
CO2 concentration (mg m-3)
4000
3500
3000
2500
2000
1500
1000
0.0
0.5
1.0
1.5
2.0
time

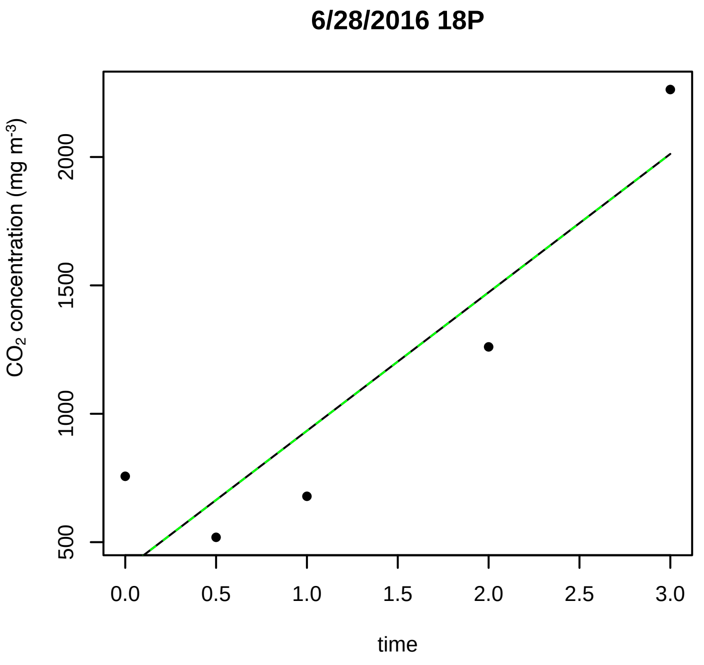
6/28/2016 18P
CO2 concentration (mg m-3)
500
1000
1500
2000
0.0
0.5
1.0
1.5
2.0
2.5
3.0
time

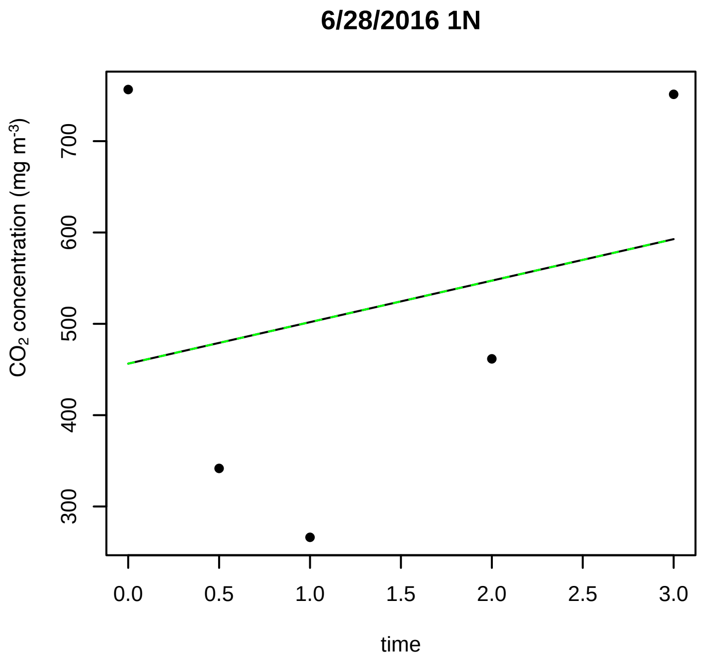
6/28/2016 1N
CO2 concentration (mg m-3)
300
400
500
600
700
0.0
0.5
1.0
1.5
2.0
2.5
3.0
time

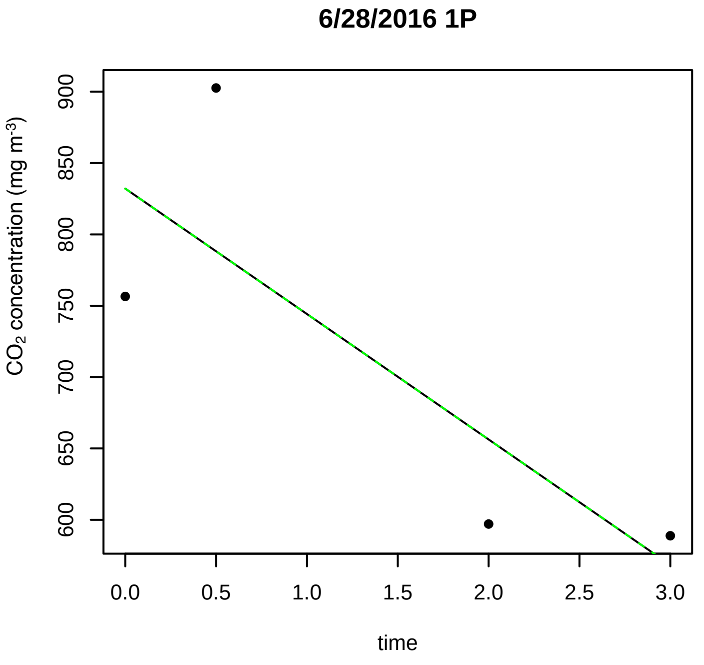
6/28/2016 1P
CO2 concentration (mg m-3)
600
650
700
750
800
850
900
0.0
0.5
1.0
1.5
2.0
2.5
3.0
time

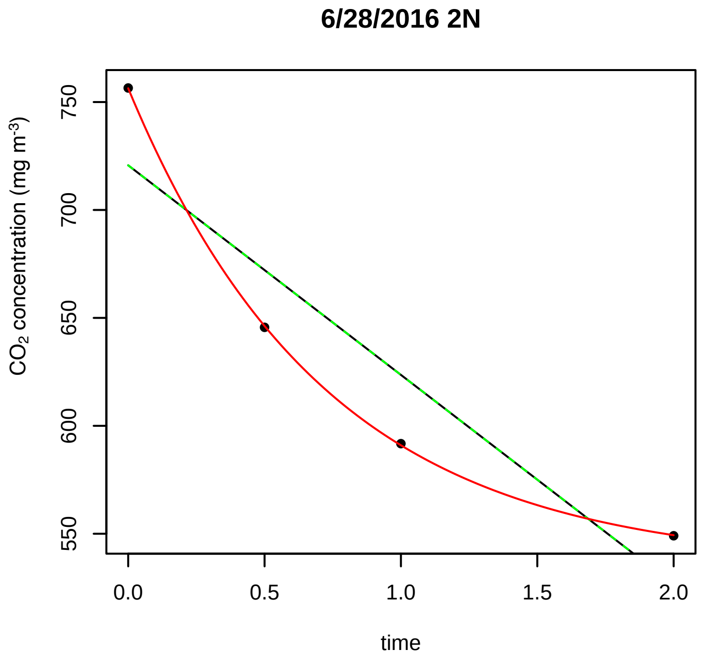
6/28/2016 2N
CO2 concentration (mg m-3)
550
600
650
700
750
0.0
0.5
1.0
1.5
2.0
time

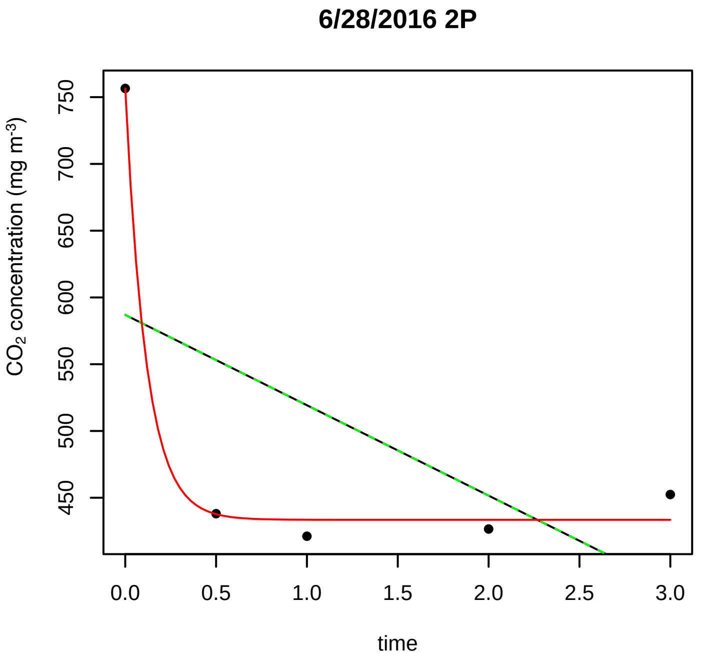
6/28/2016 2P
CO2 concentration (mg m-3)
450
500
550
600
650
700
750
0.0
0.5
1.0
1.5
2.0
2.5
3.0
time

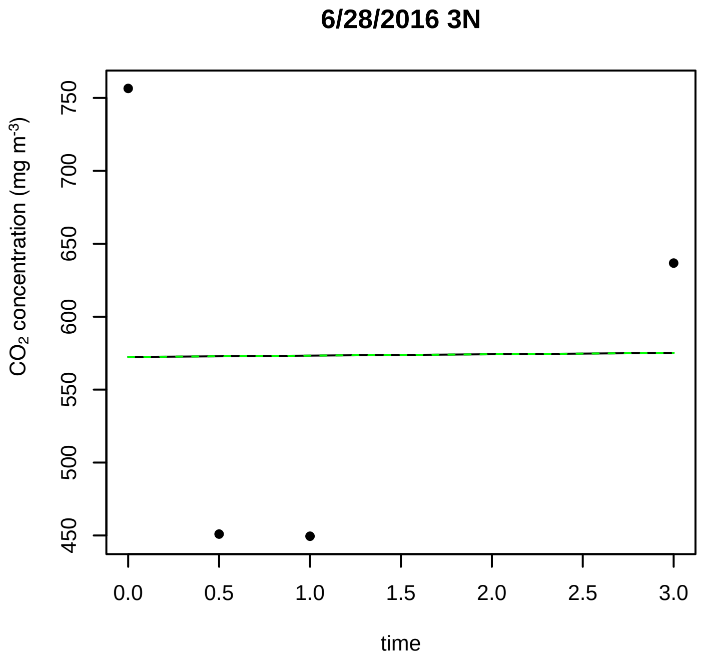
6/28/2016 3N
CO2 concentration (mg m-3)
450
500
550
600
650
700
750
0.0
0.5
1.0
1.5
2.0
2.5
3.0
time

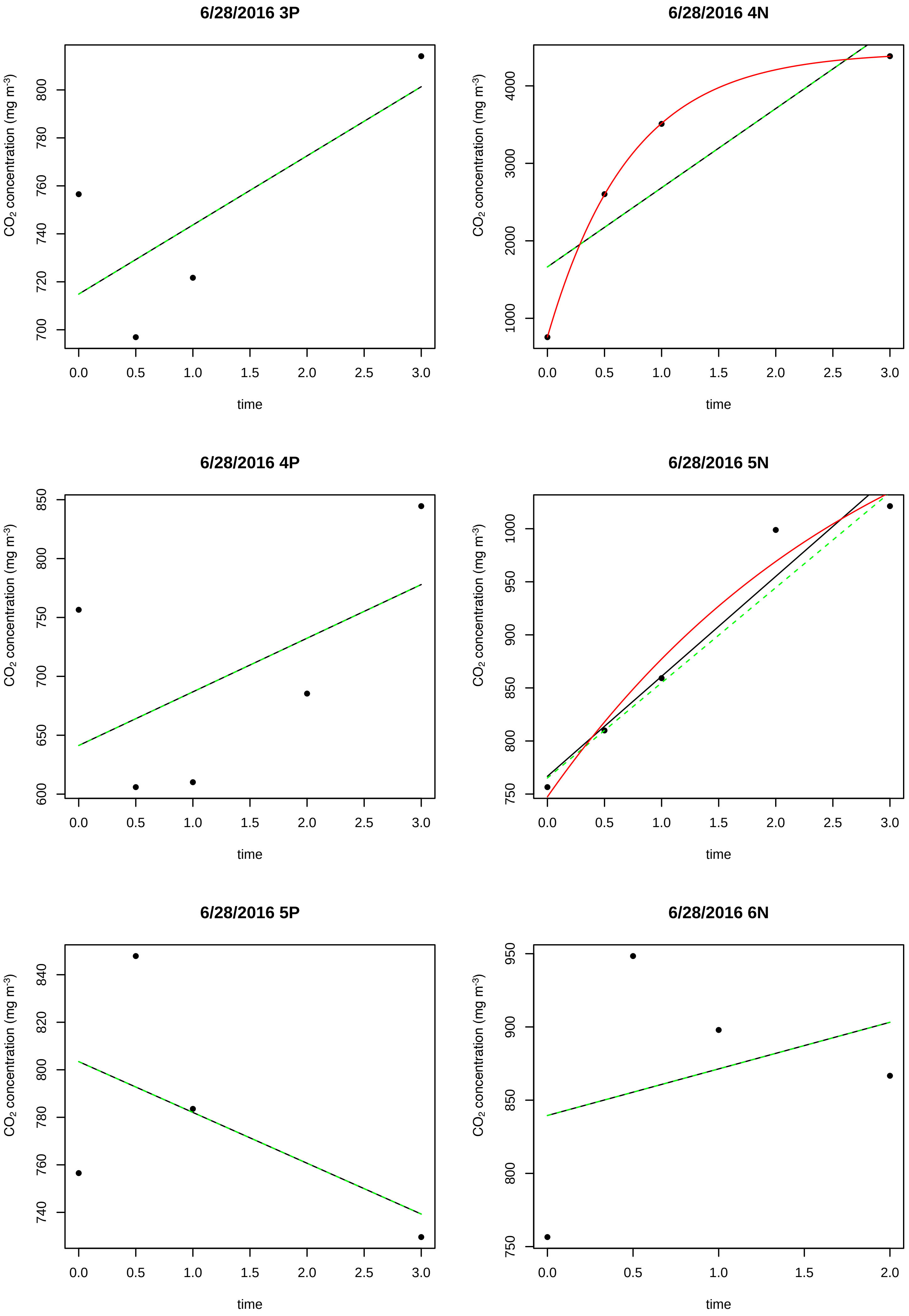
6/28/2016 3P
6/28/2016 4N
6/28/2016 4P
6/28/2016 5N
6/28/2016 5P
6/28/2016 6N
CO2 concentration (mg m-3)
time
0.0
0.5
1.0
1.5
2.0
2.5
3.0
700
720
740
760
780
800
1000
2000
3000
4000
600
650
750
850
900
950
740
760
780
800
820
840

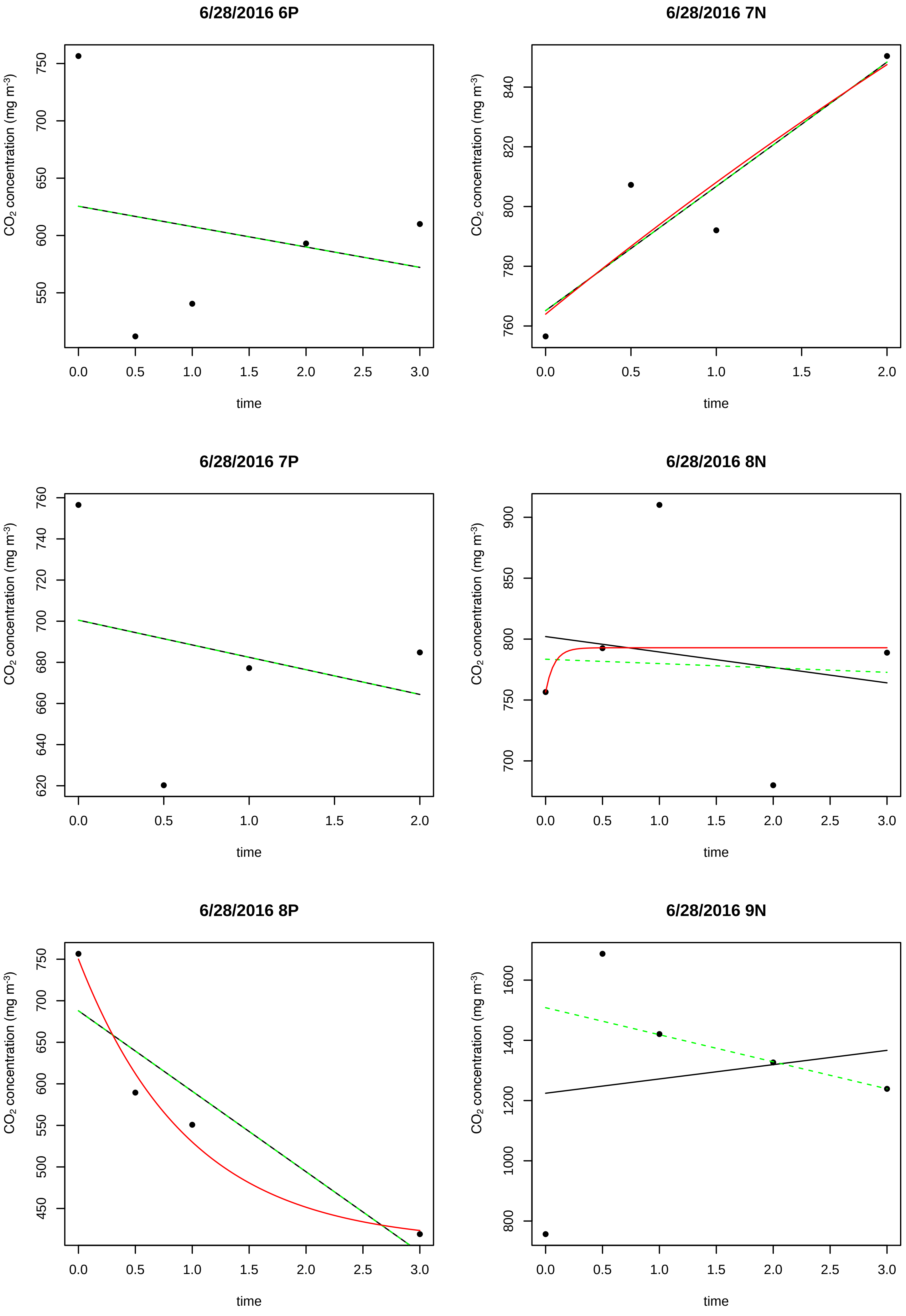
6/28/2016 6P
6/28/2016 7N
6/28/2016 7P
6/28/2016 8N
6/28/2016 8P
6/28/2016 9N
CO2 concentration (mg m-3)
time

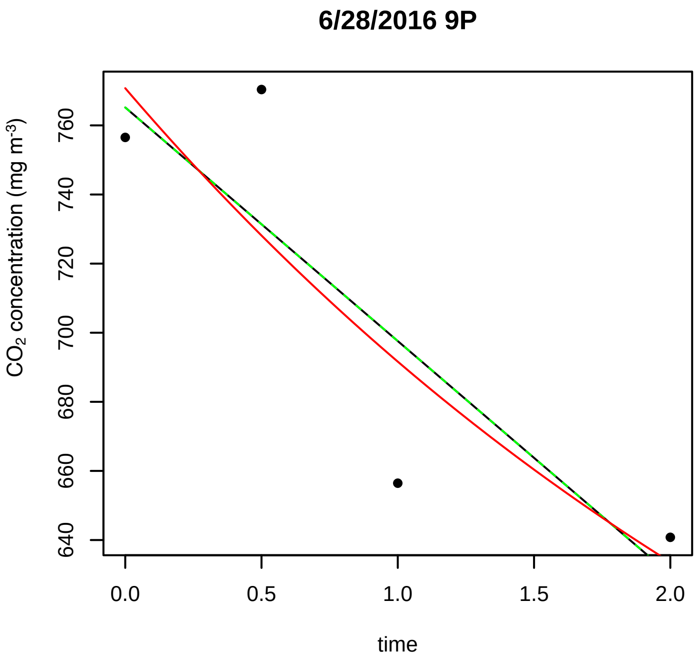
6/28/2016 9P
CO2 concentration (mg m-3)
640 660 680 700 720 740 760
0.0 0.5 1.0 1.5 2.0
time

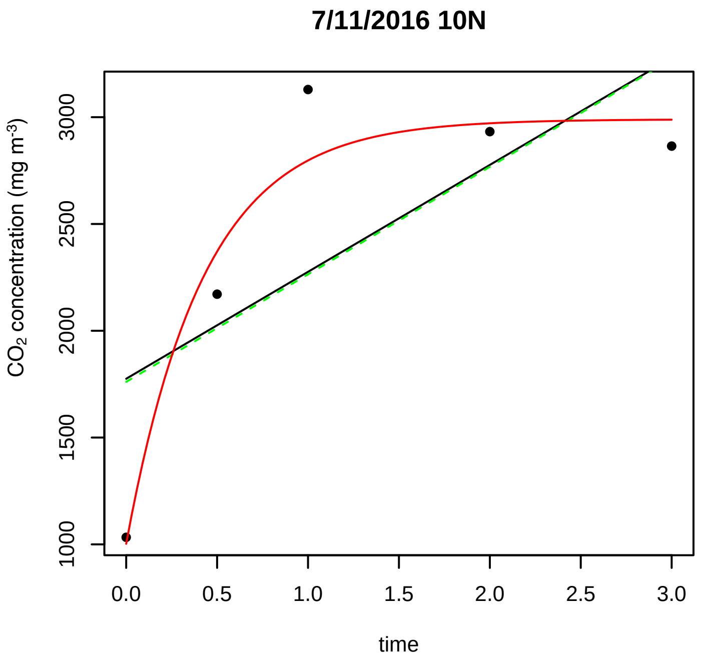
7/11/2016 10N
CO2 concentration (mg m-3)
1000 1500 2000 2500 3000
0.0 0.5 1.0 1.5 2.0 2.5 3.0
time

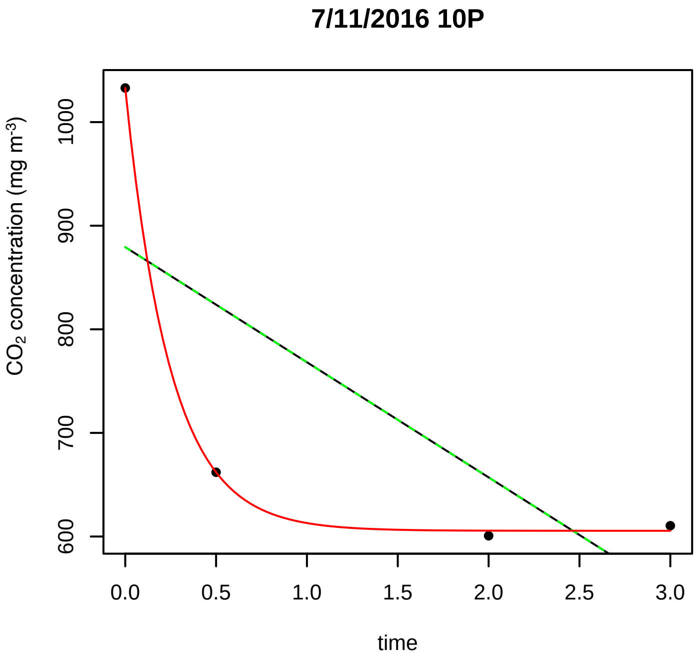
7/11/2016 10P
CO2 concentration (mg m-3)
600 700 800 900 1000
0.0 0.5 1.0 1.5 2.0 2.5 3.0
time

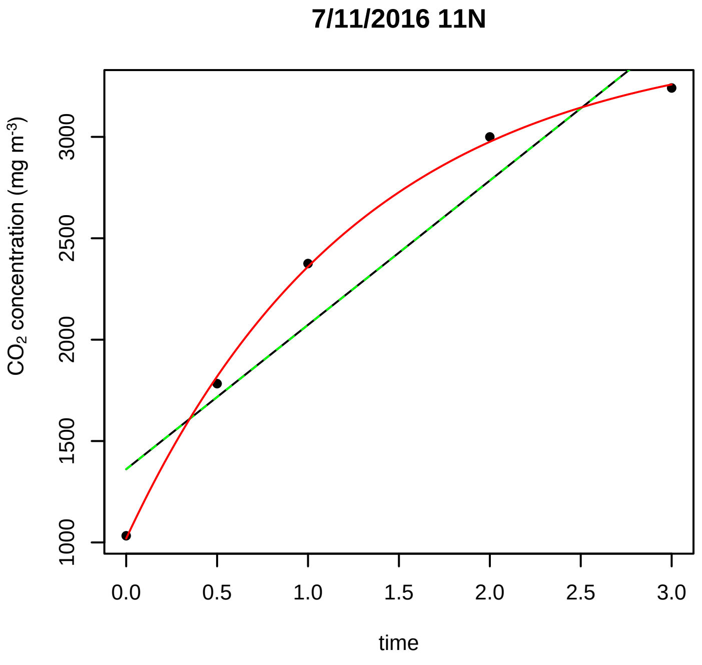
7/11/2016 11N
CO2 concentration (mg m-3)
1000 1500 2000 2500 3000
0.0 0.5 1.0 1.5 2.0 2.5 3.0
time

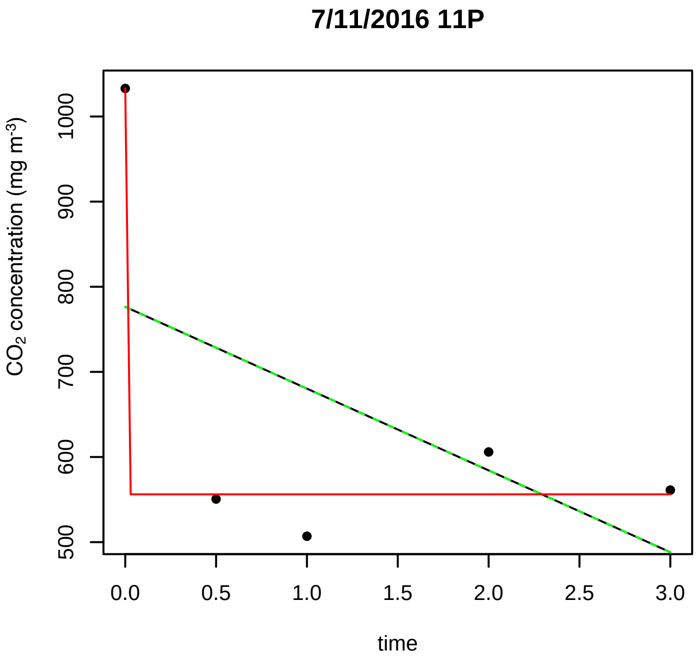
7/11/2016 11P
CO2 concentration (mg m-3)
500 600 700 800 900 1000
0.0 0.5 1.0 1.5 2.0 2.5 3.0
time

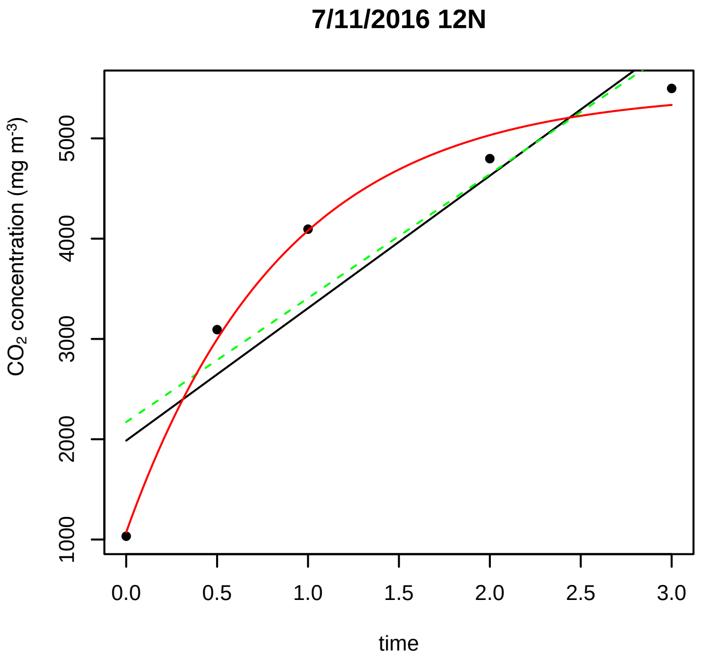
7/11/2016 12N
CO2 concentration (mg m-3)
1000 2000 3000 4000 5000
0.0 0.5 1.0 1.5 2.0 2.5 3.0
time

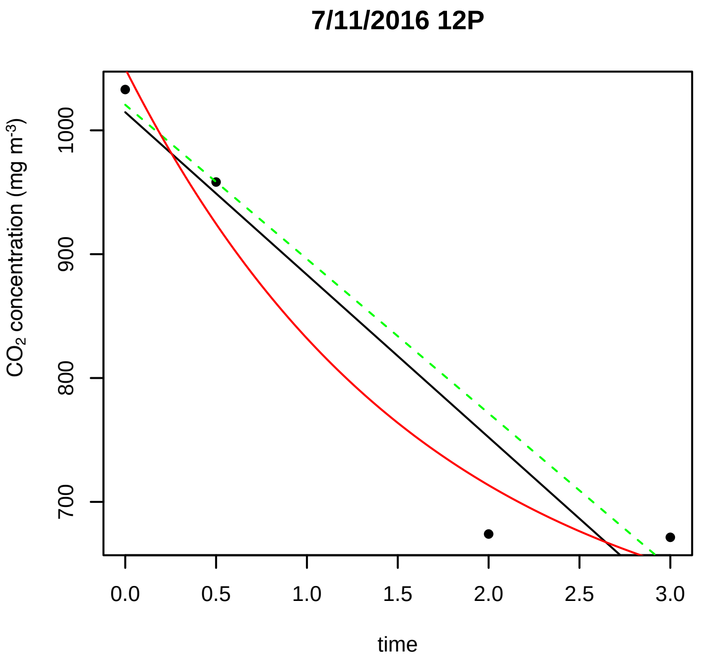
7/11/2016 12P
CO2 concentration (mg m-3)
700
800
900
1000
0.0
0.5
1.0
1.5
2.0
2.5
3.0
time

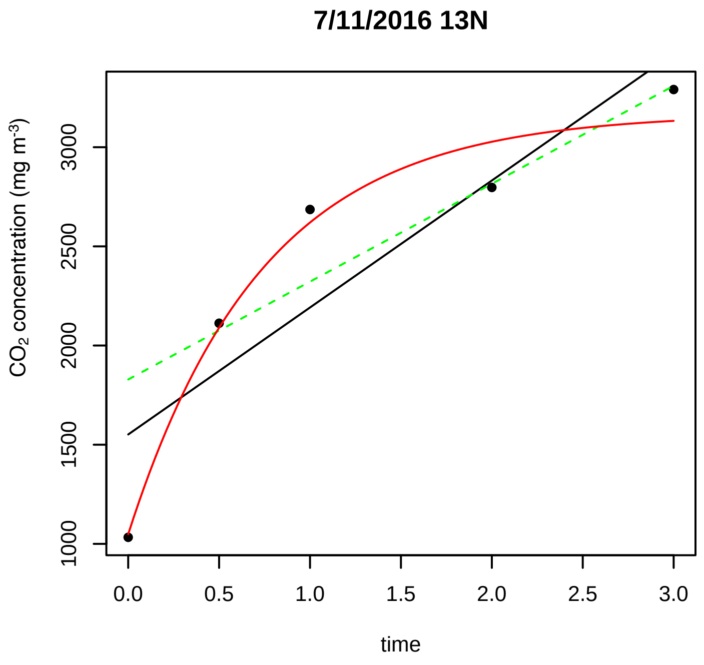
7/11/2016 13N
CO2 concentration (mg m-3)
1000
1500
2000
2500
3000
0.0
0.5
1.0
1.5
2.0
2.5
3.0
time

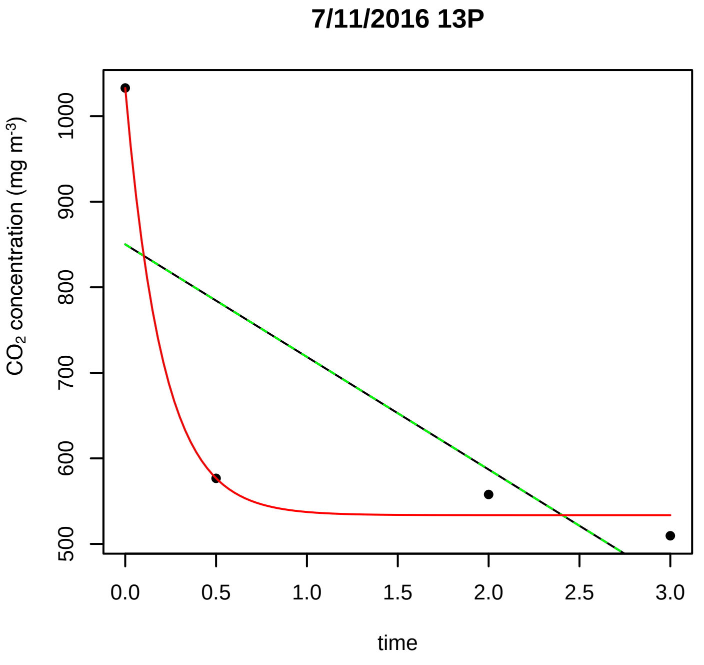
7/11/2016 13P
CO2 concentration (mg m-3)
500
600
700
800
900
1000
0.0
0.5
1.0
1.5
2.0
2.5
3.0
time

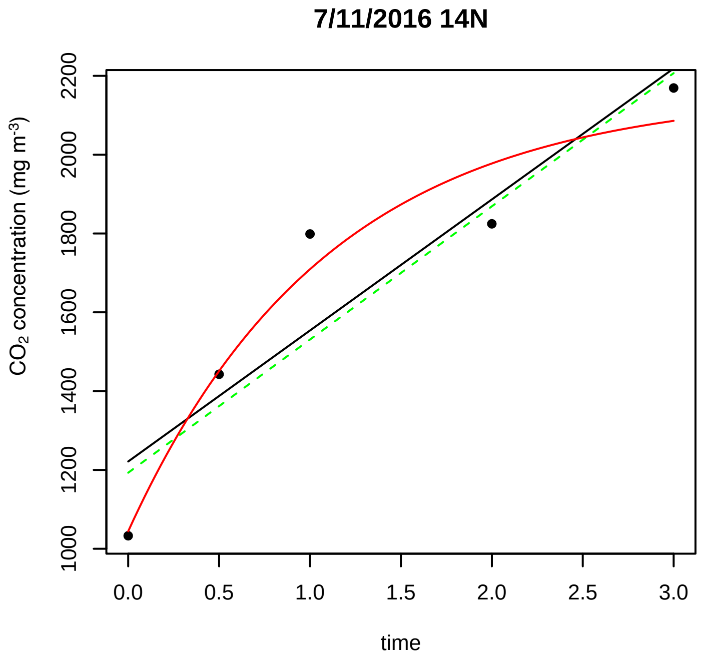
7/11/2016 14N
CO2 concentration (mg m-3)
1000
1200
1400
1600
1800
2000
2200
0.0
0.5
1.0
1.5
2.0
2.5
3.0
time

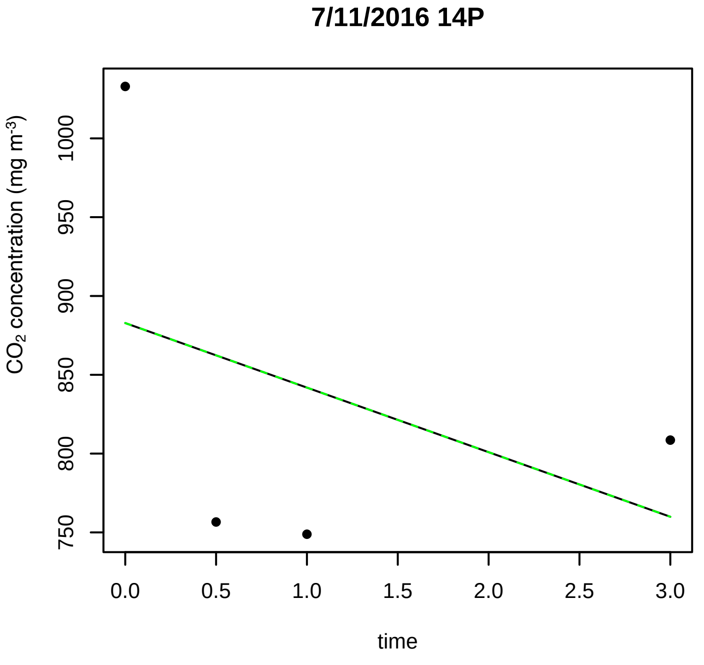
7/11/2016 14P
CO2 concentration (mg m-3)
750
800
850
900
950
1000
0.0
0.5
1.0
1.5
2.0
2.5
3.0
time

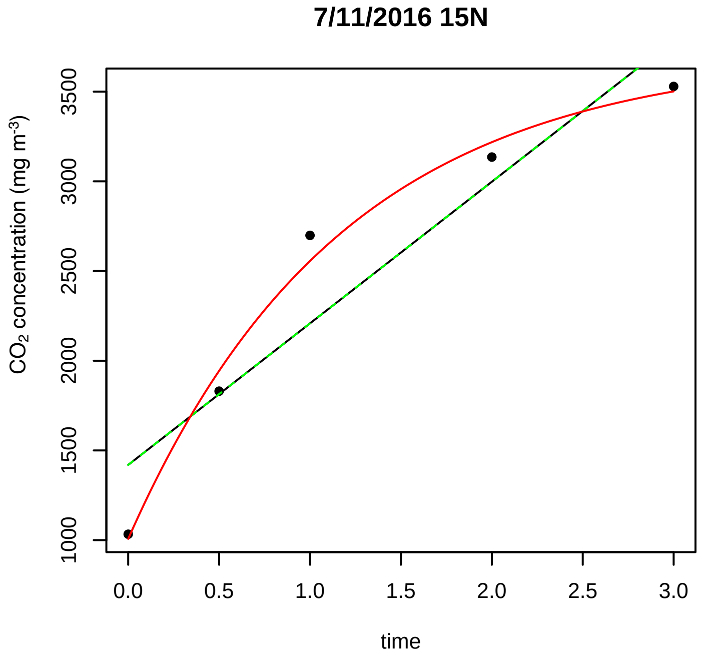
7/11/2016 15N
CO2 concentration (mg m-3)
1000
1500
2000
2500
3000
3500
0.0
0.5
1.0
1.5
2.0
2.5
3.0
time

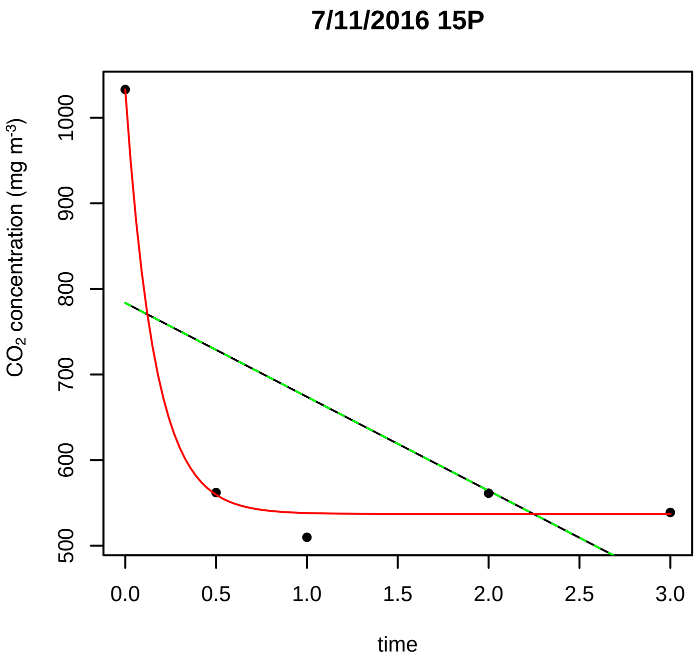
7/11/2016 15P
CO2 concentration (mg m-3)
500
600
700
800
900
1000
0.0
0.5
1.0
1.5
2.0
2.5
3.0
time

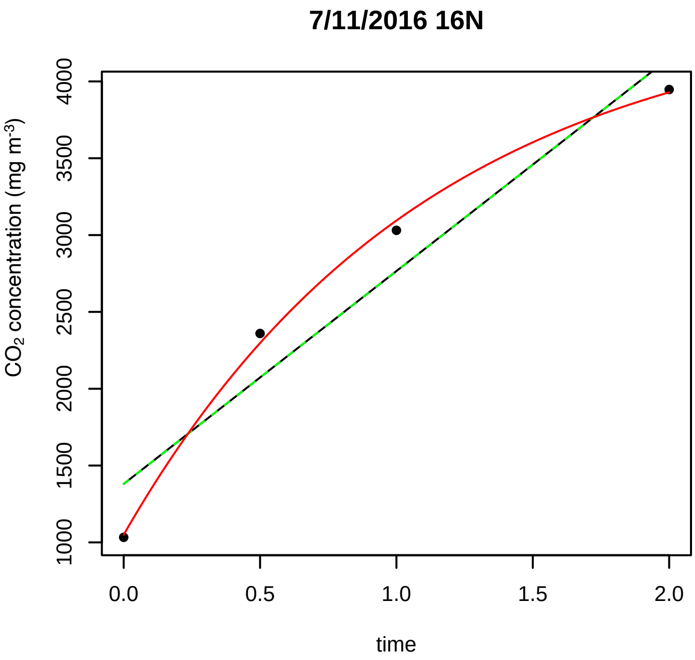
7/11/2016 16N
CO2 concentration (mg m-3)
1000
1500
2000
2500
3000
3500
4000
0.0
0.5
1.0
1.5
2.0
time

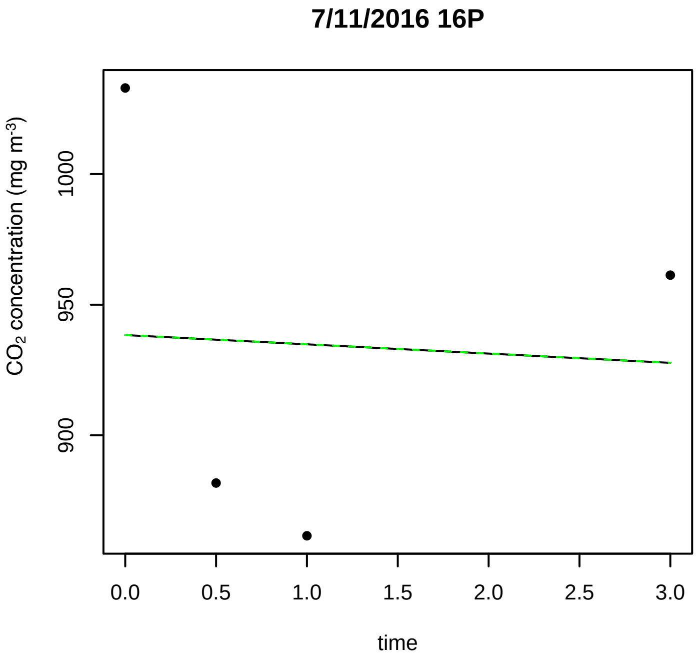
7/11/2016 16P
CO2 concentration (mg m-3)
900
950
1000
0.0
0.5
1.0
1.5
2.0
2.5
3.0
time

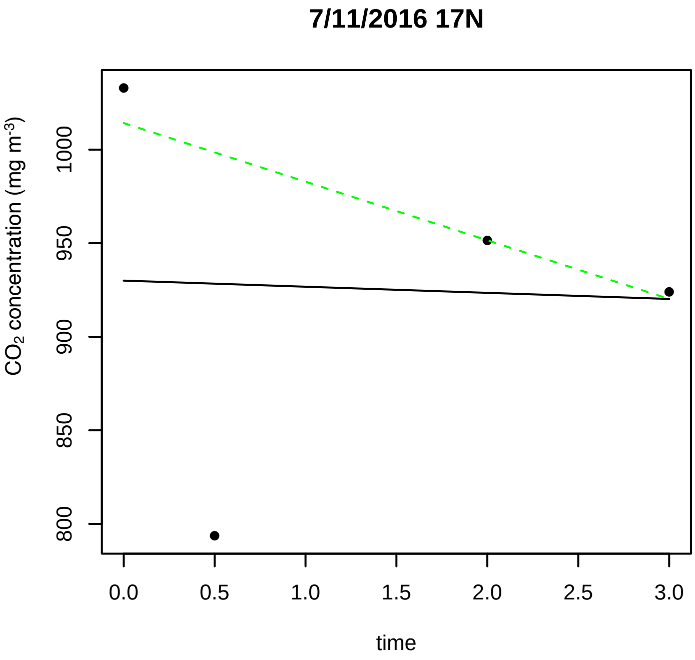
7/11/2016 17N
CO2 concentration (mg m-3)
800
850
900
950
1000
0.0
0.5
1.0
1.5
2.0
2.5
3.0
time

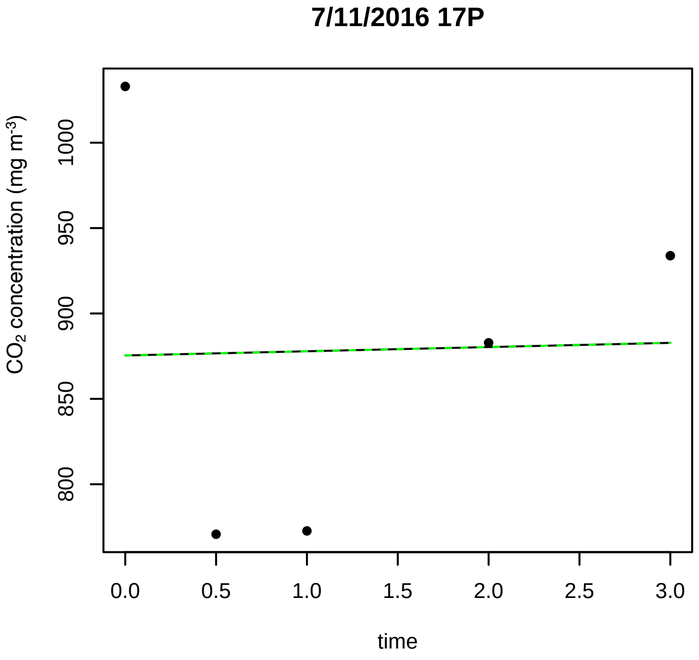
7/11/2016 17P
CO2 concentration (mg m-3)
800
850
900
950
1000
0.0
0.5
1.0
1.5
2.0
2.5
3.0
time

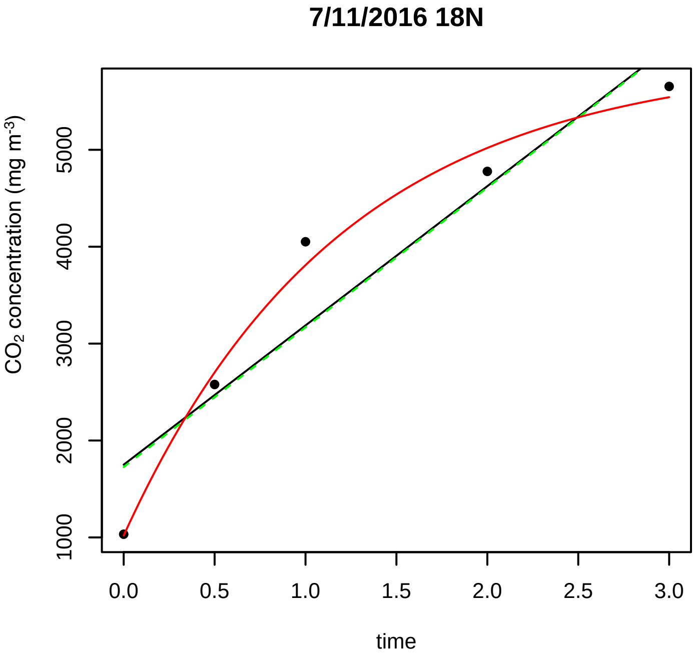
7/11/2016 18N
CO2 concentration (mg m-3)
1000
2000
3000
4000
5000
0.0
0.5
1.0
1.5
2.0
2.5
3.0
time

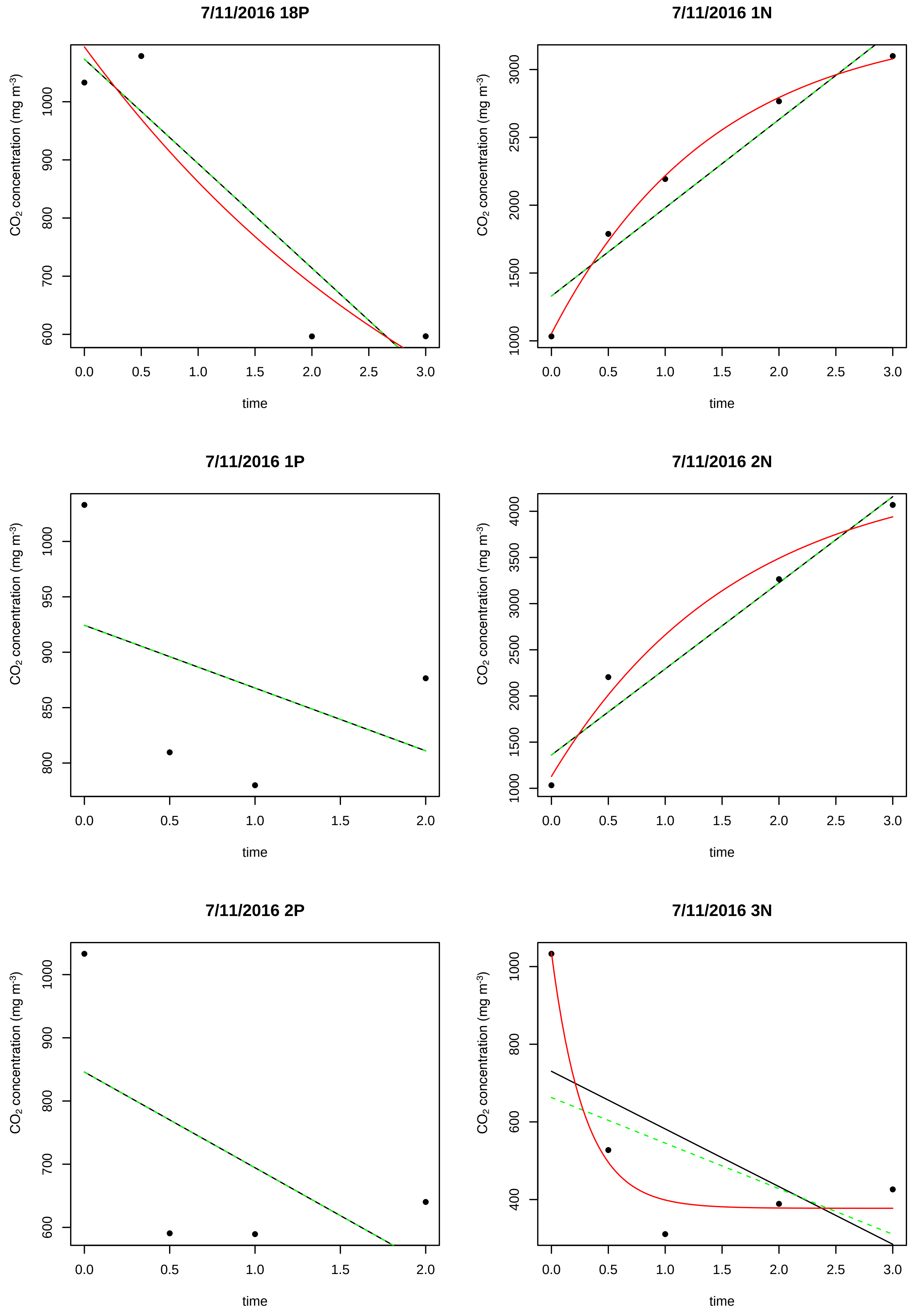
7/11/2016 18P
CO2 concentration (mg m-3)
600
700
800
900
1000
0.0
0.5
1.0
1.5
2.0
2.5
3.0
time
7/11/2016 1N
CO2 concentration (mg m-3)
1000
1500
2000
2500
3000
0.0
0.5
1.0
1.5
2.0
2.5
3.0
time
7/11/2016 1P
CO2 concentration (mg m-3)
800
850
900
950
1000
0.0
0.5
1.0
1.5
2.0
time
7/11/2016 2N
CO2 concentration (mg m-3)
1000
1500
2000
2500
3000
3500
4000
0.0
0.5
1.0
1.5
2.0
2.5
3.0
time
7/11/2016 2P
CO2 concentration (mg m-3)
600
700
800
900
1000
0.0
0.5
1.0
1.5
2.0
time
7/11/2016 3N
CO2 concentration (mg m-3)
400
600
800
1000
0.0
0.5
1.0
1.5
2.0
2.5
3.0
time

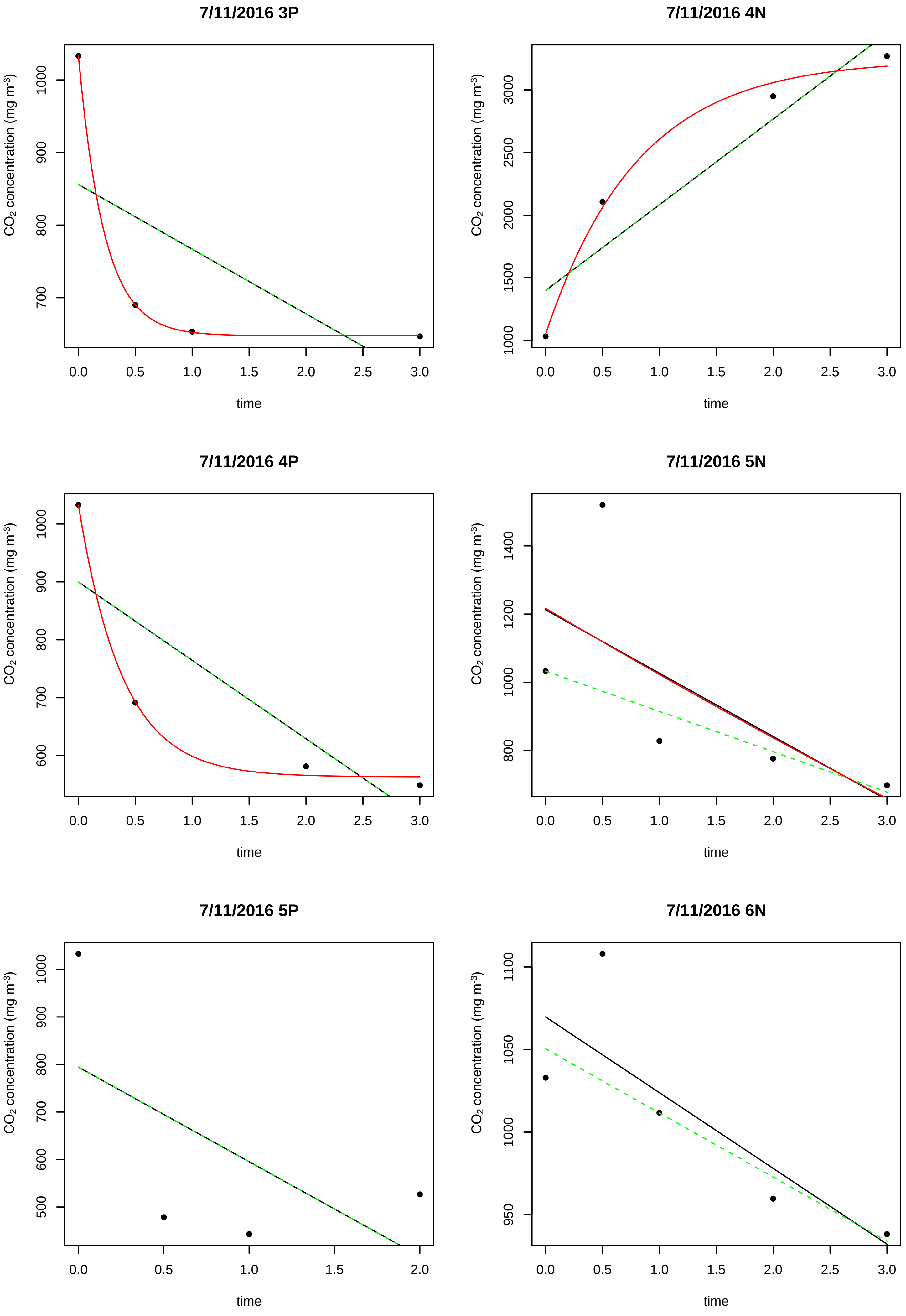

7/11/2016 3P
7/11/2016 4N
7/11/2016 4P
7/11/2016 5N
7/11/2016 5P
7/11/2016 6N
CO2 concentration (mg m-3)
time

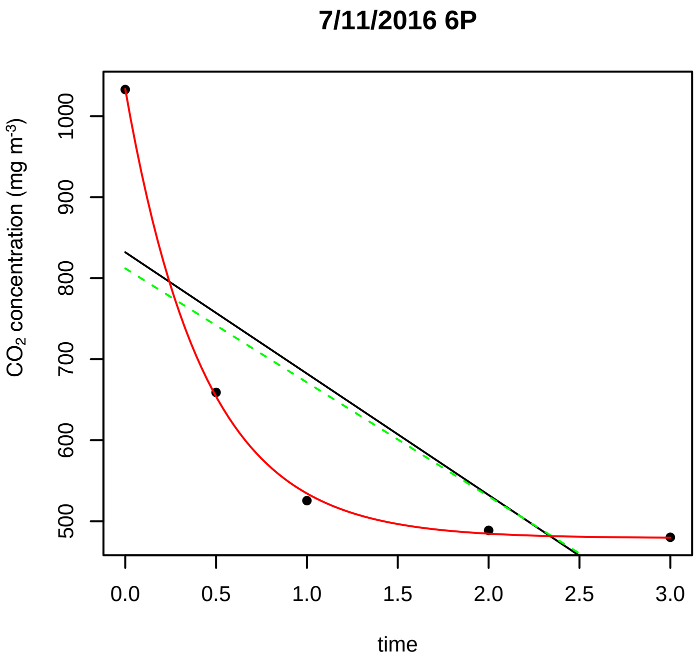
7/11/2016 6P
CO2 concentration (mg m-3)
500 600 700 800 900 1000
0.0 0.5 1.0 1.5 2.0 2.5 3.0
time

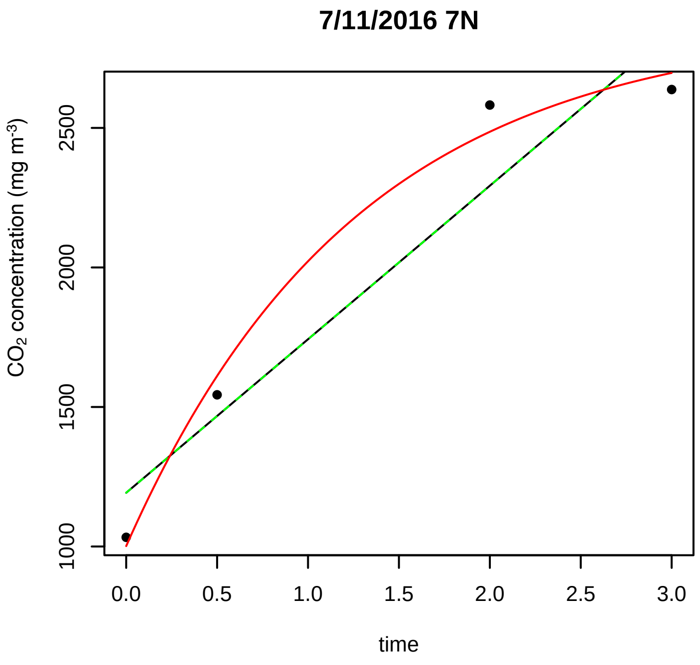
7/11/2016 7N
CO2 concentration (mg m-3)
1000 1500 2000 2500
0.0 0.5 1.0 1.5 2.0 2.5 3.0
time

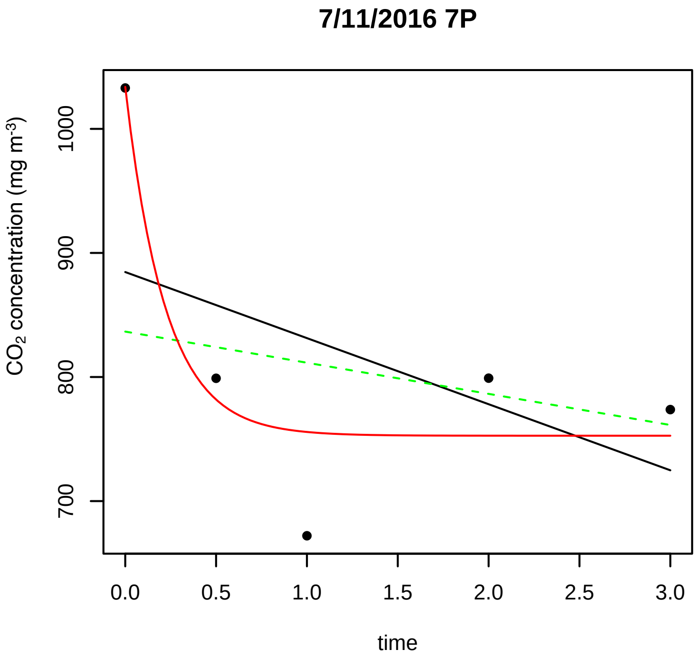
7/11/2016 7P
CO2 concentration (mg m-3)
700 800 900 1000
0.0 0.5 1.0 1.5 2.0 2.5 3.0
time

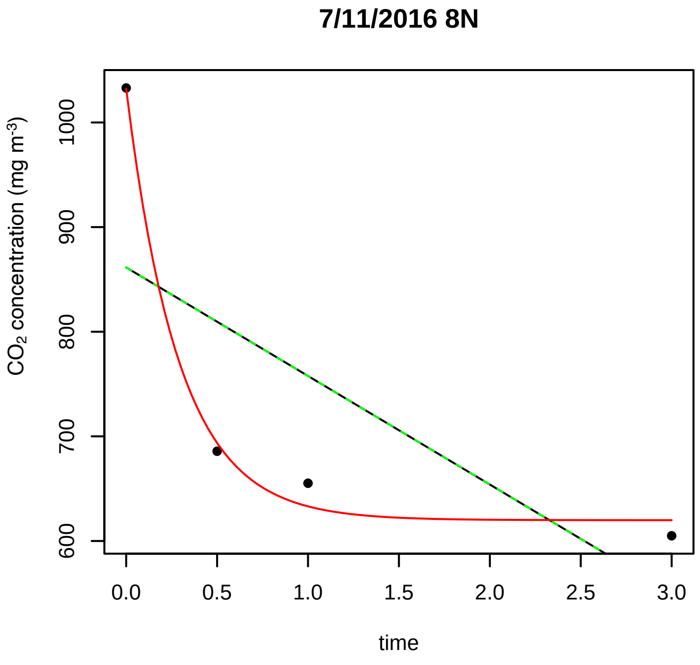
7/11/2016 8N
CO2 concentration (mg m-3)
600 700 800 900 1000
0.0 0.5 1.0 1.5 2.0 2.5 3.0
time

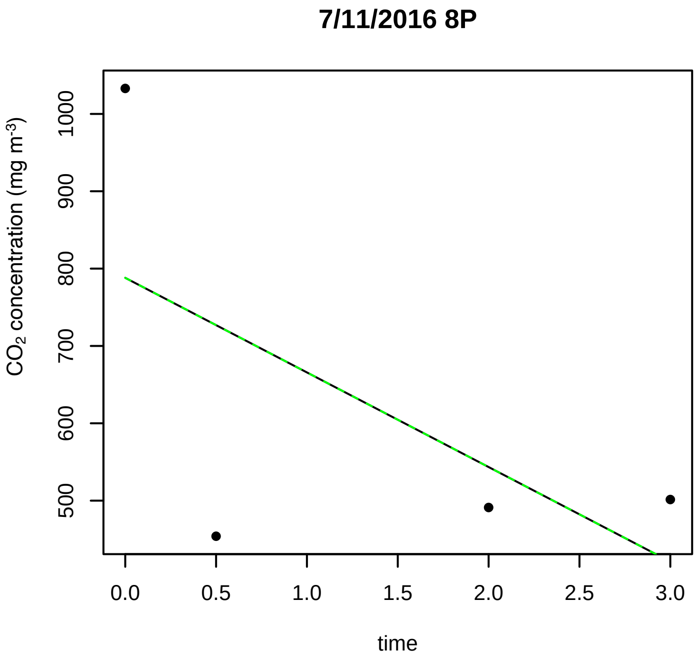
7/11/2016 8P
CO2 concentration (mg m-3)
500 600 700 800 900 1000
0.0 0.5 1.0 1.5 2.0 2.5 3.0
time

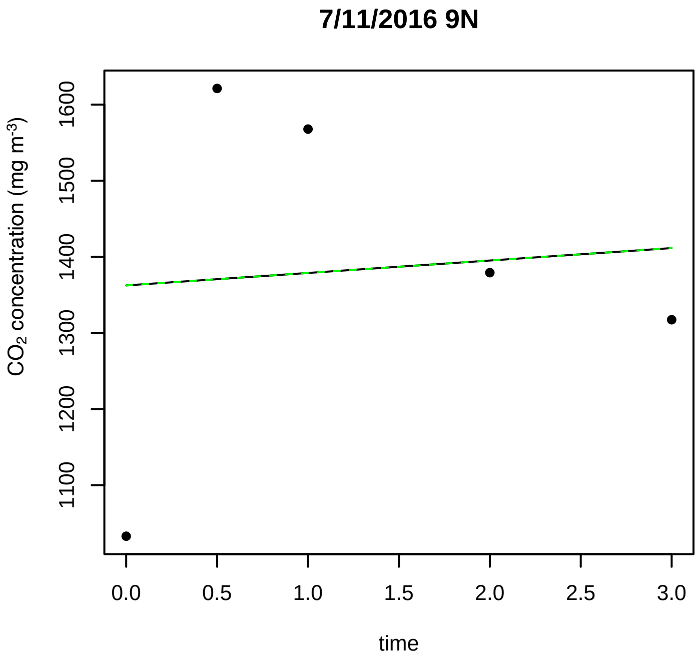
7/11/2016 9N
CO2 concentration (mg m-3)
1100 1200 1300 1400 1500 1600
0.0 0.5 1.0 1.5 2.0 2.5 3.0
time

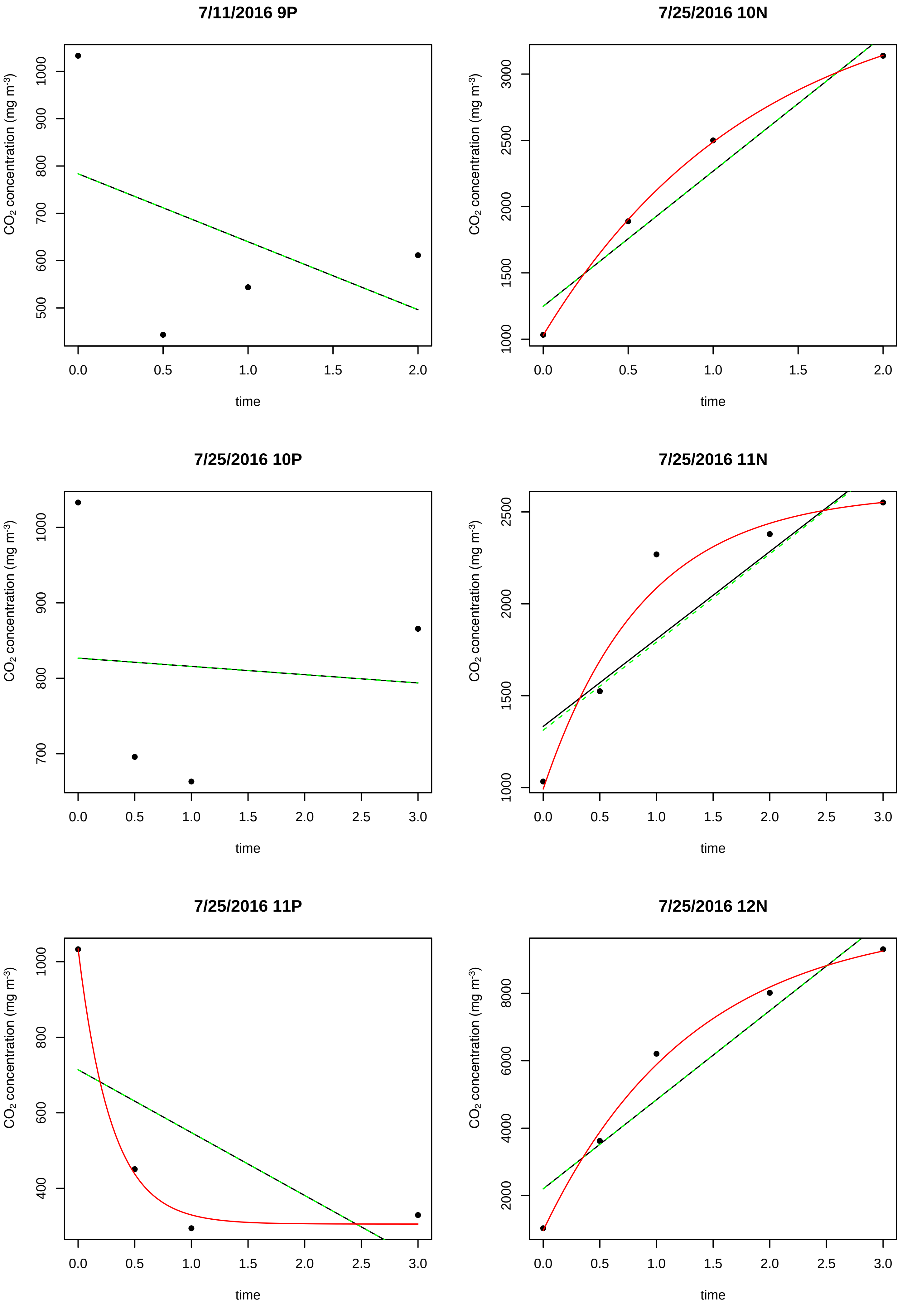
7/11/2016 9P
7/25/2016 10N
7/25/2016 10P
7/25/2016 11N
7/25/2016 11P
7/25/2016 12N
CO2 concentration (mg m-3)
time
0.0
0.5
1.0
1.5
2.0
2.5
3.0
500
600
700
800
900
1000
1500
2000
2500
3000
400
2000
4000
6000
8000

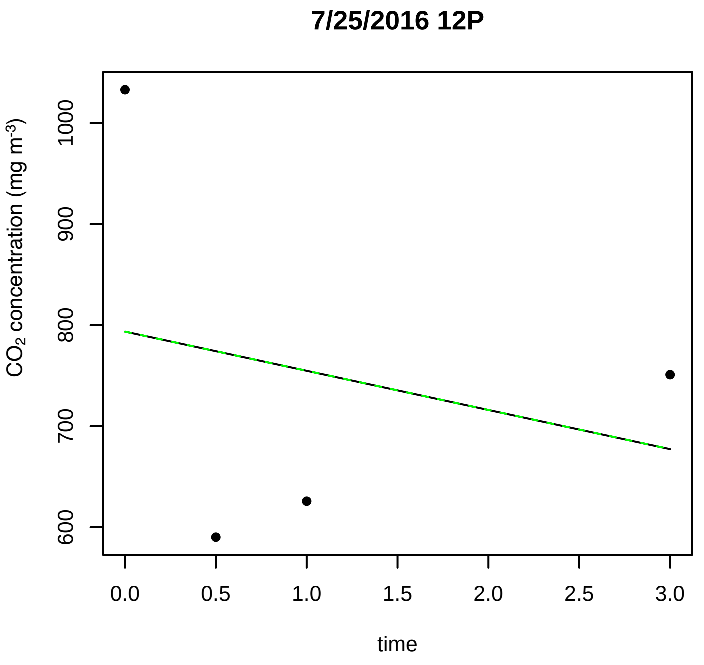
7/25/2016 12P
$CO_2$ concentration (mg m$^{-3}$)
600
700
800
900
1000
0.0
0.5
1.0
1.5
2.0
2.5
3.0
time

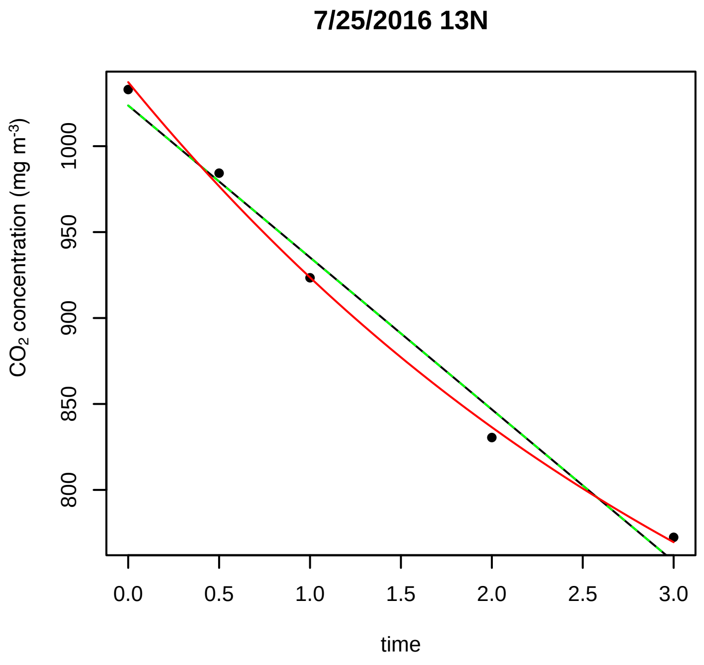
7/25/2016 13N
$CO_2$ concentration (mg m$^{-3}$)
800
850
900
950
1000
0.0
0.5
1.0
1.5
2.0
2.5
3.0
time

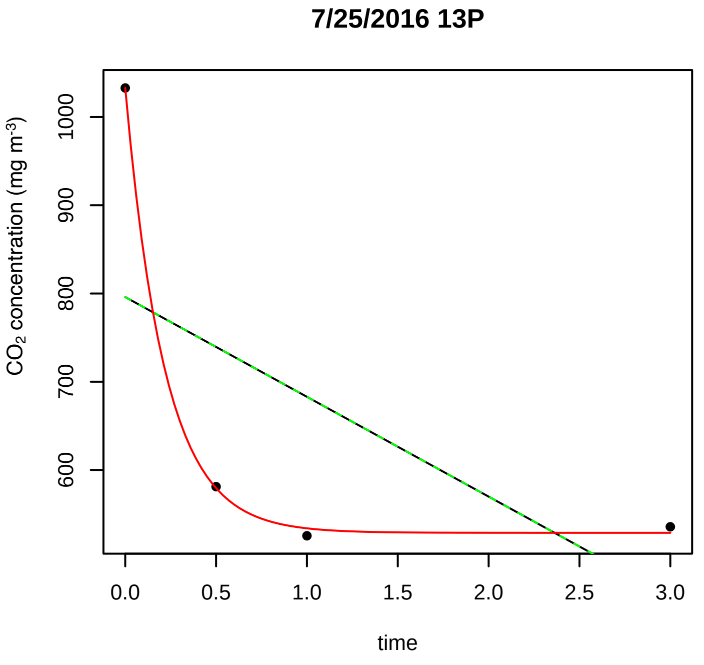
7/25/2016 13P
$CO_2$ concentration (mg m$^{-3}$)
600
700
800
900
1000
0.0
0.5
1.0
1.5
2.0
2.5
3.0
time

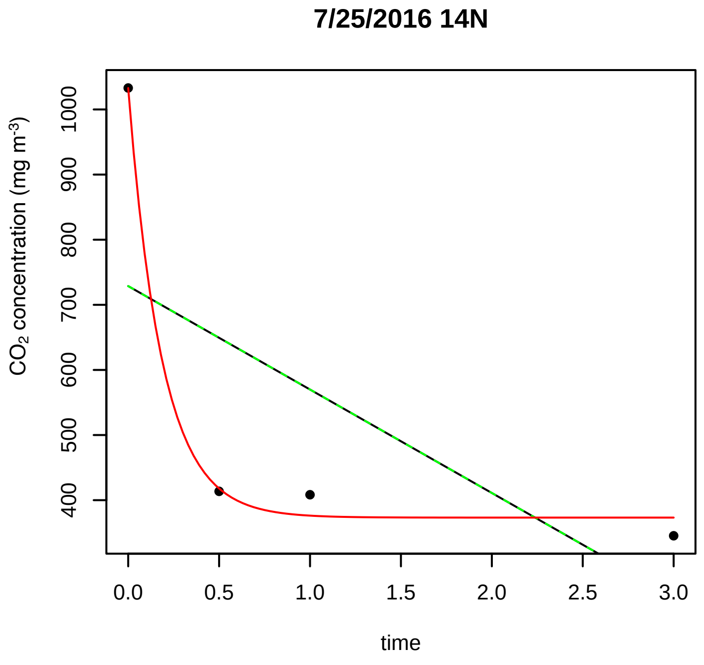
7/25/2016 14N
$CO_2$ concentration (mg m$^{-3}$)
400
500
600
700
800
900
1000
0.0
0.5
1.0
1.5
2.0
2.5
3.0
time

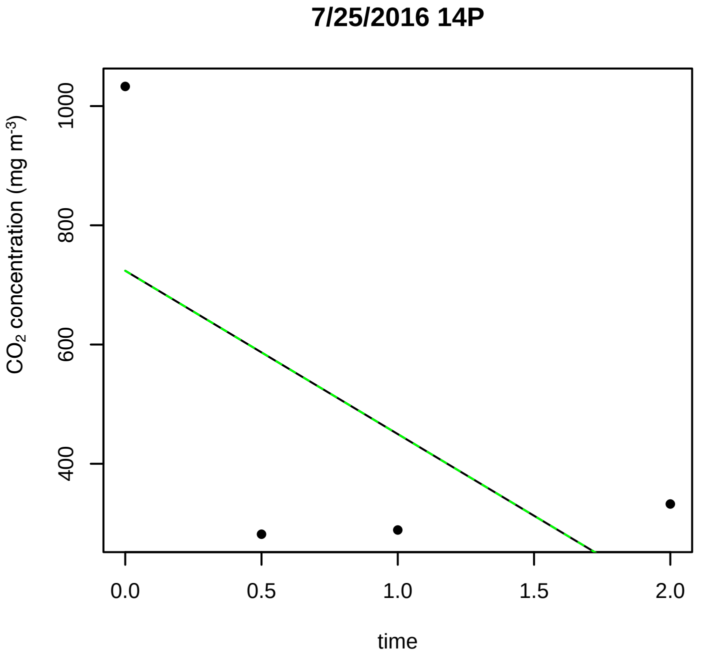
7/25/2016 14P
$CO_2$ concentration (mg m$^{-3}$)
400
600
800
1000
0.0
0.5
1.0
1.5
2.0
time

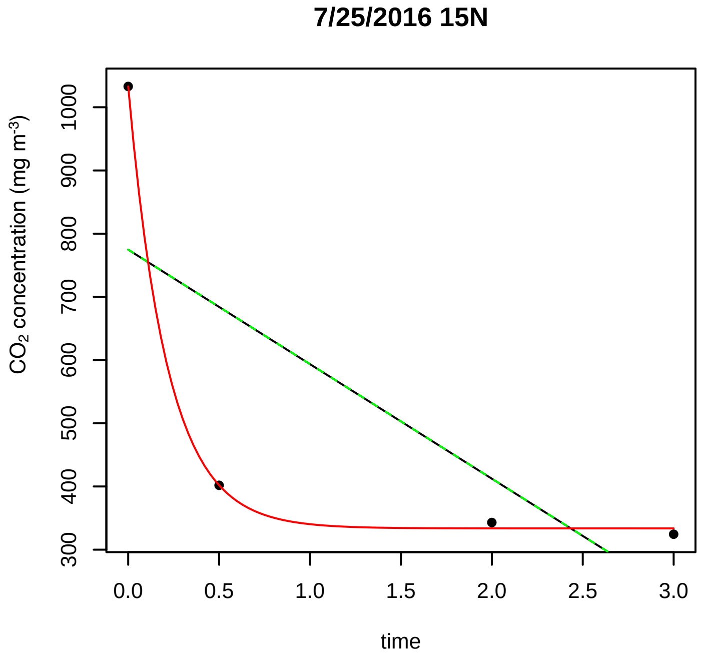
7/25/2016 15N
$CO_2$ concentration (mg m$^{-3}$)
300
400
500
600
700
800
900
1000
0.0
0.5
1.0
1.5
2.0
2.5
3.0
time

**7/25/2016 15P**

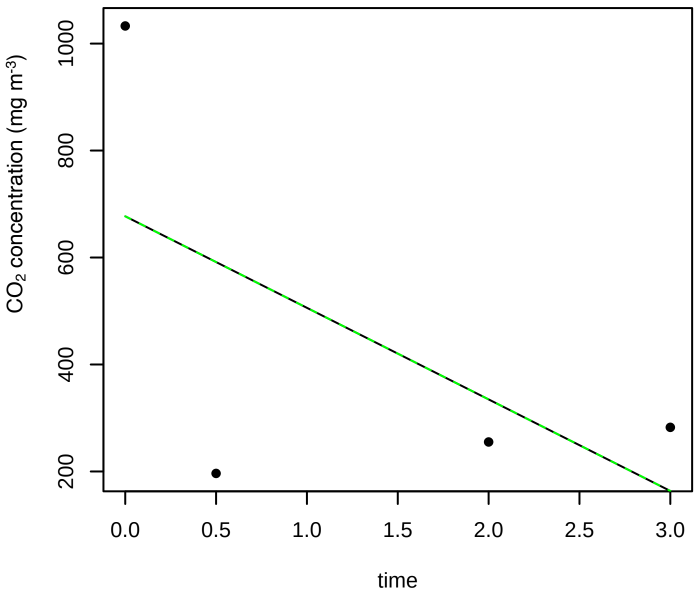


**7/25/2016 16N**

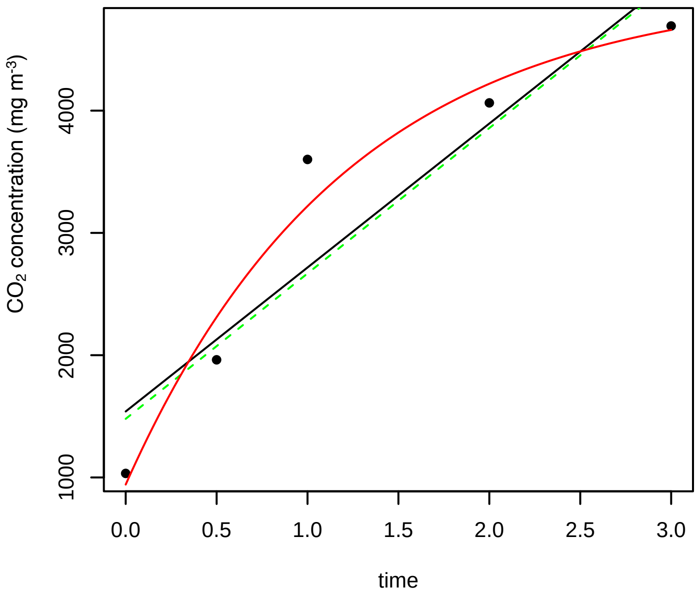


**7/25/2016 16P**

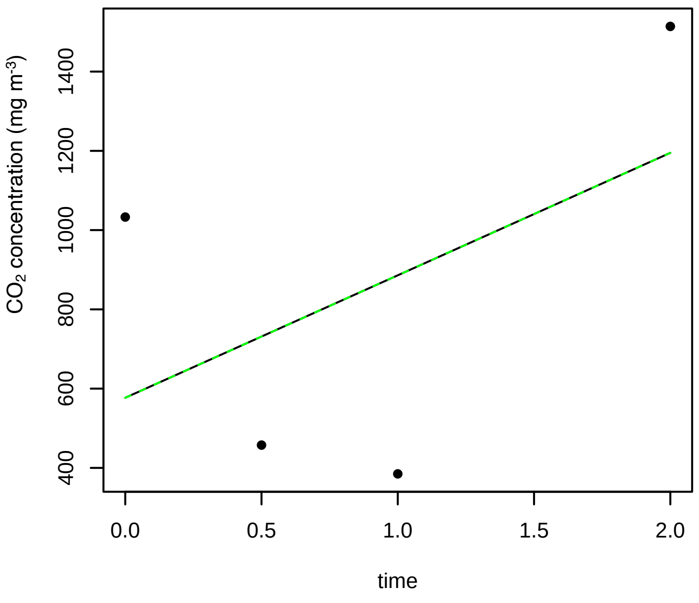


**7/25/2016 17N**

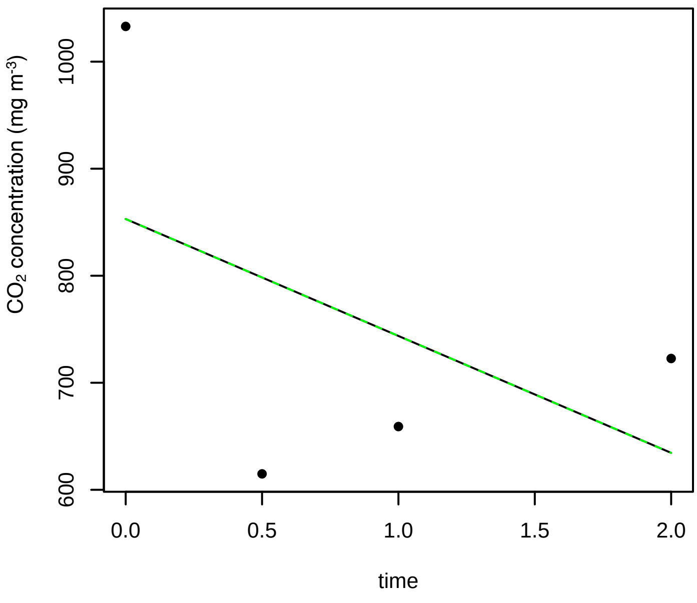


**7/25/2016 17P**

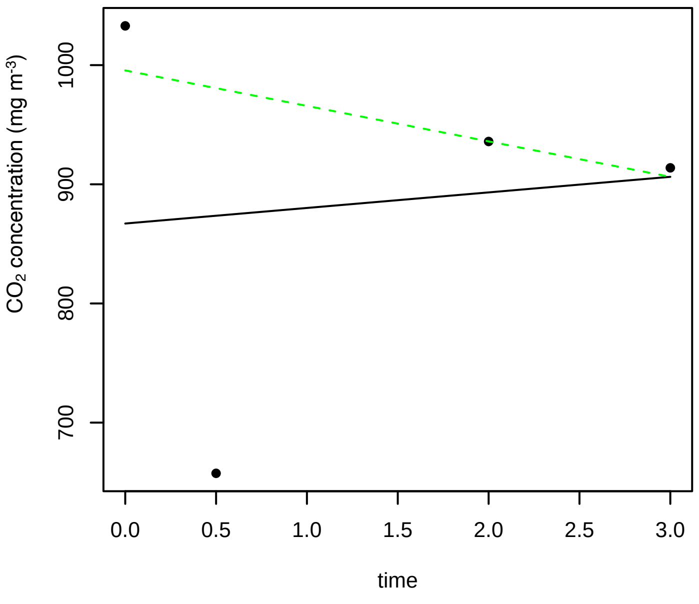


**7/25/2016 18N**

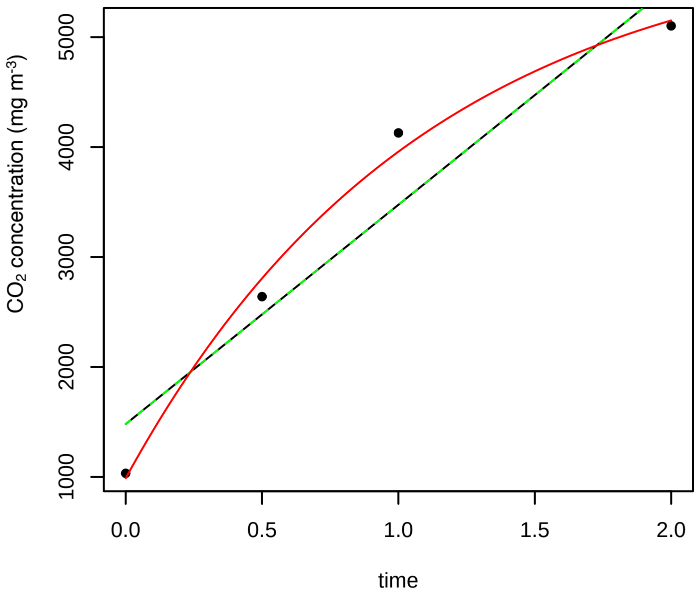

7/25/2016 18P

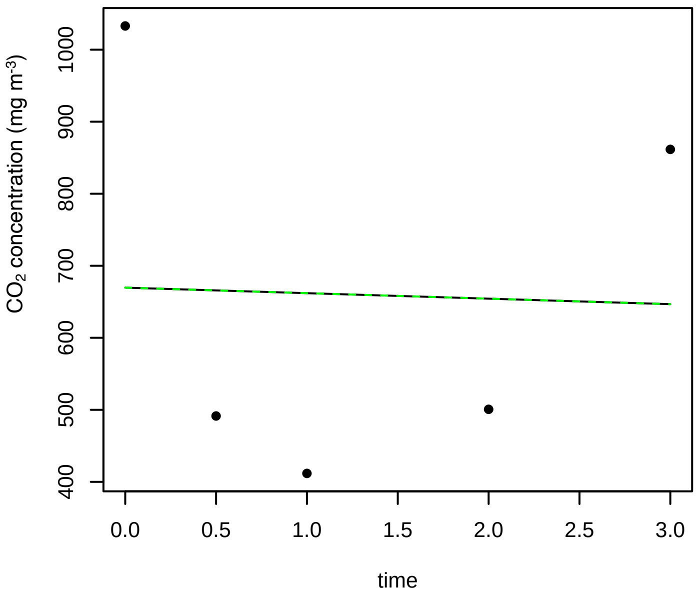


7/25/2016 1N

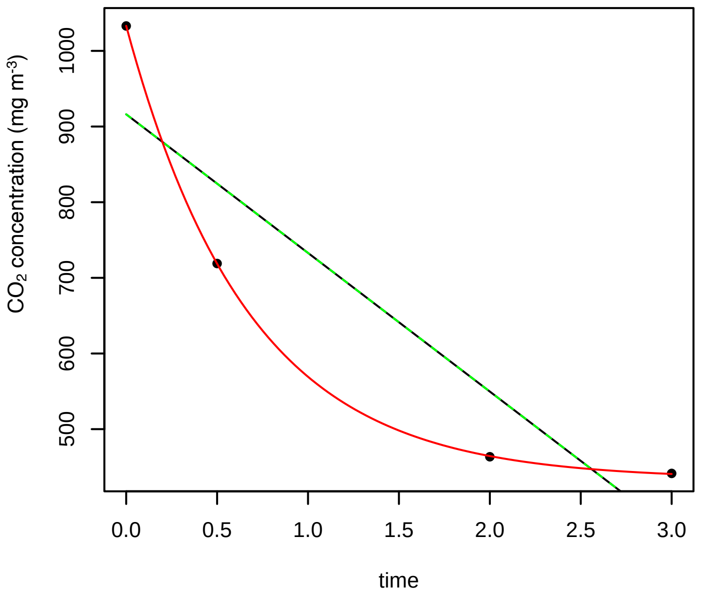


7/25/2016 1P

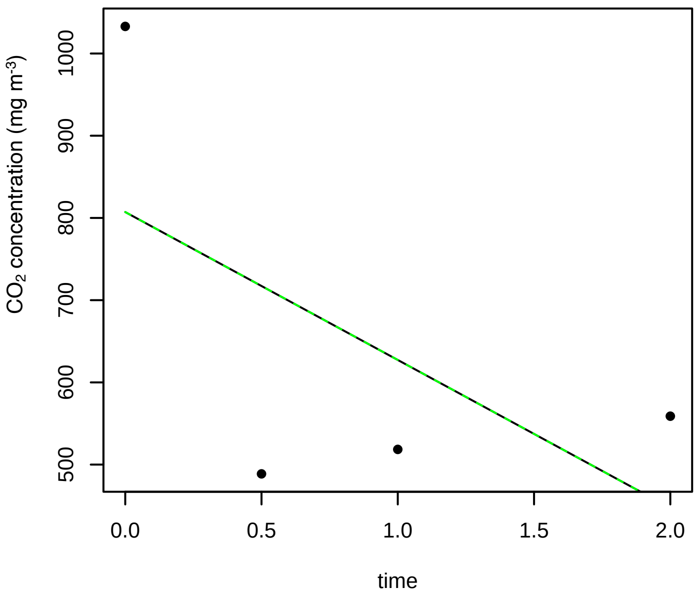


7/25/2016 2N

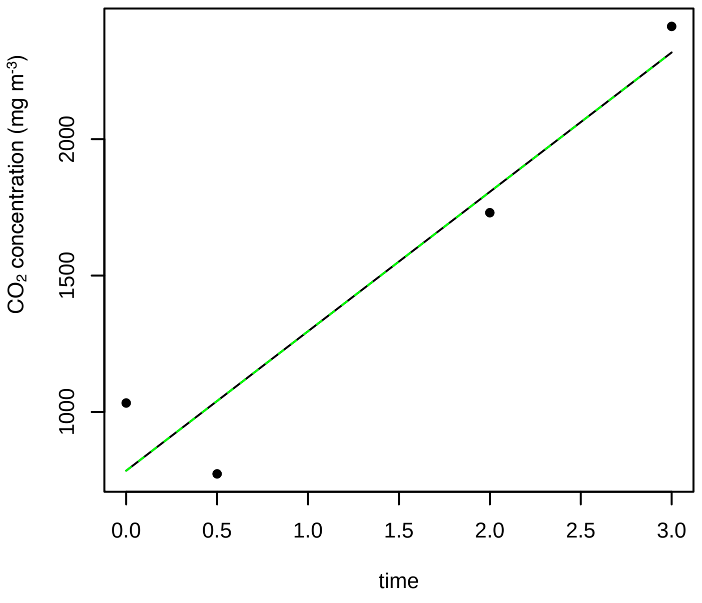


7/25/2016 2P

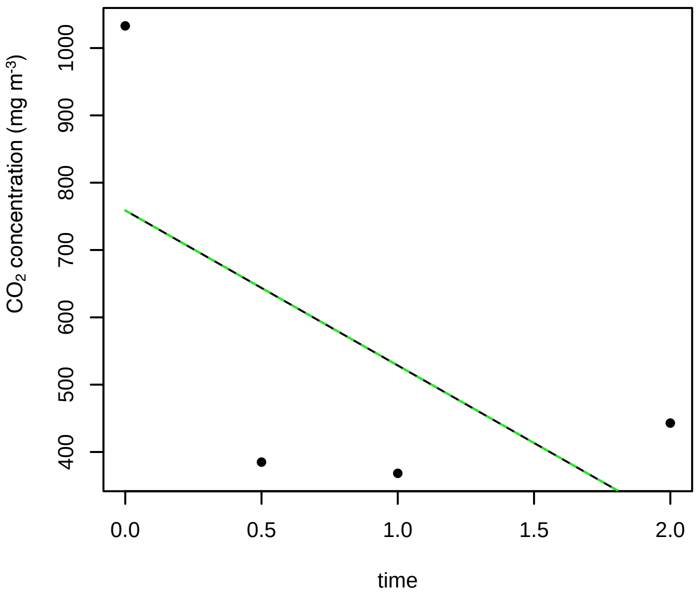


7/25/2016 3N

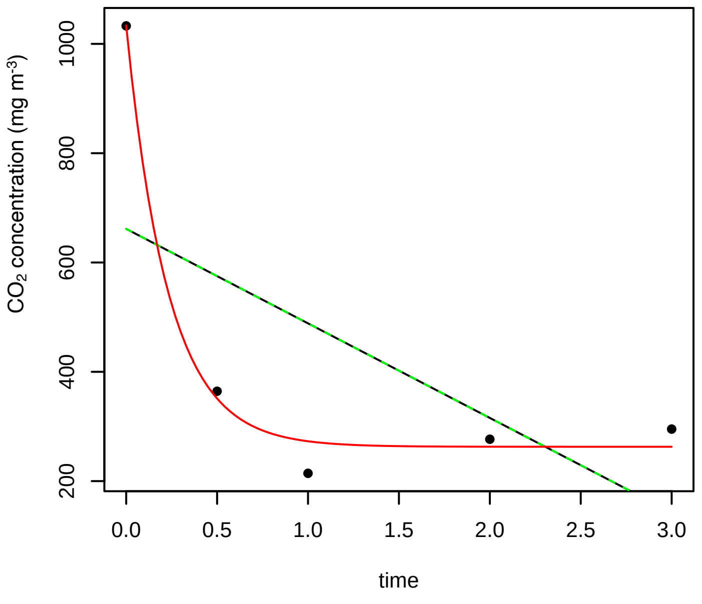

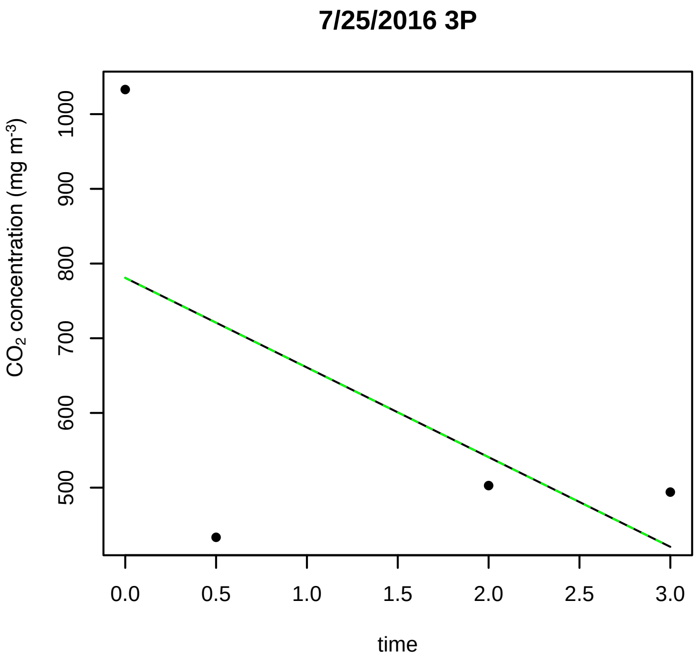
7/25/2016 3P
CO2 concentration (mg m-3)
500
600
700
800
900
1000
0.0
0.5
1.0
1.5
2.0
2.5
3.0
time

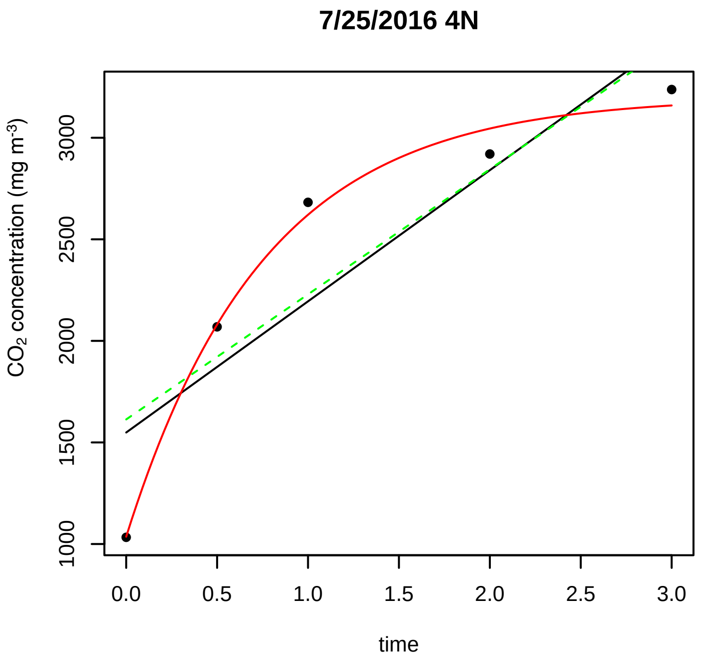
7/25/2016 4N
CO2 concentration (mg m-3)
1000
1500
2000
2500
3000
0.0
0.5
1.0
1.5
2.0
2.5
3.0
time

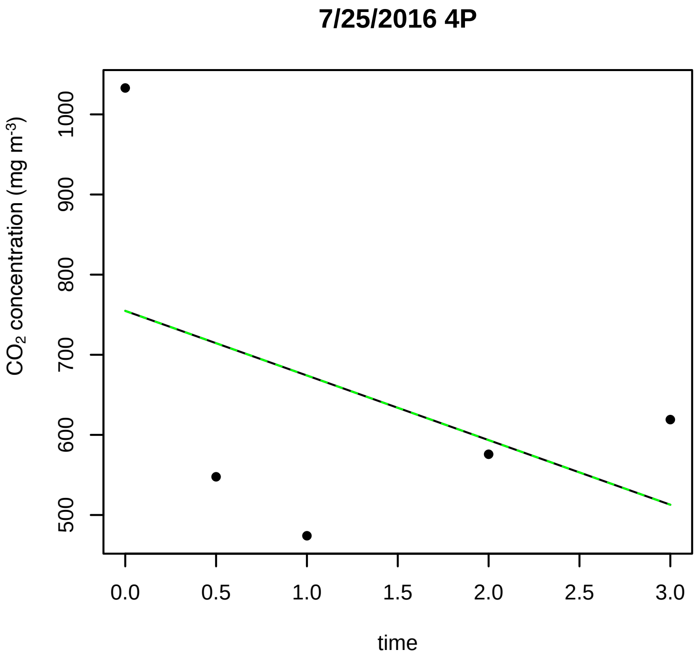
7/25/2016 4P
CO2 concentration (mg m-3)
500
600
700
800
900
1000
0.0
0.5
1.0
1.5
2.0
2.5
3.0
time

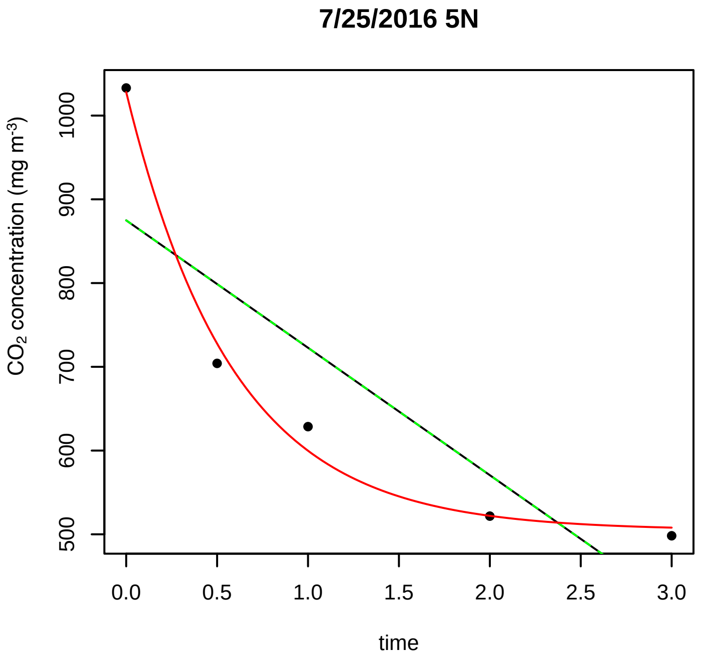
7/25/2016 5N
CO2 concentration (mg m-3)
500
600
700
800
900
1000
0.0
0.5
1.0
1.5
2.0
2.5
3.0
time

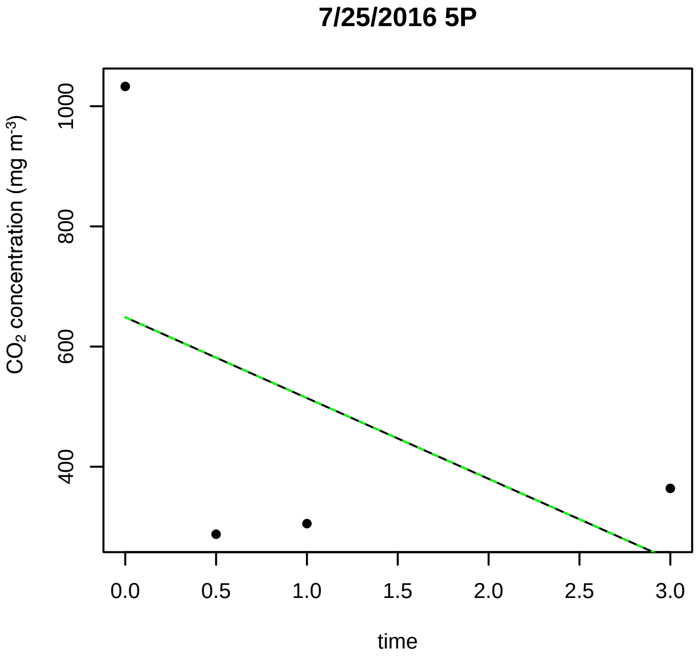
7/25/2016 5P
CO2 concentration (mg m-3)
400
600
800
1000
0.0
0.5
1.0
1.5
2.0
2.5
3.0
time

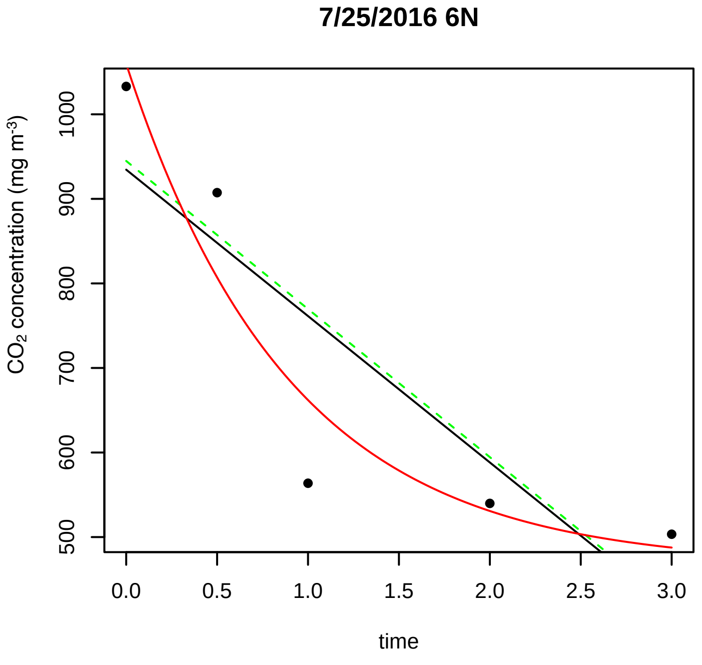
7/25/2016 6N
CO2 concentration (mg m-3)
500
600
700
800
900
1000
0.0
0.5
1.0
1.5
2.0
2.5
3.0
time

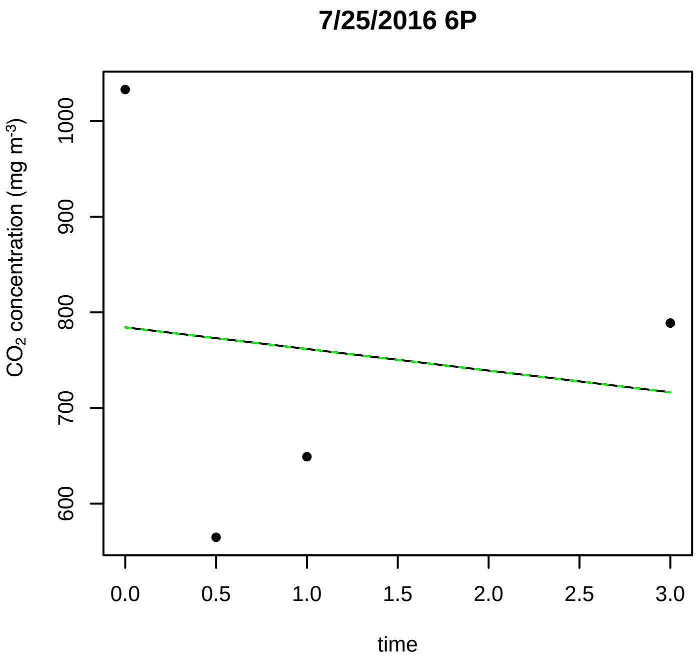
7/25/2016 6P
CO2 concentration (mg m-3)
1000
900
800
700
600
0.0
0.5
1.0
1.5
2.0
2.5
3.0
time

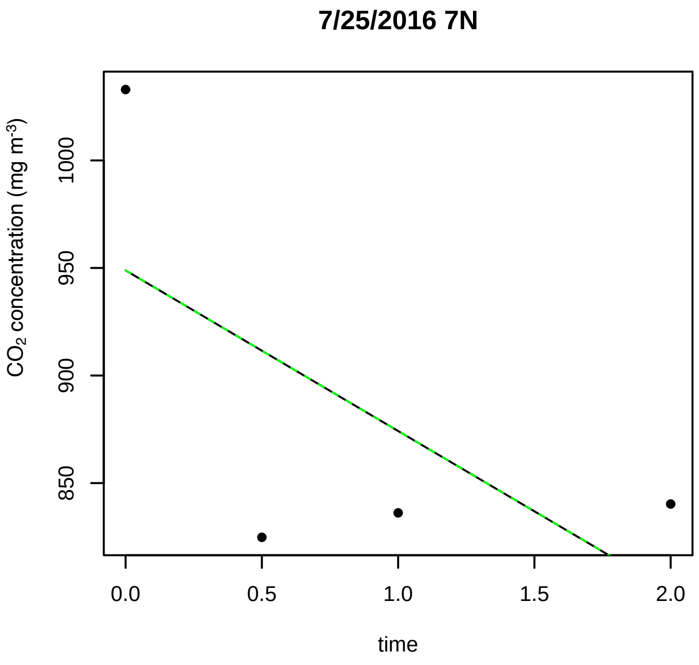
7/25/2016 7N
CO2 concentration (mg m-3)
1000
950
900
850
0.0
0.5
1.0
1.5
2.0
time

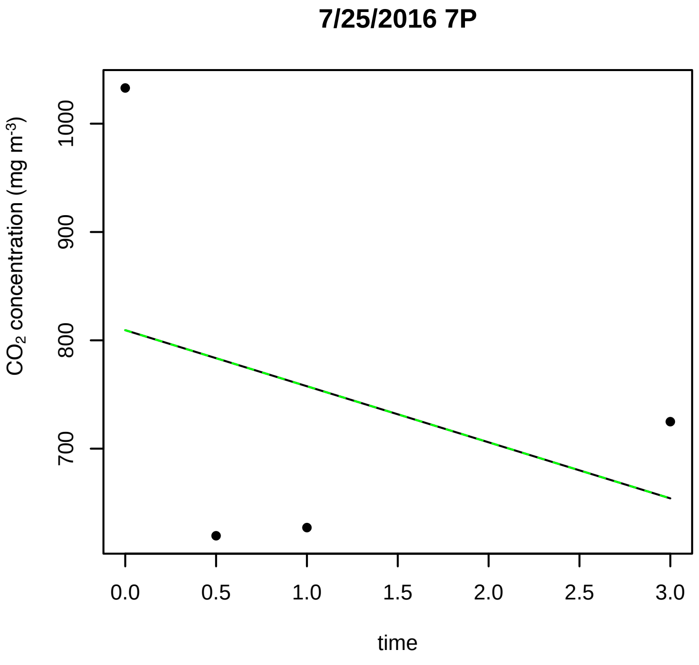
7/25/2016 7P
CO2 concentration (mg m-3)
1000
900
800
700
0.0
0.5
1.0
1.5
2.0
2.5
3.0
time

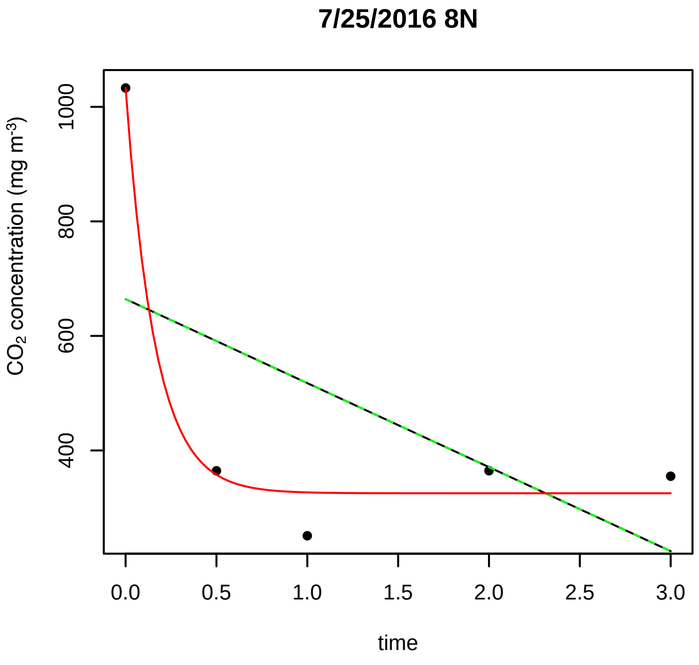
7/25/2016 8N
CO2 concentration (mg m-3)
1000
800
600
400
0.0
0.5
1.0
1.5
2.0
2.5
3.0
time

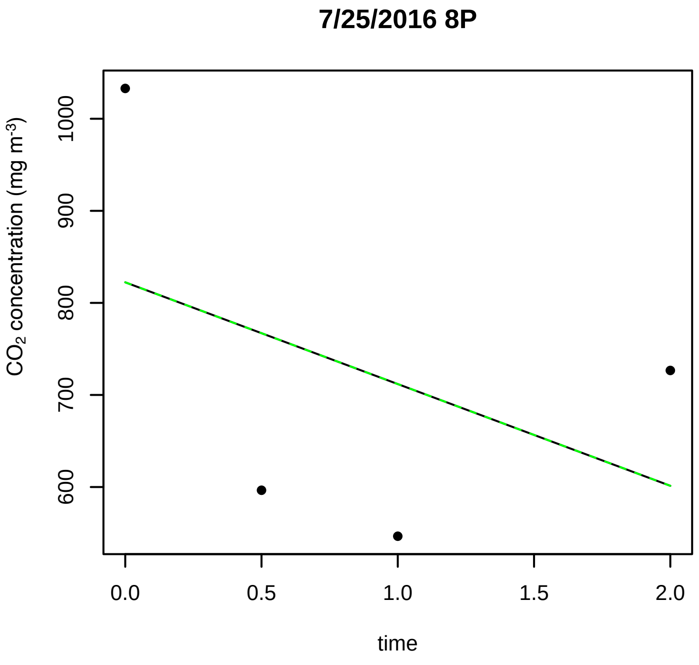
7/25/2016 8P
CO2 concentration (mg m-3)
1000
900
800
700
600
0.0
0.5
1.0
1.5
2.0
time

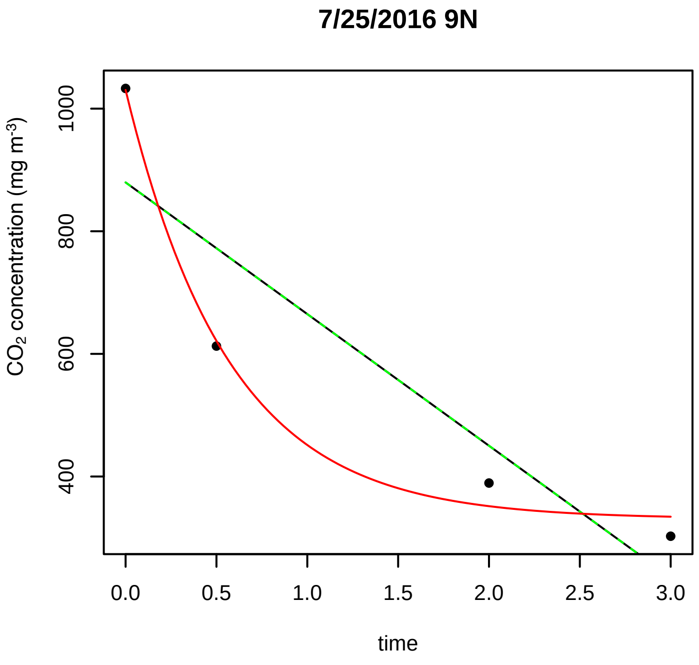
7/25/2016 9N
CO2 concentration (mg m-3)
1000
800
600
400
0.0
0.5
1.0
1.5
2.0
2.5
3.0
time

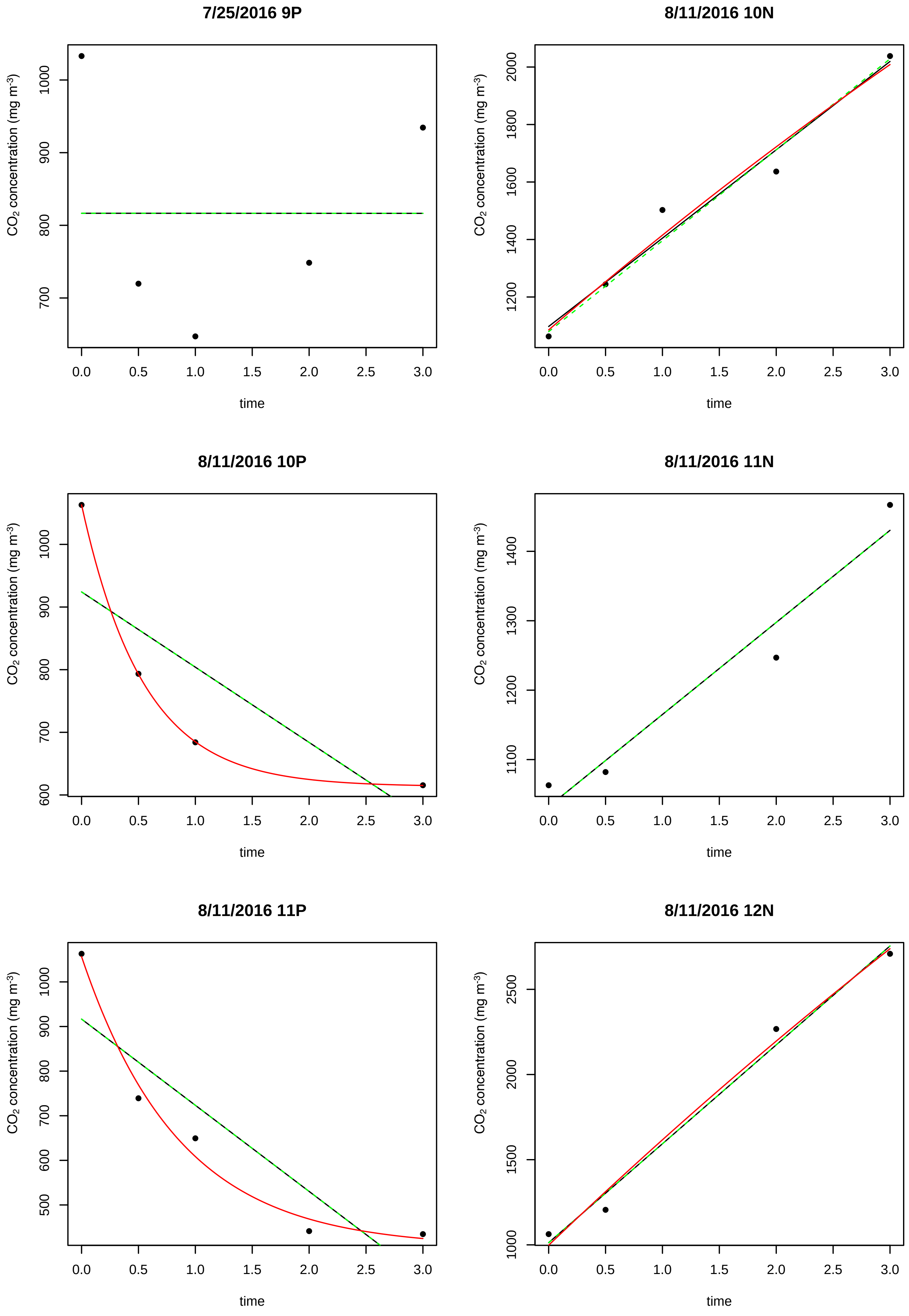

7/25/2016 9P
8/11/2016 10N
8/11/2016 10P
8/11/2016 11N
8/11/2016 11P
8/11/2016 12N
CO2 concentration (mg m-3)
time

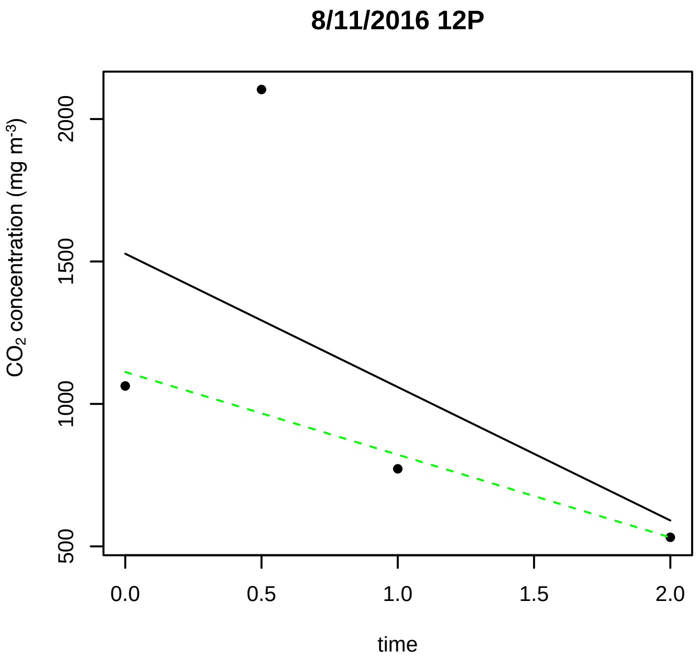
8/11/2016 12P
$CO_2$ concentration (mg m$^{-3}$)
time

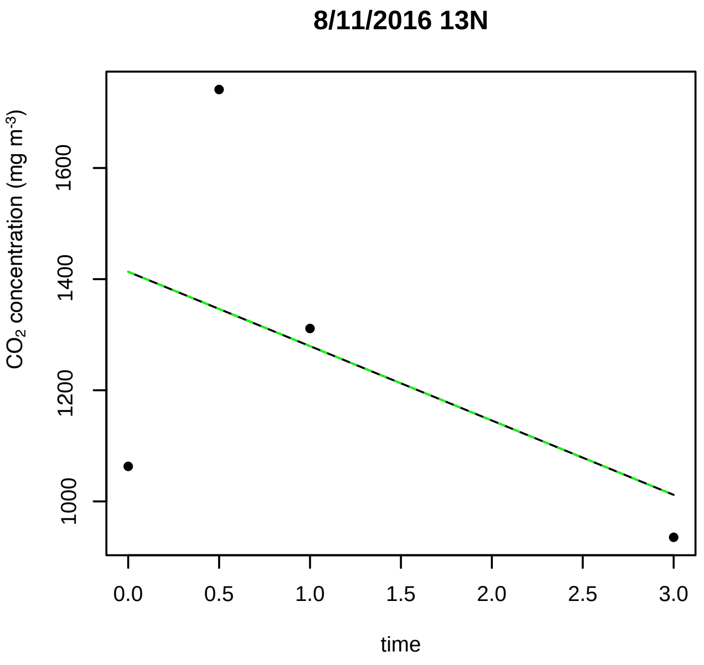
8/11/2016 13N
$CO_2$ concentration (mg m$^{-3}$)
time

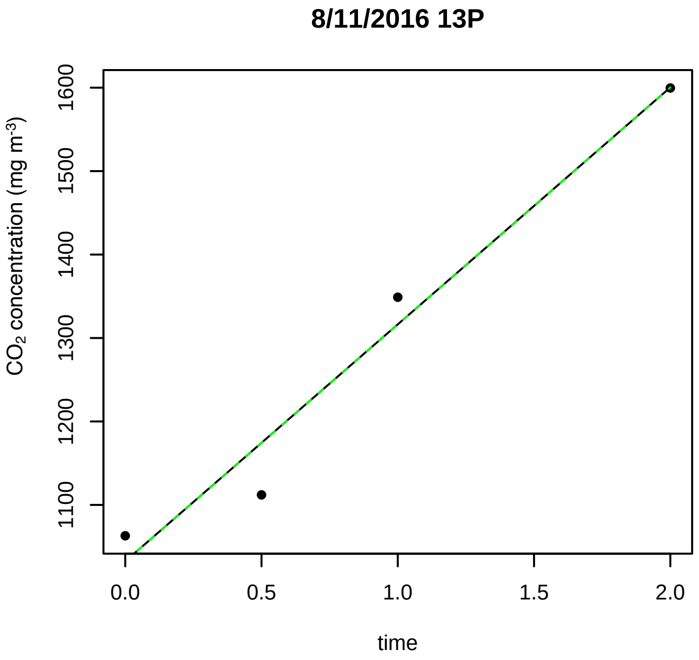
8/11/2016 13P
$CO_2$ concentration (mg m$^{-3}$)
time

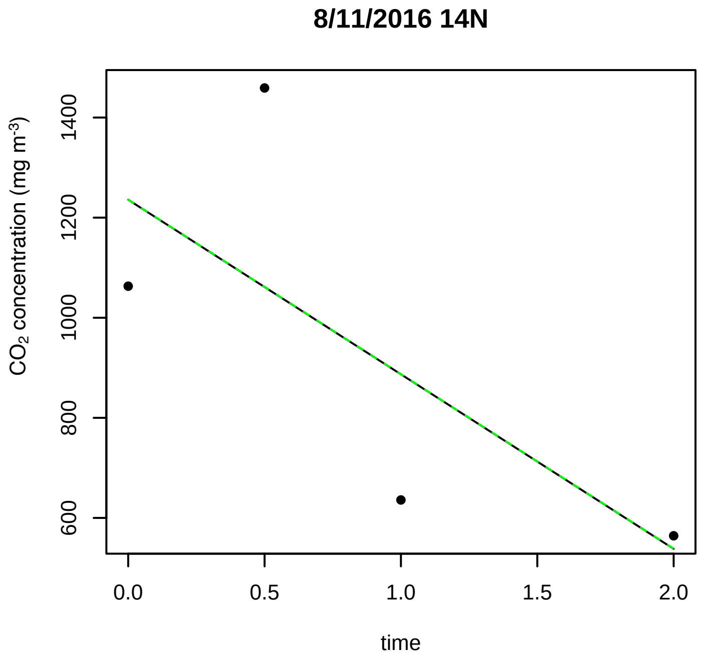
8/11/2016 14N
$CO_2$ concentration (mg m$^{-3}$)
time

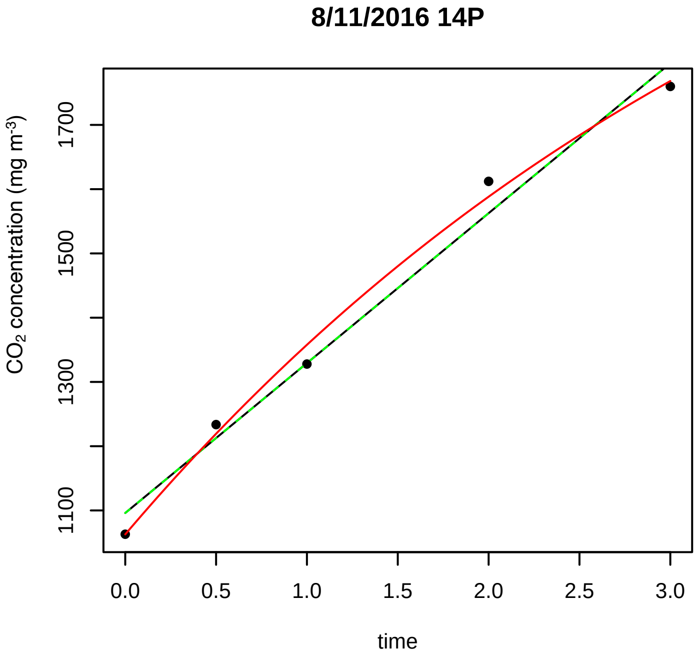
8/11/2016 14P
$CO_2$ concentration (mg m$^{-3}$)
time

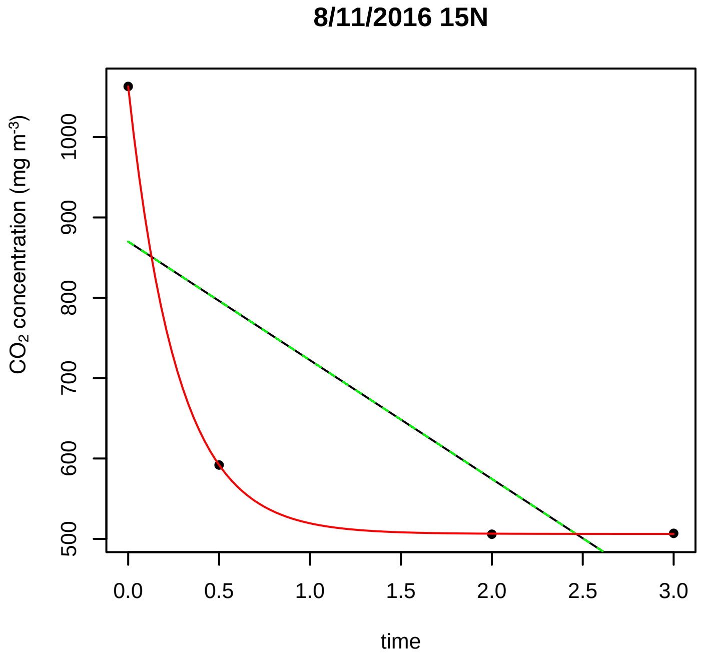
8/11/2016 15N
$CO_2$ concentration (mg m$^{-3}$)
time

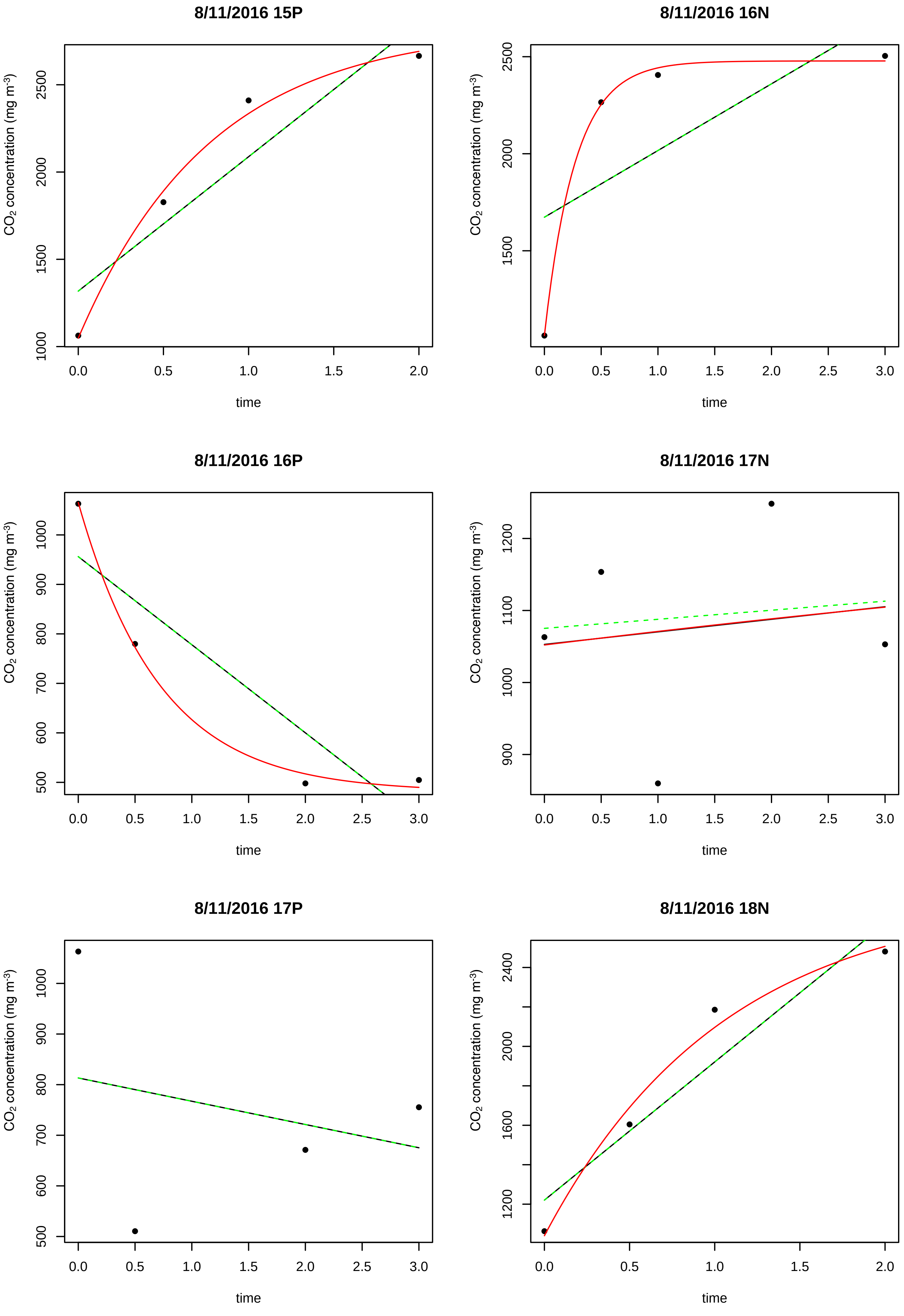
8/11/2016 15P
8/11/2016 16N
8/11/2016 16P
8/11/2016 17N
8/11/2016 17P
8/11/2016 18N
CO2 concentration (mg m-3)
time

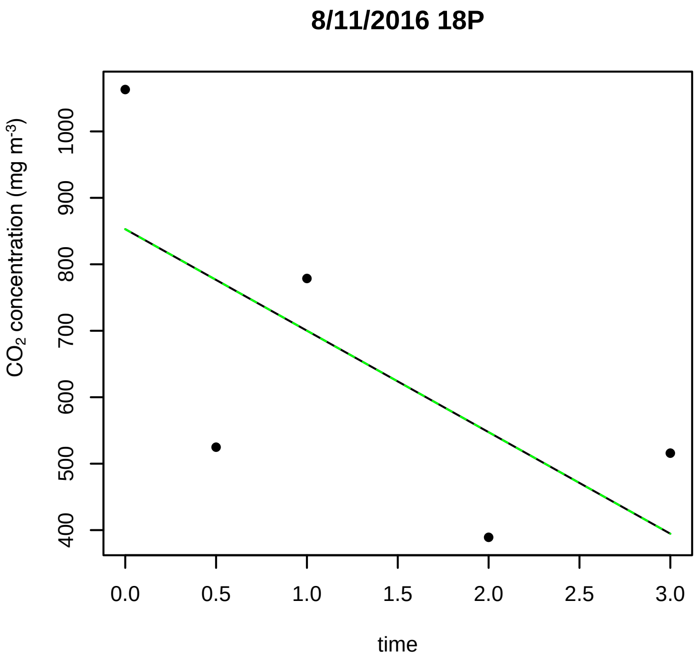
8/11/2016 18P
CO2 concentration (mg m-3)
time

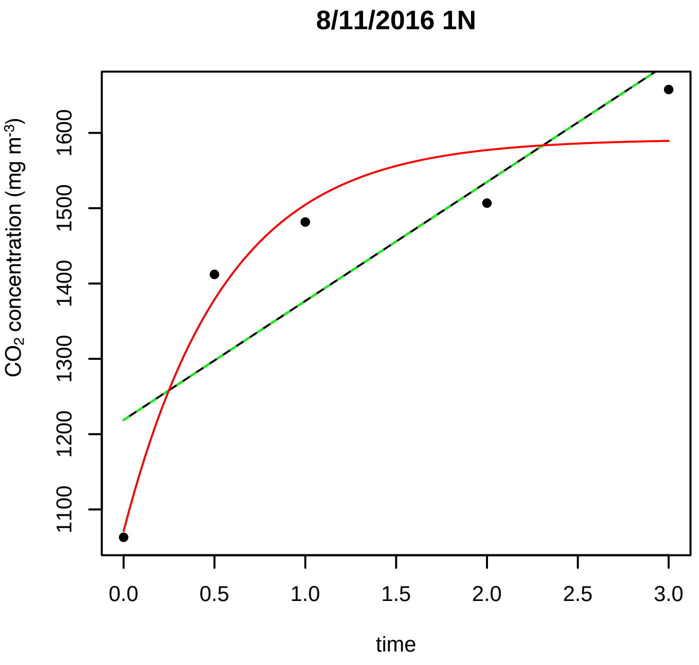
8/11/2016 1N
CO2 concentration (mg m-3)
time

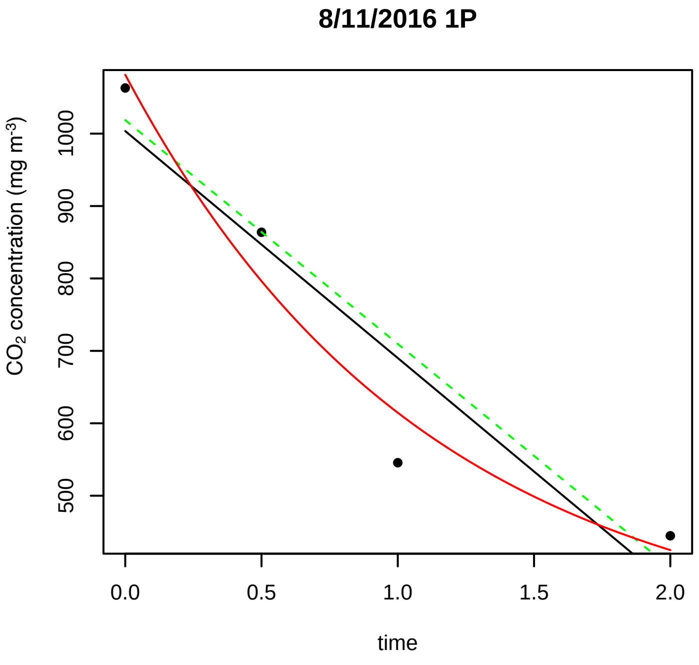
8/11/2016 1P
CO2 concentration (mg m-3)
time

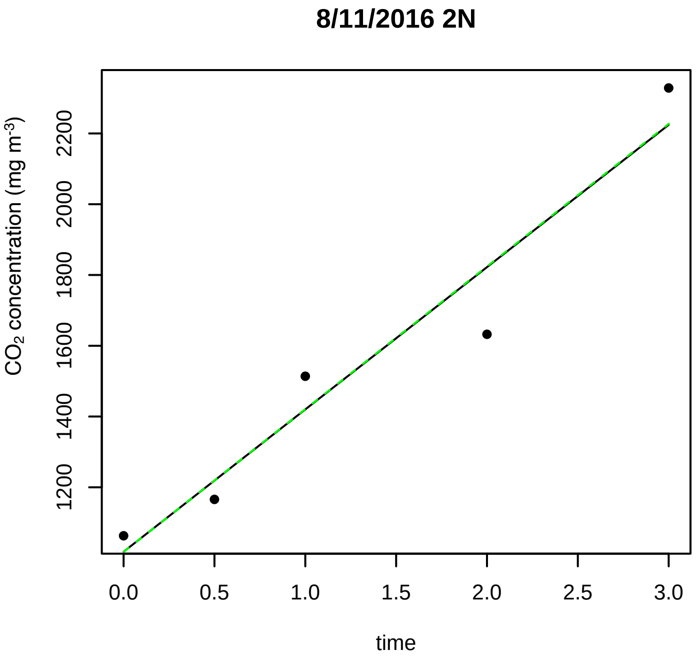
8/11/2016 2N
CO2 concentration (mg m-3)
time

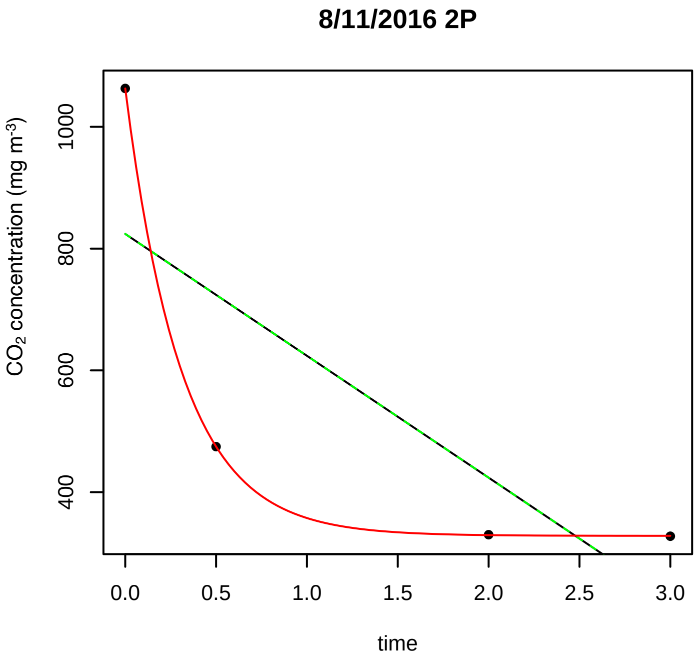
8/11/2016 2P
CO2 concentration (mg m-3)
time

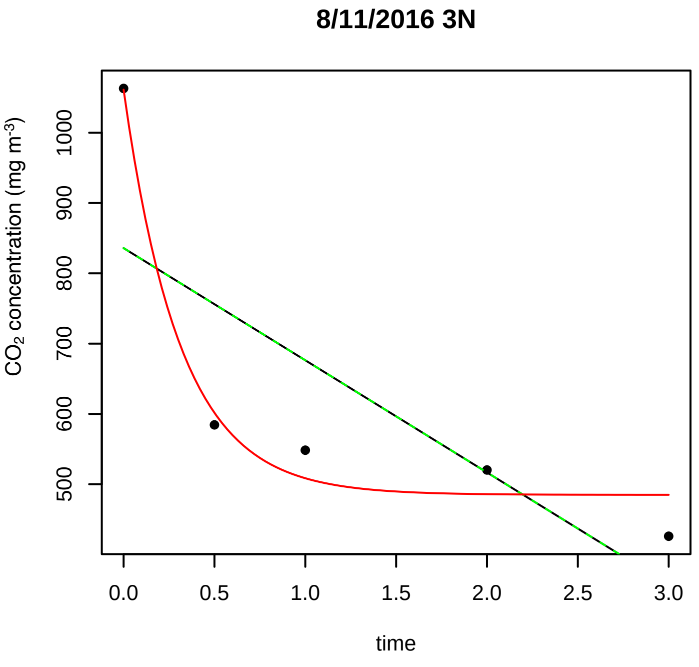
8/11/2016 3N
CO2 concentration (mg m-3)
time

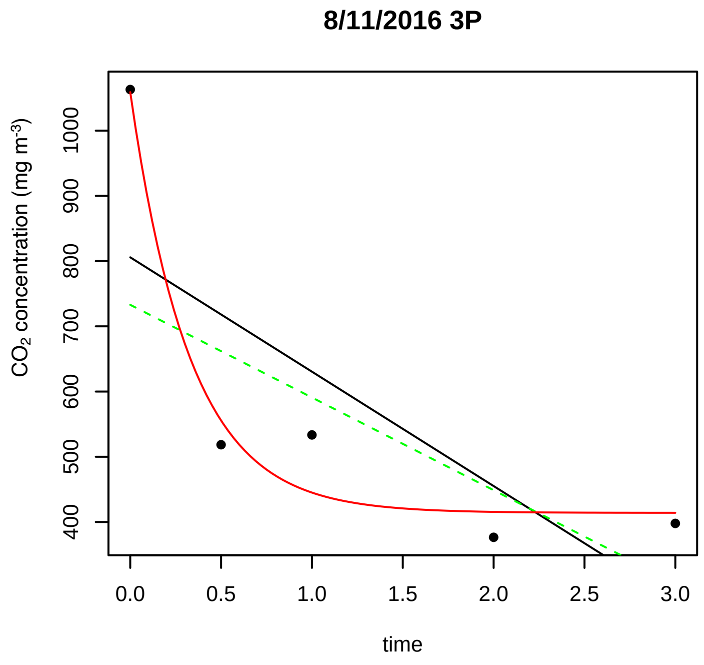
8/11/2016 3P
CO2 concentration (mg m-3)
400 500 600 700 800 900 1000
0.0 0.5 1.0 1.5 2.0 2.5 3.0
time

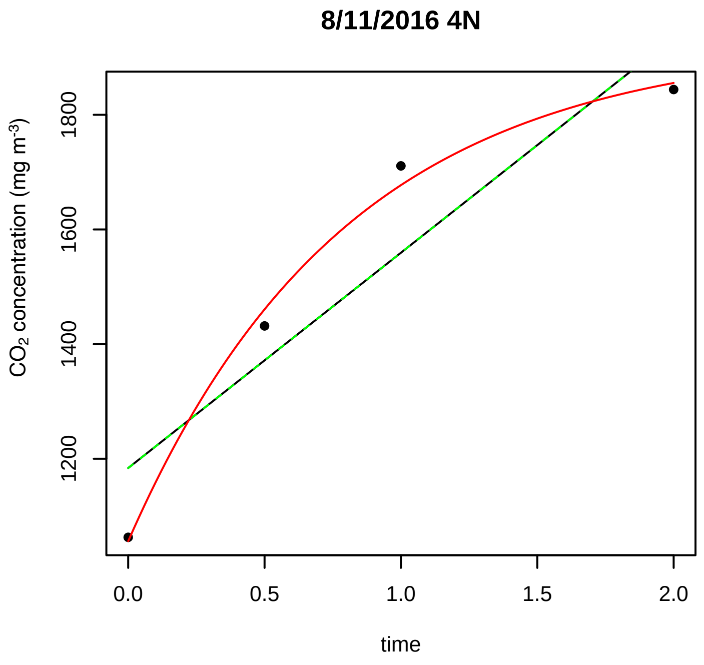
8/11/2016 4N
CO2 concentration (mg m-3)
1200 1400 1600 1800
0.0 0.5 1.0 1.5 2.0
time

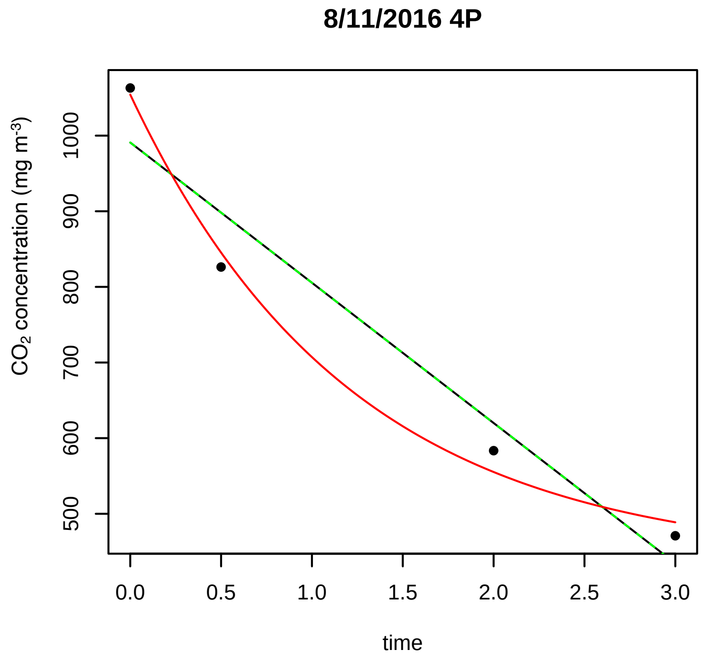
8/11/2016 4P
CO2 concentration (mg m-3)
500 600 700 800 900 1000
0.0 0.5 1.0 1.5 2.0 2.5 3.0
time

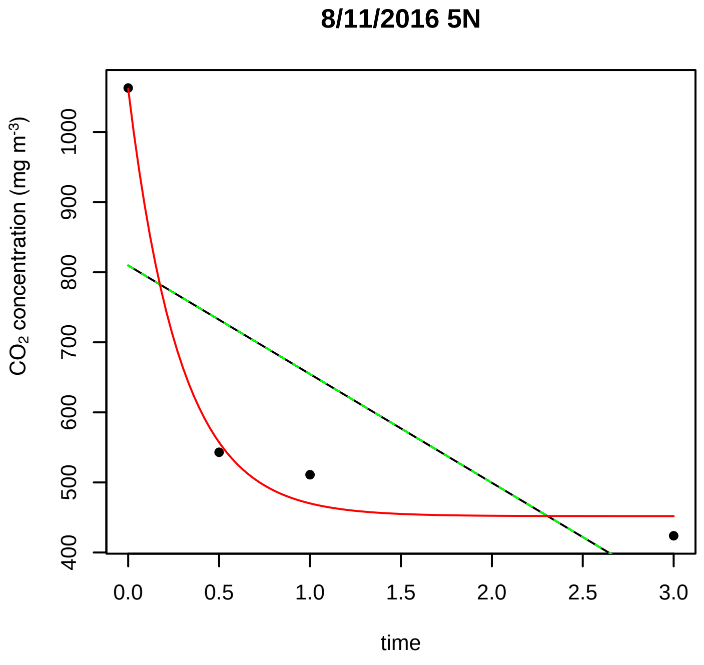
8/11/2016 5N
CO2 concentration (mg m-3)
400 500 600 700 800 900 1000
0.0 0.5 1.0 1.5 2.0 2.5 3.0
time

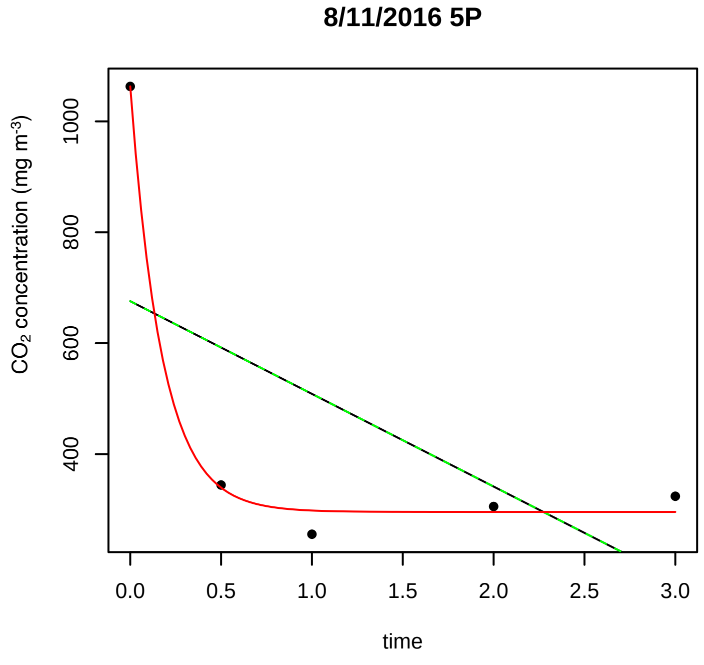
8/11/2016 5P
CO2 concentration (mg m-3)
400 600 800 1000
0.0 0.5 1.0 1.5 2.0 2.5 3.0
time

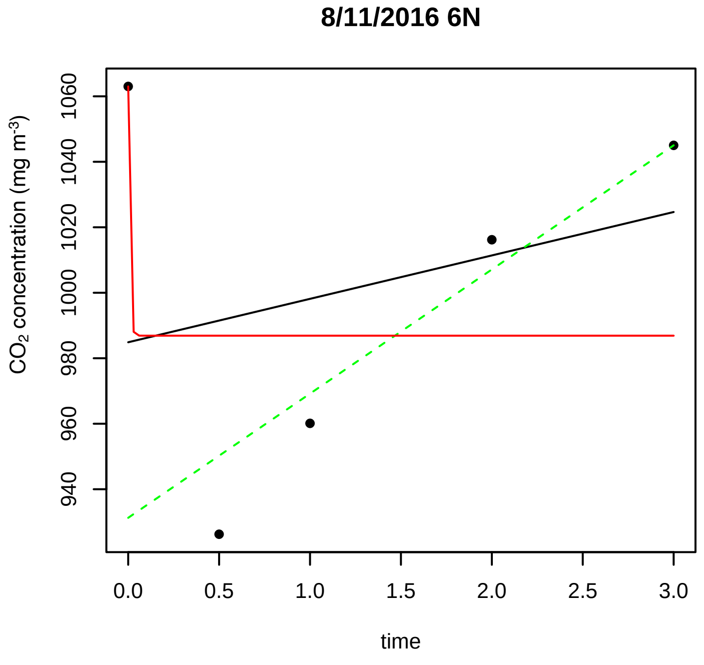
8/11/2016 6N
CO2 concentration (mg m-3)
940 960 980 1000 1020 1040 1060
0.0 0.5 1.0 1.5 2.0 2.5 3.0
time

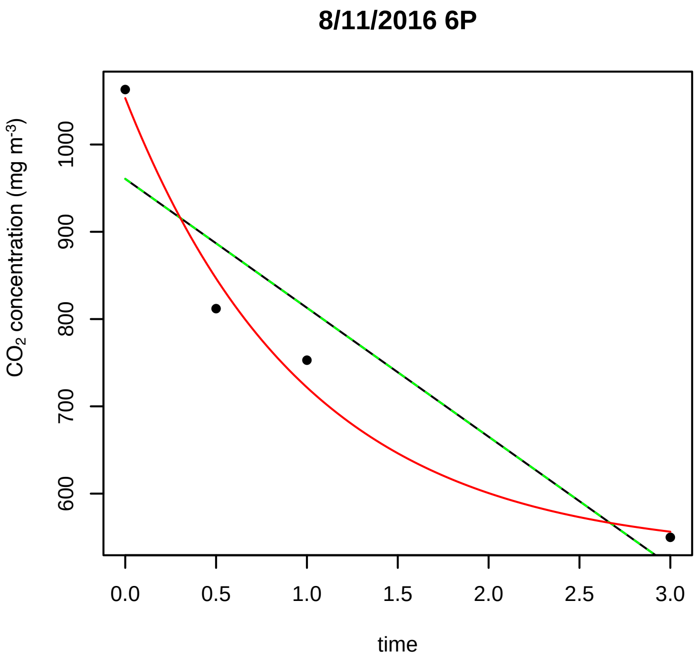
8/11/2016 6P
CO2 concentration (mg m-3)
600
700
800
900
1000
0.0
0.5
1.0
1.5
2.0
2.5
3.0
time

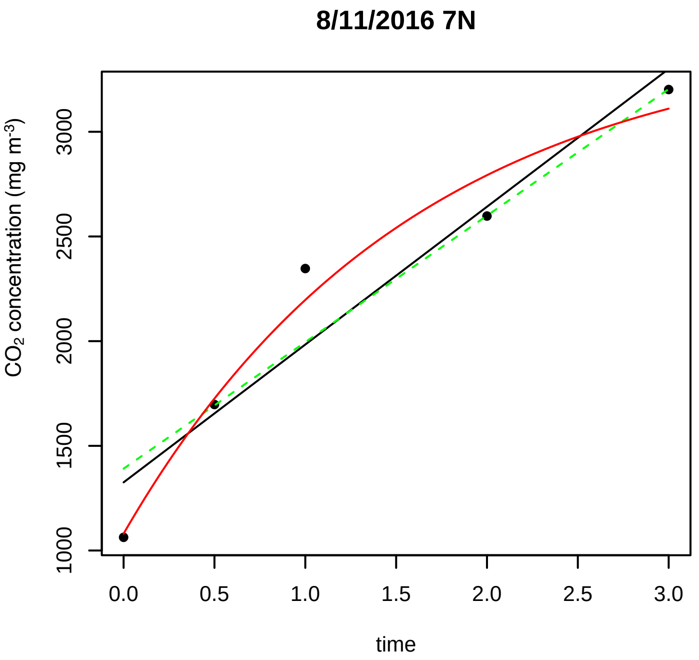
8/11/2016 7N
CO2 concentration (mg m-3)
1000
1500
2000
2500
3000
0.0
0.5
1.0
1.5
2.0
2.5
3.0
time

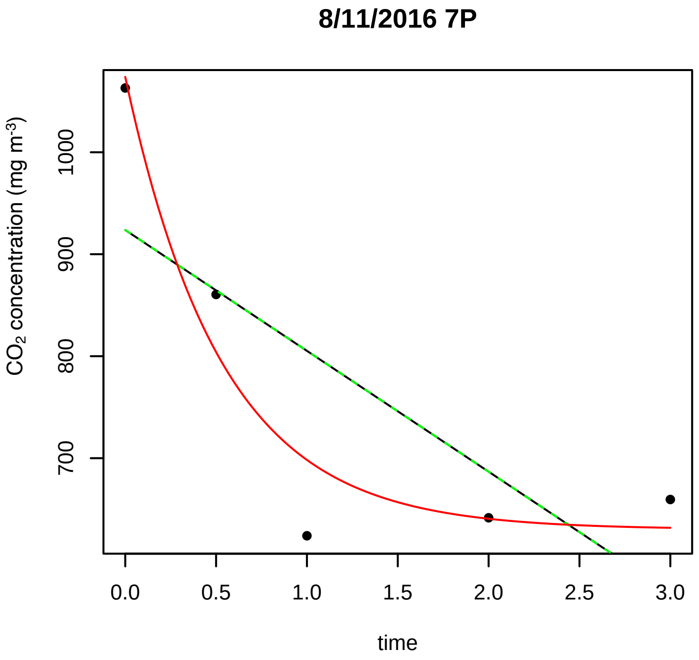
8/11/2016 7P
CO2 concentration (mg m-3)
700
800
900
1000
0.0
0.5
1.0
1.5
2.0
2.5
3.0
time

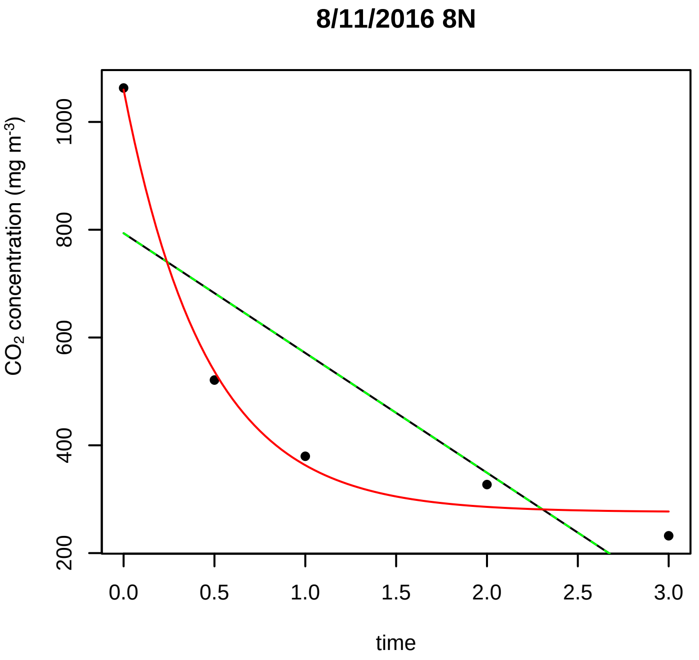
8/11/2016 8N
CO2 concentration (mg m-3)
200
400
600
800
1000
0.0
0.5
1.0
1.5
2.0
2.5
3.0
time

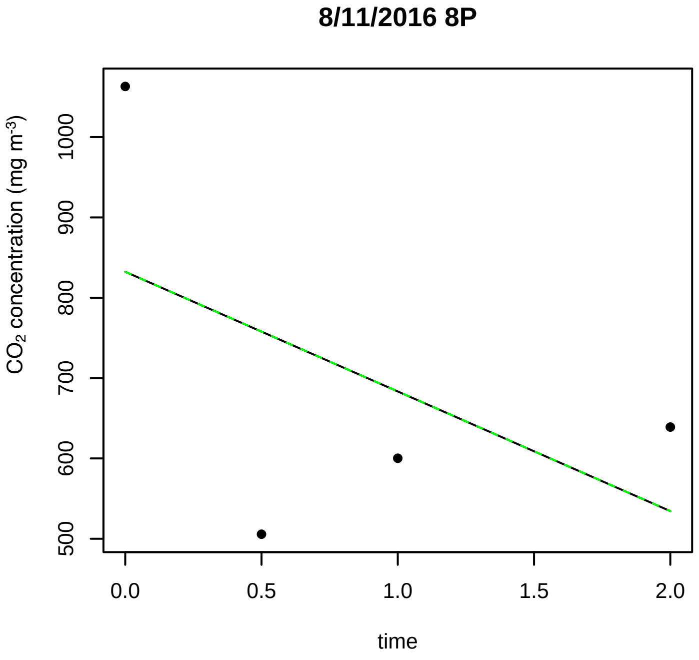
8/11/2016 8P
CO2 concentration (mg m-3)
500
600
700
800
900
1000
0.0
0.5
1.0
1.5
2.0
time

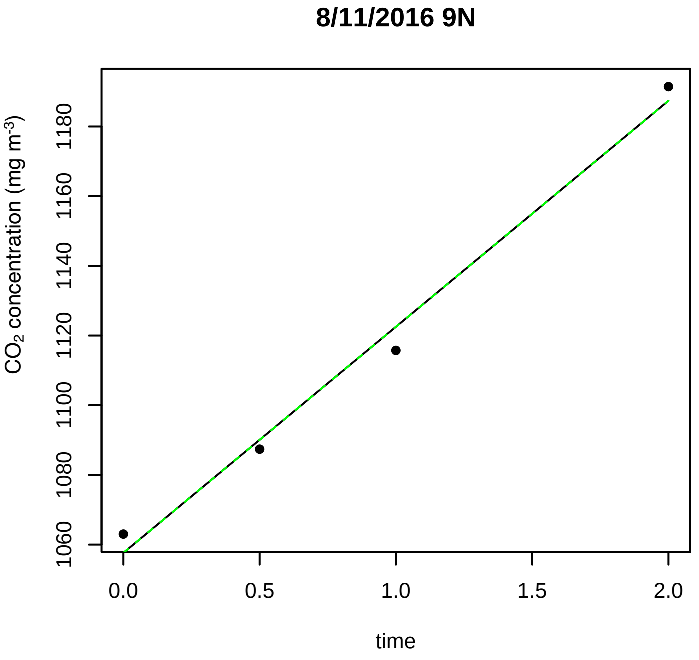
8/11/2016 9N
CO2 concentration (mg m-3)
1060
1080
1100
1120
1140
1160
1180
0.0
0.5
1.0
1.5
2.0
time

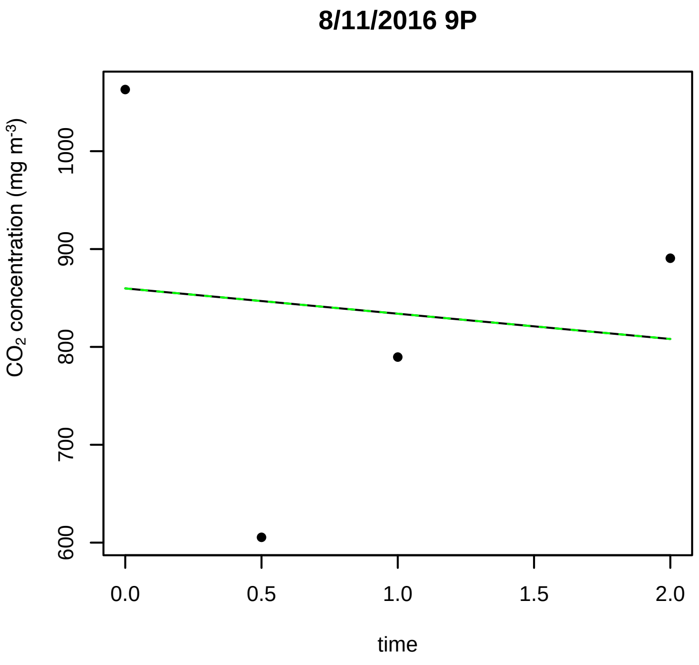

8/11/2016 9P
CO2 concentration (mg m-3)
1000
900
800
700
600
0.0
0.5
1.0
1.5
2.0
time

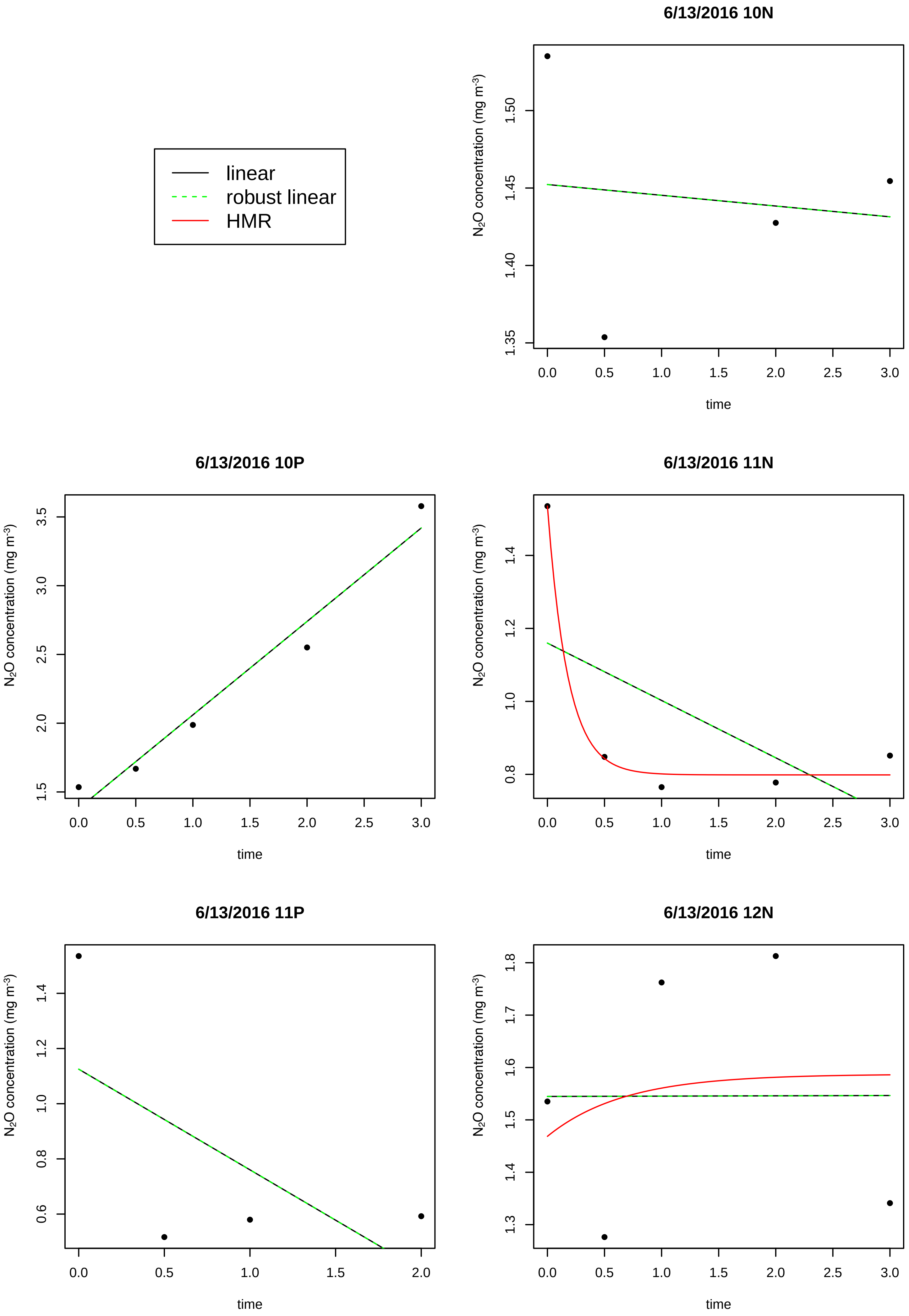

linear
robust linear
HMR
6/13/2016 10N
6/13/2016 10P
6/13/2016 11N
6/13/2016 11P
6/13/2016 12N
N2O concentration (mg m-3)
time

6/13/2016 12P

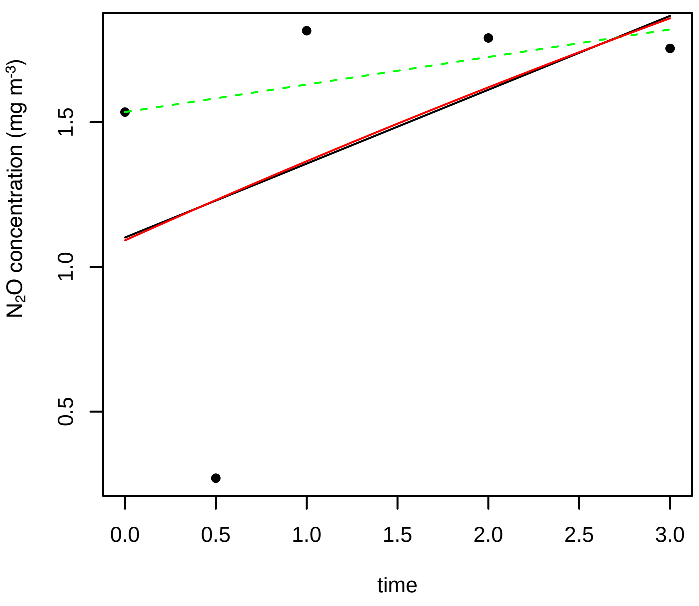


6/13/2016 13N

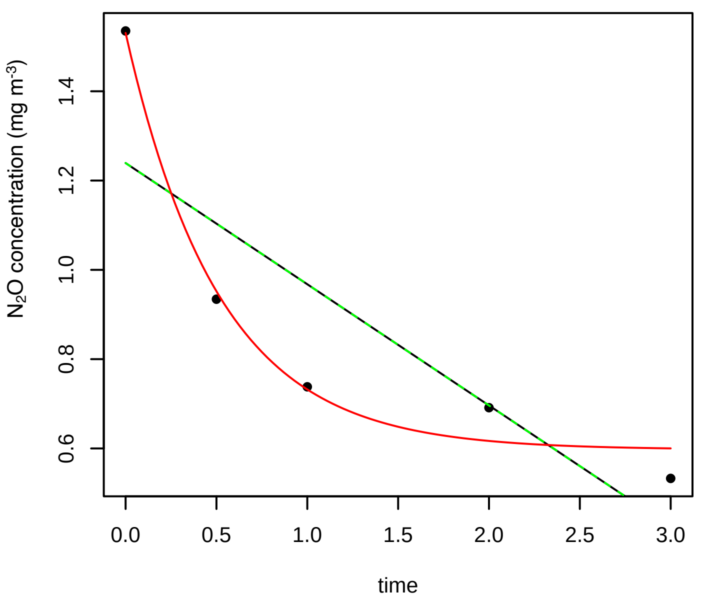


6/13/2016 13P

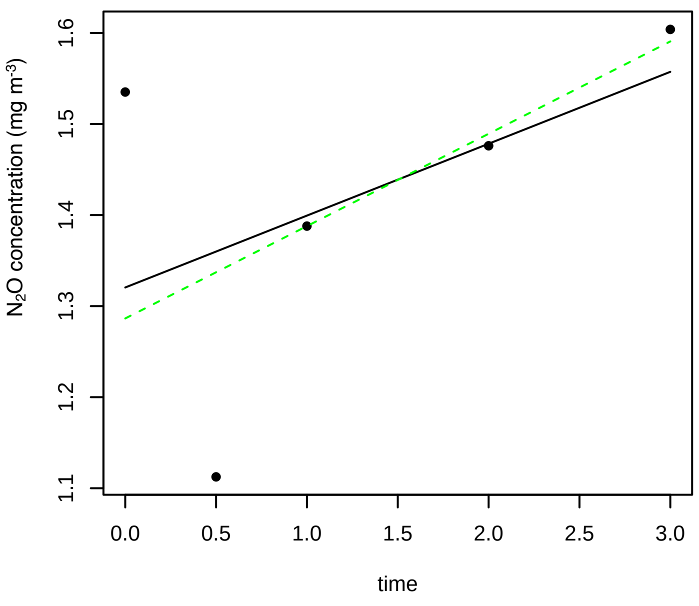


6/13/2016 14N

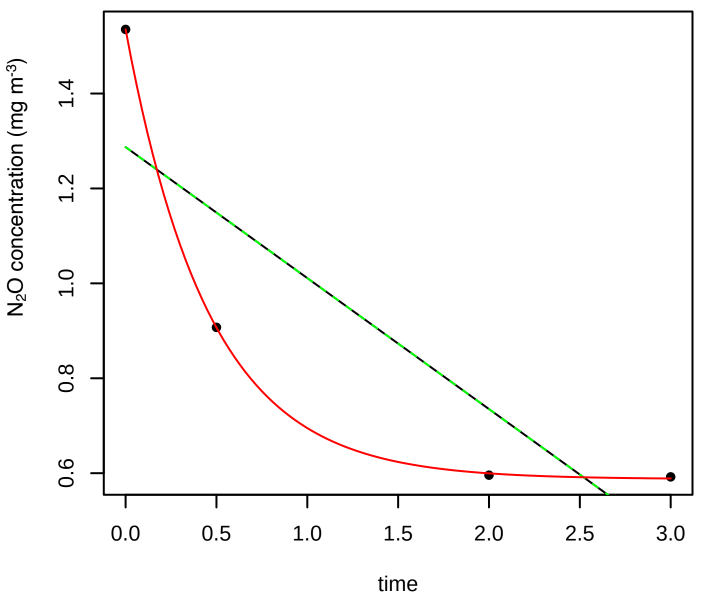


6/13/2016 14P

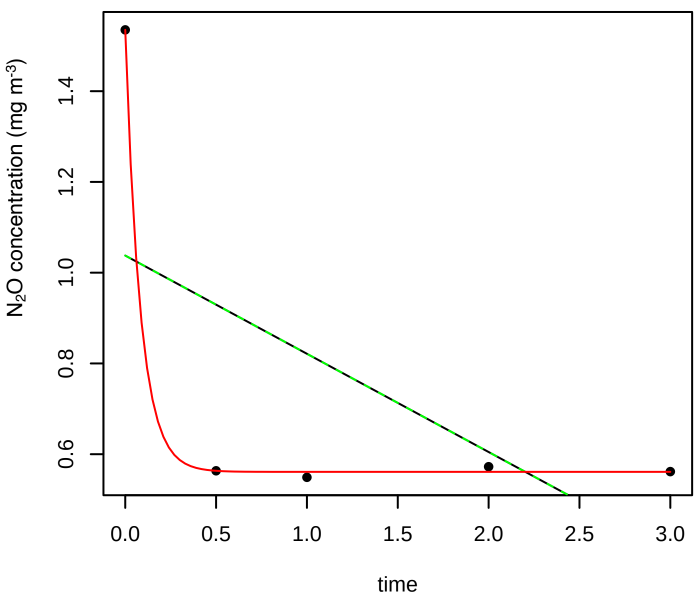


6/13/2016 15N

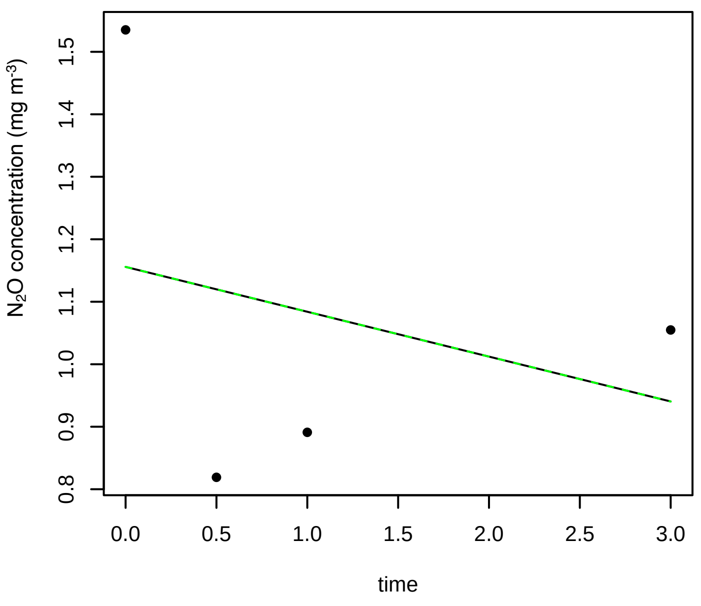

**6/13/2016 15P**

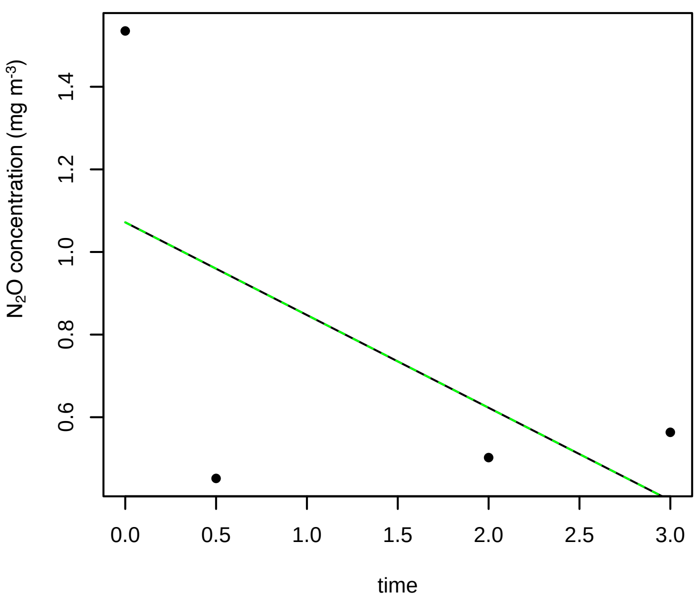


**6/13/2016 16N**

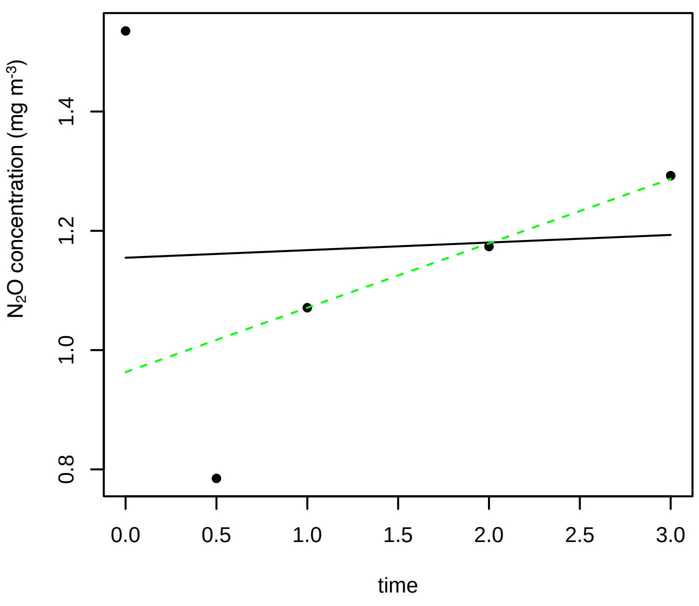


**6/13/2016 16P**

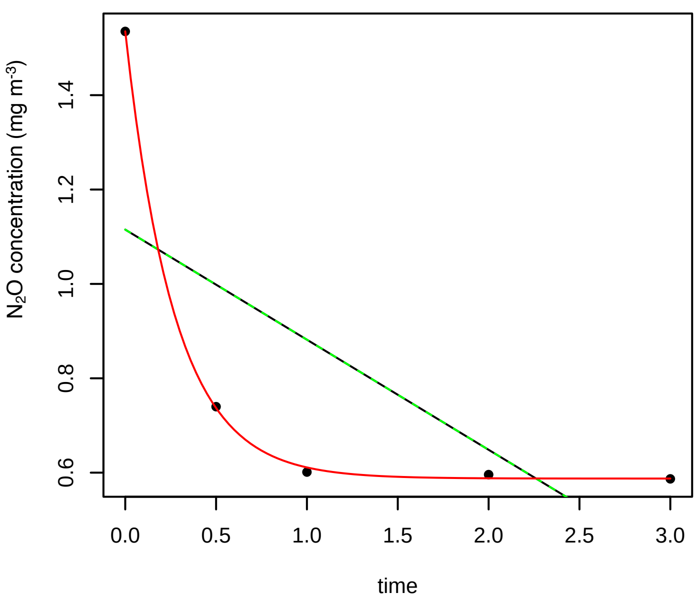


**6/13/2016 17N**

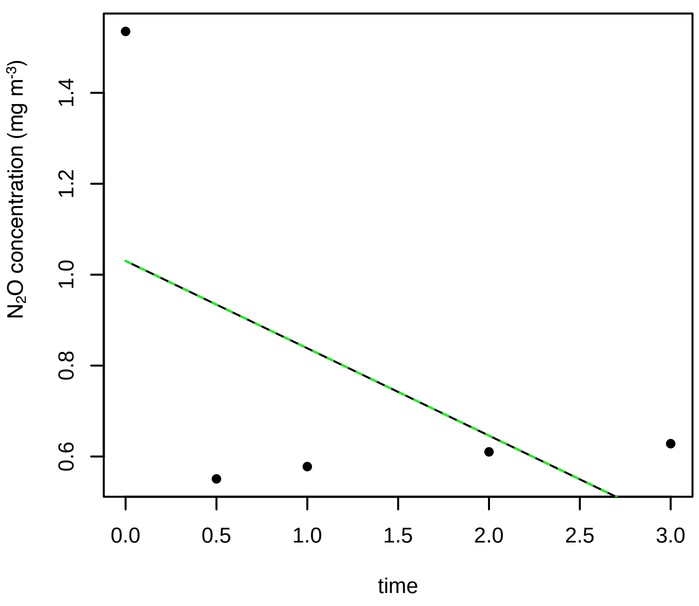


**6/13/2016 17P**

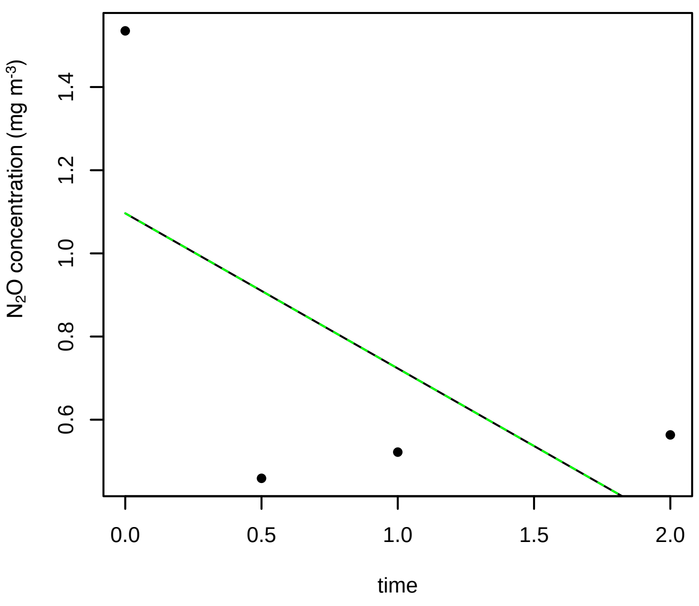


**6/13/2016 18N**

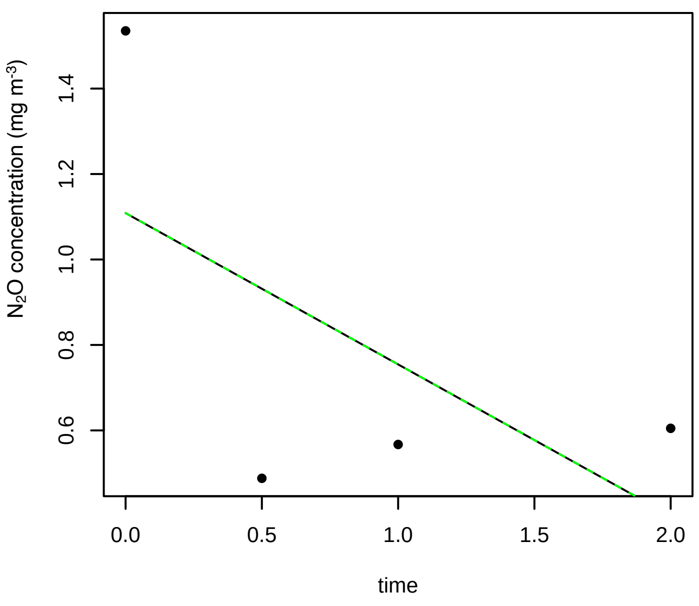

## 6/13/2016 18P

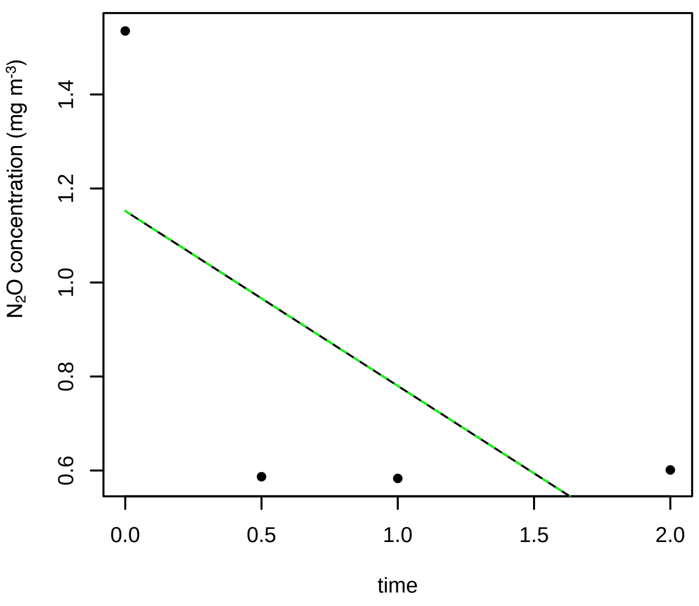


## 6/13/2016 1N

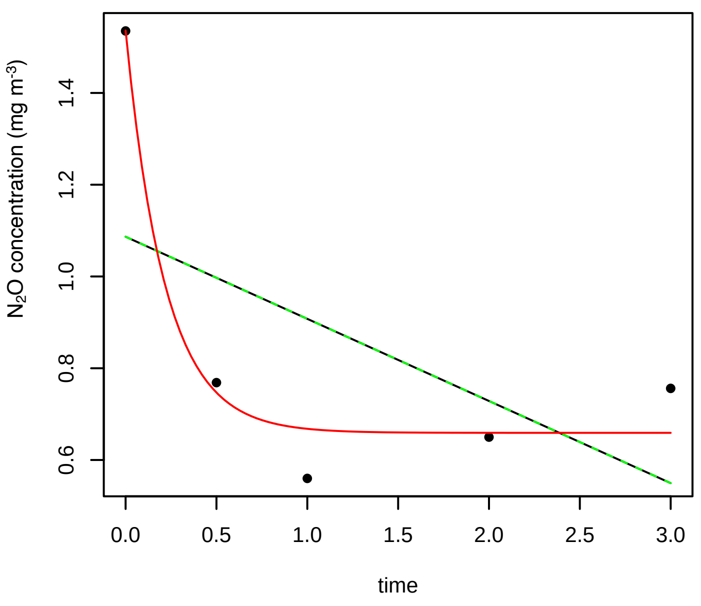


## 6/13/2016 1P

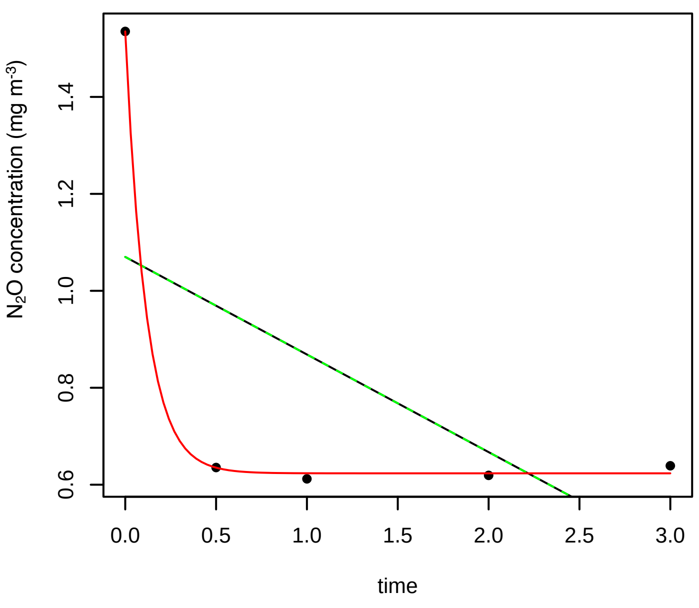


## 6/13/2016 2N

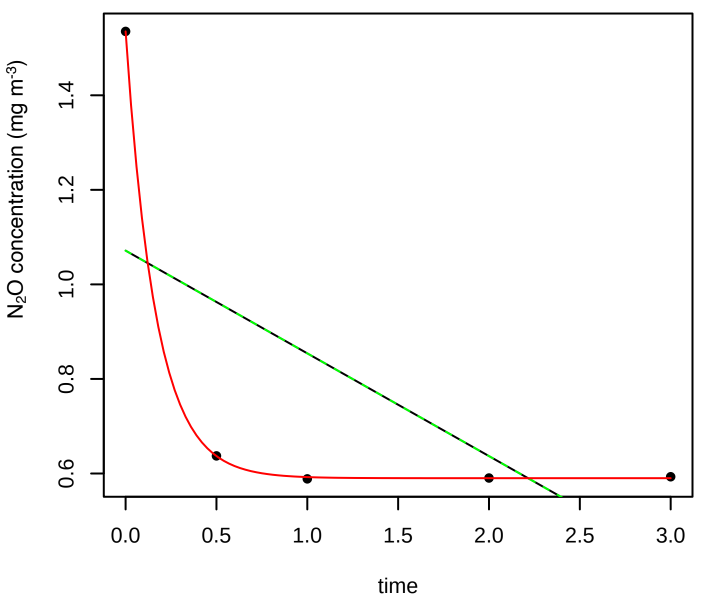


## 6/13/2016 2P

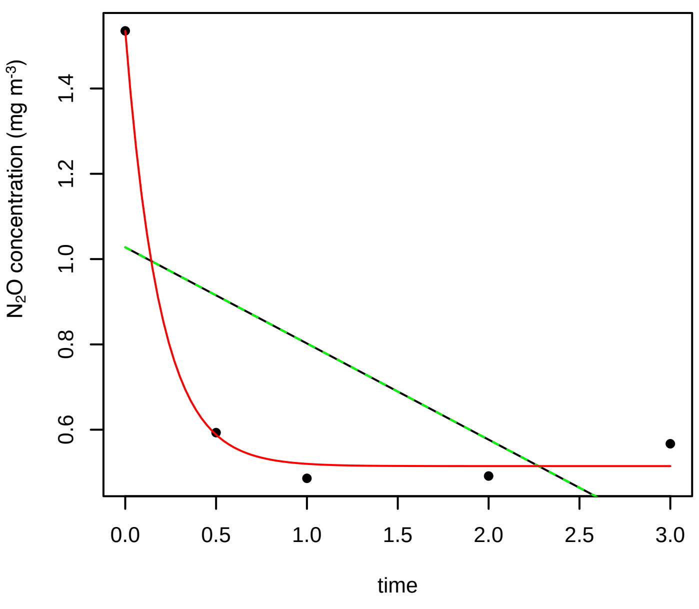


## 6/13/2016 3N

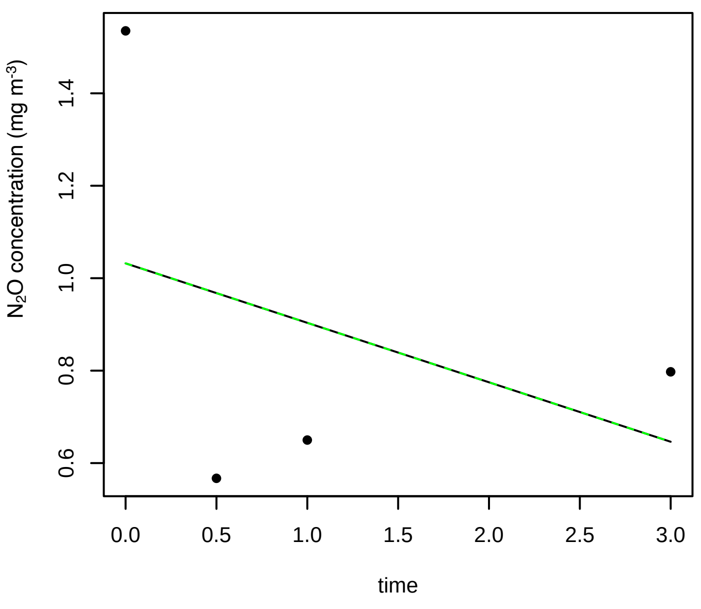

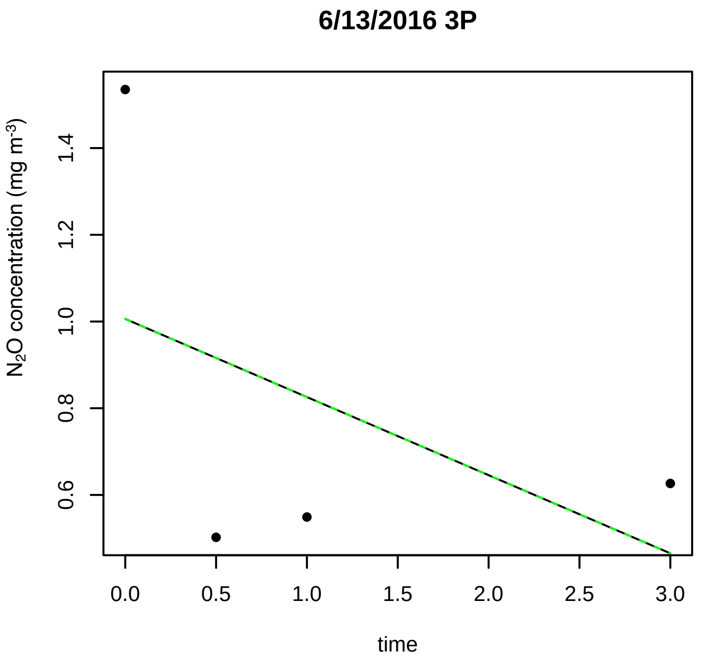
6/13/2016 3P
N2O concentration (mg m-3)
time

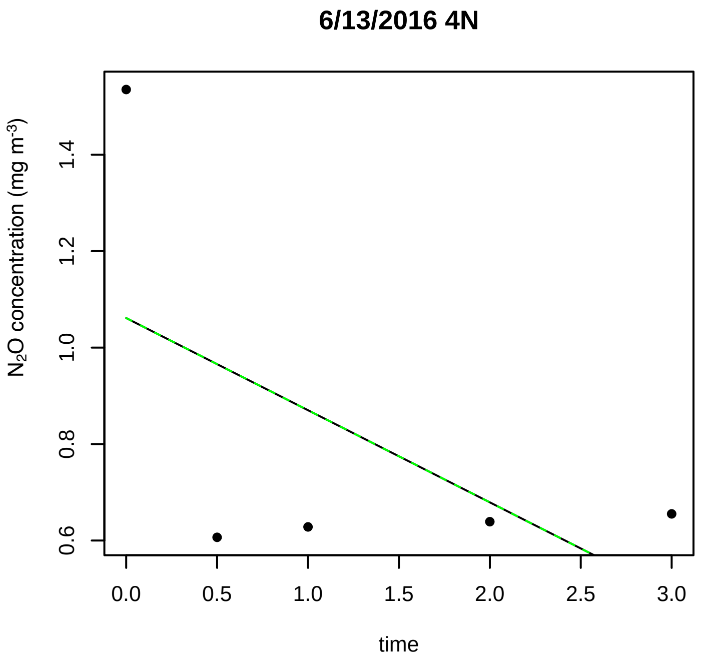
6/13/2016 4N
N2O concentration (mg m-3)
time

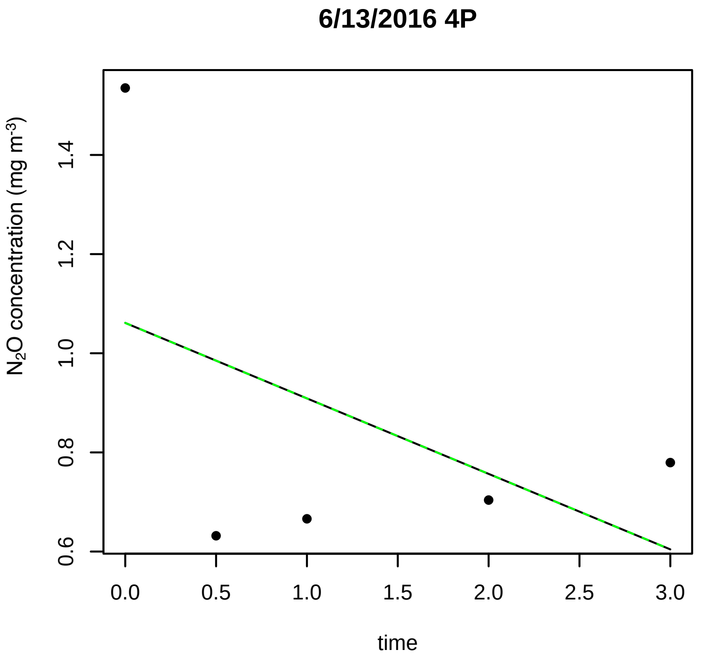
6/13/2016 4P
N2O concentration (mg m-3)
time

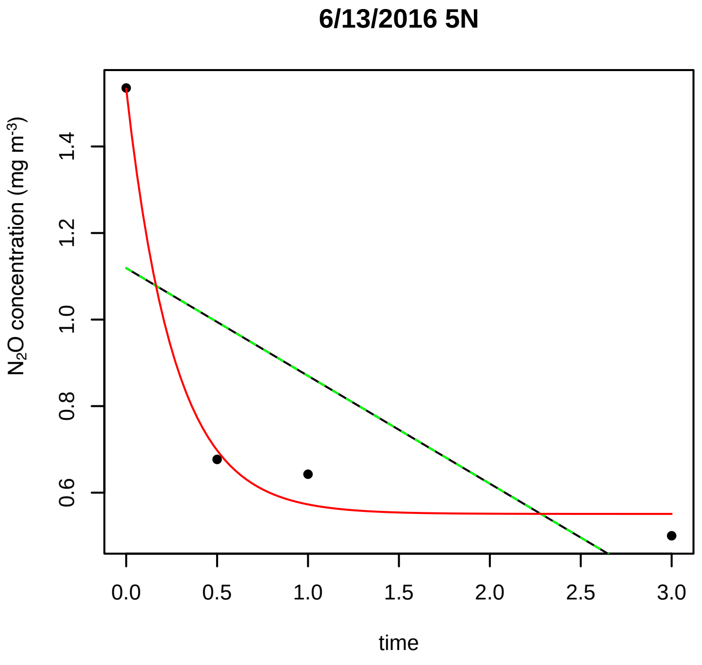
6/13/2016 5N
N2O concentration (mg m-3)
time

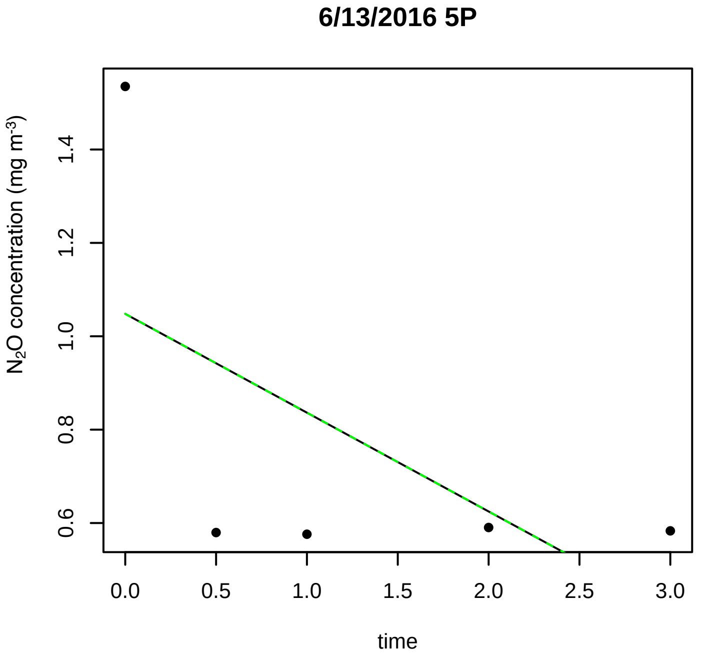
6/13/2016 5P
N2O concentration (mg m-3)
time

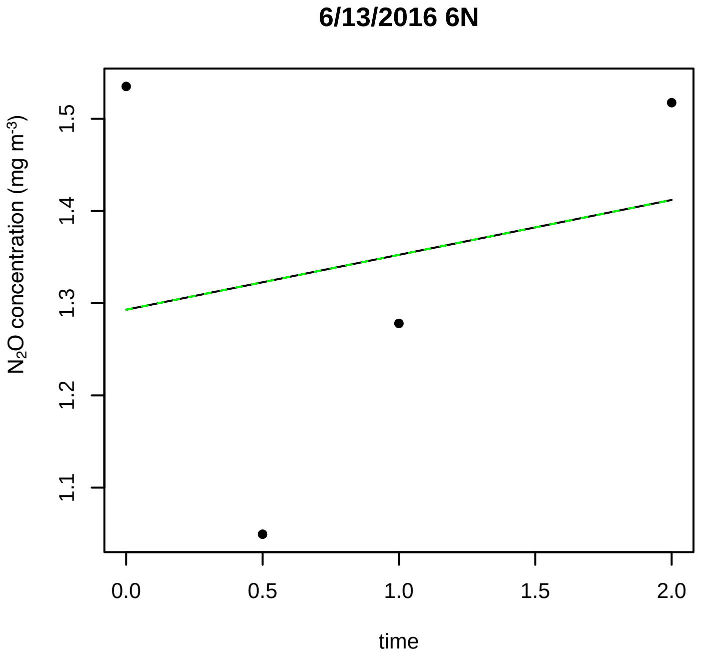
6/13/2016 6N
N2O concentration (mg m-3)
time

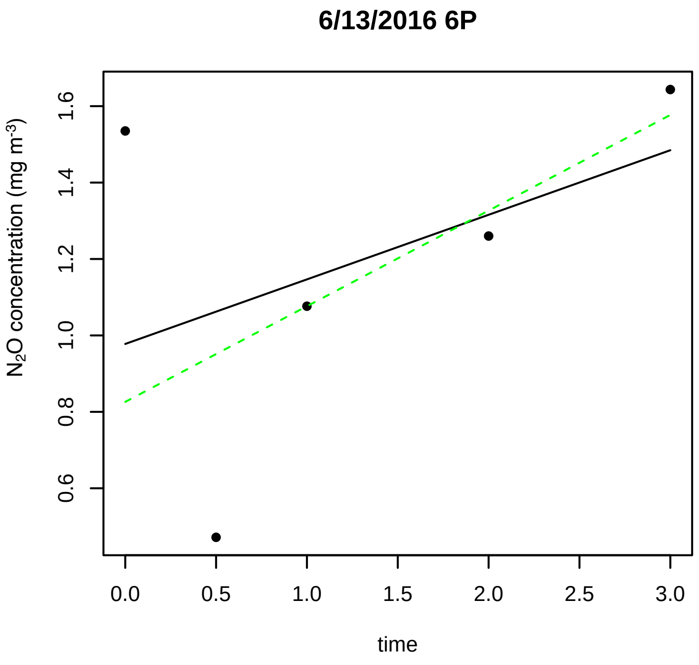
6/13/2016 6P
N2O concentration (mg m-3)
time
0.0 0.5 1.0 1.5 2.0 2.5 3.0
0.6 0.8 1.0 1.2 1.4 1.6

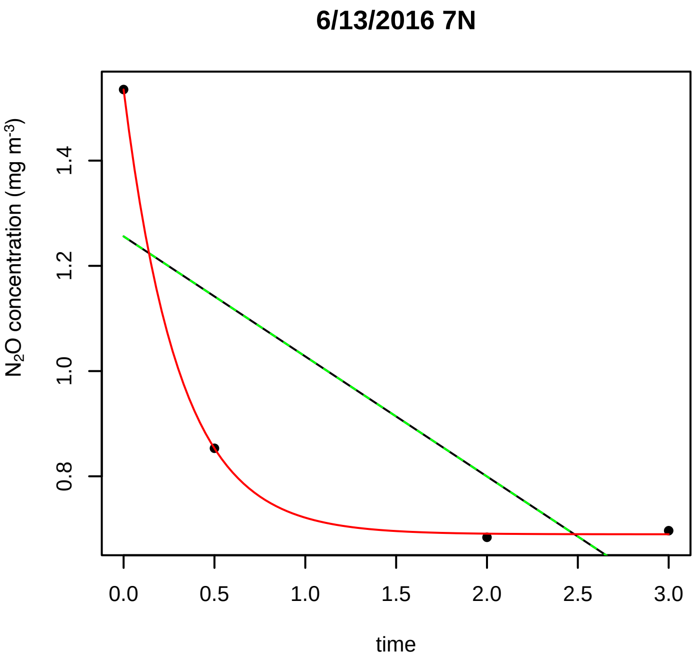
6/13/2016 7N
N2O concentration (mg m-3)
time
0.0 0.5 1.0 1.5 2.0 2.5 3.0
0.8 1.0 1.2 1.4

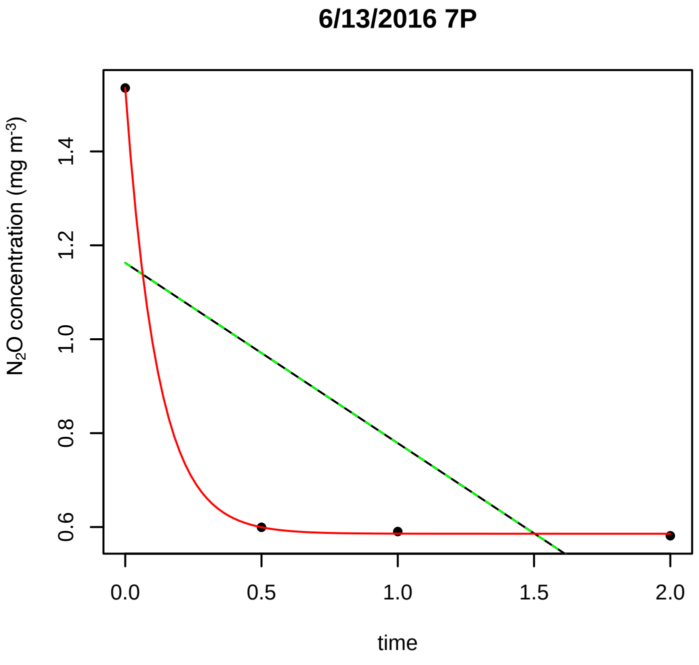
6/13/2016 7P
N2O concentration (mg m-3)
time
0.0 0.5 1.0 1.5 2.0
0.6 0.8 1.0 1.2 1.4

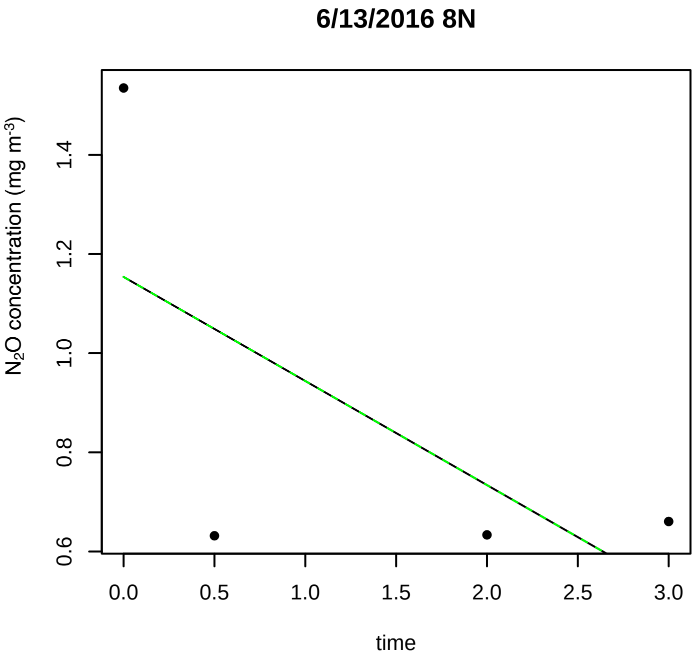
6/13/2016 8N
N2O concentration (mg m-3)
time
0.0 0.5 1.0 1.5 2.0 2.5 3.0
0.6 0.8 1.0 1.2 1.4

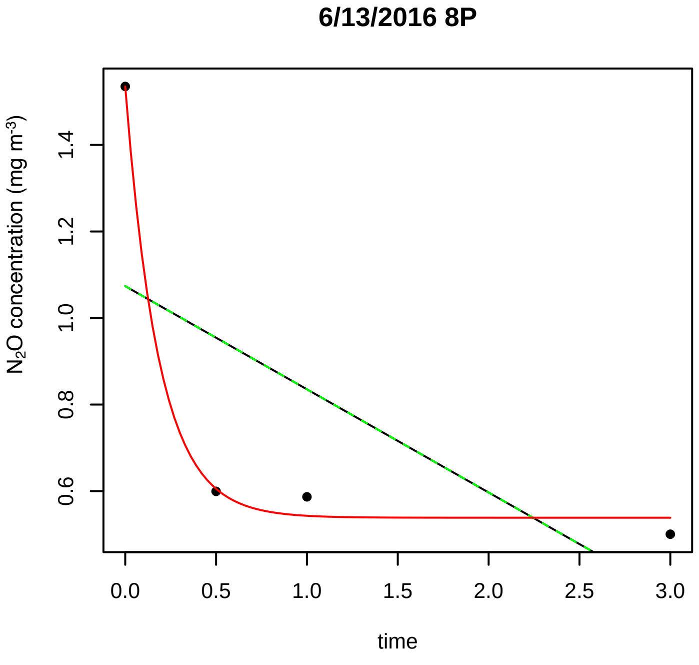
6/13/2016 8P
N2O concentration (mg m-3)
time
0.0 0.5 1.0 1.5 2.0 2.5 3.0
0.6 0.8 1.0 1.2 1.4

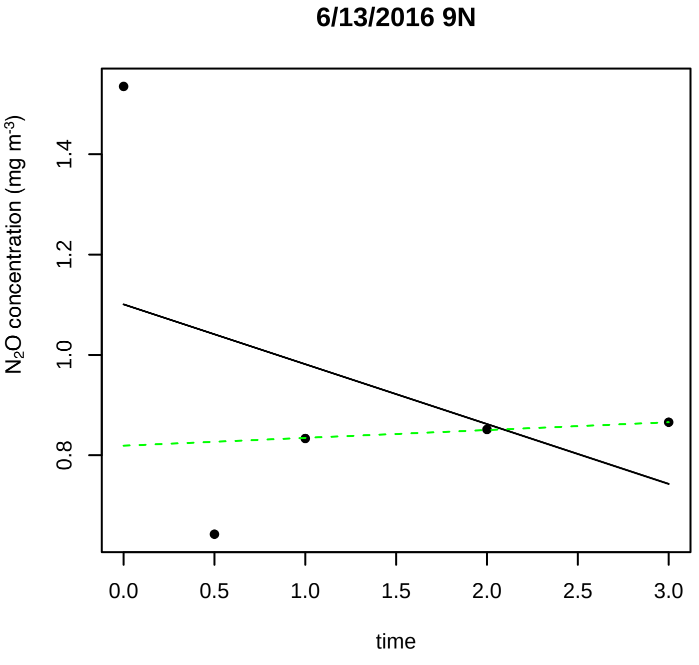
6/13/2016 9N
N2O concentration (mg m-3)
time
0.0 0.5 1.0 1.5 2.0 2.5 3.0
0.8 1.0 1.2 1.4

### 6/13/2016 9P

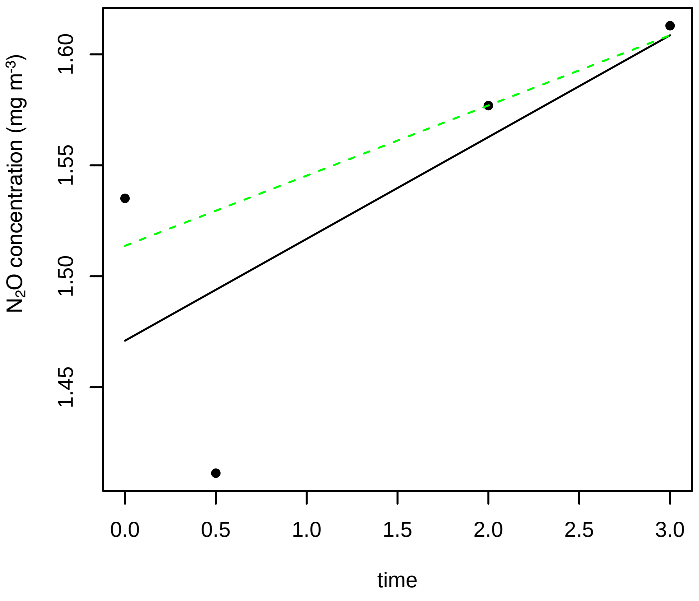


### 6/28/2016 10N

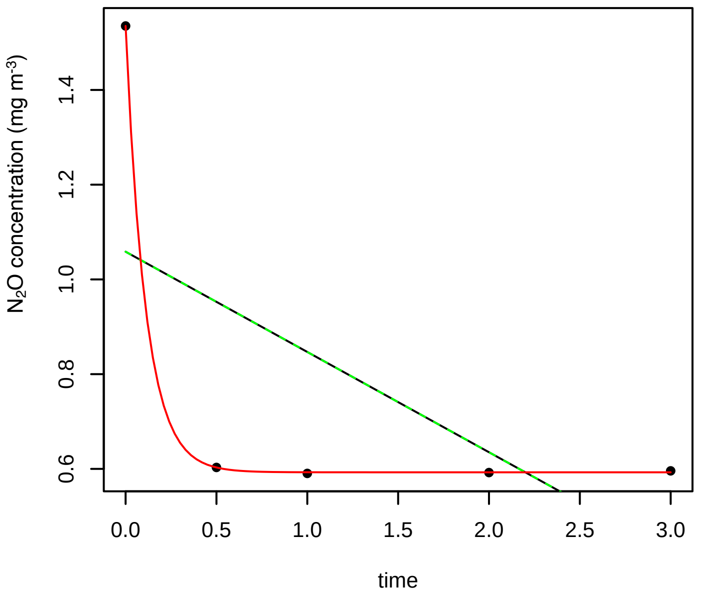


### 6/28/2016 10P

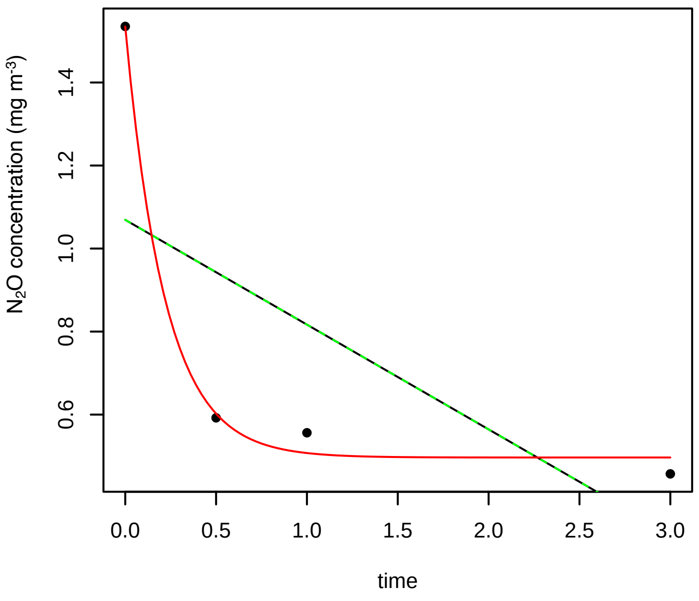


### 6/28/2016 11N

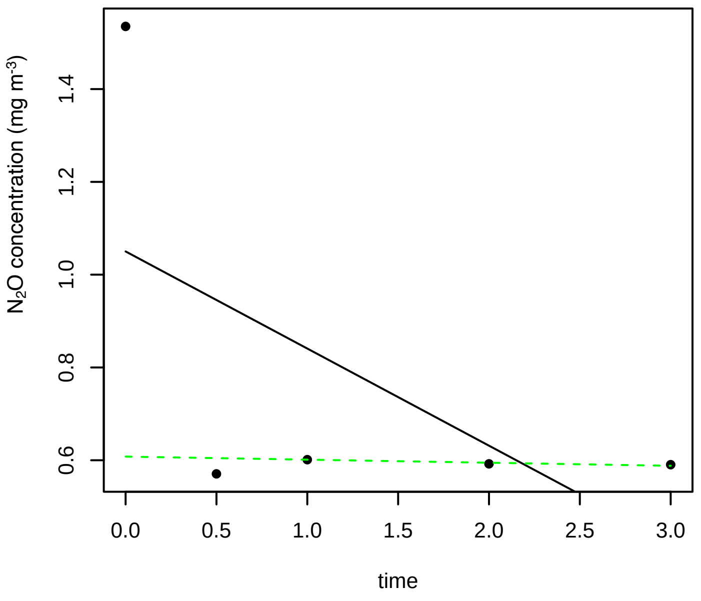


### 6/28/2016 11P

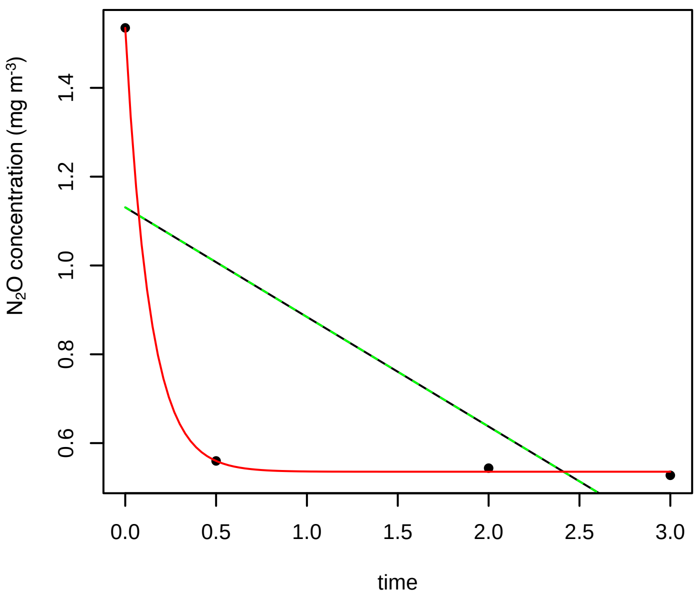


### 6/28/2016 12N

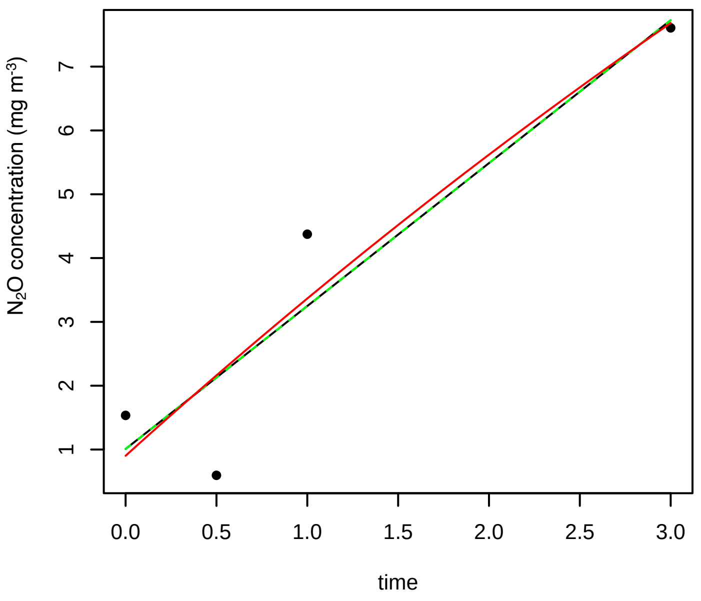

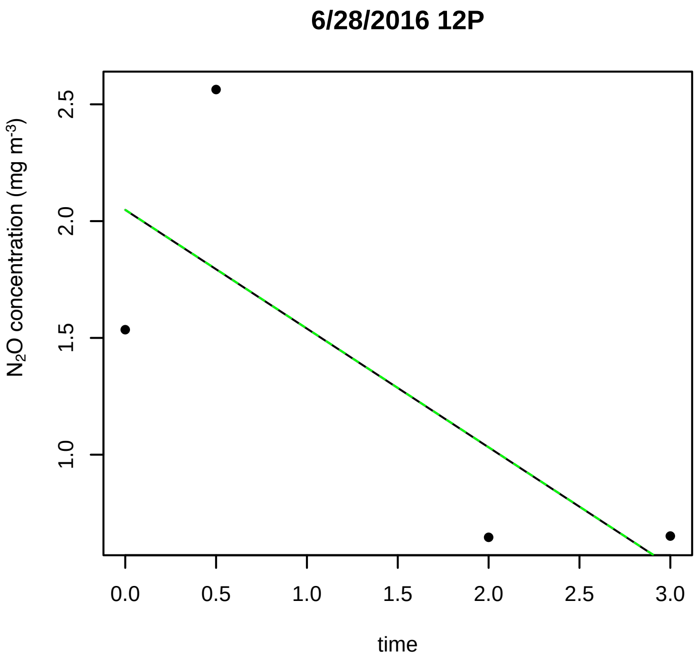
6/28/2016 12P
$N_2O$ concentration (mg $m^{-3}$)
time

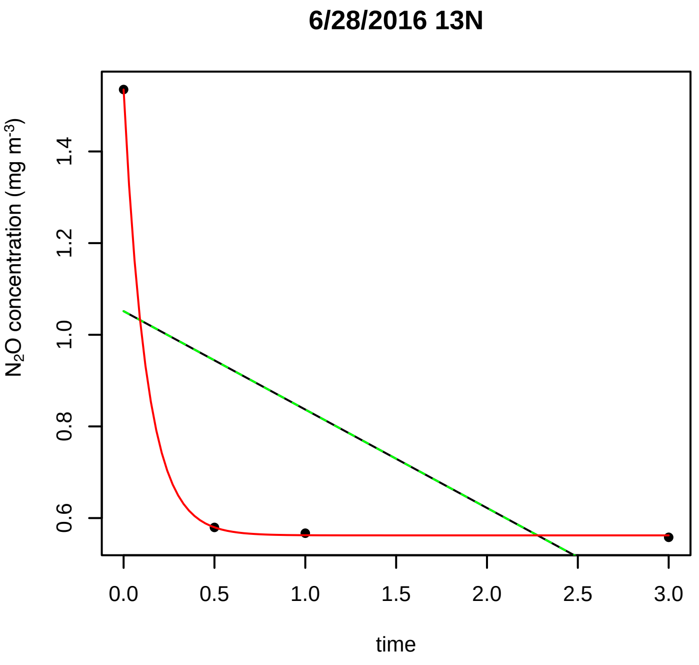
6/28/2016 13N
$N_2O$ concentration (mg $m^{-3}$)
time

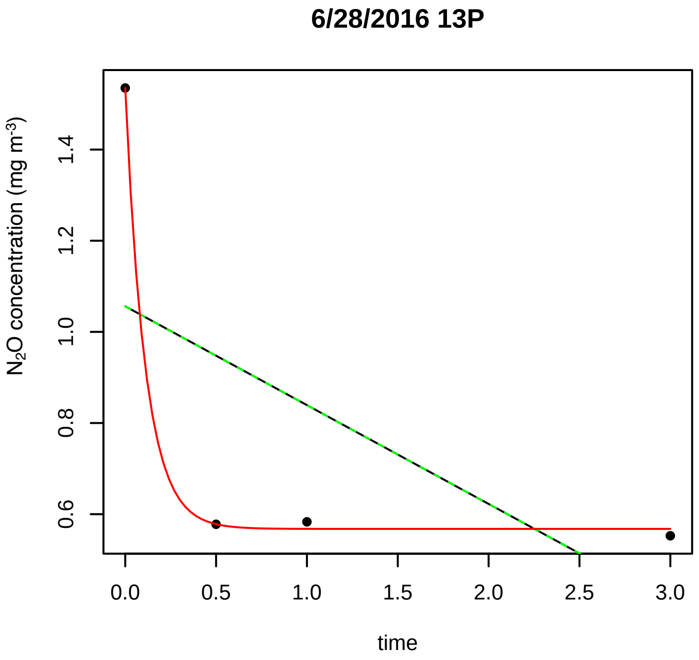
6/28/2016 13P
$N_2O$ concentration (mg $m^{-3}$)
time

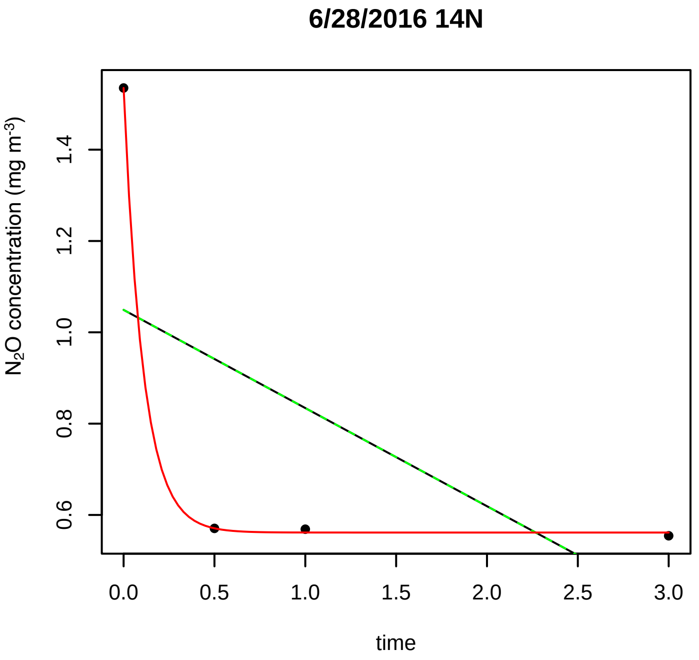
6/28/2016 14N
$N_2O$ concentration (mg $m^{-3}$)
time

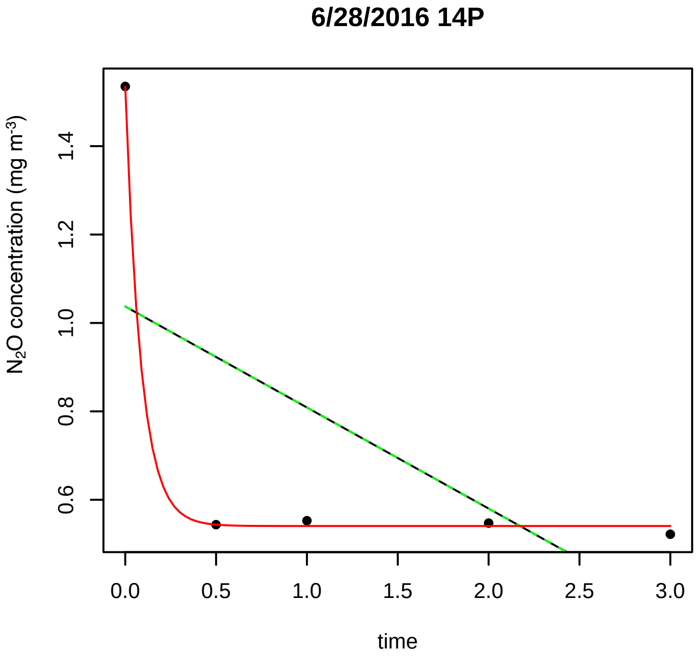
6/28/2016 14P
$N_2O$ concentration (mg $m^{-3}$)
time

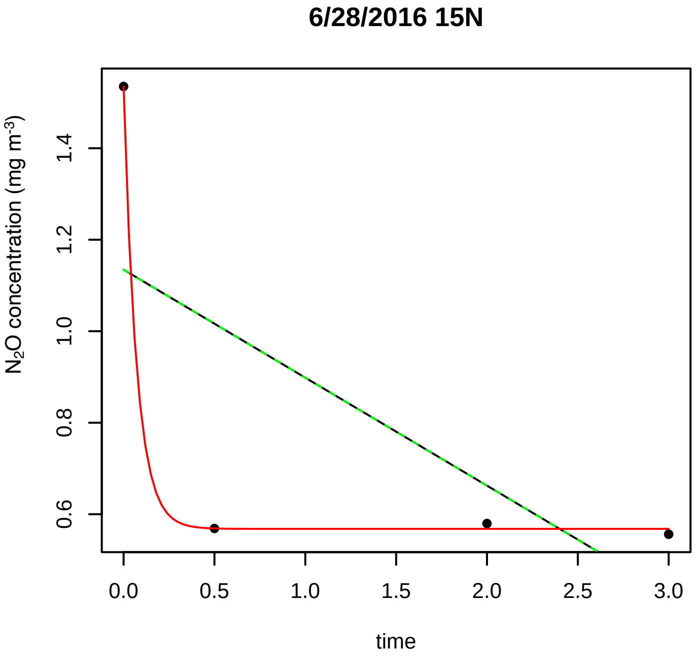
6/28/2016 15N
$N_2O$ concentration (mg $m^{-3}$)
time

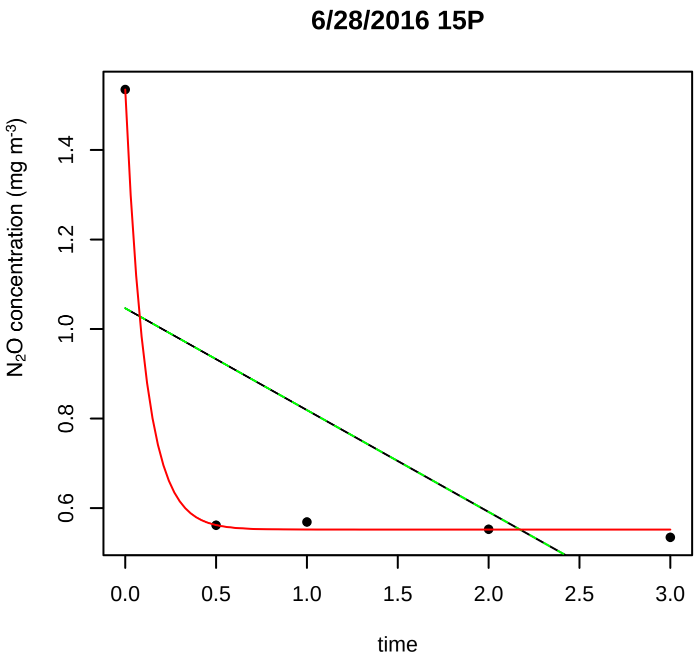
6/28/2016 15P
N2O concentration (mg m-3)
time

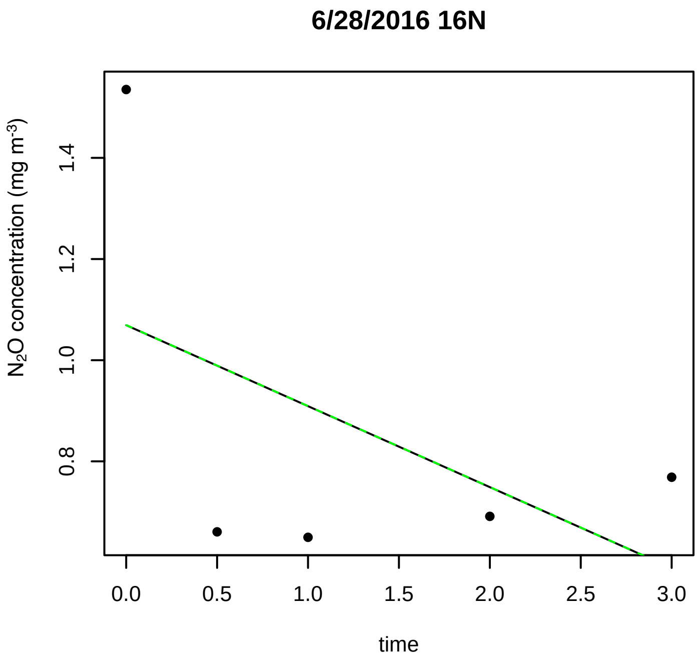
6/28/2016 16N
N2O concentration (mg m-3)
time

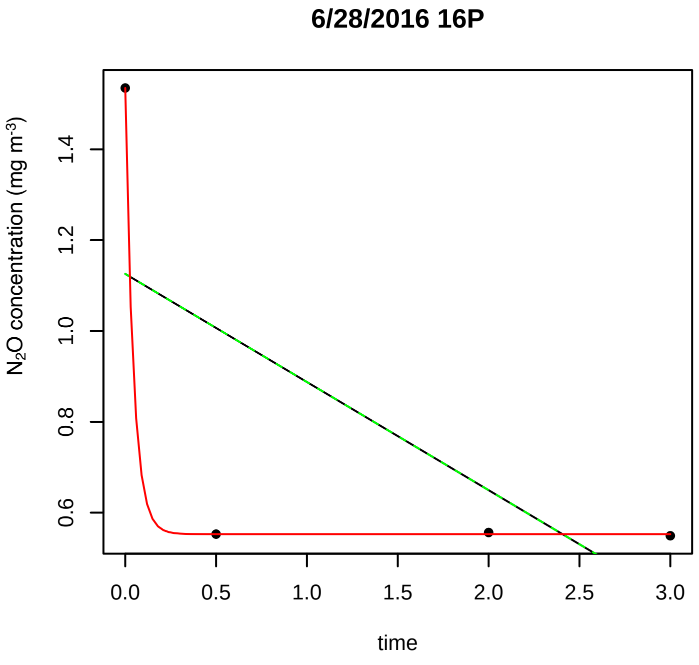
6/28/2016 16P
N2O concentration (mg m-3)
time

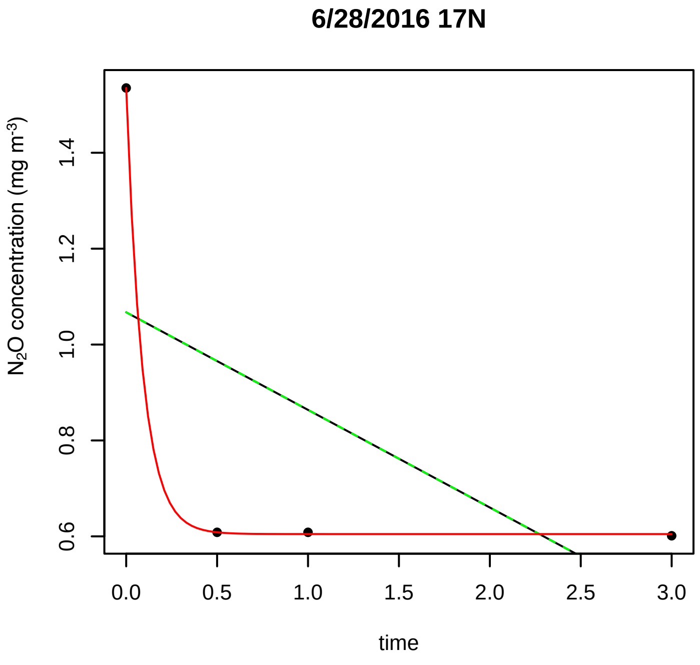
6/28/2016 17N
N2O concentration (mg m-3)
time

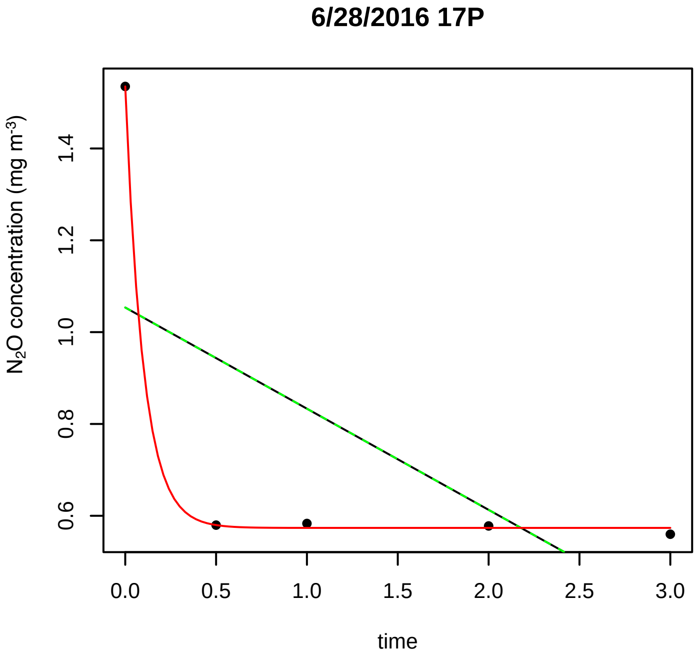
6/28/2016 17P
N2O concentration (mg m-3)
time

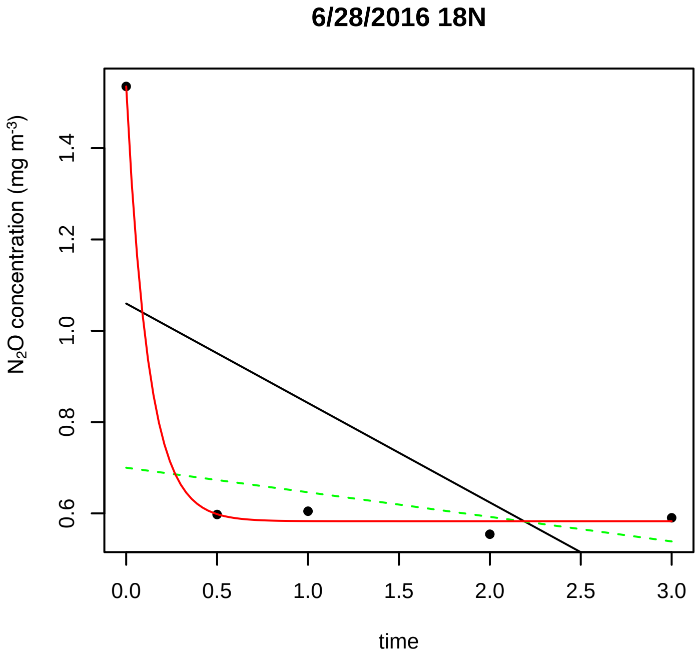
6/28/2016 18N
N2O concentration (mg m-3)
time

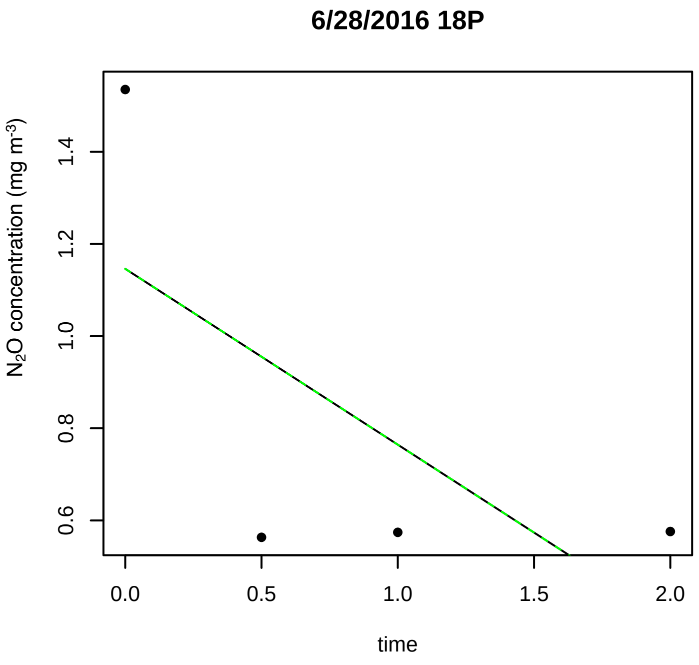
6/28/2016 18P
N2O concentration (mg m-3)
time

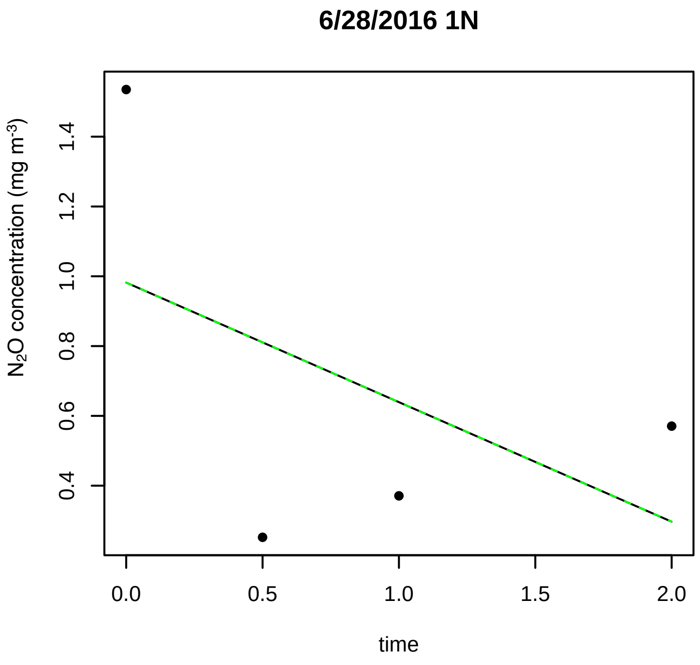
6/28/2016 1N
N2O concentration (mg m-3)
time

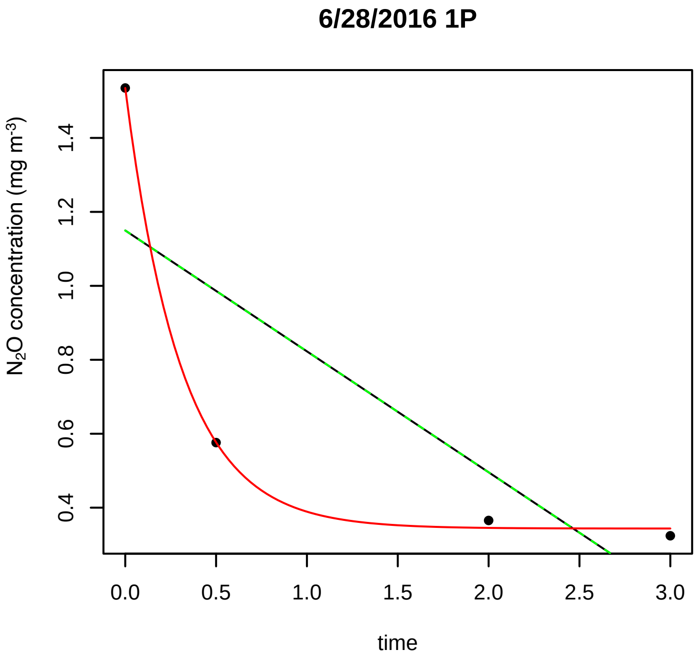
6/28/2016 1P
N2O concentration (mg m-3)
time

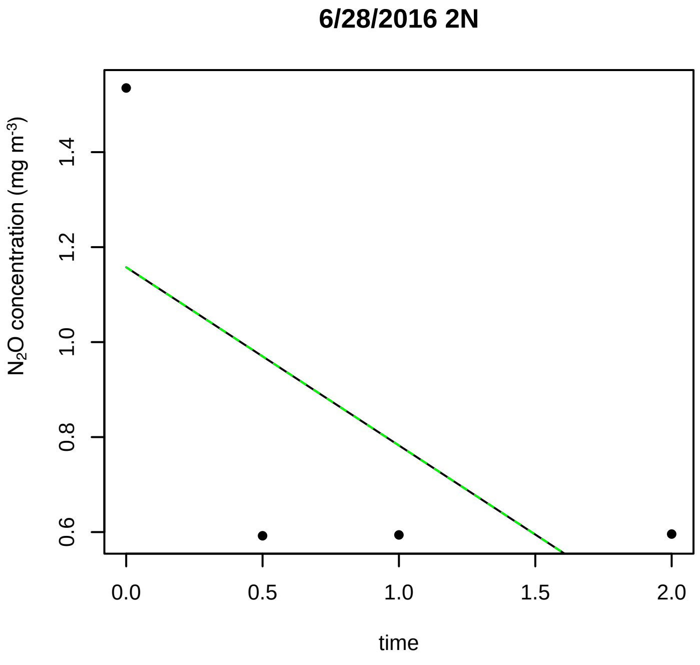
6/28/2016 2N
N2O concentration (mg m-3)
time

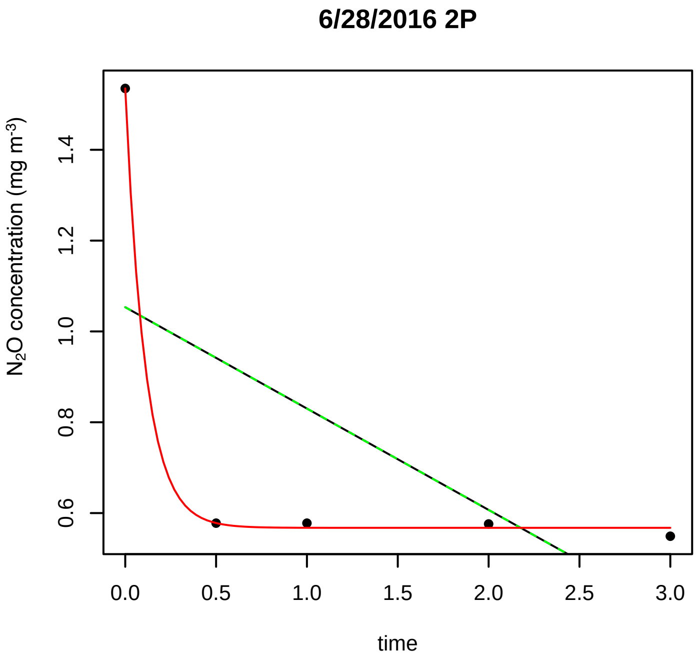
6/28/2016 2P
N2O concentration (mg m-3)
time

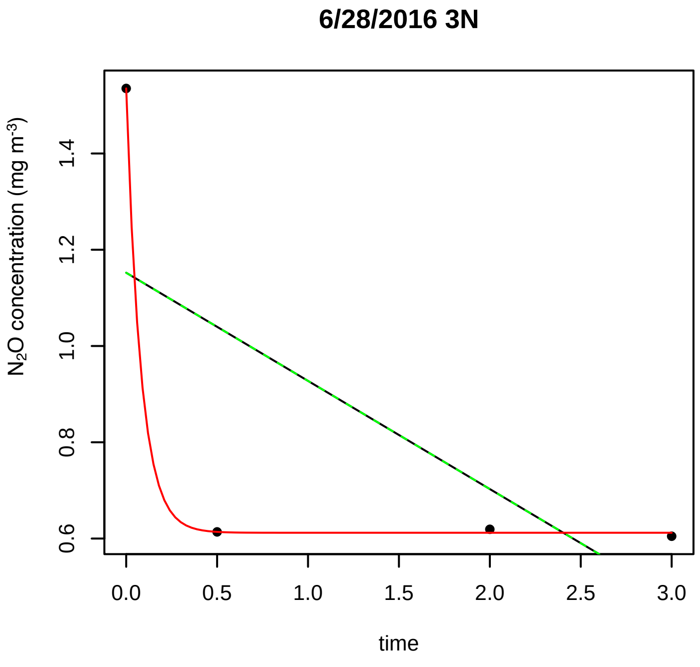
6/28/2016 3N
N2O concentration (mg m-3)
time

**6/28/2016 3P**

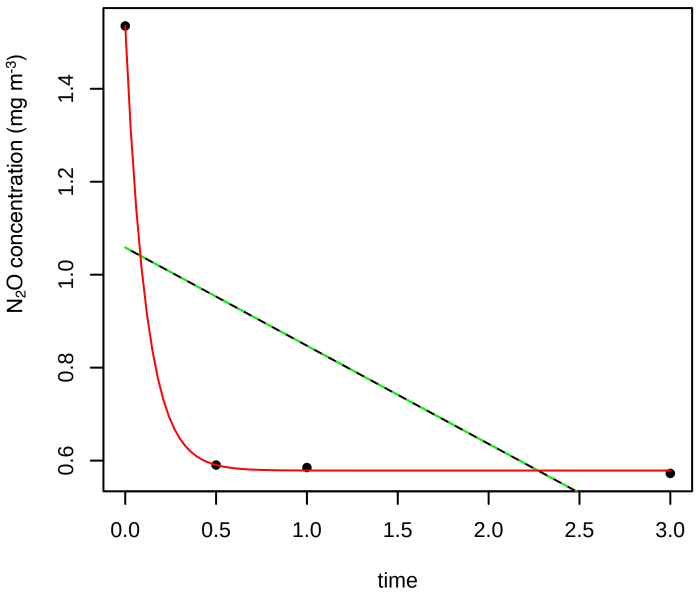


**6/28/2016 4N**

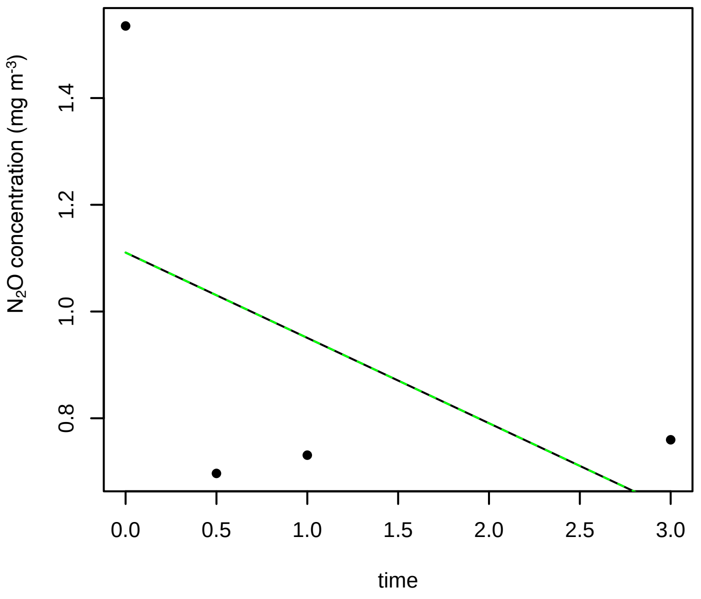


**6/28/2016 4P**

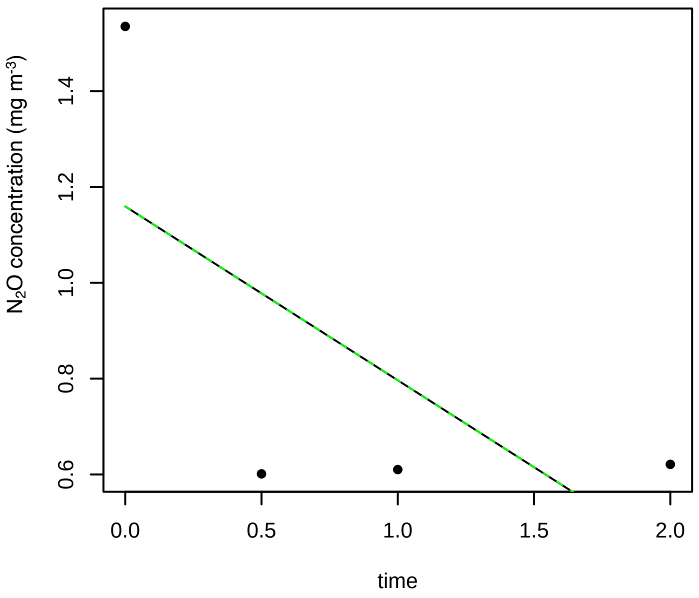


**6/28/2016 5N**

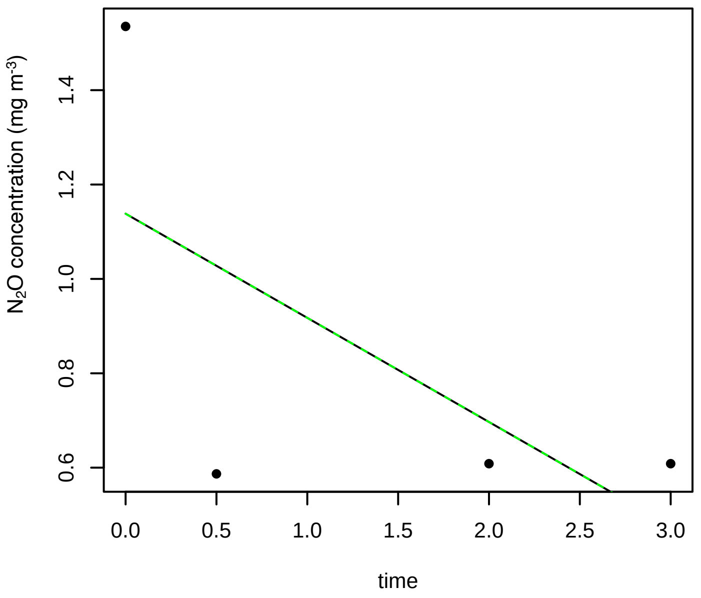


**6/28/2016 5P**

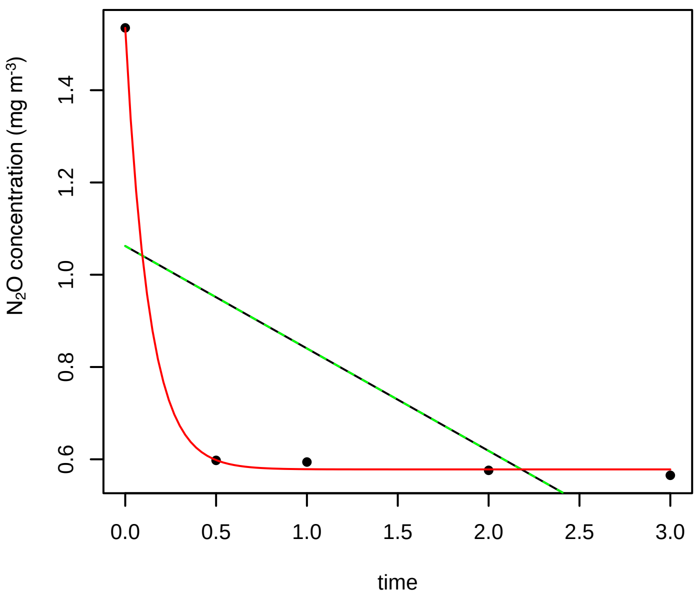


**6/28/2016 6N**

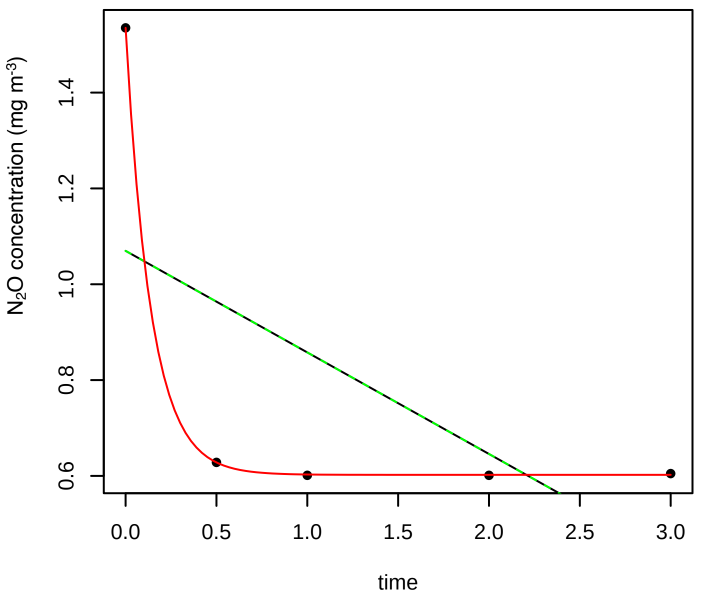

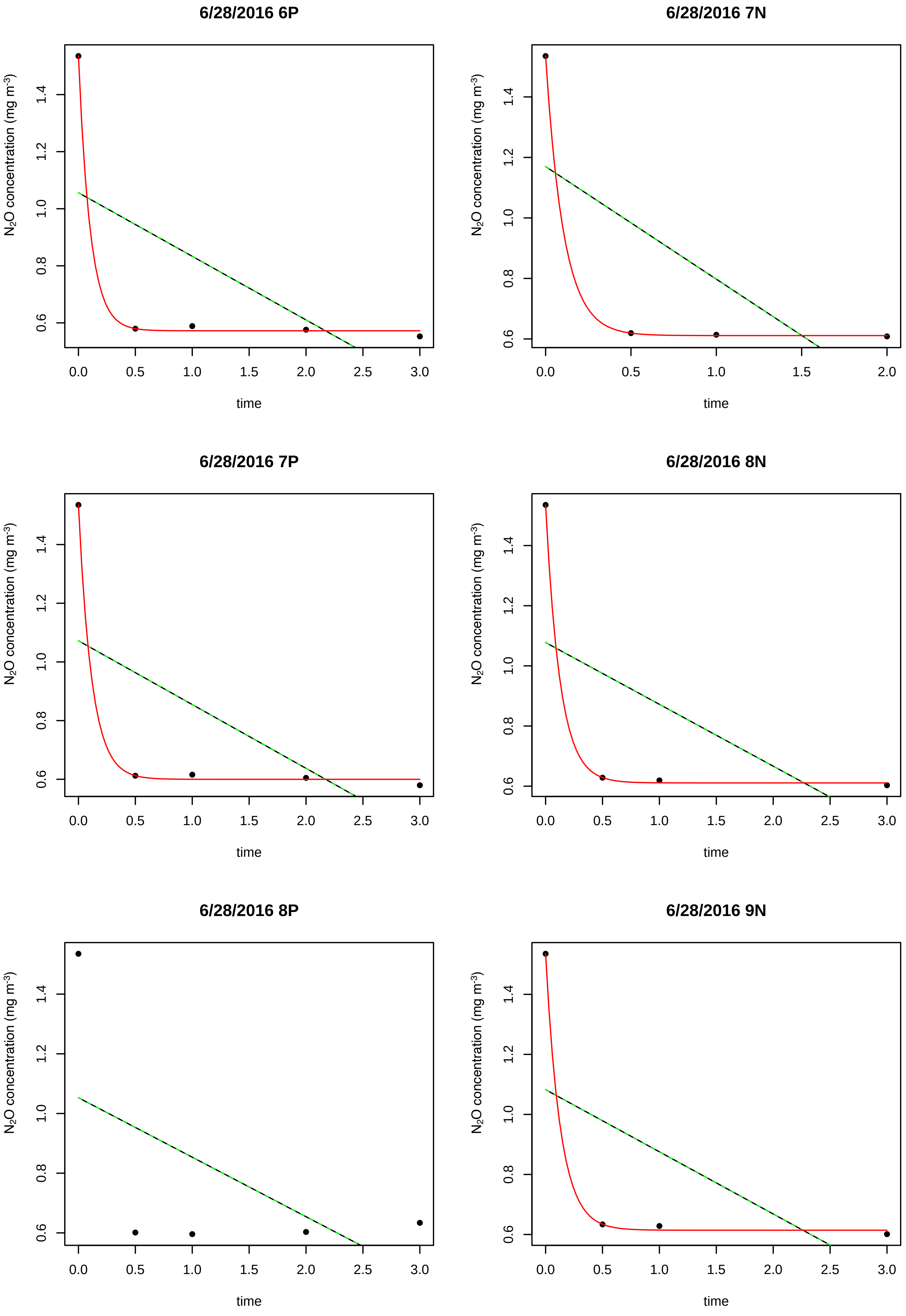
6/28/2016 6P
6/28/2016 7N
6/28/2016 7P
6/28/2016 8N
6/28/2016 8P
6/28/2016 9N
N2O concentration (mg m-3)
time
0.0
0.5
1.0
1.5
2.0
2.5
3.0
0.6
0.8
1.0
1.2
1.4

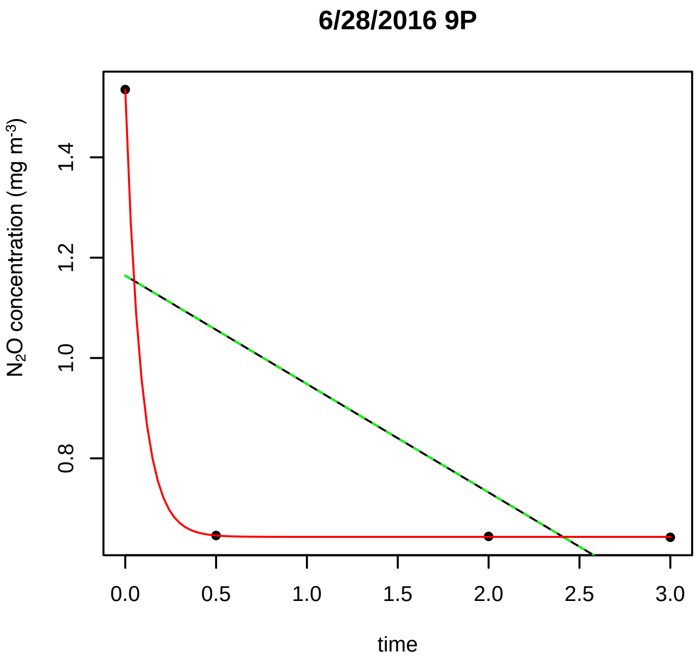
6/28/2016 9P
N$_2$O concentration (mg m$^{-3}$)
time

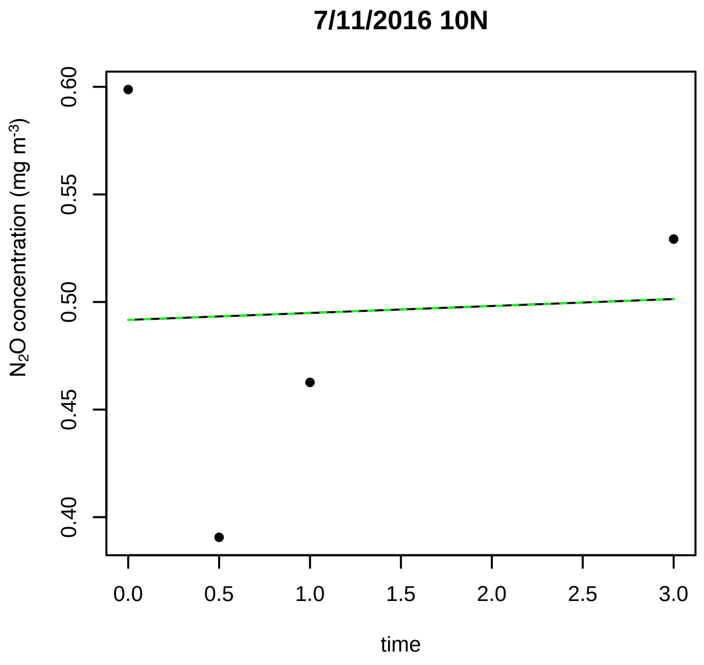
7/11/2016 10N
N$_2$O concentration (mg m$^{-3}$)
time

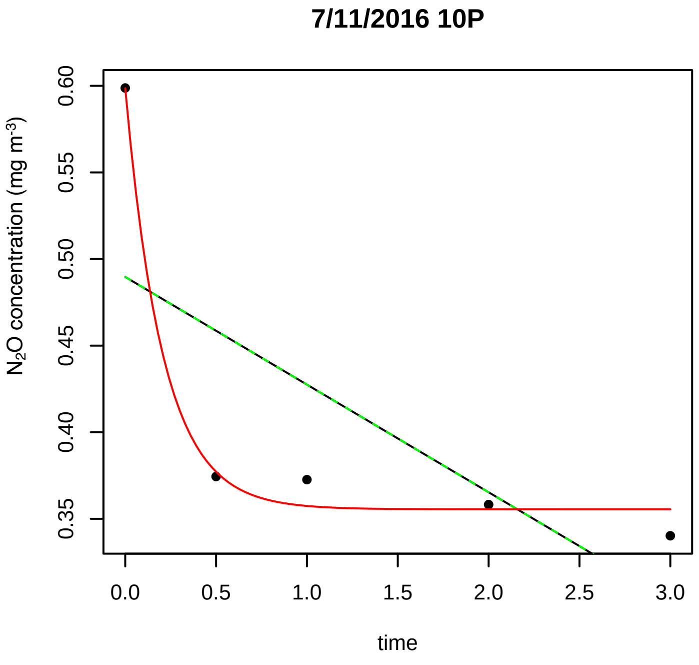
7/11/2016 10P
N$_2$O concentration (mg m$^{-3}$)
time

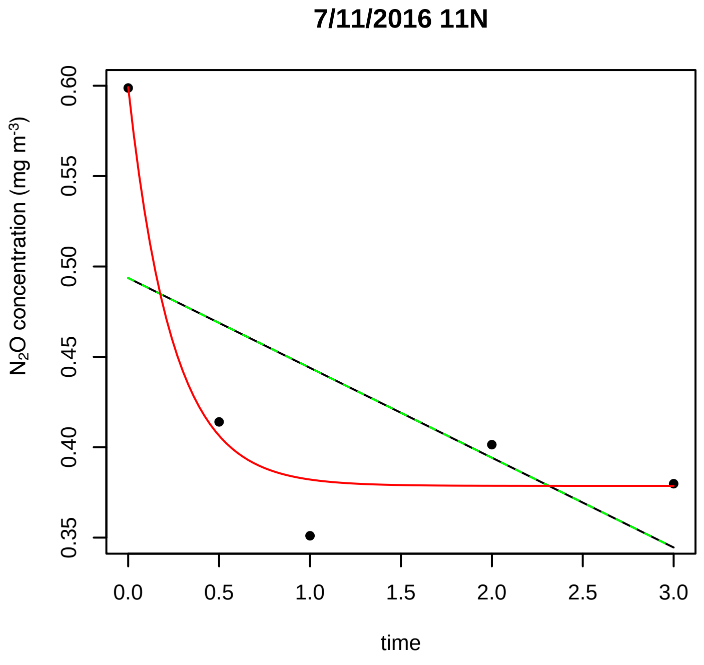
7/11/2016 11N
N$_2$O concentration (mg m$^{-3}$)
time

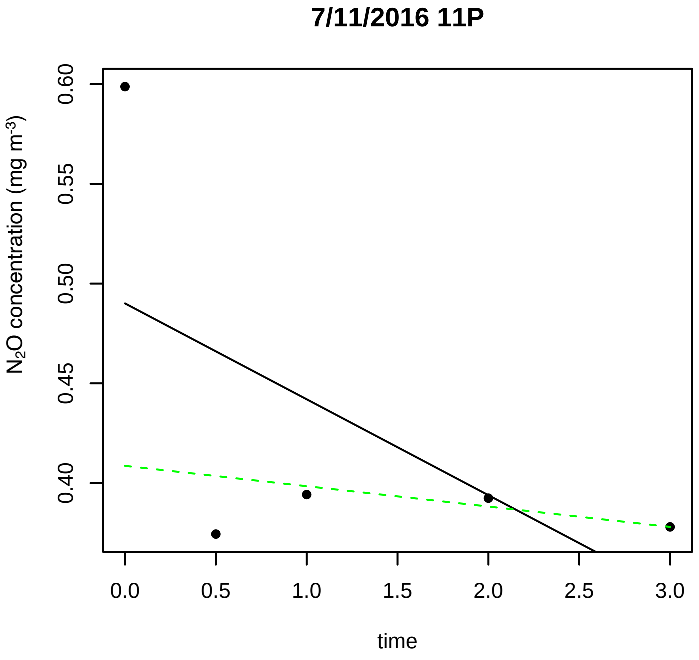
7/11/2016 11P
N$_2$O concentration (mg m$^{-3}$)
time

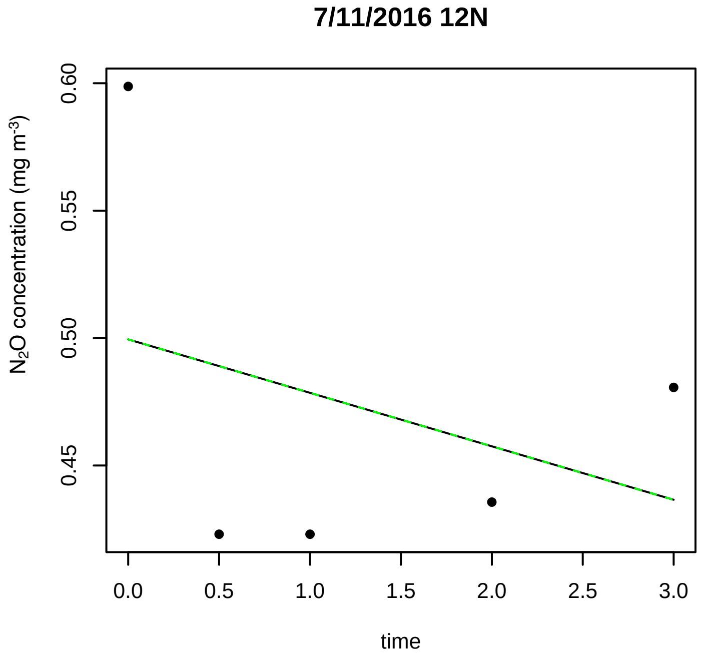
7/11/2016 12N
N$_2$O concentration (mg m$^{-3}$)
time

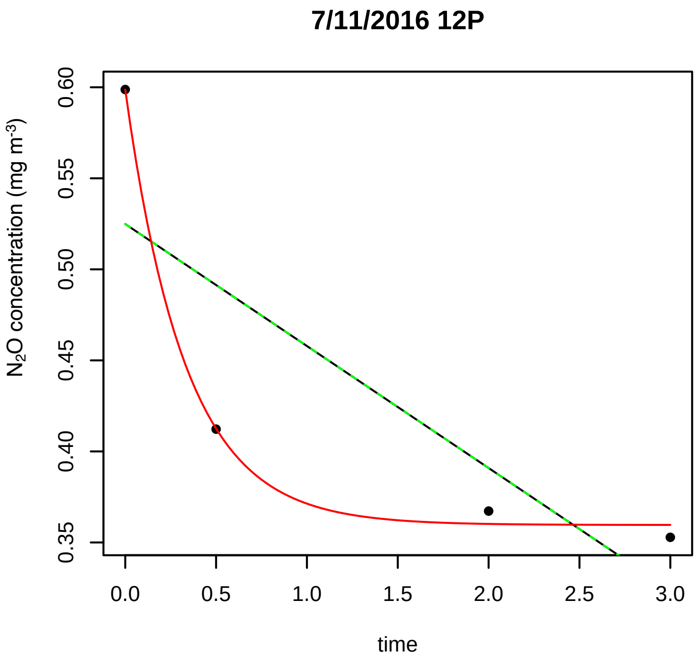
7/11/2016 12P
N2O concentration (mg m-3)
0.35
0.40
0.45
0.50
0.55
0.60
0.0
0.5
1.0
1.5
2.0
2.5
3.0
time

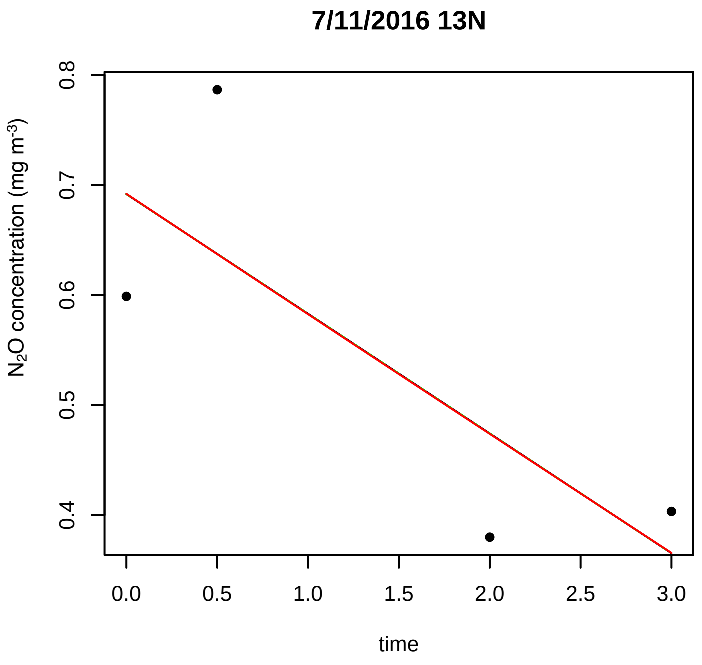
7/11/2016 13N
N2O concentration (mg m-3)
0.4
0.5
0.6
0.7
0.8
0.0
0.5
1.0
1.5
2.0
2.5
3.0
time

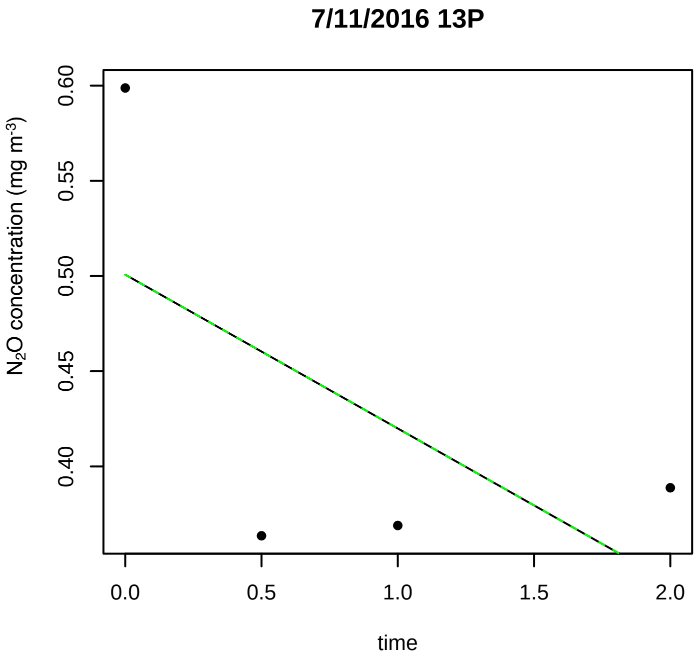
7/11/2016 13P
N2O concentration (mg m-3)
0.40
0.45
0.50
0.55
0.60
0.0
0.5
1.0
1.5
2.0
time

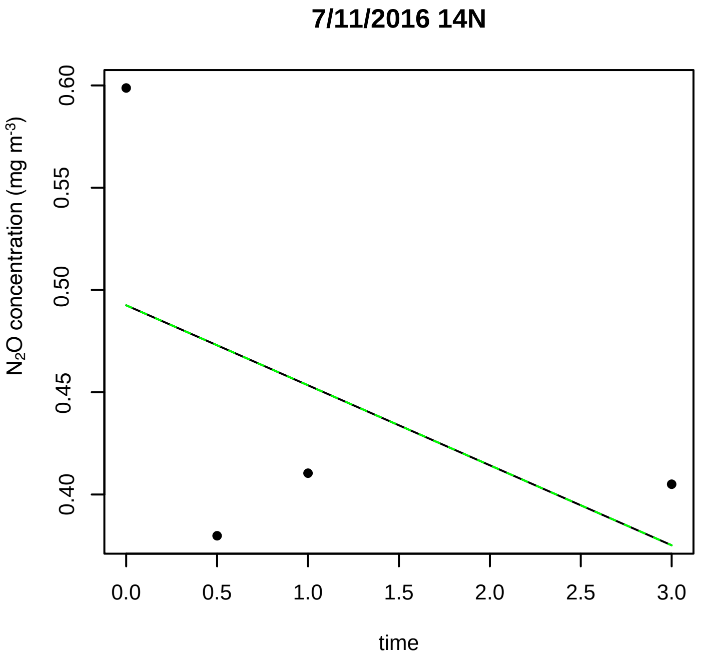
7/11/2016 14N
N2O concentration (mg m-3)
0.40
0.45
0.50
0.55
0.60
0.0
0.5
1.0
1.5
2.0
2.5
3.0
time

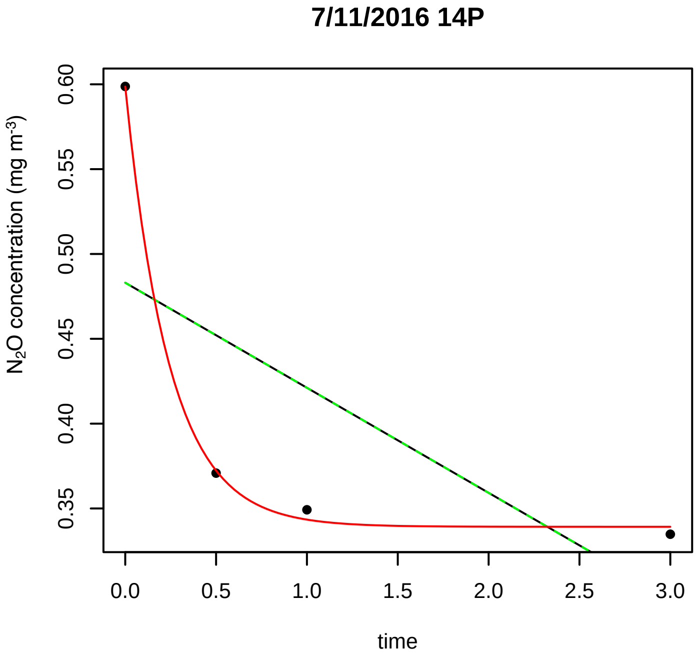
7/11/2016 14P
N2O concentration (mg m-3)
0.35
0.40
0.45
0.50
0.55
0.60
0.0
0.5
1.0
1.5
2.0
2.5
3.0
time

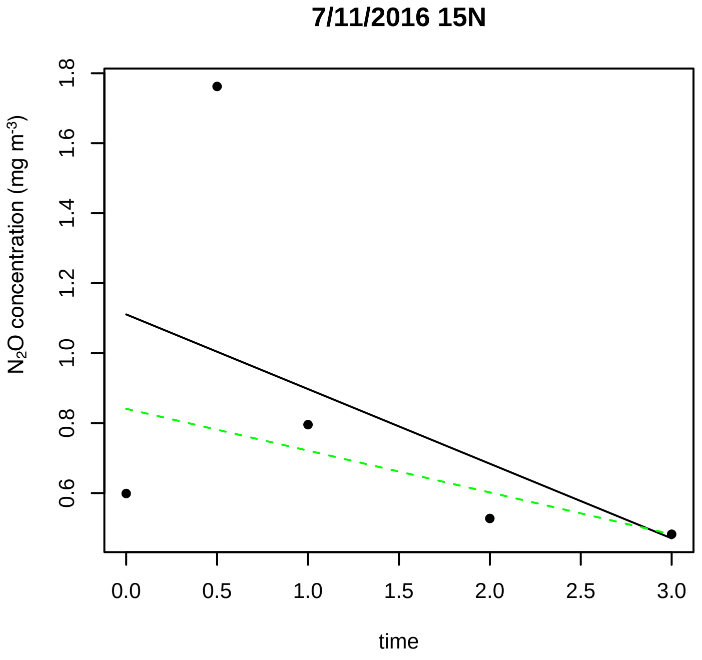
7/11/2016 15N
N2O concentration (mg m-3)
0.6
0.8
1.0
1.2
1.4
1.6
1.8
0.0
0.5
1.0
1.5
2.0
2.5
3.0
time

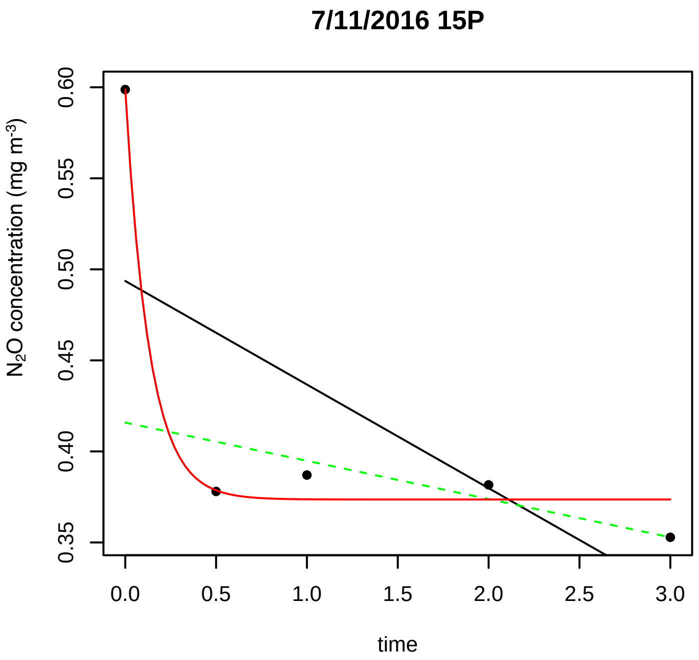
7/11/2016 15P
N2O concentration (mg m-3)
0.35
0.40
0.45
0.50
0.55
0.60
0.0
0.5
1.0
1.5
2.0
2.5
3.0
time

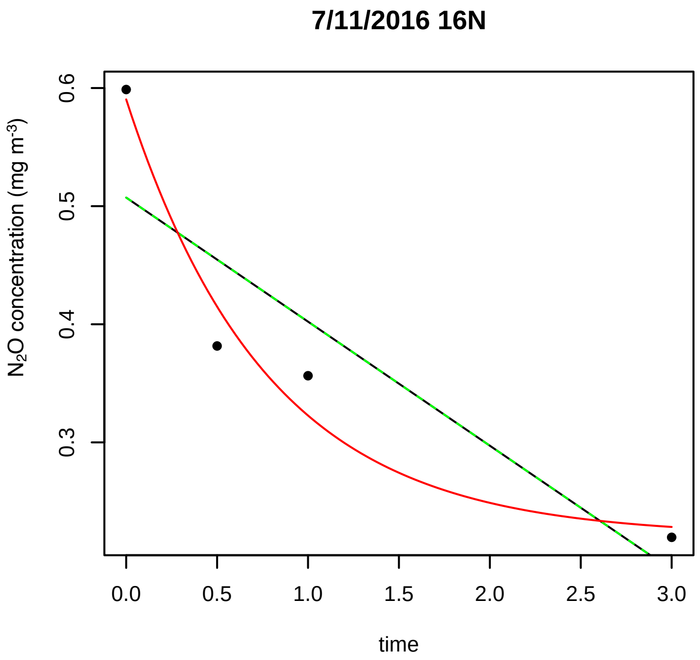
7/11/2016 16N
N2O concentration (mg m-3)
0.3
0.4
0.5
0.6
0.0
0.5
1.0
1.5
2.0
2.5
3.0
time

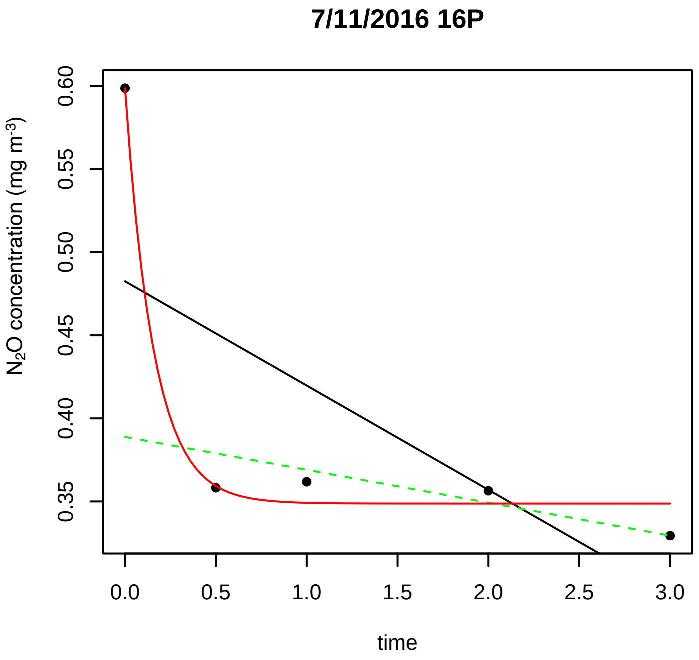
7/11/2016 16P
N2O concentration (mg m-3)
0.35
0.40
0.45
0.50
0.55
0.60
0.0
0.5
1.0
1.5
2.0
2.5
3.0
time

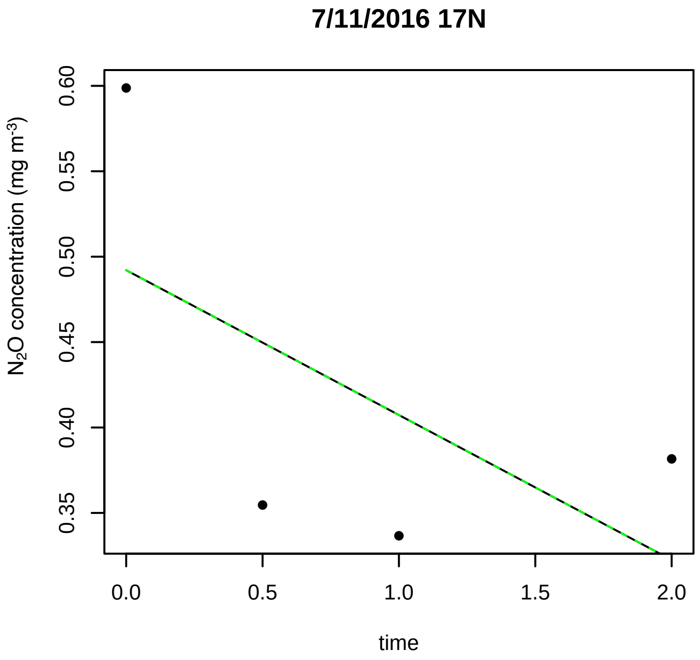
7/11/2016 17N
N2O concentration (mg m-3)
0.35
0.40
0.45
0.50
0.55
0.60
0.0
0.5
1.0
1.5
2.0
time

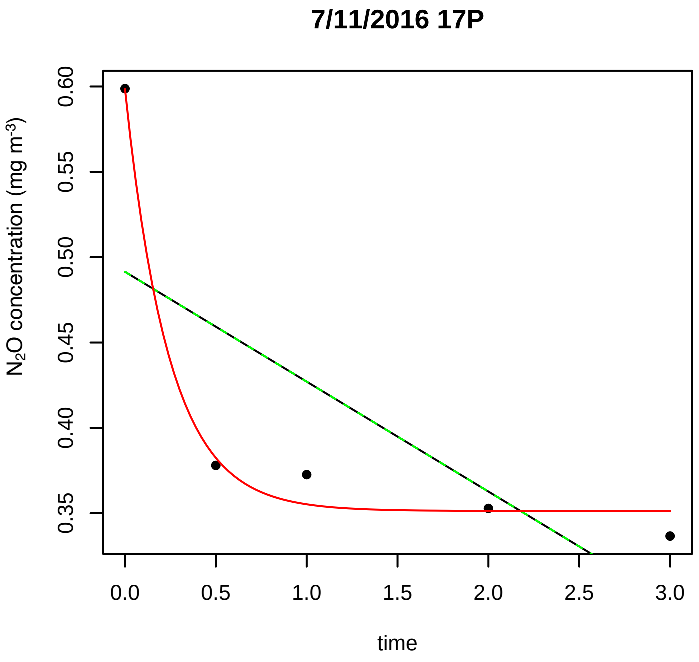
7/11/2016 17P
N2O concentration (mg m-3)
0.35
0.40
0.45
0.50
0.55
0.60
0.0
0.5
1.0
1.5
2.0
2.5
3.0
time

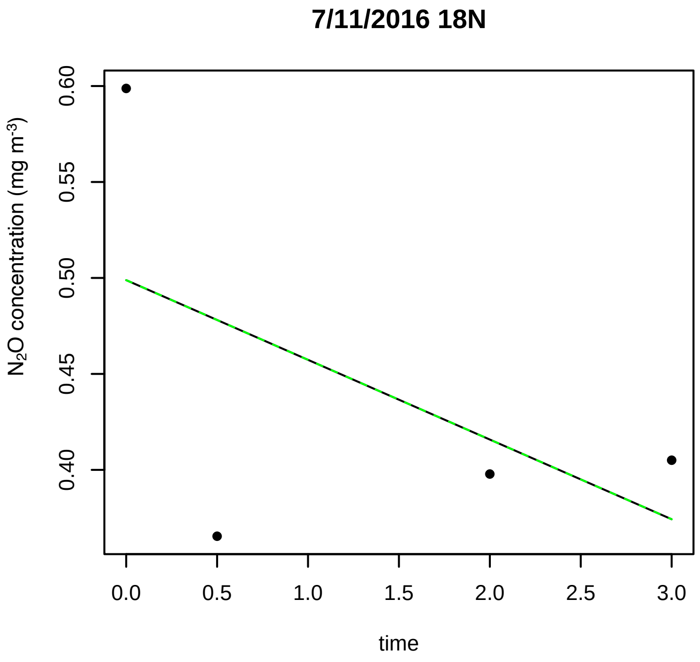
7/11/2016 18N
N2O concentration (mg m-3)
0.40
0.45
0.50
0.55
0.60
0.0
0.5
1.0
1.5
2.0
2.5
3.0
time

**7/11/2016 18P**

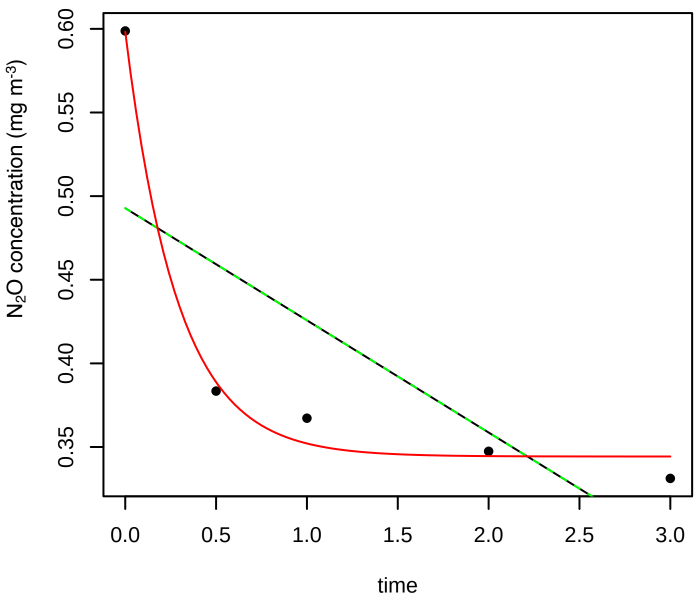


**7/11/2016 1N**

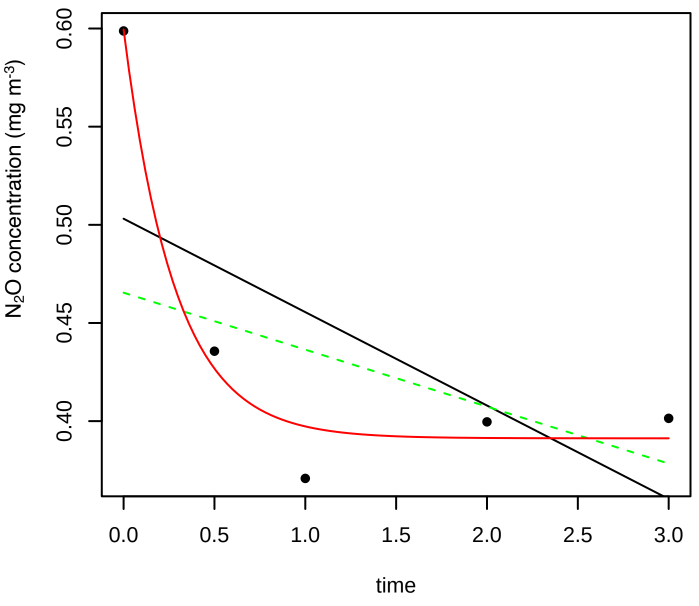


**7/11/2016 1P**

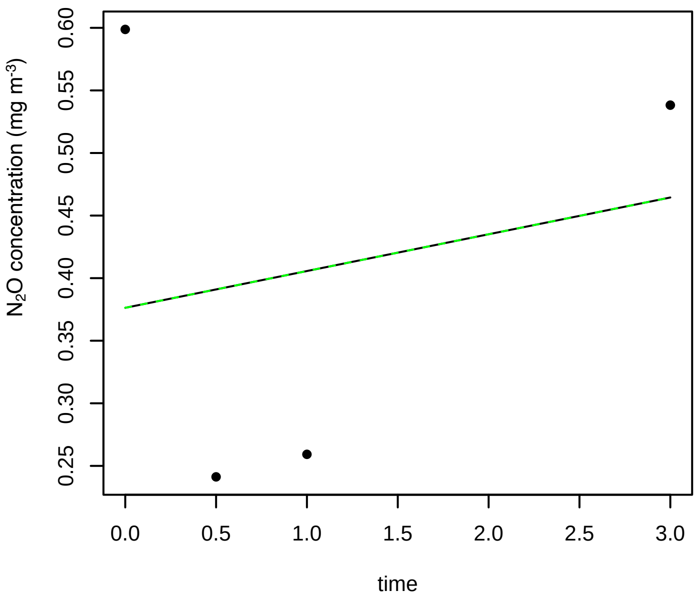


**7/11/2016 2N**

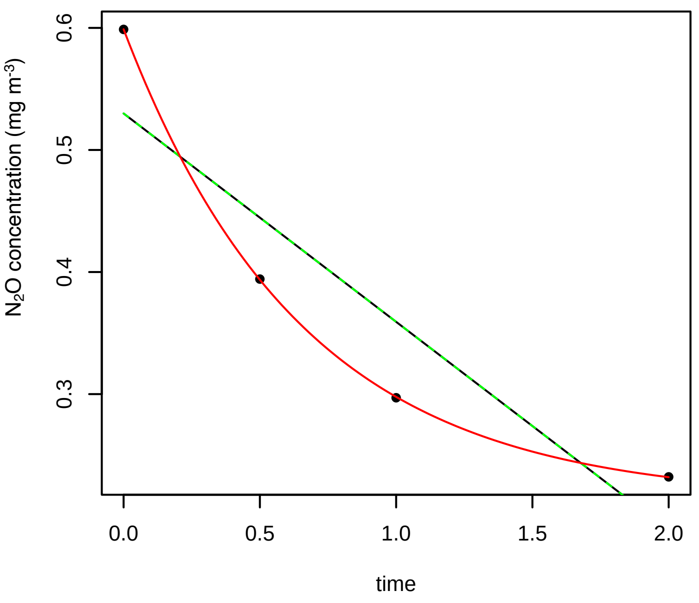


**7/11/2016 2P**

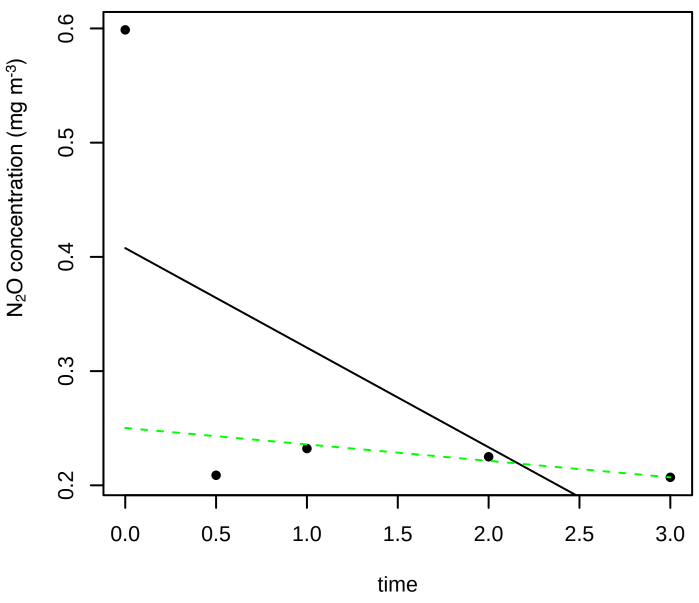


**7/11/2016 3N**

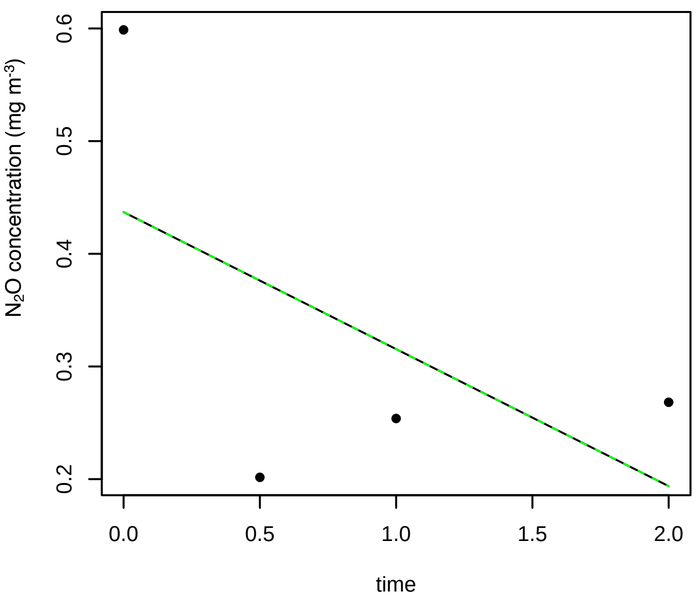

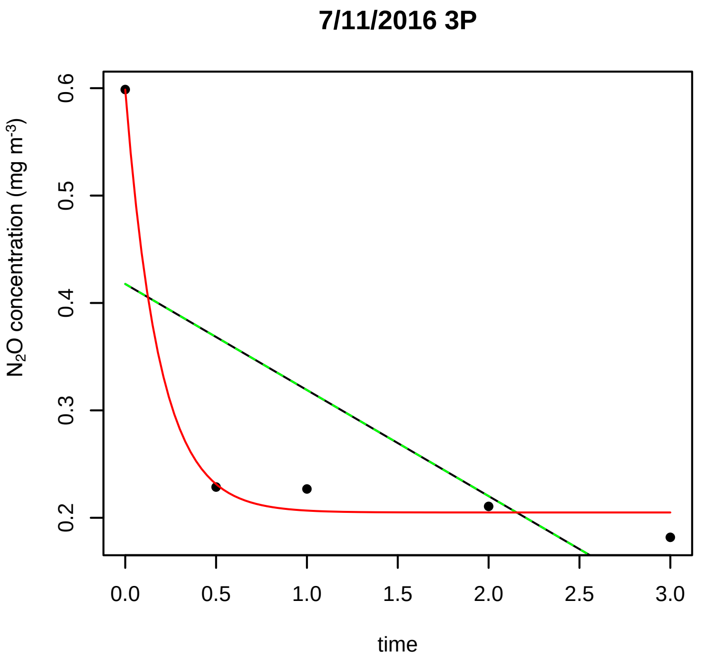
7/11/2016 3P
N2O concentration (mg m-3)
time

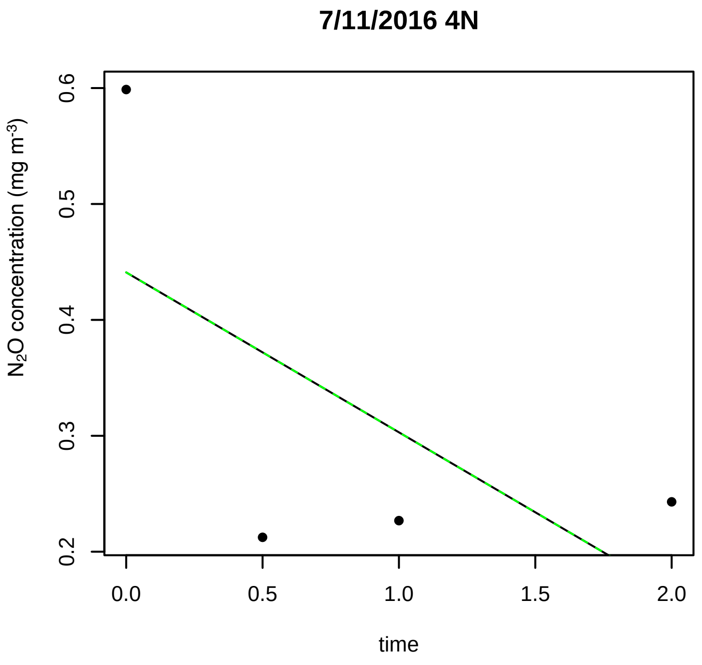
7/11/2016 4N
N2O concentration (mg m-3)
time

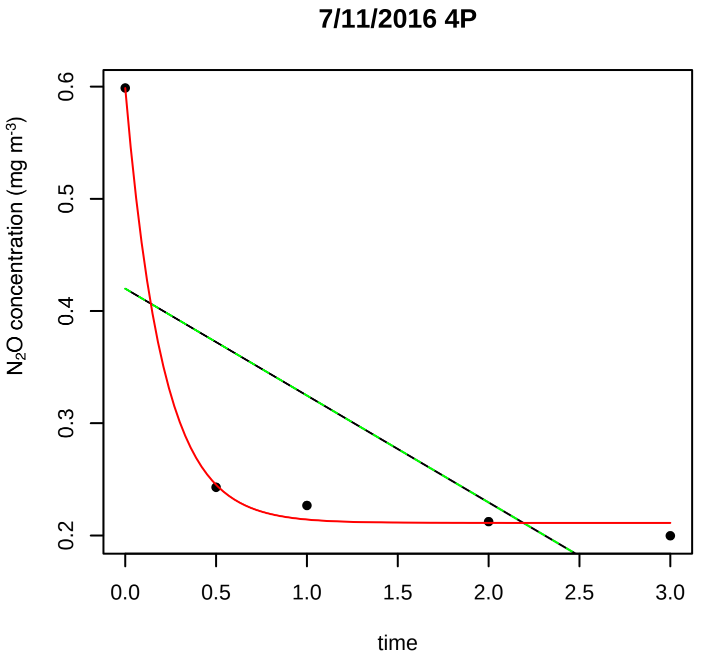
7/11/2016 4P
N2O concentration (mg m-3)
time

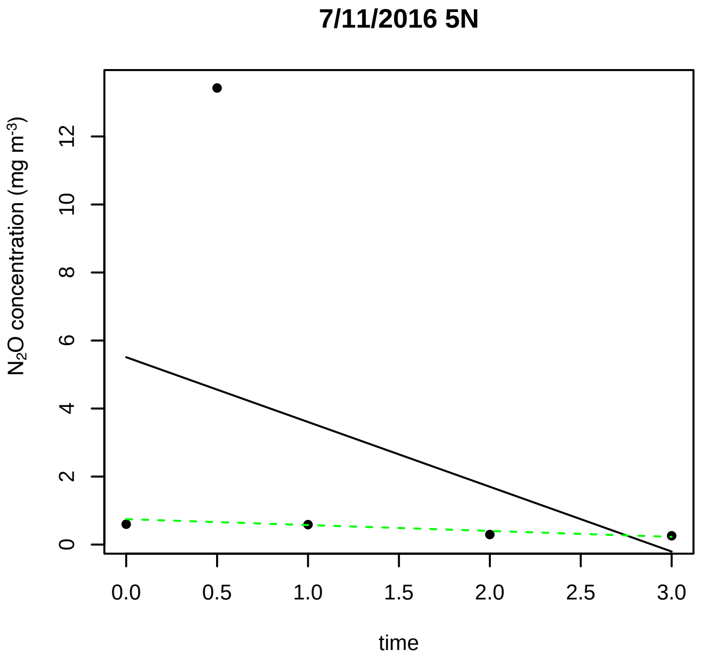
7/11/2016 5N
N2O concentration (mg m-3)
time

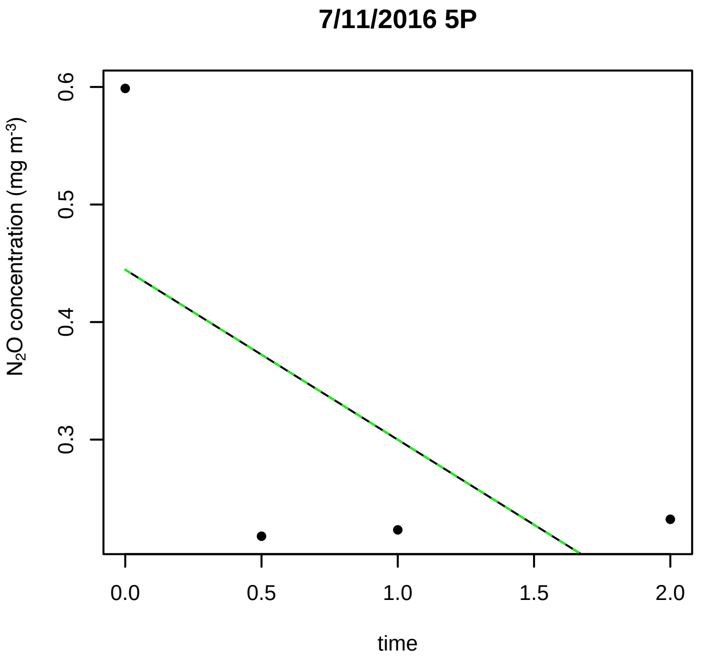
7/11/2016 5P
N2O concentration (mg m-3)
time

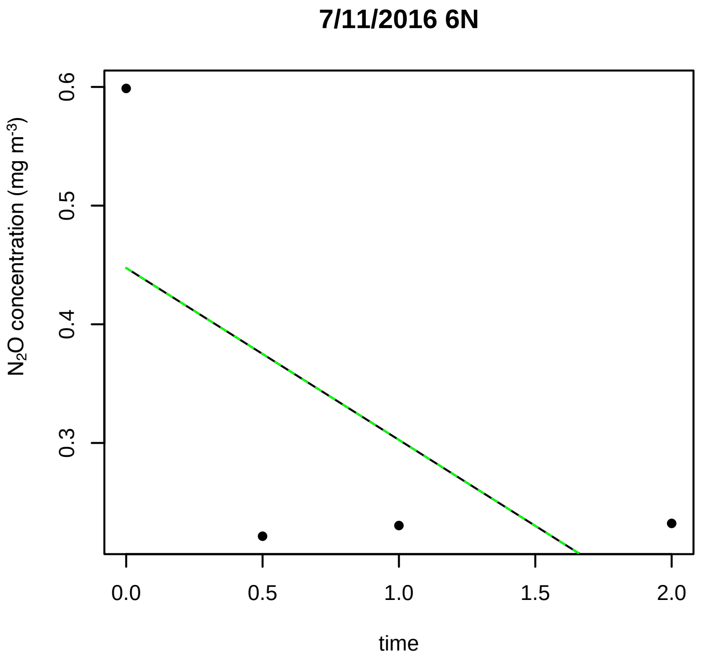
7/11/2016 6N
N2O concentration (mg m-3)
time

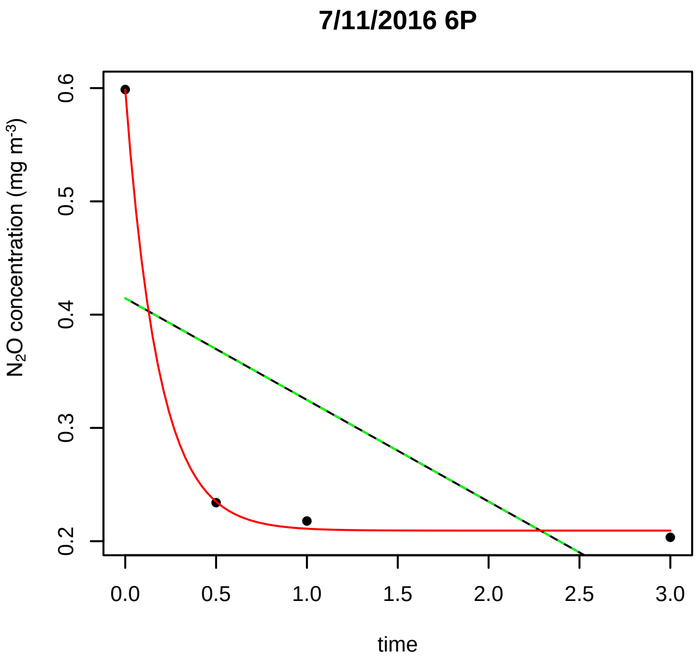
7/11/2016 6P
N2O concentration (mg m-3)
0.2
0.3
0.4
0.5
0.6
0.0
0.5
1.0
1.5
2.0
2.5
3.0
time

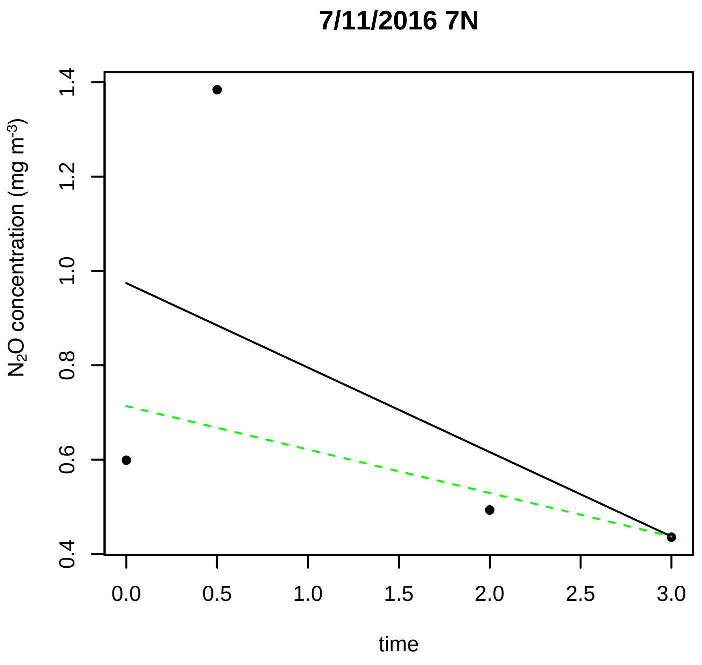
7/11/2016 7N
N2O concentration (mg m-3)
0.4
0.6
0.8
1.0
1.2
1.4
0.0
0.5
1.0
1.5
2.0
2.5
3.0
time

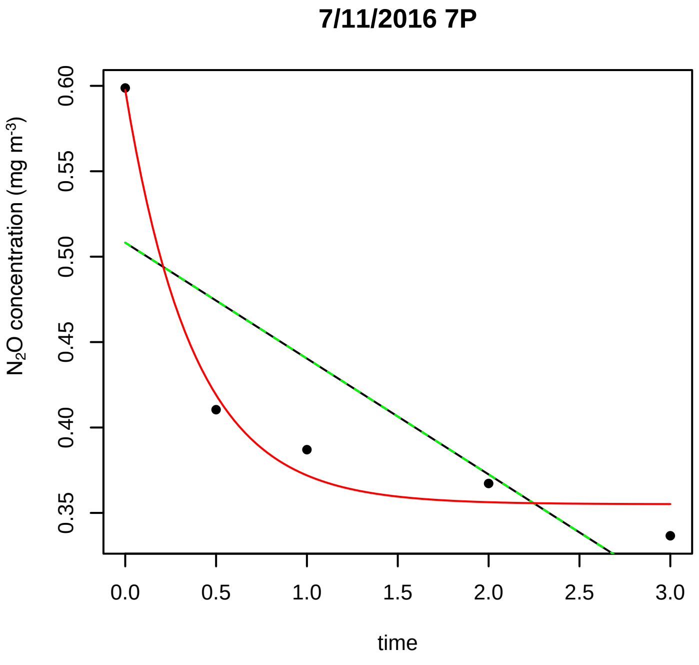
7/11/2016 7P
N2O concentration (mg m-3)
0.35
0.40
0.45
0.50
0.55
0.60
0.0
0.5
1.0
1.5
2.0
2.5
3.0
time

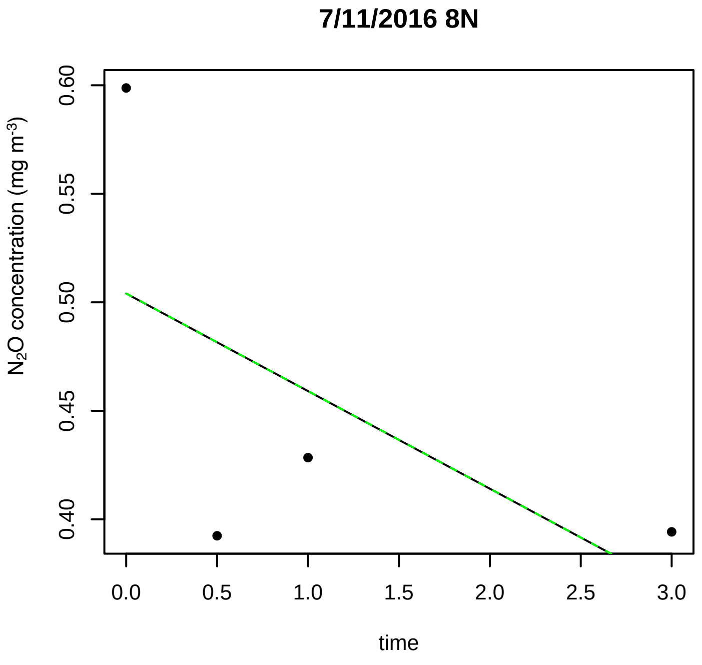
7/11/2016 8N
N2O concentration (mg m-3)
0.40
0.45
0.50
0.55
0.60
0.0
0.5
1.0
1.5
2.0
2.5
3.0
time

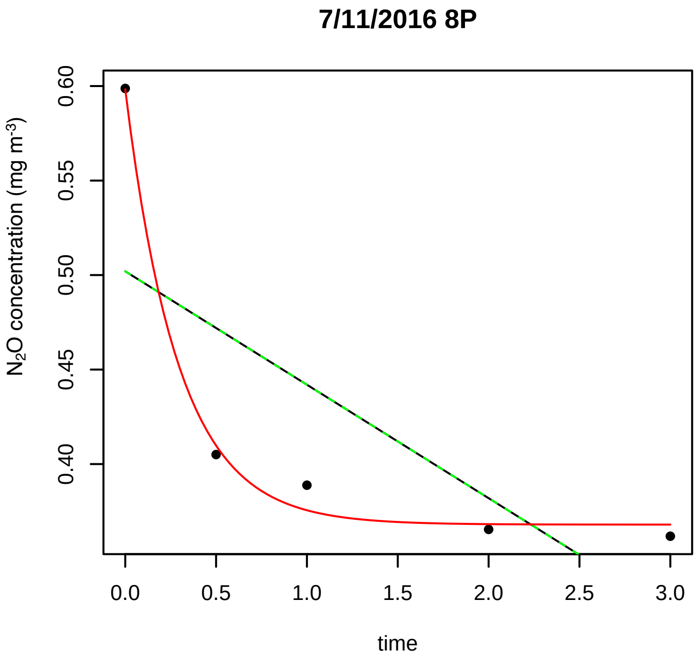
7/11/2016 8P
N2O concentration (mg m-3)
0.40
0.45
0.50
0.55
0.60
0.0
0.5
1.0
1.5
2.0
2.5
3.0
time

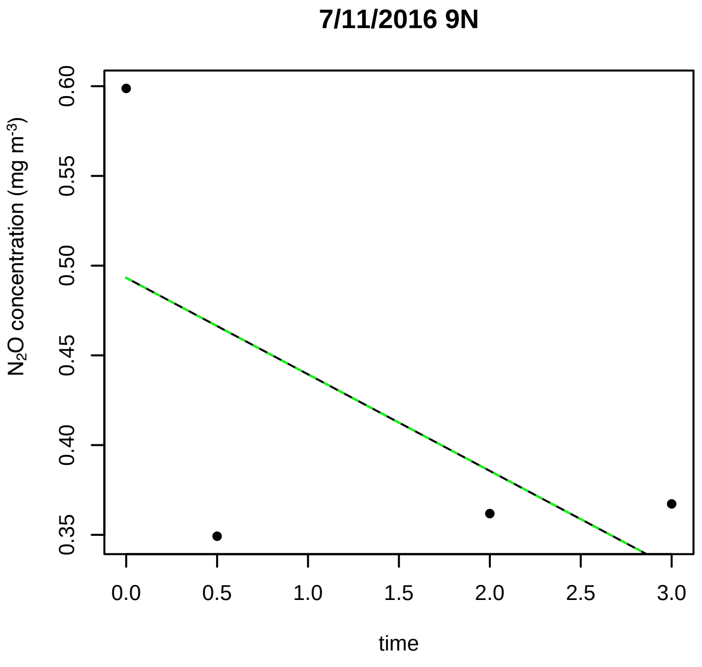
7/11/2016 9N
N2O concentration (mg m-3)
0.35
0.40
0.45
0.50
0.55
0.60
0.0
0.5
1.0
1.5
2.0
2.5
3.0
time

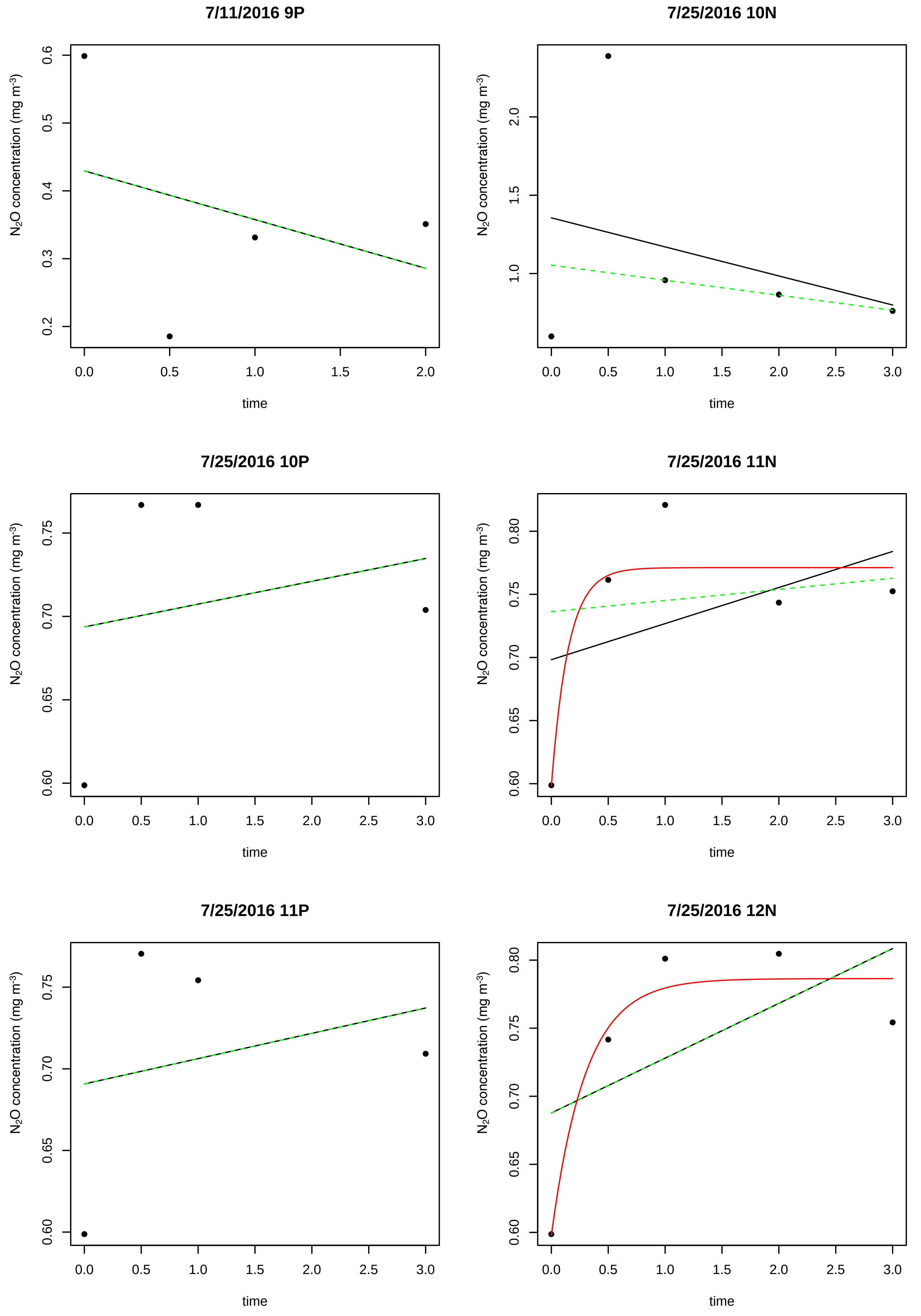
7/11/2016 9P
7/25/2016 10N
7/25/2016 10P
7/25/2016 11N
7/25/2016 11P
7/25/2016 12N
$N_2O$ concentration (mg $m^{-3}$)
time

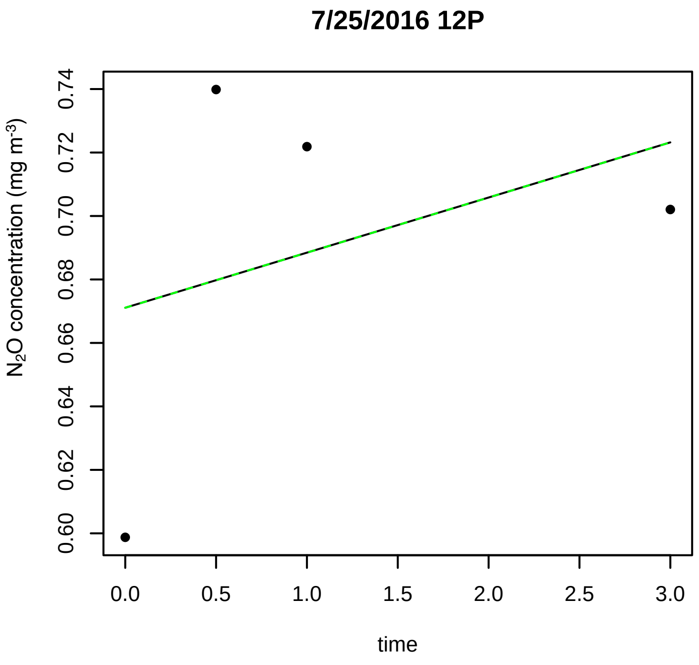
7/25/2016 12P
N2O concentration (mg m-3)
0.60
0.62
0.64
0.66
0.68
0.70
0.72
0.74
0.0
0.5
1.0
1.5
2.0
2.5
3.0
time

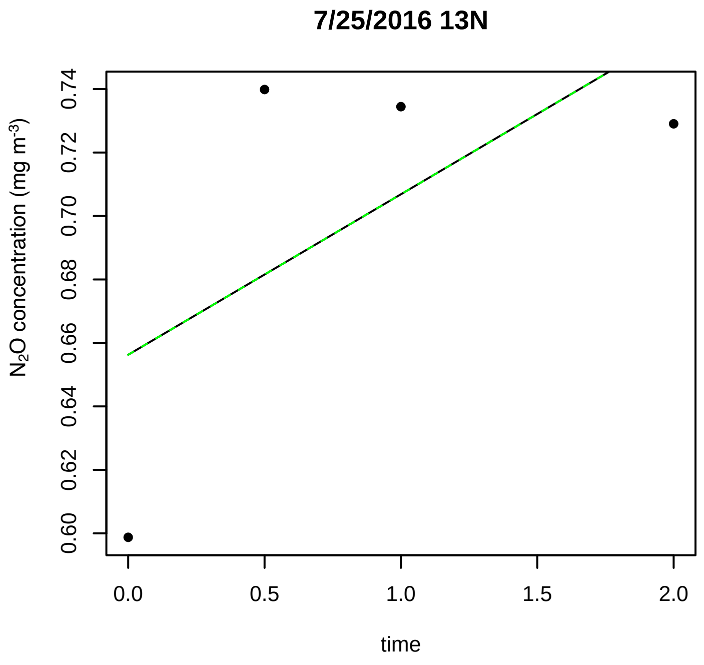
7/25/2016 13N
N2O concentration (mg m-3)
0.60
0.62
0.64
0.66
0.68
0.70
0.72
0.74
0.0
0.5
1.0
1.5
2.0
time

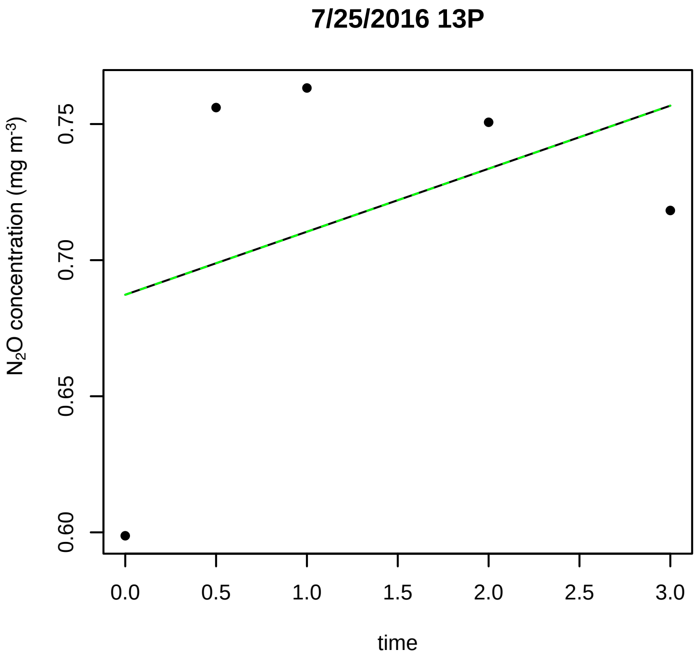
7/25/2016 13P
N2O concentration (mg m-3)
0.60
0.65
0.70
0.75
0.0
0.5
1.0
1.5
2.0
2.5
3.0
time

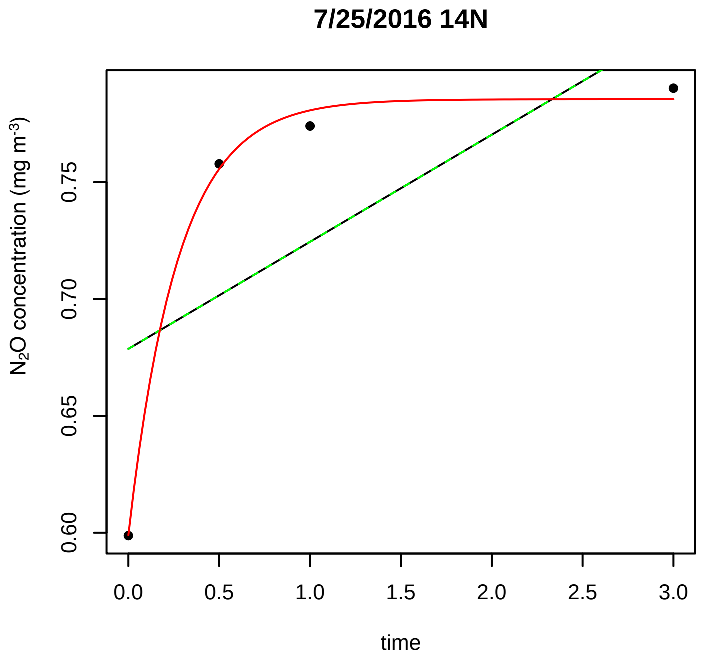
7/25/2016 14N
N2O concentration (mg m-3)
0.60
0.65
0.70
0.75
0.0
0.5
1.0
1.5
2.0
2.5
3.0
time

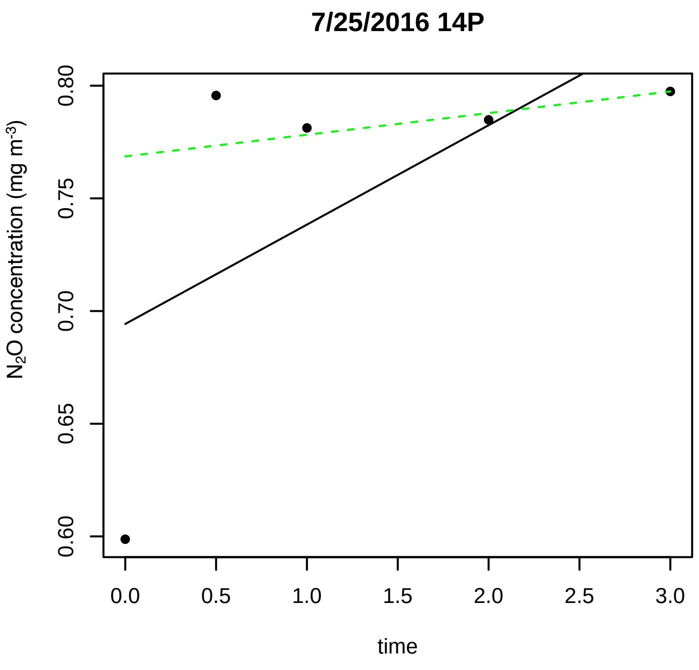
7/25/2016 14P
N2O concentration (mg m-3)
0.60
0.65
0.70
0.75
0.80
0.0
0.5
1.0
1.5
2.0
2.5
3.0
time

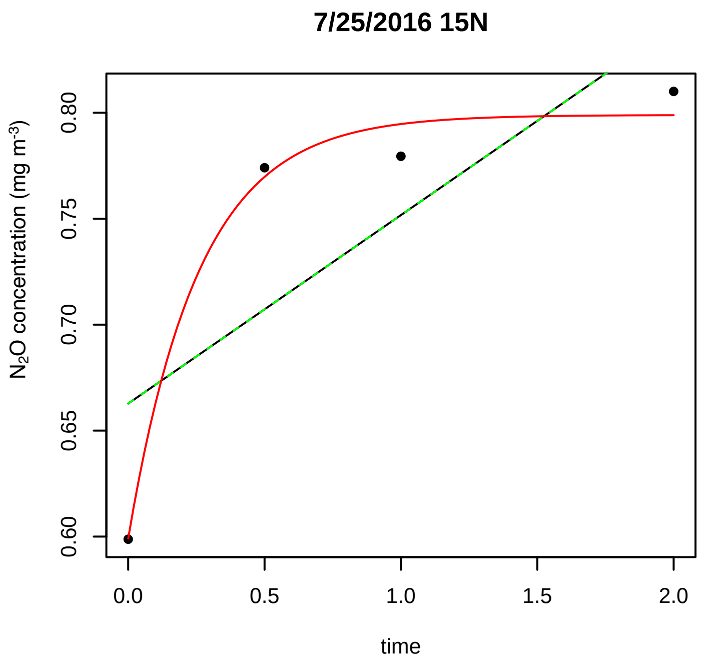
7/25/2016 15N
N2O concentration (mg m-3)
0.60
0.65
0.70
0.75
0.80
0.0
0.5
1.0
1.5
2.0
time

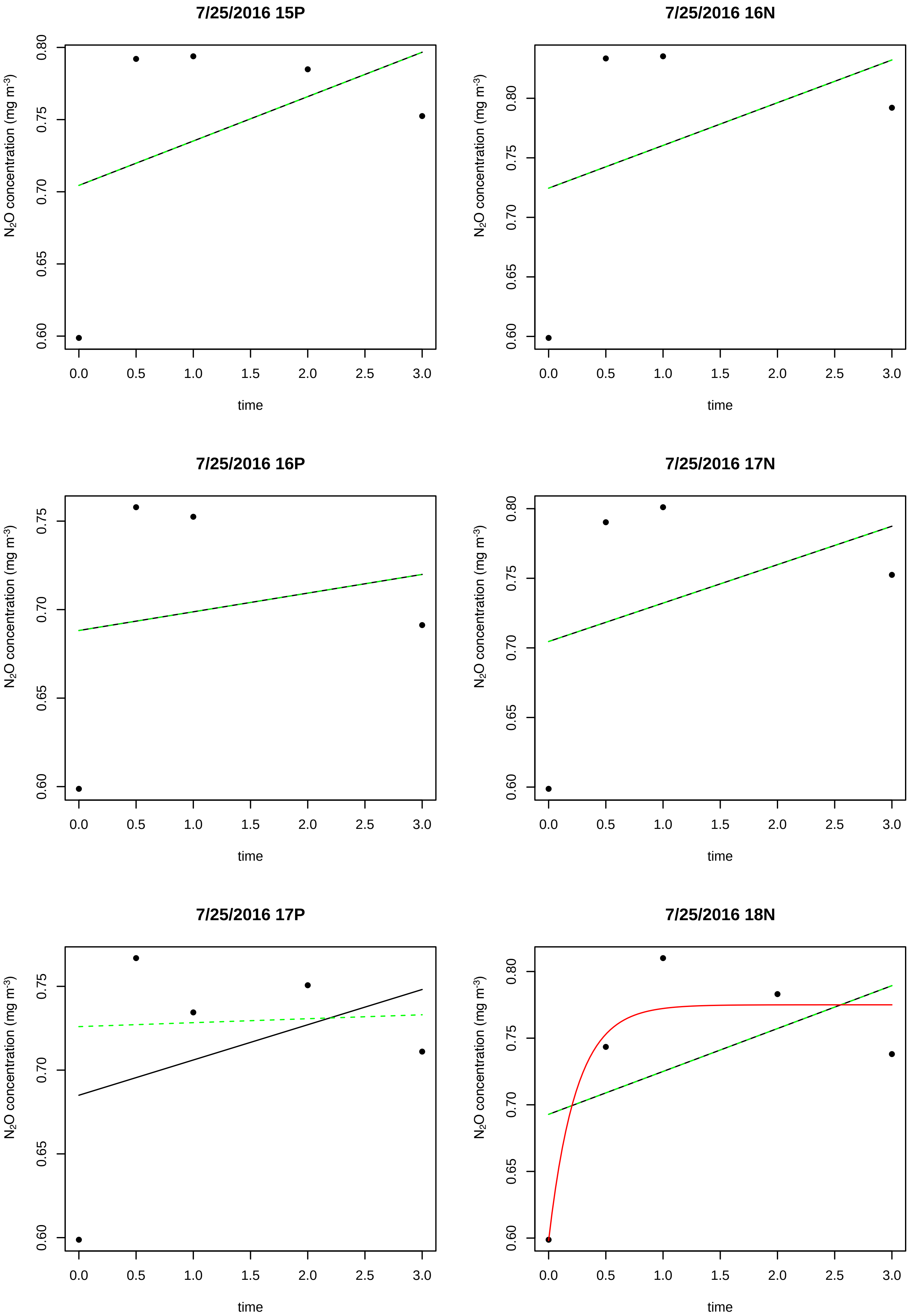
7/25/2016 15P
7/25/2016 16N
7/25/2016 16P
7/25/2016 17N
7/25/2016 17P
7/25/2016 18N
N$_2$O concentration (mg m$^{-3}$)
time
0.60
0.65
0.70
0.75
0.80
0.0
0.5
1.0
1.5
2.0
2.5
3.0

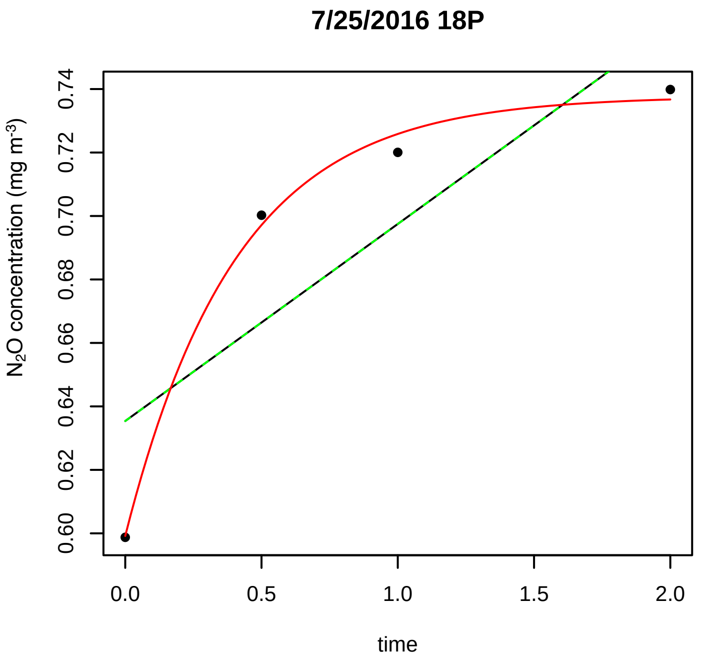
7/25/2016 18P
$N_2O$ concentration (mg $m^{-3}$)
0.60
0.62
0.64
0.66
0.68
0.70
0.72
0.74
0.0
0.5
1.0
1.5
2.0
time

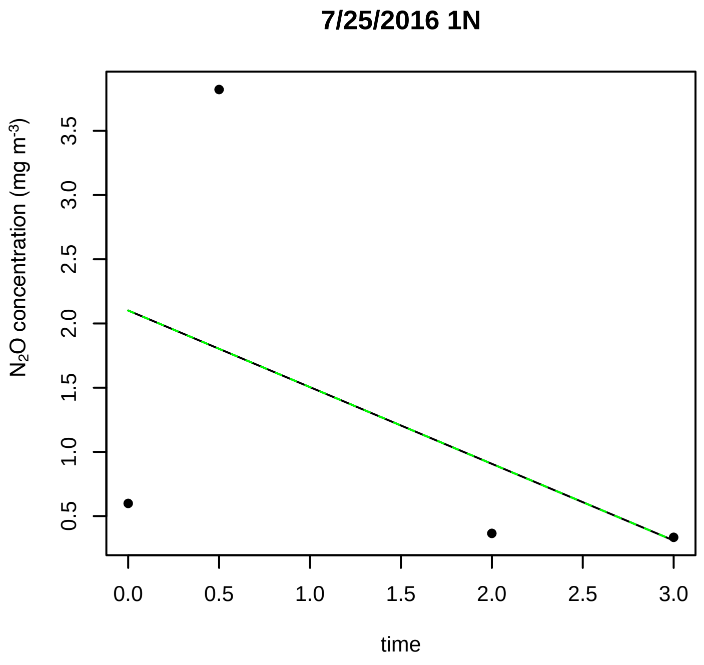
7/25/2016 1N
$N_2O$ concentration (mg $m^{-3}$)
0.5
1.0
1.5
2.0
2.5
3.0
3.5
0.0
0.5
1.0
1.5
2.0
2.5
3.0
time

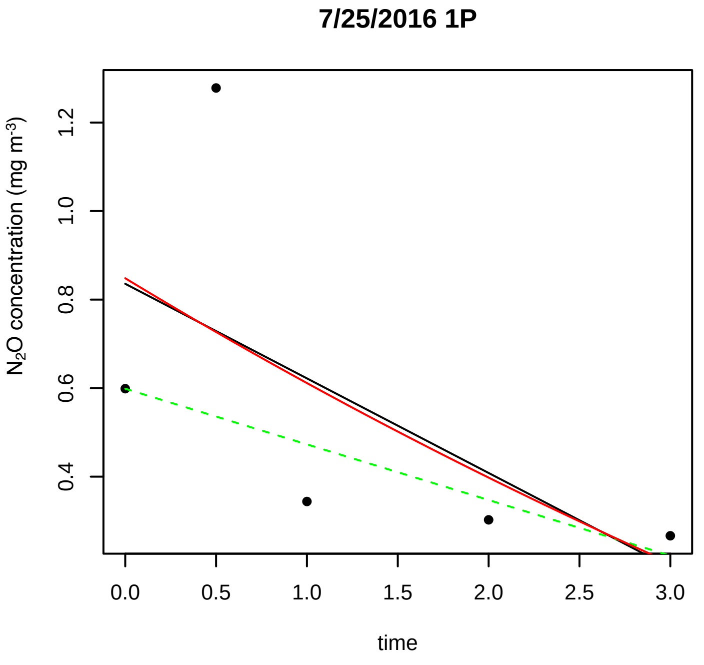
7/25/2016 1P
$N_2O$ concentration (mg $m^{-3}$)
0.4
0.6
0.8
1.0
1.2
0.0
0.5
1.0
1.5
2.0
2.5
3.0
time

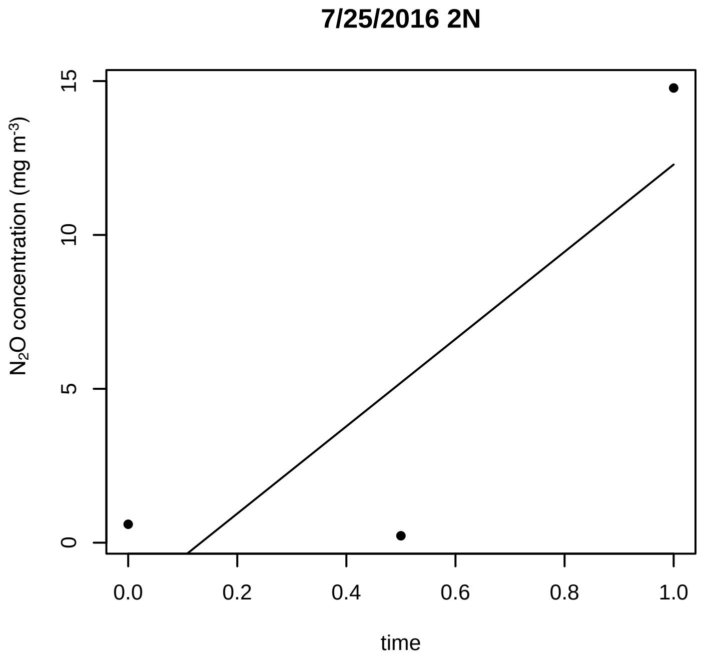
7/25/2016 2N
$N_2O$ concentration (mg $m^{-3}$)
0
5
10
15
0.0
0.2
0.4
0.6
0.8
1.0
time

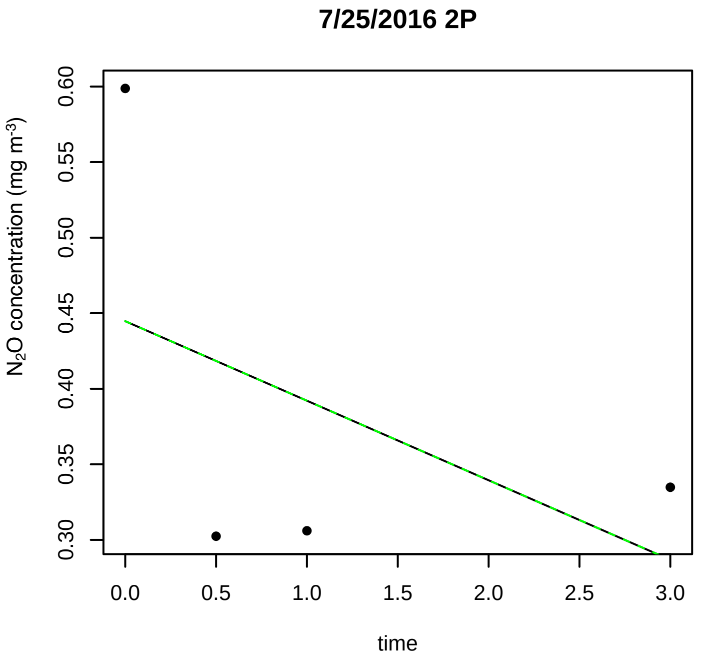
7/25/2016 2P
$N_2O$ concentration (mg $m^{-3}$)
0.30
0.35
0.40
0.45
0.50
0.55
0.60
0.0
0.5
1.0
1.5
2.0
2.5
3.0
time

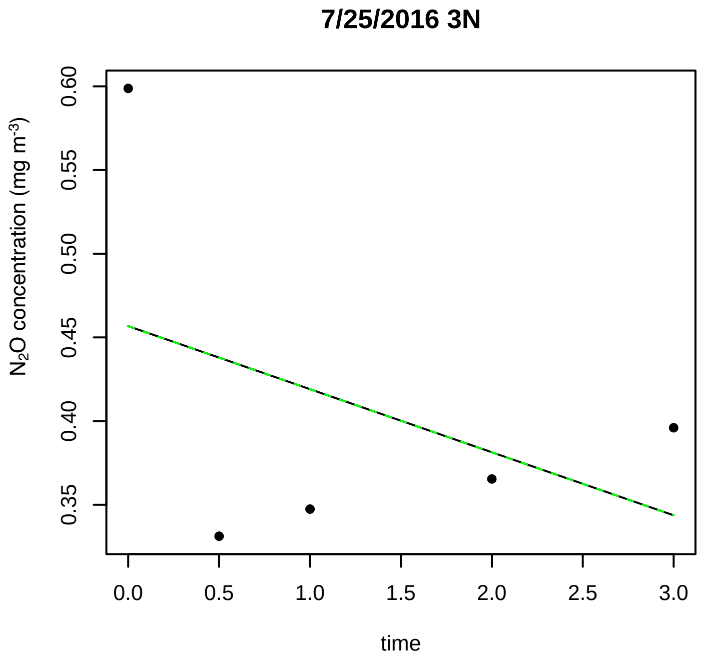
7/25/2016 3N
$N_2O$ concentration (mg $m^{-3}$)
0.35
0.40
0.45
0.50
0.55
0.60
0.0
0.5
1.0
1.5
2.0
2.5
3.0
time

**7/25/2016 3P**

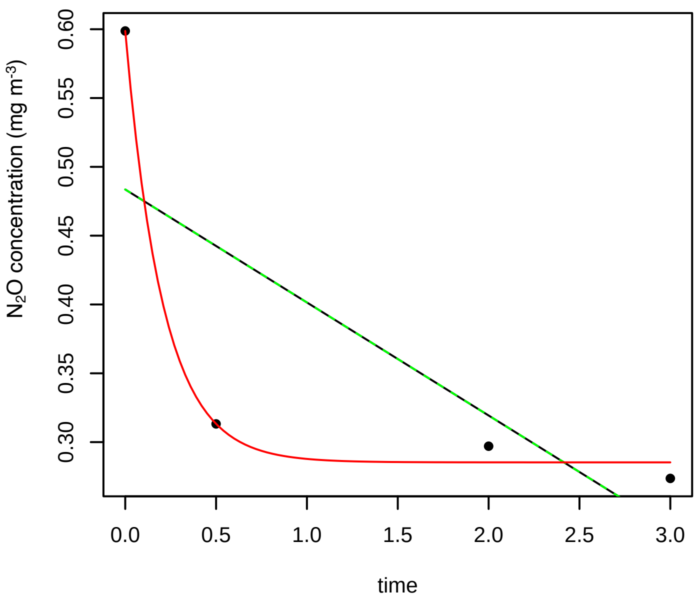


**7/25/2016 4N**

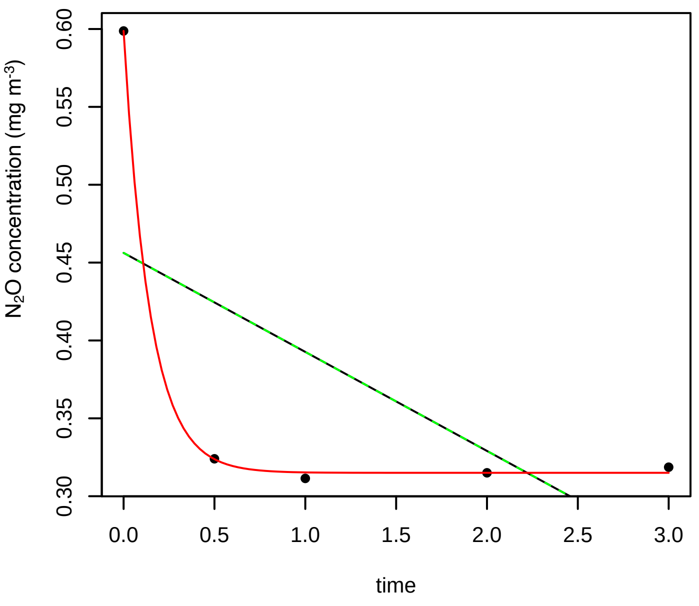


**7/25/2016 4P**

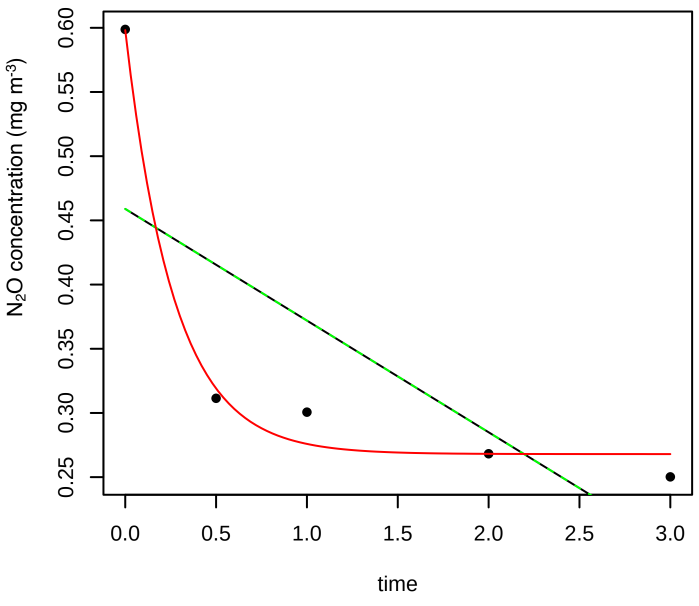


**7/25/2016 5N**

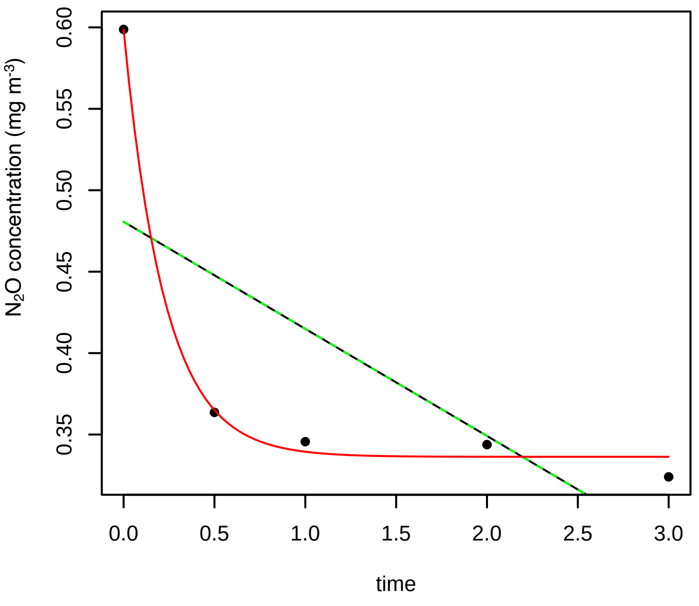


**7/25/2016 5P**

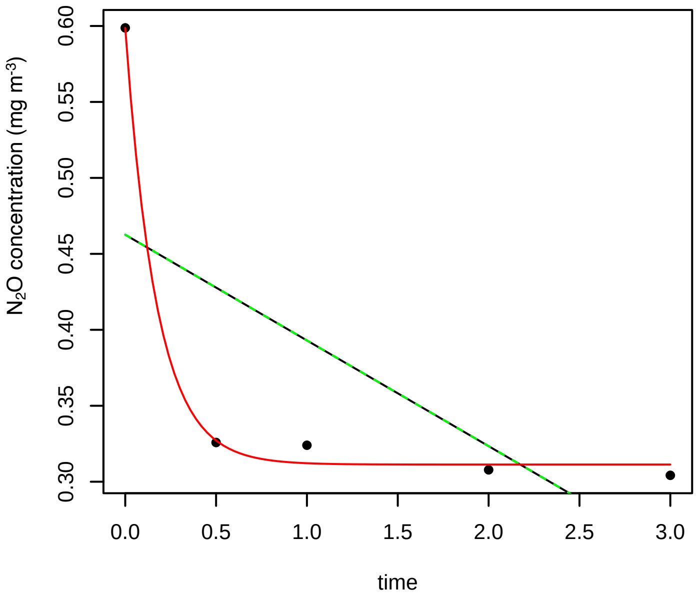


**7/25/2016 6N**

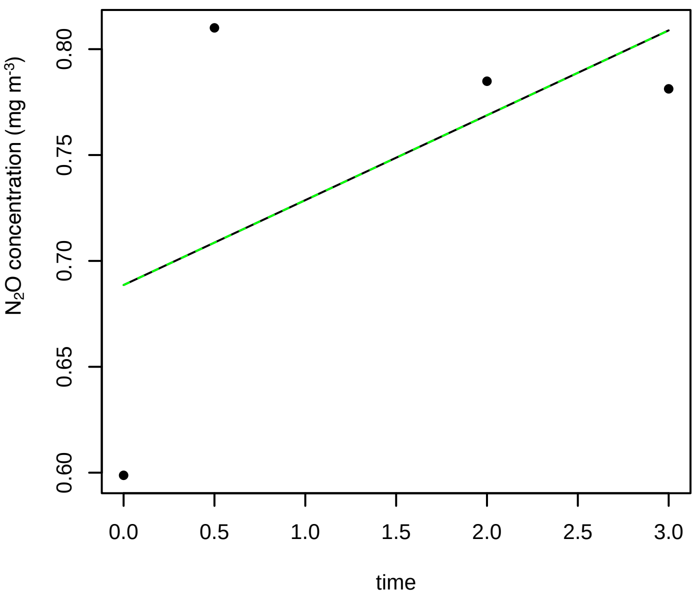

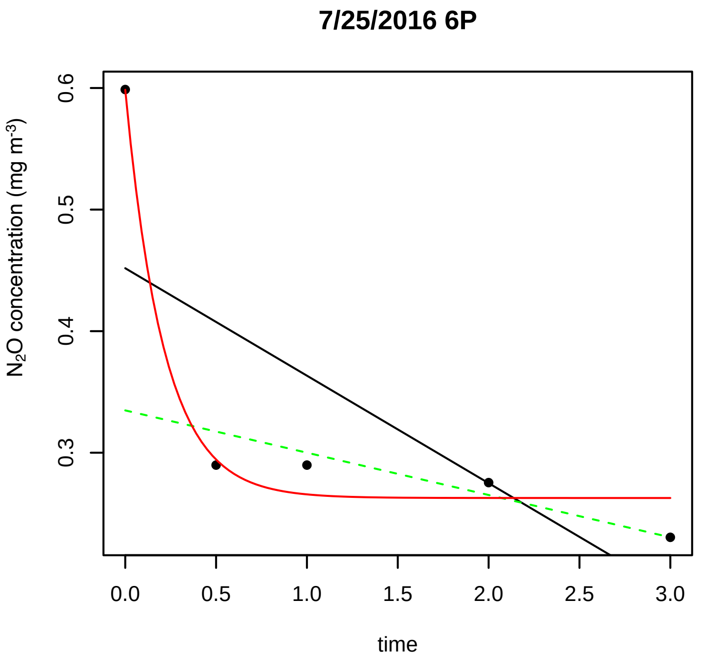
7/25/2016 6P
N2O concentration (mg m-3)
0.3
0.4
0.5
0.6
0.0
0.5
1.0
1.5
2.0
2.5
3.0
time

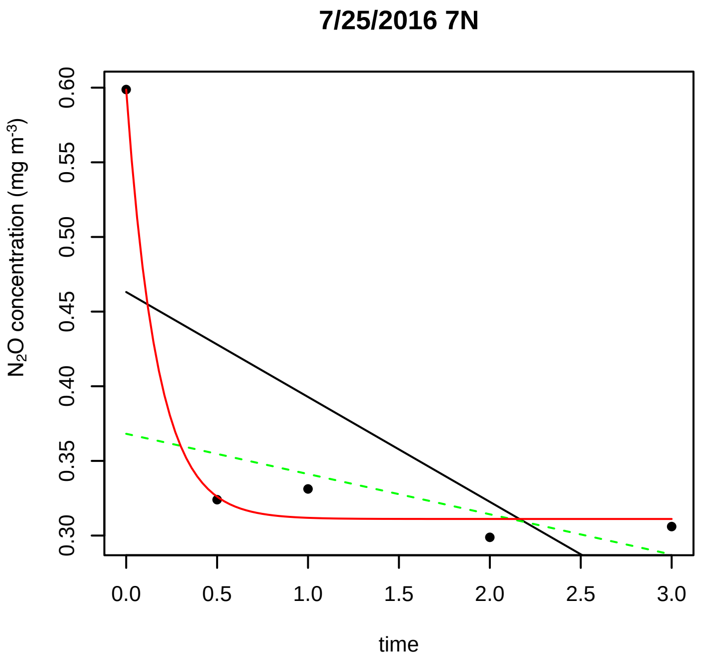
7/25/2016 7N
N2O concentration (mg m-3)
0.30
0.35
0.40
0.45
0.50
0.55
0.60
0.0
0.5
1.0
1.5
2.0
2.5
3.0
time

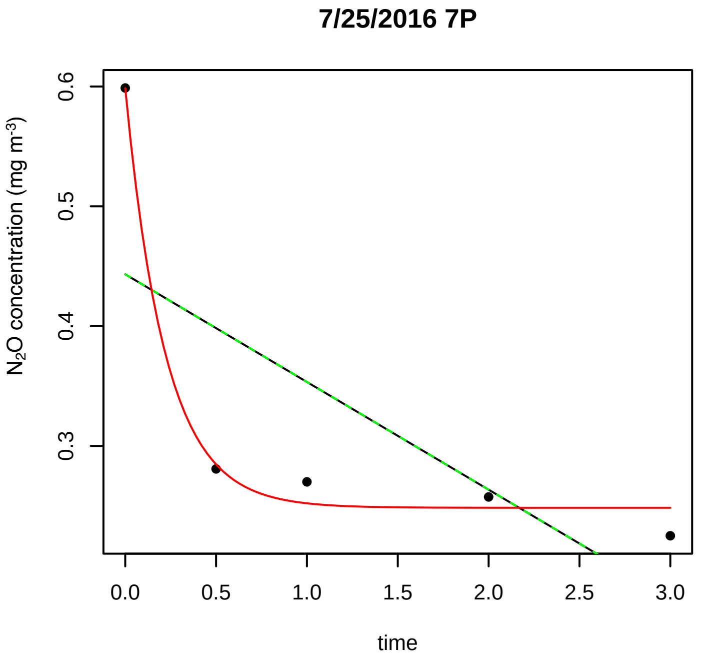
7/25/2016 7P
N2O concentration (mg m-3)
0.3
0.4
0.5
0.6
0.0
0.5
1.0
1.5
2.0
2.5
3.0
time

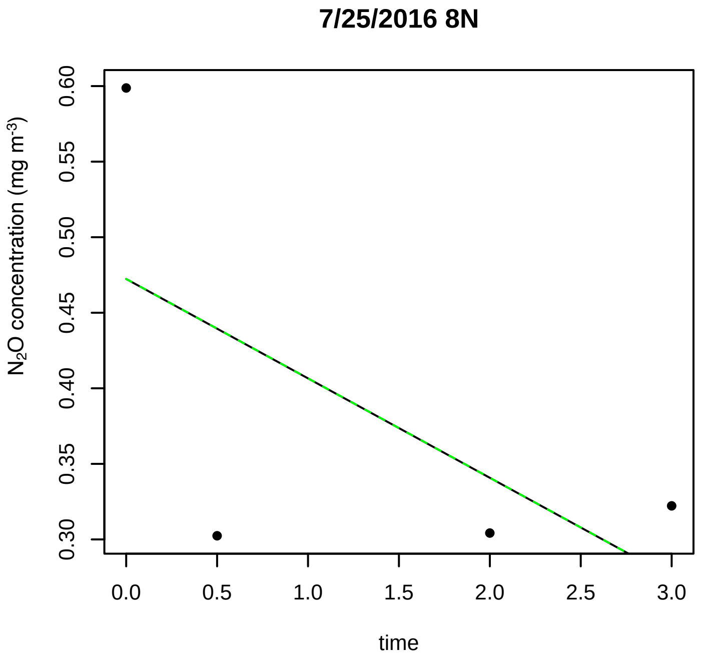
7/25/2016 8N
N2O concentration (mg m-3)
0.30
0.35
0.40
0.45
0.50
0.55
0.60
0.0
0.5
1.0
1.5
2.0
2.5
3.0
time

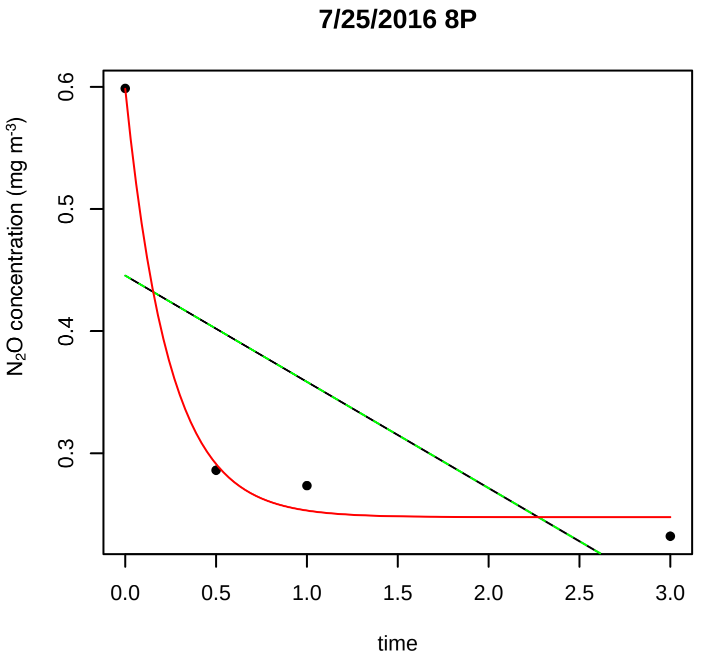
7/25/2016 8P
N2O concentration (mg m-3)
0.3
0.4
0.5
0.6
0.0
0.5
1.0
1.5
2.0
2.5
3.0
time

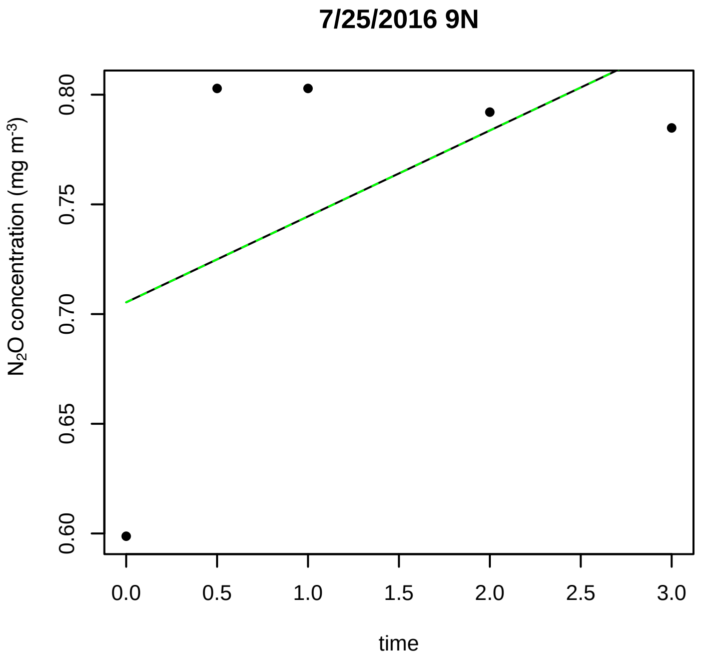
7/25/2016 9N
N2O concentration (mg m-3)
0.60
0.65
0.70
0.75
0.80
0.0
0.5
1.0
1.5
2.0
2.5
3.0
time

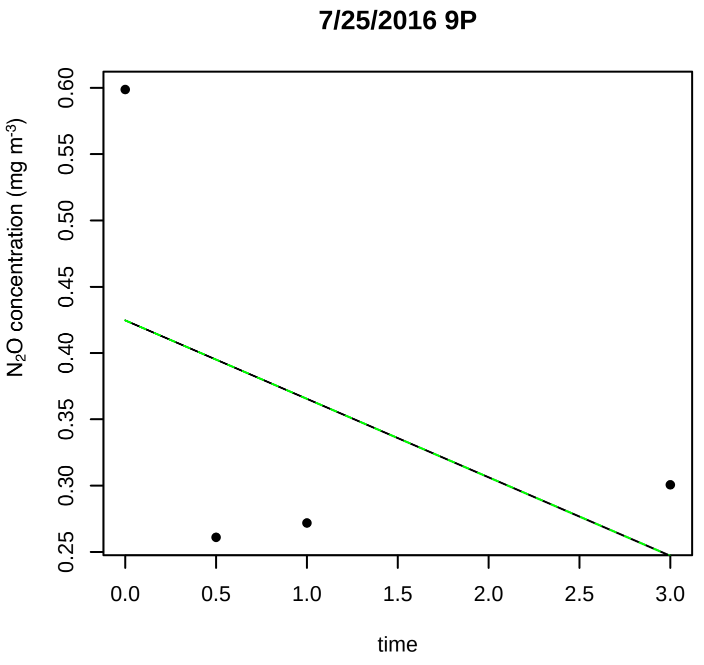
7/25/2016 9P
N2O concentration (mg m-3)
time

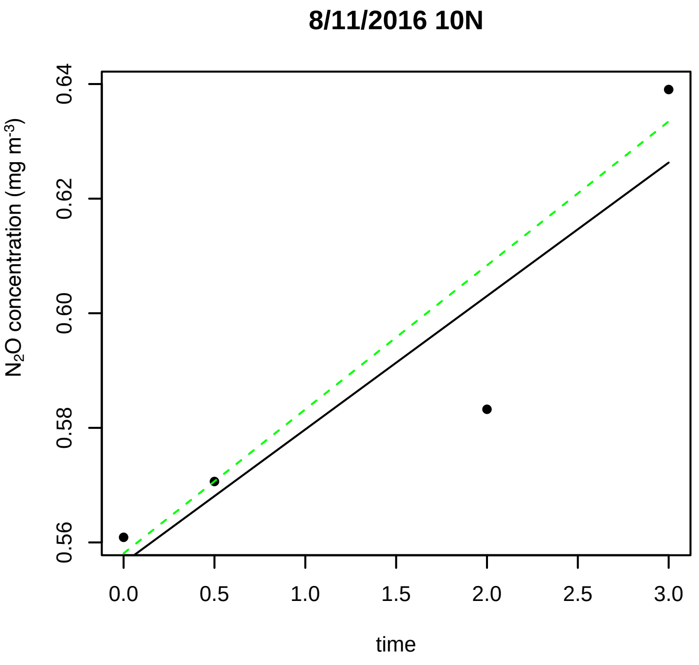
8/11/2016 10N
N2O concentration (mg m-3)
time

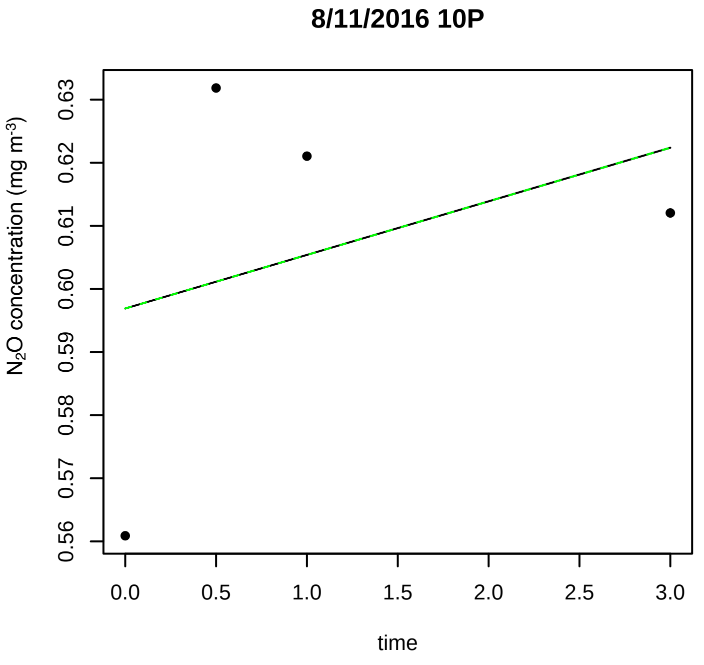
8/11/2016 10P
N2O concentration (mg m-3)
time

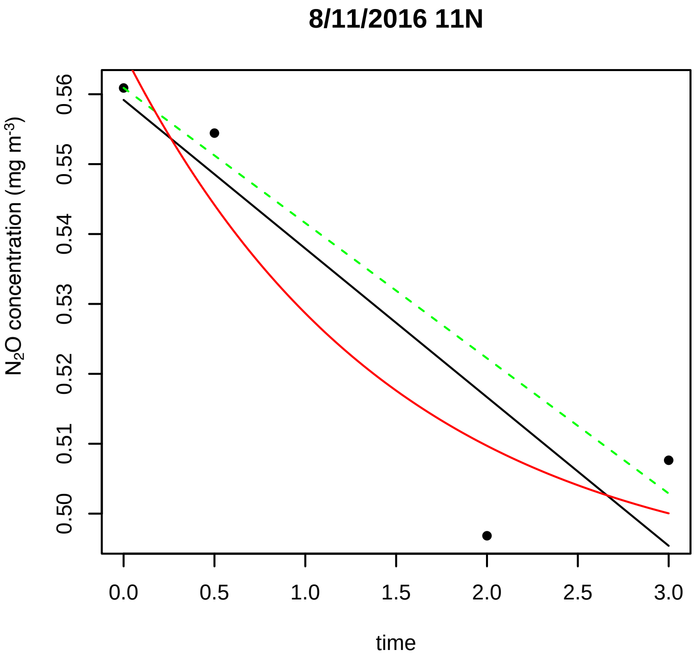
8/11/2016 11N
N2O concentration (mg m-3)
time

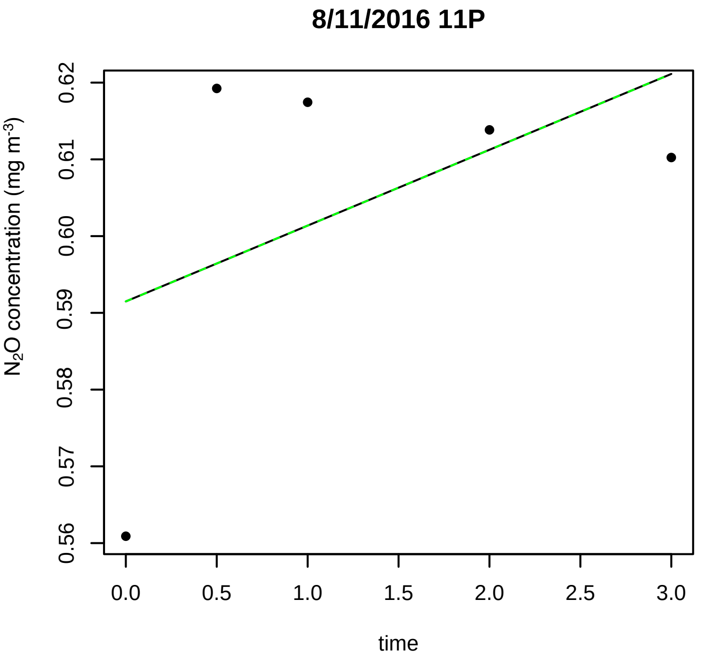
8/11/2016 11P
N2O concentration (mg m-3)
time

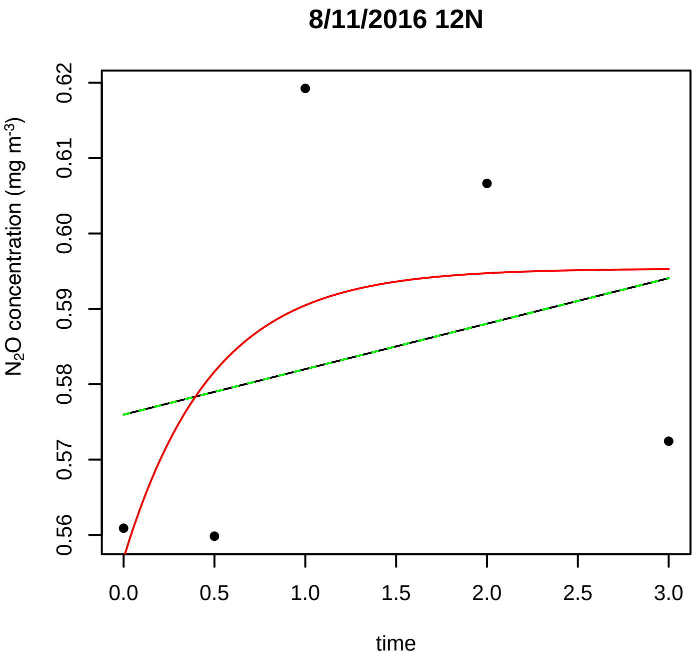
8/11/2016 12N
N2O concentration (mg m-3)
time

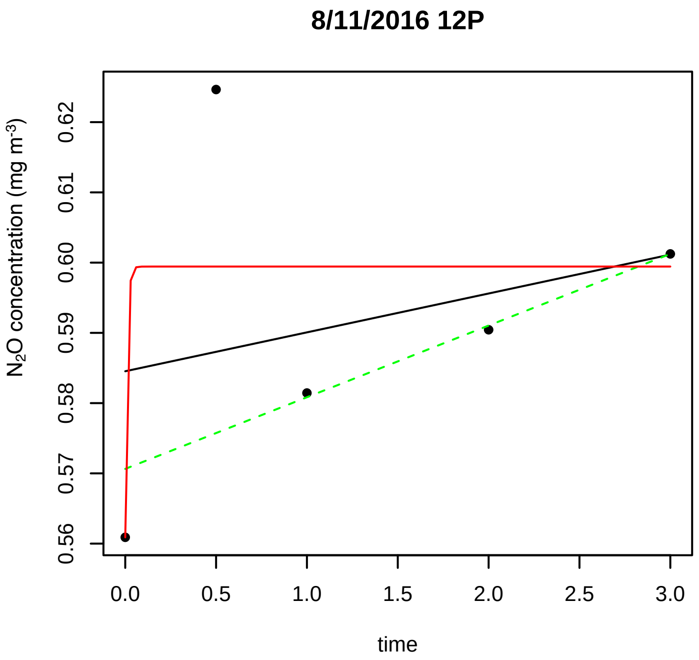
8/11/2016 12P
N2O concentration (mg m-3)
0.56
0.57
0.58
0.59
0.60
0.61
0.62
0.0
0.5
1.0
1.5
2.0
2.5
3.0
time

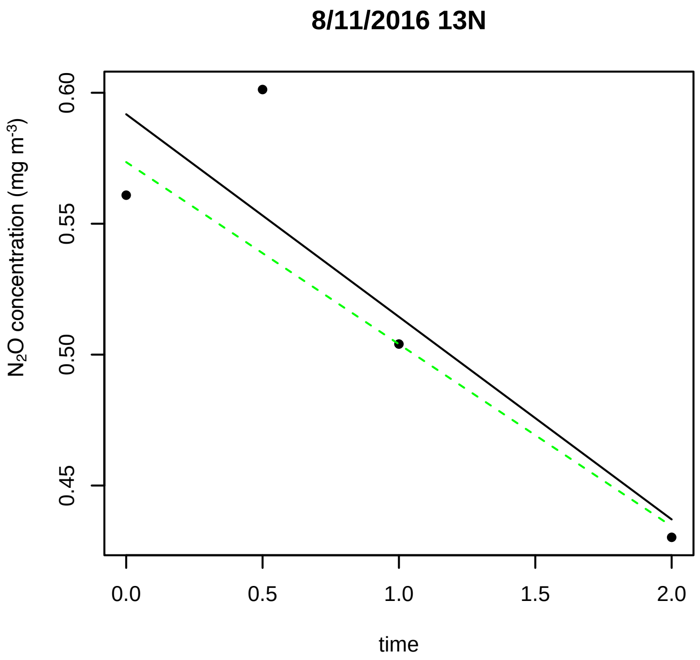
8/11/2016 13N
N2O concentration (mg m-3)
0.45
0.50
0.55
0.60
0.0
0.5
1.0
1.5
2.0
time

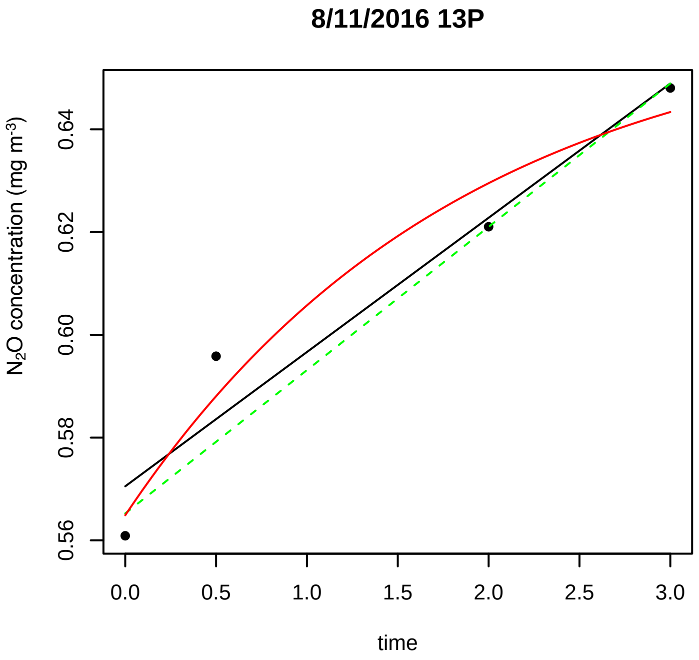
8/11/2016 13P
N2O concentration (mg m-3)
0.56
0.58
0.60
0.62
0.64
0.0
0.5
1.0
1.5
2.0
2.5
3.0
time

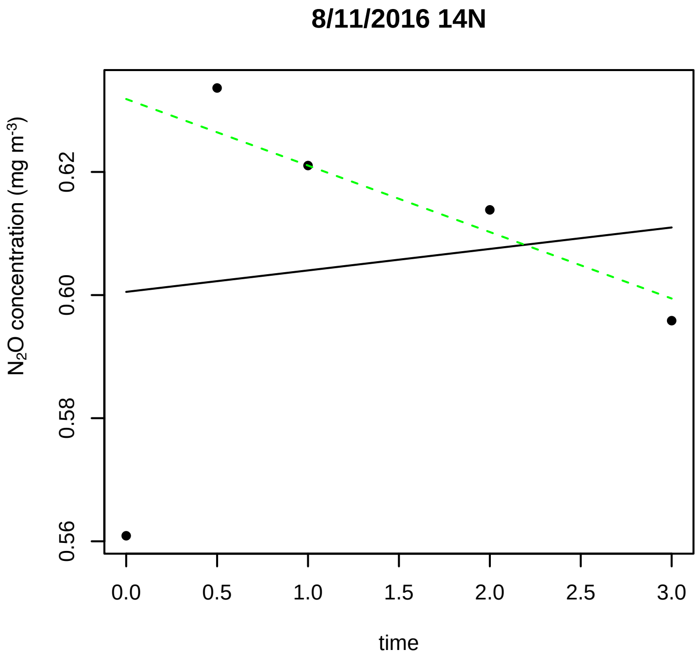
8/11/2016 14N
N2O concentration (mg m-3)
0.56
0.58
0.60
0.62
0.0
0.5
1.0
1.5
2.0
2.5
3.0
time

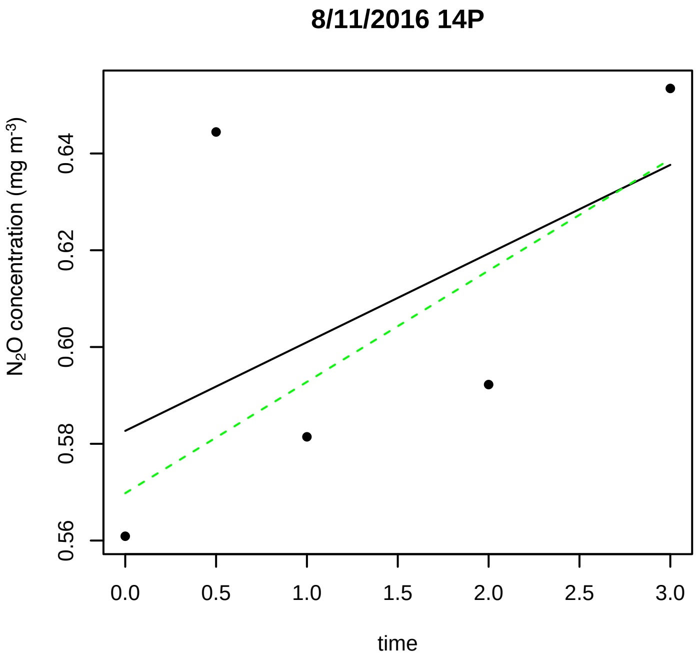
8/11/2016 14P
N2O concentration (mg m-3)
0.56
0.58
0.60
0.62
0.64
0.0
0.5
1.0
1.5
2.0
2.5
3.0
time

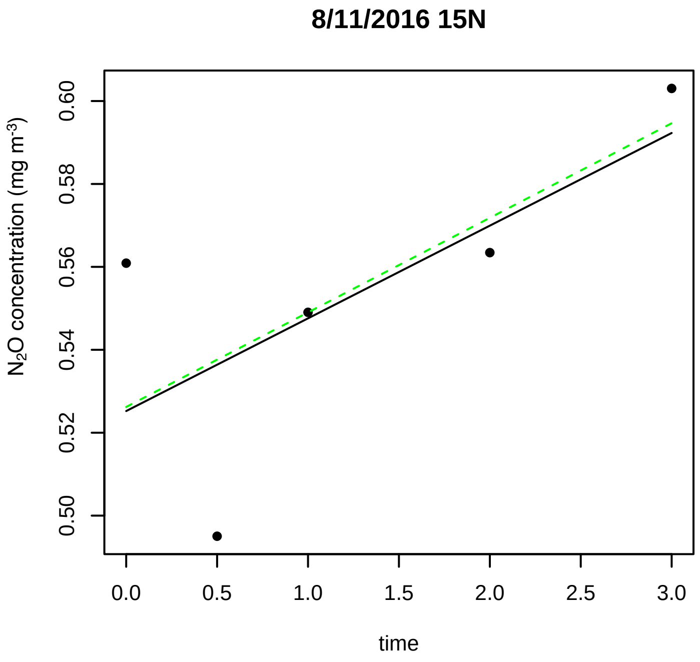
8/11/2016 15N
N2O concentration (mg m-3)
0.50
0.52
0.54
0.56
0.58
0.60
0.0
0.5
1.0
1.5
2.0
2.5
3.0
time

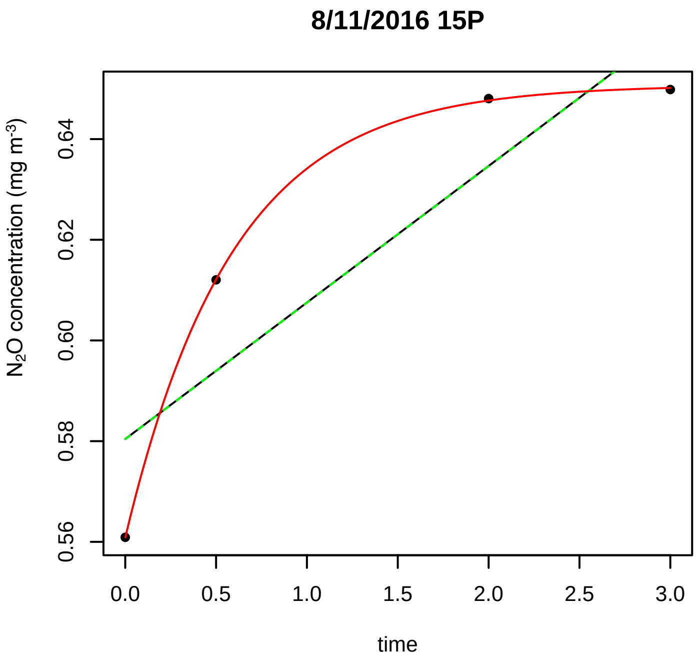
8/11/2016 15P
N2O concentration (mg m-3)
0.56
0.58
0.60
0.62
0.64
0.0
0.5
1.0
1.5
2.0
2.5
3.0
time

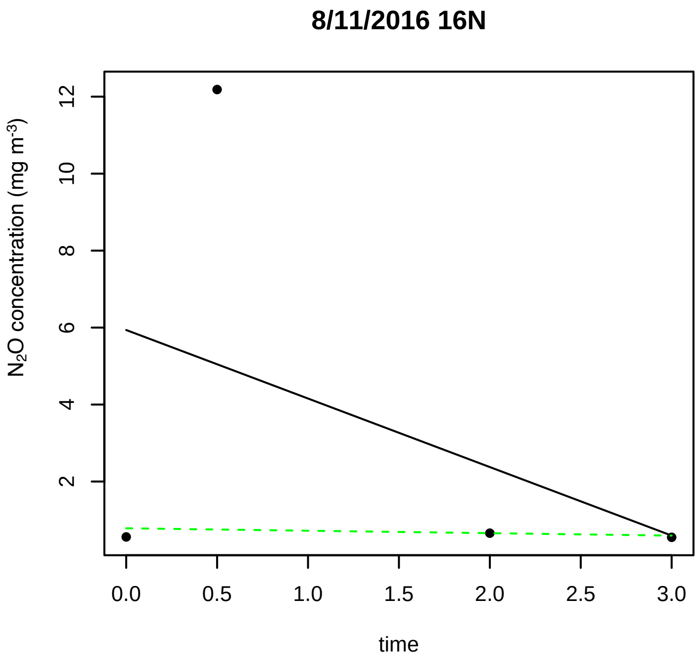
8/11/2016 16N
N2O concentration (mg m-3)
2
4
6
8
10
12
0.0
0.5
1.0
1.5
2.0
2.5
3.0
time

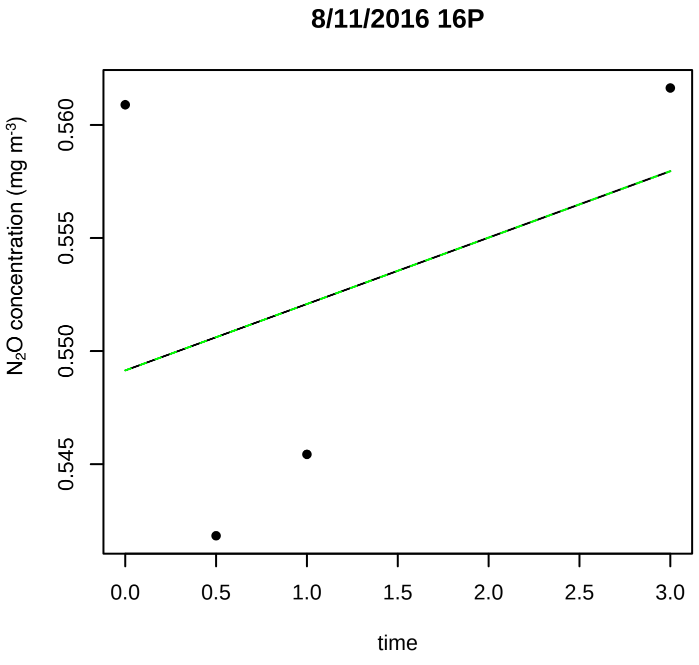
8/11/2016 16P
N2O concentration (mg m-3)
0.545
0.550
0.555
0.560
0.0
0.5
1.0
1.5
2.0
2.5
3.0
time

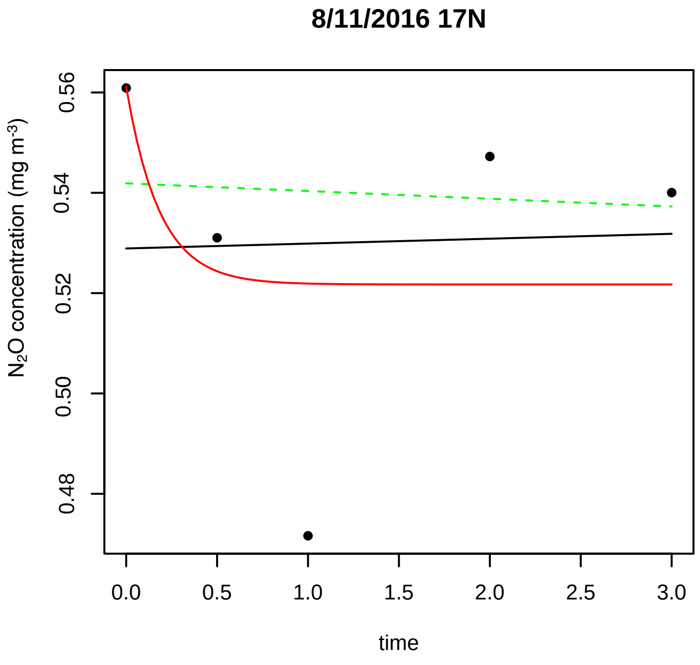
8/11/2016 17N
N2O concentration (mg m-3)
0.48
0.50
0.52
0.54
0.56
0.0
0.5
1.0
1.5
2.0
2.5
3.0
time

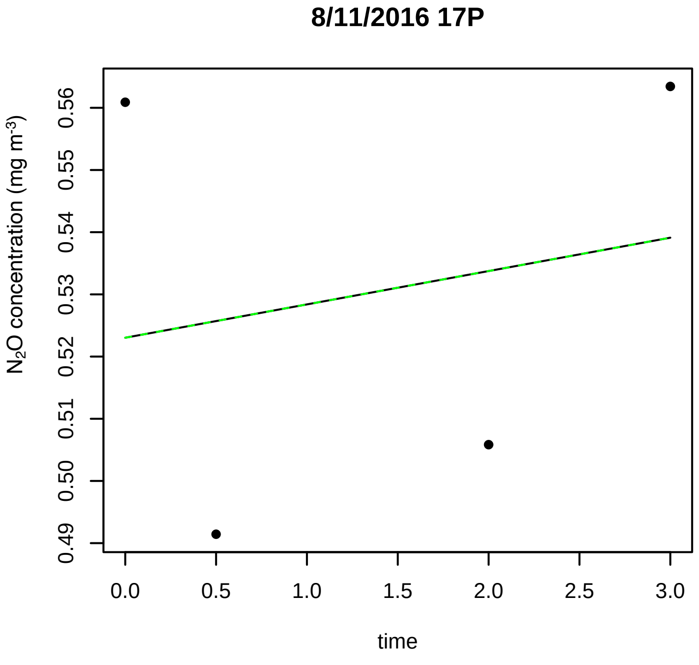
8/11/2016 17P
N2O concentration (mg m-3)
0.49
0.50
0.51
0.52
0.53
0.54
0.55
0.56
0.0
0.5
1.0
1.5
2.0
2.5
3.0
time

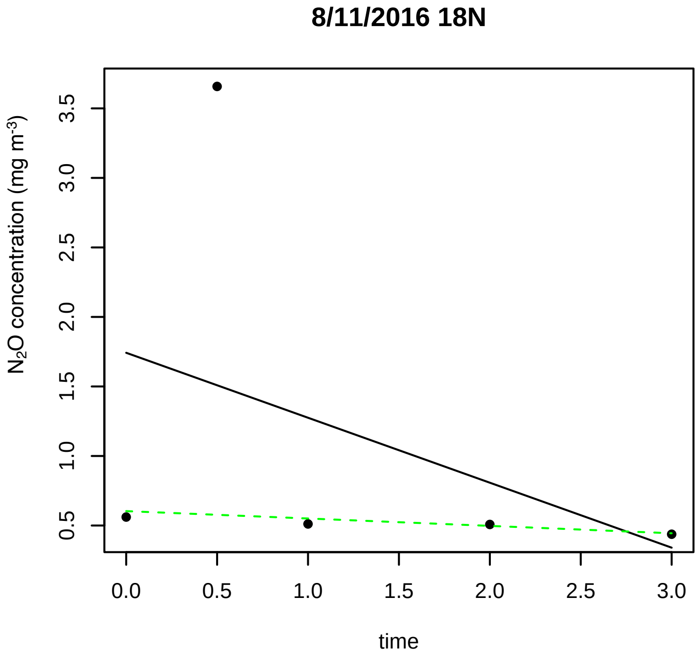
8/11/2016 18N
N2O concentration (mg m-3)
0.5
1.0
1.5
2.0
2.5
3.0
3.5
0.0
0.5
1.0
1.5
2.0
2.5
3.0
time

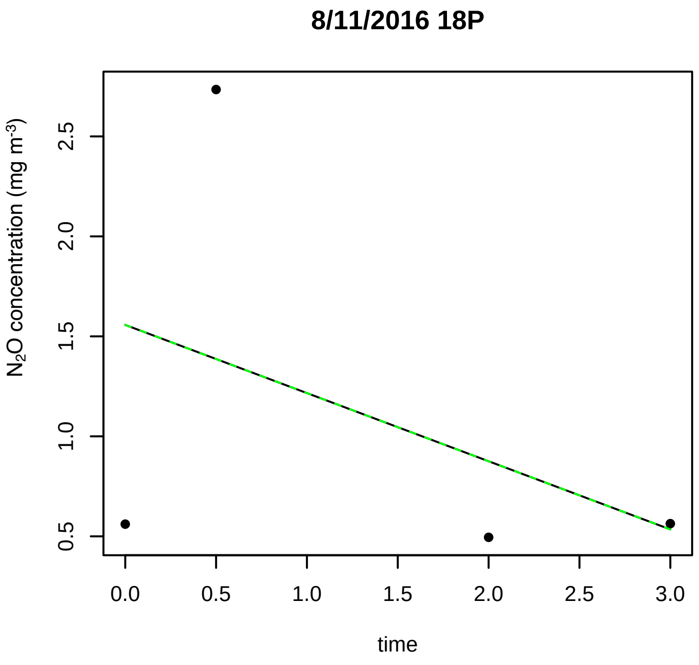
8/11/2016 18P
N$_2$O concentration (mg m$^{-3}$)
0.5
1.0
1.5
2.0
2.5
time
0.0
0.5
1.0
1.5
2.0
2.5
3.0

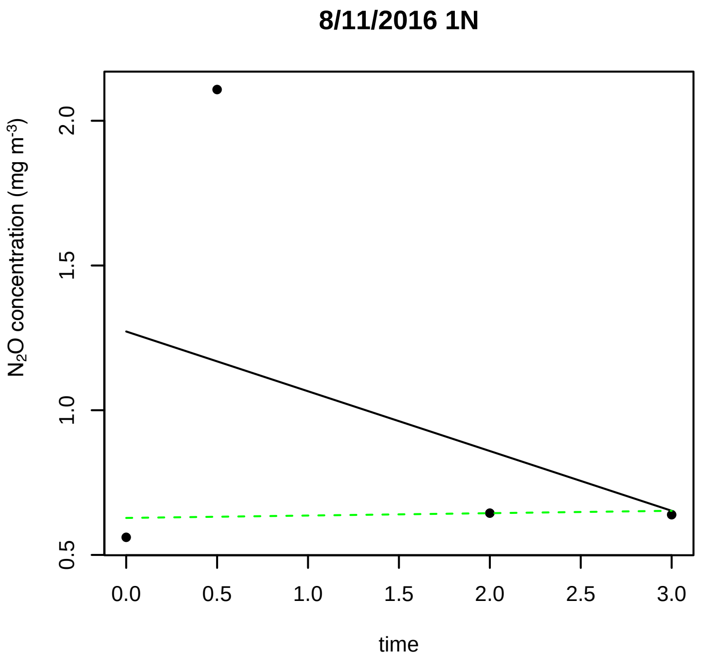
8/11/2016 1N
N$_2$O concentration (mg m$^{-3}$)
0.5
1.0
1.5
2.0
time
0.0
0.5
1.0
1.5
2.0
2.5
3.0

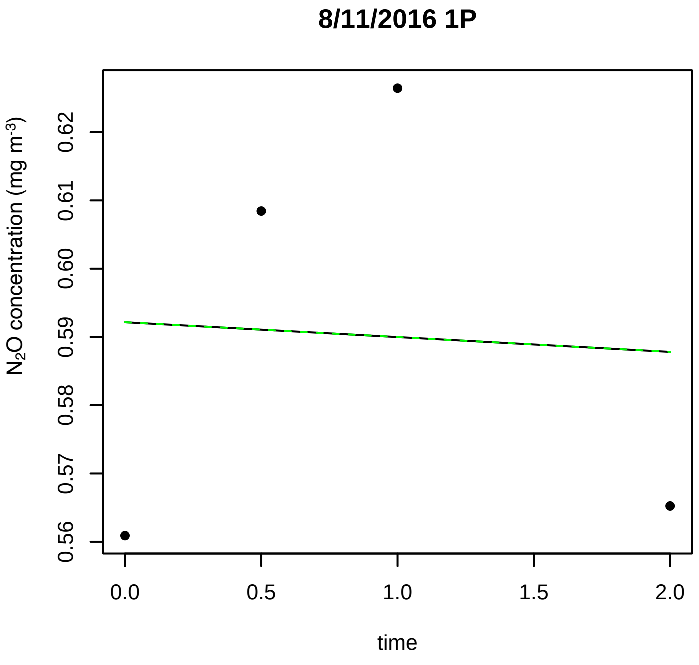
8/11/2016 1P
N$_2$O concentration (mg m$^{-3}$)
0.56
0.57
0.58
0.59
0.60
0.61
0.62
time
0.0
0.5
1.0
1.5
2.0

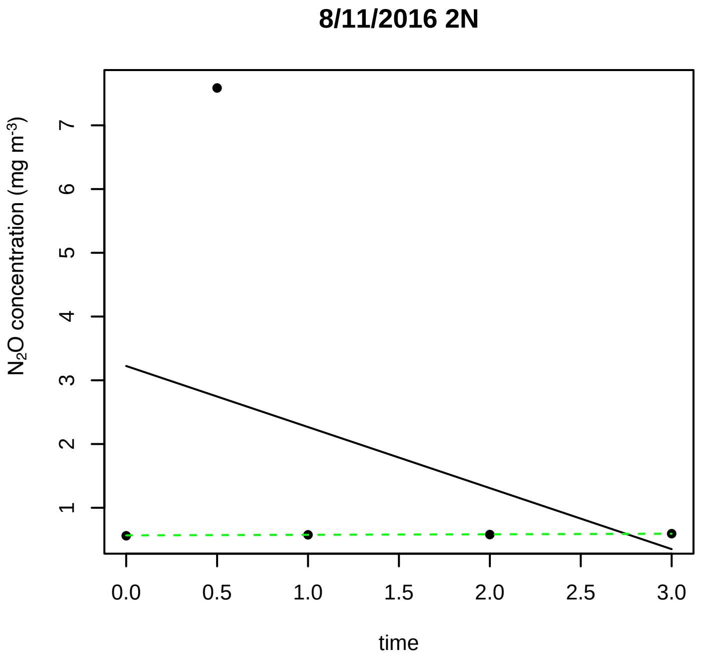
8/11/2016 2N
N$_2$O concentration (mg m$^{-3}$)
1
2
3
4
5
6
7
time
0.0
0.5
1.0
1.5
2.0
2.5
3.0

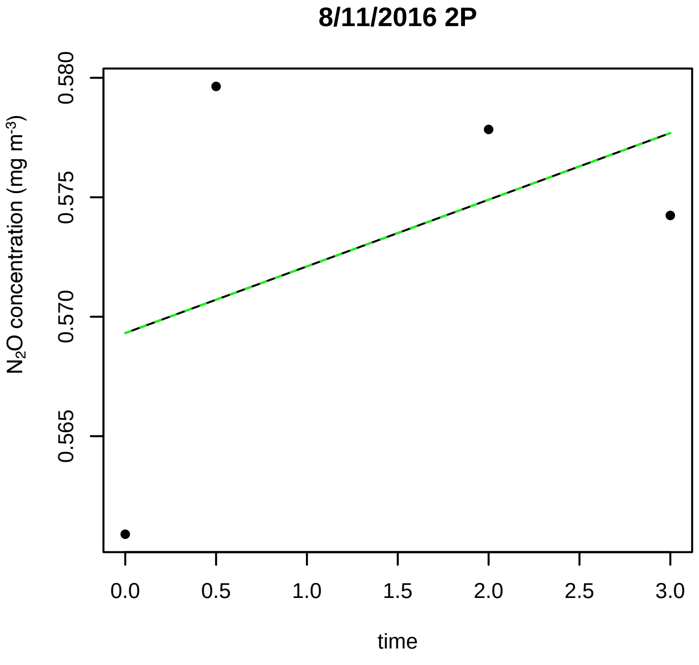
8/11/2016 2P
N$_2$O concentration (mg m$^{-3}$)
0.565
0.570
0.575
0.580
time
0.0
0.5
1.0
1.5
2.0
2.5
3.0

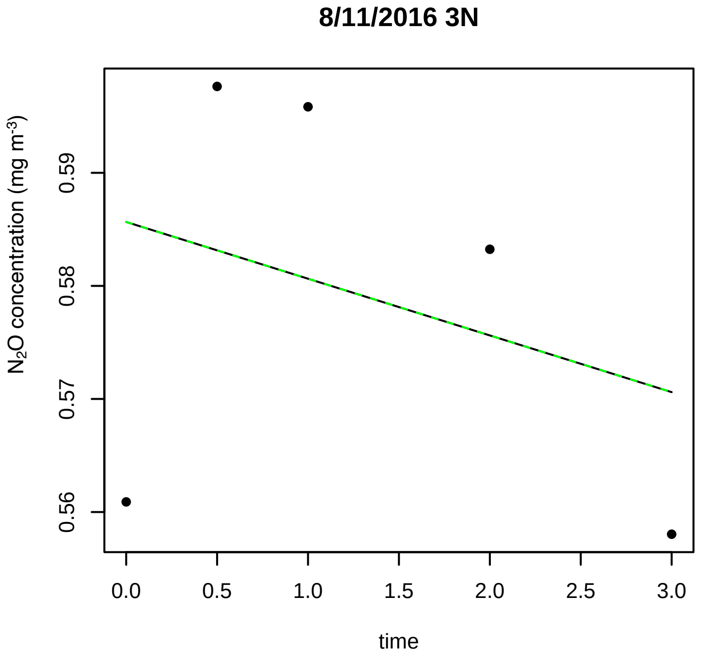
8/11/2016 3N
N$_2$O concentration (mg m$^{-3}$)
0.56
0.57
0.58
0.59
time
0.0
0.5
1.0
1.5
2.0
2.5
3.0

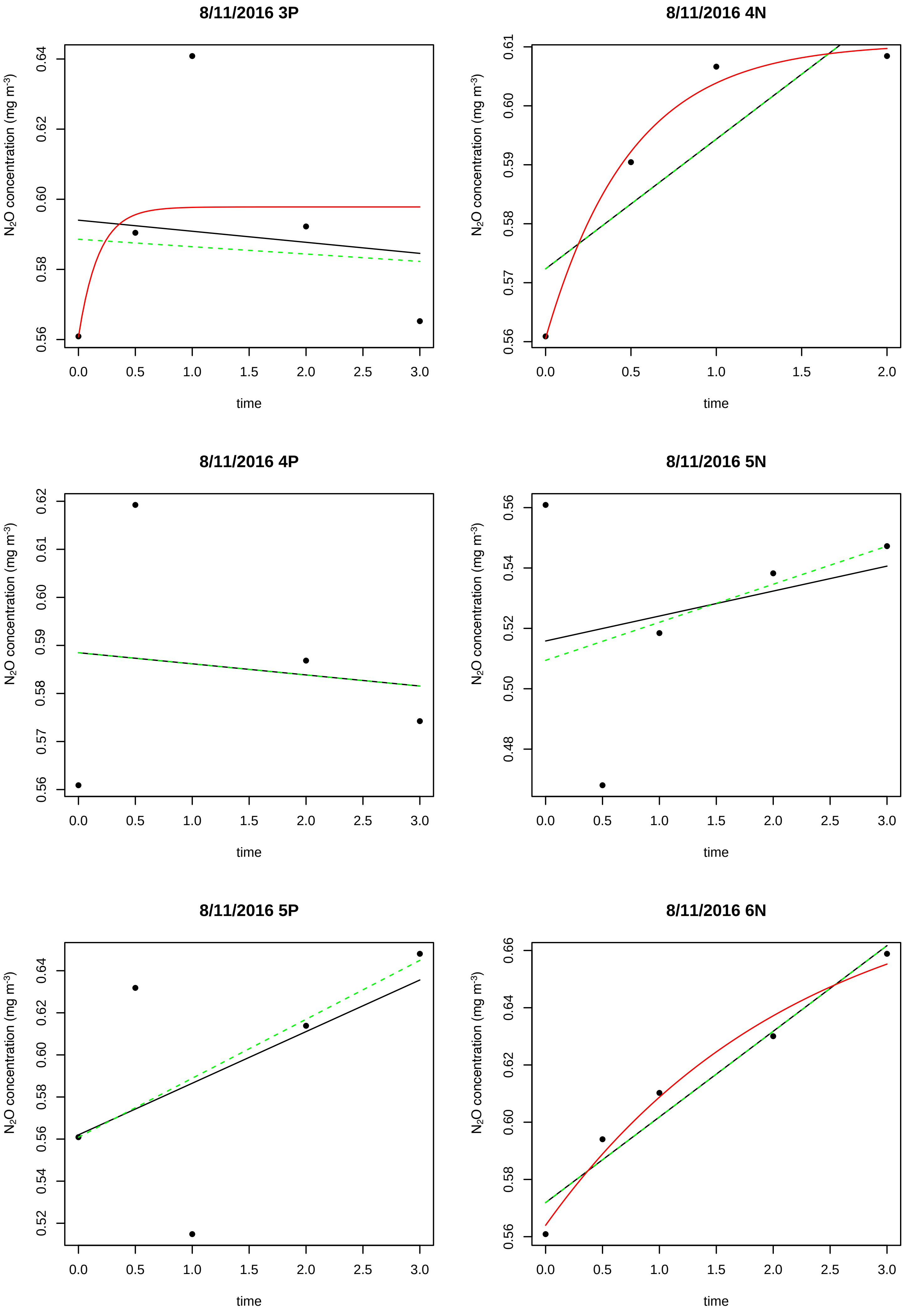
8/11/2016 3P
8/11/2016 4N
8/11/2016 4P
8/11/2016 5N
8/11/2016 5P
8/11/2016 6N
$N_2O$ concentration (mg $m^{-3}$)
time

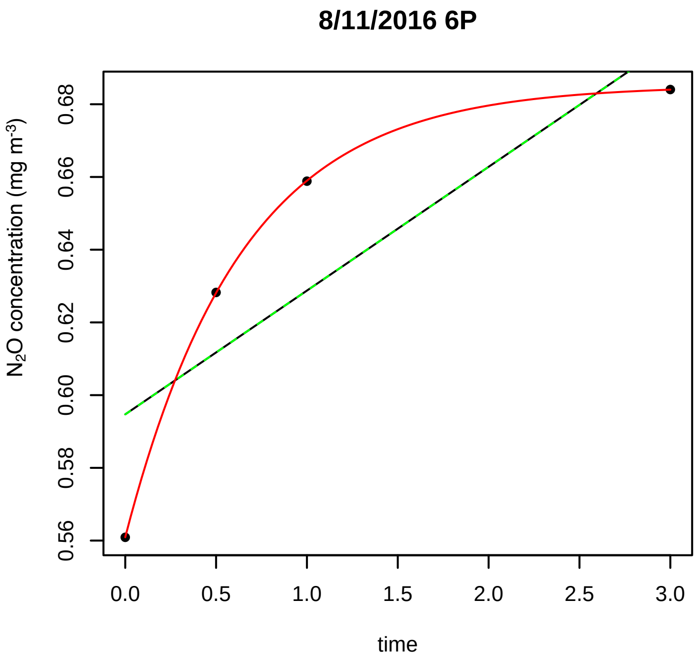
8/11/2016 6P
N2O concentration (mg m-3)
0.56
0.58
0.60
0.62
0.64
0.66
0.68
0.0
0.5
1.0
1.5
2.0
2.5
3.0
time

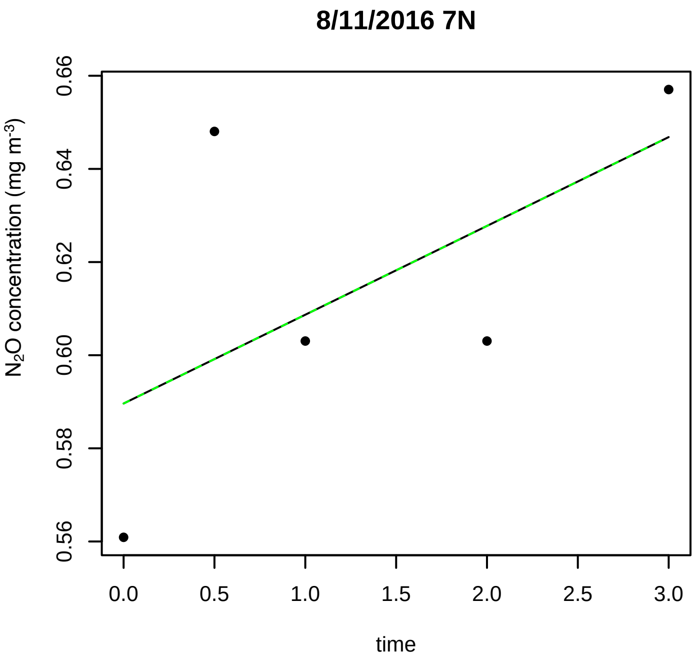
8/11/2016 7N
N2O concentration (mg m-3)
0.56
0.58
0.60
0.62
0.64
0.66
0.0
0.5
1.0
1.5
2.0
2.5
3.0
time

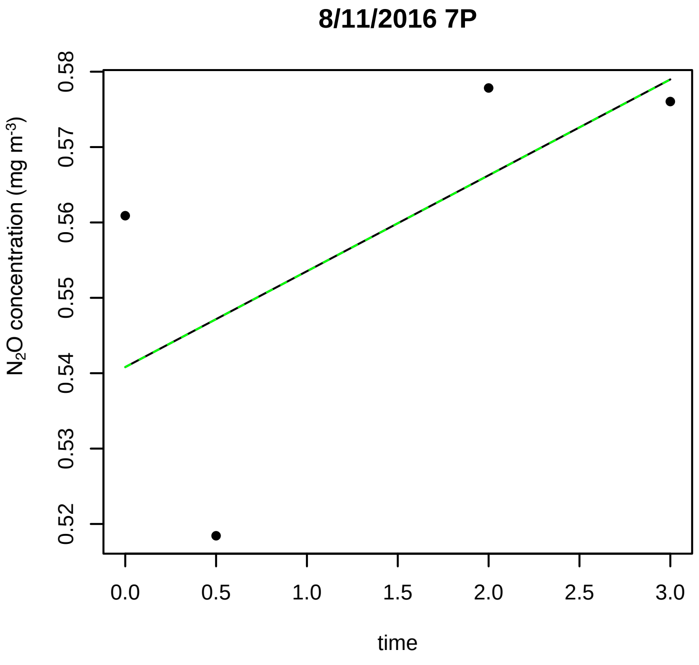
8/11/2016 7P
N2O concentration (mg m-3)
0.52
0.53
0.54
0.55
0.56
0.57
0.58
0.0
0.5
1.0
1.5
2.0
2.5
3.0
time

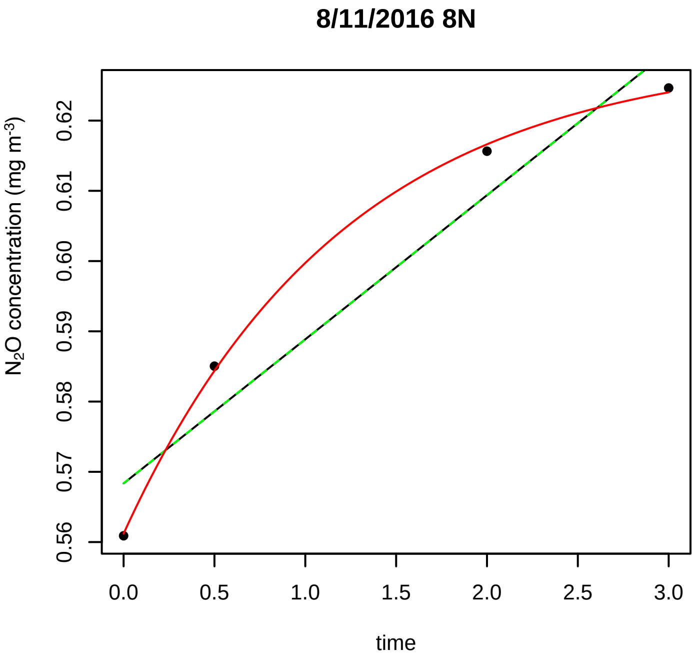
8/11/2016 8N
N2O concentration (mg m-3)
0.56
0.57
0.58
0.59
0.60
0.61
0.62
0.0
0.5
1.0
1.5
2.0
2.5
3.0
time

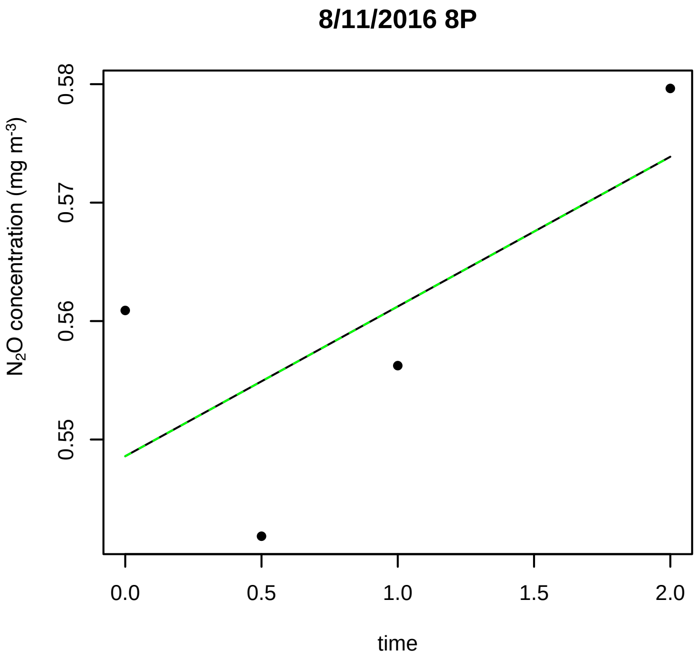
8/11/2016 8P
N2O concentration (mg m-3)
0.55
0.56
0.57
0.58
0.0
0.5
1.0
1.5
2.0
time

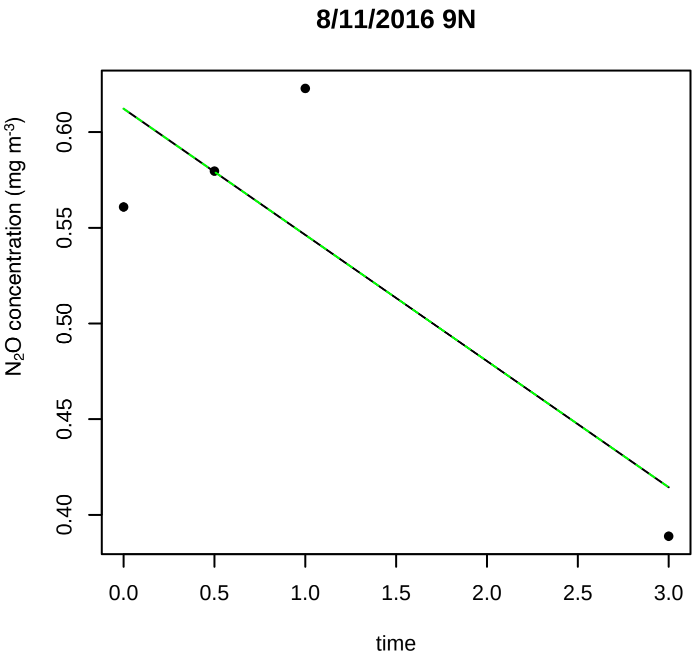
8/11/2016 9N
N2O concentration (mg m-3)
0.40
0.45
0.50
0.55
0.60
0.0
0.5
1.0
1.5
2.0
2.5
3.0
time

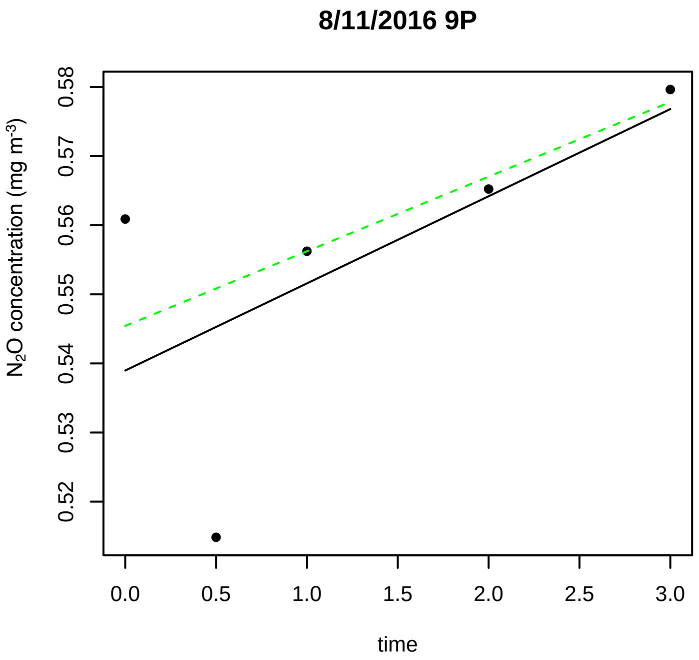

8/11/2016 9P
N2O concentration (mg m-3)
0.52
0.53
0.54
0.55
0.56
0.57
0.58
0.0
0.5
1.0
1.5
2.0
2.5
3.0
time
